# Dynamic chromatin organization and regulatory interactions in human endothelial cell differentiation

**DOI:** 10.1101/2022.04.15.488491

**Authors:** Kris G Alavattam, Katie A Mitzelfelt, Giancarlo Bonora, Paul A Fields, Xiulan Yang, Han Sheng Chiu, Lil Pabon, Alessandro Bertero, Nathan J Palpant, William S Noble, Charles E Murry

## Abstract

**Background:** Vascular endothelial cells are a mesoderm-derived lineage with many essential functions, including angiogenesis and coagulation. However, the gene regulatory mechanisms that underpin endothelial specialization are largely unknown, as are the roles of 3D chromatin organization in regulating endothelial cell transcription.

**Methods:** To investigate the relationships between 3D chromatin organization and gene expression in endothelial cell differentiation, we induced endothelial cell differentiation from human pluripotent stem cells and performed Hi-C and RNA-seq assays at specific timepoints in differentiation.

**Results:** Our analyses reveal that long-range intrachromosomal contacts increase over the course of endothelial cell differentiation, as do genomic compartment transitions between active and inactive states. These compartmental states are tightly associated with endothelial transcription. Dynamic topologically associating domain (TAD) boundaries strengthen and converge on an endothelial cell state, and nascent TAD boundaries are linked to the expression of genes that support endothelial cell specification. Relatedly, chromatin pairwise point interactions (DNA loops) increase in frequency during differentiation and are linked to the expression of genes with essential roles in vascular biology, including *MECOM, TFPI*, and *KDR*. To identify forms of regulation specific to endothelial cell differentiation, we compared the functional chromatin dynamics of endothelial cells with those of developing cardiomyocytes. Cardiomyocytes exhibit greater long-range *cis* interactions than endothelial cells, whereas endothelial cells have increased local intra-TAD interactions and much more abundant pairwise point interactions.

**Conclusions:** Genome topology changes dynamically during endothelial differentiation, including acquisition of long-range *cis* interactions and new TAD boundaries, interconversion of hetero- and euchromatin, and formation of DNA loops. These chromatin dynamics guide transcription in the development of endothelial cells and promote the divergence of endothelial cells from related cell types such as cardiomyocytes.

## Introduction

Endothelial cells, a mesoderm-derived cell population, line the entirety of the circulatory system. Their functions are complex and critical, including angiogenesis, blood clotting, barrier function, vasomotor function, and fluid/nutrient filtration. Endothelial cell dysfunction is a prominent feature of many pathological conditions, including nearly all cardiovascular diseases^1^. Complex transcriptional changes mediate both endothelial cell development and dysfunction^2,3^. It remains largely unknown what brings about such changes in gene expression—a knowledge gap that must be filled in order to, for example, develop means of regenerating damaged endothelium or modulating endothelial function.

The role of 3D chromatin organization in gene expression is an active area of study^4^. However, to date, few reports have examined the genome organization of endothelial cells^5–7^, and there are many gaps in our current understanding. We developed a novel stepwise protocol to induce endothelial differentiation from human pluripotent stem cells and, taking advantage of *in situ* DNase Hi-C^8^, we investigated global chromatin organization at specific time points in endothelial cell development. In performing RNA-seq, we correlated and contextualized Hi-C data with endothelial cell gene expression. Our results show that dynamic changes in chromatin organization, including genomic compartmentalization and topologically associating domains, are associated with corresponding changes in transcription. Our investigation of pairwise point interactions, which arise from chromatin looping mechanisms, reveals that point-interaction anchors are associated with the expression of essential genes in active (“A” type) chromatin compartments. Altogether, this study provides a comprehensive look at dynamic 3D chromatin organization during human endothelial cell development and uncovers important relationships between 3D chromatin organization and transcription.

## Methods

### Cell culture

Human pluripotent stem cells (hPSCs) from the RUES2 line (RUESe002-A; WiCell) were maintained on recombinant human Laminin-521 matrix (rhLaminin521; Biolamina) in Essential 8 (E8) media (ThermoFisher) at a density of 0.5 µg/cm^2^. Cells were passaged with Versene (ThermoFisher) and seeded overnight with 10 μM Y-27632 (ROCK inhibitor; Tocris). Karyotyping was performed by Diagnostic Cytogenetics Incorporated, Seattle, WA, and cells were found to contain no clonal abnormalities (Supplemental Figure 1A). Prior to directed endothelial cell differentiation (day −1), hPSCs were re-seeded with 3.0 × 10^5^ cells per well of a 12-well plate coated with 2 µg/cm^2^ rhLaminin521; then, the cells were immersed in E8 supplemented with 10 µM Y-27632. The following day (day 0), differentiation was induced with 7 µM CHIR99021 (a GSK3 inhibitor; Cayman) in RPMI (ThermoFisher) supplemented with 500 µg/mL BSA (Sigma, A9418) and 213 µg/mL ascorbic acid (Sigma, A8960). Seventy-two hours later (day 3), the media was switched to Stempro34 (Invitrogen) containing 300 ng/mL VEGF (Peprotech), 5 ng/mL bFGF (R&D), 10 ng/mL BMP4 (Preprotech), 4 × 10^−4^ M monothioglycerol (Sigma), 50 µg/mL ascorbic acid (Sigma), 2 mM L-glutamine (Invitrogen), and penicillin-streptomycin (Invitrogen). Forty-eight hours later (day 5), cells were passaged with 0.25% Trypsin (ThermoFisher) and re-seeded onto 0.2% gelatin-coated 10-cm dishes with 5.0 × 10^5^ cells per dish in Endothelial Cell Growth Medium (EGM; Lonza) containing 20 ng/mL VEGF, 20 ng/mL bFGF, and 1 µM CHIR99021. Cells were maintained on these dishes in EGM and above-described factors until day 14 (Figure 1A).

**Figure 1.**
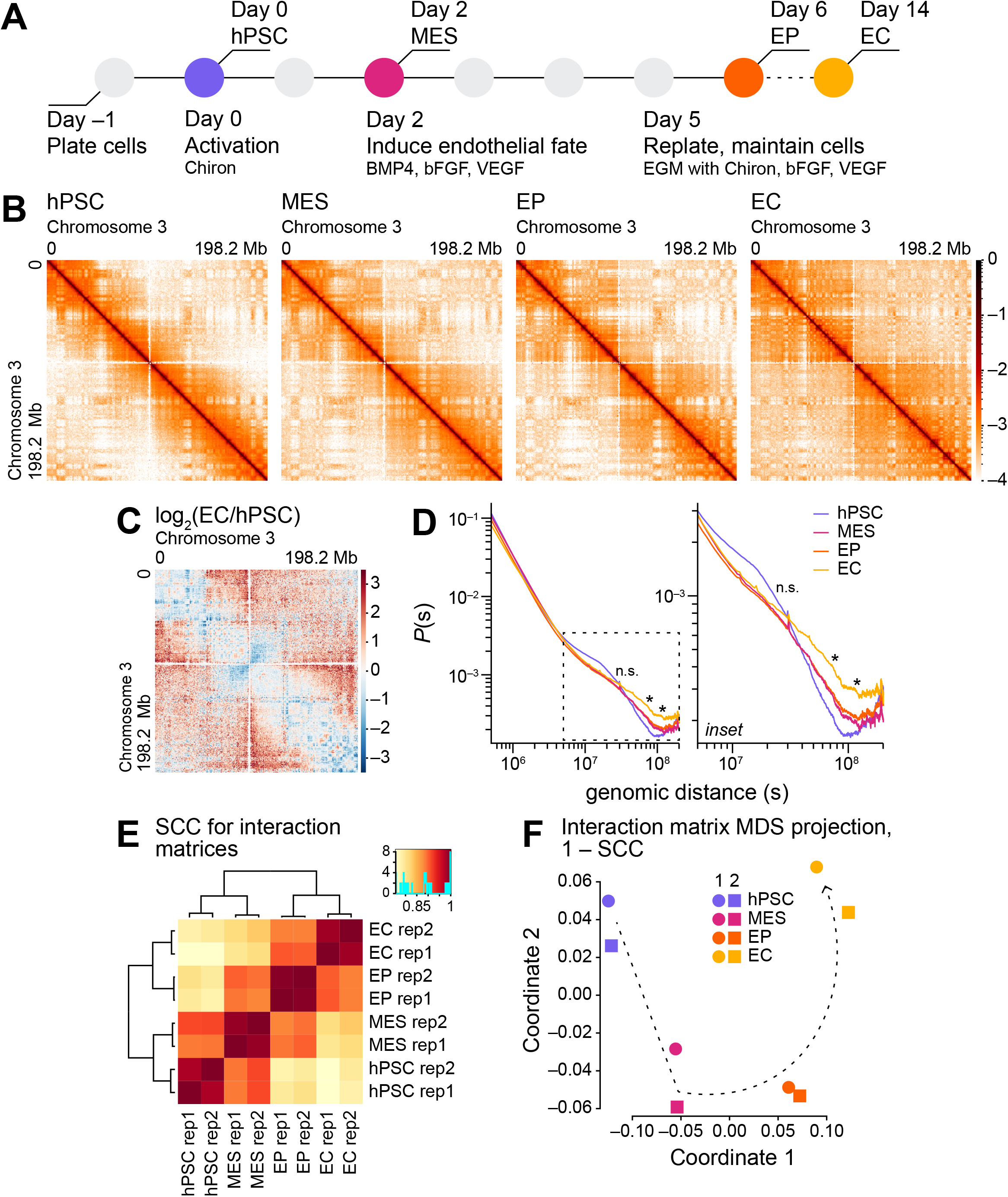
**A**. Schematic of the endothelial cell differentiation protocol. hPSC: human pluripotent stem cell; MES: mesoderm cell; EP: endothelial cell progenitor; EC: endothelial cell; BMP4: bone morphogenetic protein 4; bFGF: basic fibroblast growth factor; VEGF: vascular endothelial growth factor; EGM: endothelial cell growth medium. **B**. Heatmaps of normalized Hi-C interaction frequencies (500-kb resolution, chromosome 3) in hPSC, MES, EP, and EC. **C**. log_2_ ratios of normalized Hi-C interaction frequencies (500-kb resolution, chromosome 3) for successive cell types in endothelial cell differentiation. Red, observed/expected interactions higher in EC; blue, observed/expected interactions higher in hPSC. **D**. *Cis* interaction frequency probabilities *P* stratified by genomic distance *s* (Mb) over 0.5–120 Mb for Hi-C samples (500-kb resolution, autosomes; left). *P* stratified by s over 50–120 Mb (inset, right). P-value significantly different at *s* of 30, 75, and 105 Mb via t-tests between hPSC and EC with false discovery rate correction, * < 0.05, n.s. (not significantly different). **E**. Hierarchically clustered heatmap of HiCRep stratum-adjusted correlation coefficients (SCC)^14^ for Hi-C samples (500-kb resolution, autosomes). **F**. Multidimensional scaling (MDS) projection of SCC for Hi-C samples (500-kb resolution); similarity measure: 1 – SCC.

### Flow cytometry

To evaluate the purity of differentiating cells, cell aliquots were collected at days 6 and 14 of differentiation (Figure 1A). The cells were washed in Dulbecco’s phosphate-buffered saline (DPBS) with 5% fetal bovine serum (FBS) and resuspended in a solution of Dulbecco’s Modified Eagle’s Medium (DMEM; Corning) containing mouse anti-human CD34-PerCP (BD Biosciences 340430, 1:5) and mouse anti-human CD31-FITC (BD Biosciences 555445, 1:5) antibodies, or a solution of DMEM and antibody-appropriate isotype controls. All staining was performed on live cells. Staining was performed on ice for 45 minutes in darkness. The stained cells were washed in DPBS and fixed with 4% paraformaldehyde (Affymetrix) prior to flow cytometry analyses. Flow cytometry was performed using a FACS Canto II cell analysis instrument (BD Biosciences); flow cytometry data were analyzed using FlowJo Software (FlowJo, LLC). Gates were set such that isotype controls contained 5% positive cells (Supplemental Figure 1B, 1C).

### Hi-C: Sample preparation, wetwork, library generation, and sequencing

*In situ* DNase Hi-C [12, 31] was performed on 2−3 × 10^6^ cells from two independent differentiations at the following time points: day 0, a pluripotent cell type (hPSC); day 2, a mesodermal cell type (MES); day 6, an endothelial-progenitor cell type (EP); and day 14, an endothelial cell type (EC; Figure 1A). To prepare samples for Hi-C wetwork, plated cells were washed three times with DPBS and then fixed with a mixture of fresh RPMI containing 2% formaldehyde (diluted from a 37% formaldehyde solution); fixation took place for 10 minutes at room temperature with orbital rotation. Formaldehyde was quenched with 1% 2.5 M glycine for 5 minutes at room temperature and then 15 minutes at 4 °C. Afterwards, cells were treated with 0.25% trypsin for 5 minutes at 37 °C, washed with RPMI containing 10% FBS, and subsequently scraped off their plates. Cells were washed once with DPBS, flash frozen in liquid nitrogen, and stored at -80 °C until the time of Hi-C wetwork.

To perform *in situ* DNase Hi-C, frozen samples were thawed and lysed in 500 μL lysis buffer comprised of 10 mM Tris-HCl (pH 8.0), 10 mM NaCl, 0.2% Igepal CA-630, 1× protease inhibitor, and double-distilled water (ddH_2_O). Then, nuclei were resuspended in 300 µL DNase buffer with 0.2% SDS and MnCl_2_, and incubated at 37 °C for 60 minutes with periodic vortexing. Afterwards, nuclei were subjected to an additional 300 µL DNase buffer containing 2% Triton X-100 and RNase A, and incubated for another 10 minutes. Six units of DNase (ThermoFisher, EN0525) were added and incubated for 7 minutes at room temperature. The reaction was stopped with 30 µL of 0.5 M EDTA and 15 µL of 10% SDS. Nuclei were collected and resuspended in 150 µL water and combined with 300 μL AMPure XP beads (Beckman). DNA-end repair with T4 DNA Polymerase (ThermoFisher, EP0062) and Klenow (ThermoFisher, EP0052) was performed *in situ*, as was subsequent dA-tailing with Klenow exo- (ThermoFisher, EP0422). Then, biotinylated oligonucleotides (adapters) were ligated to DNA at 16 °C overnight. To remove unbound adapters, nuclei were washed once with AMPure buffer (20% PEG in 2.5 M NaCl), then twice with 80% ethanol. To carry out adapter phosphorylation and ligation, PNK treatment was performed for 4 hours at room temperature. To de-crosslink the DNA, samples were treated with Proteinase K overnight at 62 °C. The next day, DNA precipitation was performed with 0.055 mg/mL glycogen, 10% volume 3 M NaOAc (pH 5.2), and 100% volume isopropanol for 2 hours at -80 °C. To purify the DNA, it was resuspended in 100 μL water and combined with 100 µL of AMPure beads. The pull-down of biotin adapter-containing DNA was performed using MyOne C1 Beads (ThermoFisher, 65001) for 30 minutes at room temperature with rotation. Afterwards, samples were washed four times with bind-and-wash buffer (5 mM Tris-HCl pH 8.0, 0.5 mM EDTA, 1 M NaCl, and 0.05% Tween-20) followed by two elution-buffer washes. On-bead DNA underwent end repair using the reagents in a Fast DNA End Repair Kit (ThermoFisher, K0771), and this was followed by dA-tailing using Klenow Exo- (ThermoFisher, EP0422); between each reaction, the DNA was washed four times with bind-and-wash buffer and twice with Tris-EDTA buffer.

Sequencing Y-adapters were ligated at room temperature for 60 minutes. To amplify the Hi-C libraries, the DNA underwent 10 PCR cycles using Kapa HiFi ReadyStart Master Mix (Roche, KK2602) with barcode-containing primers. Libraries were purified with 0.8× volumes of Ampure XP beads and quantified with a Qubit prior to sequencing. The libraries were paired-end sequenced using a NextSeq 500 (Illumina) in a high-output run with 150 cycles, 75 cycles for each end.

### RNA-seq: Sample preparation, wetwork, library generation, and sequencing

Cell samples from two independent differentiations were collected in Buffer RLT (QIAGEN) at the the time points described above (Figure 1A). Samples were stored at −80 °C prior to RNA purification, which was performed with an RNeasy Mini Kit (QIAGEN) with on-column DNase digestion. RNA-seq libraries were prepared from total RNA (≥200 nucleotides in length) using the TruSeq Stranded Total RNA Ribo-Zero H/M/R kit (Illumina, RS-122-2201). Libraries were paired-end sequenced on a NextSeq 500 (Illumina) in a high-output run with 150 cycles, 75 cycles for each end. The RNA-seq datasets exhibit high percentages of unique, paired alignments (Supplemental Table 1).

### Hi-C: Data-sourcing, alignment, processing, and quality control

*In situ* DNase Hi-C datasets for cardiomyocyte samples differentiated from RUES2 hPSCs were obtained from published work (GEO GSE106690)^9^. As with the endothelial cell data generated for this study, the cardiomyocyte data are comprised of samples from two independent differentiations at the the following time points: day 0, hPSC; day 2, MES; day 5, a cardiomyocyte-progenitor cell type (CP); and day 14, a cardiomyocyte cell type (CM).

Reads were aligned to a *Homo sapiens* reference genome (Ensembl 83) with BWA-MEM (0.7.13-r1126)^10^’^11^ using default parameters. Each read-pair end was aligned individually. The Hi-C datasets exhibit high percentages of unique, paired alignments (Supplemental Table 2, 3). Primary alignments were extracted and sorted with Samtools (version 1.2)^12^. Then, the alignments were processed with HiC-Pro (version 2.7.6)^13^, filtering for MAPQ scores ≥30 and excluding read pairs that mapped within 1 kb of each other; PCR duplicates, defined as sequence matches with the exact same starts and ends, were excluded. HiC-Pro allValidPairs files and unbalanced matrices were generated at the following resolutions: 40, 100, and 500 kb. To assess contact-matrix similarity, HiCRep was used (Supplemental Table 4)^14^. HiCRep stratum-adjusted correlation coefficients (SCCs) were calculated using the following parameters: resol = 500000, ubr = 5000000, h = 1. All other parameters were set to default values.

To facilitate genomic binning at resolutions finer than 40 kb, biological replicates were merged with HiC-Pro, thereby increasing the sequencing depth for each sample. The merged Hi-C datasets are comprised of 111.4 million unique, valid read pairs for hPSC from endothelial cell differentiation; 120.0 million unique, valid read pairs for MES from endothelial cell differentiation; 128.7 million unique, valid read pairs for EP; 132.9 million unique, valid read pairs for EC; 138.5 million unique, valid read pairs for hPSC from cardiomyocyte differentiation; 143.3 million unique, valid read pairs for MES from cardiomyocyte differentiation; 161.6 million unique, valid read pairs for CP; and 185.0 million unique, valid read pairs for CM. HiC-Pro allValidPairs files and unbalanced text matrices were generated for pooled replicates at the following resolutions: 10, 20, 40, 100, and 500 kb.

Using cooler^15^ and `HiCExplorer`^16–18^ hicConvertFormat, HiC-Pro matrices were converted to the cooler format (`.cool`). Read pairs not aligned to autosomes or chromosome X were excluded from `.cool` files with HiCExplorer `hicAdjustMatrix`. Cooler-formatted matrices were balanced using Sinkhorn balancing^19^ such that the sum of every row and column is equal; to do so, the cooler balance command was called with default parameters.

### Hi-C: Generation and visualization of Hi-C heatmaps

To generate and visualize chromatin-contact heatmaps, HiCExplorer `hicPiotMatrix` was used with contact matrices. To aid visualization, matrix values were log-transformed. To generate and visualize differential interactions between samples, log_2_-ratio matrices were generated with HiCExplorer `hicCompareMatrices` using `.cool` files from two datasets; the log_2_-ratio matrices were plotted with `hicPiotMatrix.` To generate and visualize Pearson correlation coefficient heatmaps, contact matrices were input into HiCExplorer hicPCA for conversion to distance-normalized matrices^20^ (i.e., matrices taken from dividing observed interactions by expected interactions) and then Pearson correlation coefficient matrices; `hicPiotMatrix` was used to visualize the Pearson correlation coefficient matrices.

### Hi-C: *Cis* contact-decay curve analyses

*Cis* contact-decay curves were generated by aggregating normalized counts as a function of distance at 500-kb intervals using HiCExplorer `hicPiotDistVsCounts.` Adjusted p-values were obtained from t-tests between hPSC and EC.

### Hi-C: Genomic compartment analyses

To calculate principal component (PC) scores for genomic compartment assignment, contact matrices were distance-normalized, transformed into Pearson correlation coefficient matrices, and eigen-decomposed using HOMER^21,22^. HiC-Pro `validPairs` files were used as inputs. Eigenvectors were generated for contact matrices at 100- and 500-kb resolutions. Eigenvectors were assessed, and the first PC (PC1) was found to represent the genomic compartment profile (data not shown); subsequent principal components represented profiles distinct from genomic compartments (data not shown). Per convention^20^, PC1 orientation and binwise genomic-compartment assignments were based on biological features: bins with higher gene densities and increased enrichment for transcription were assigned to the “A” compartment type; all other bins were assigned to the “B” compartment type. Using oriented PC1 scores, sample similarity was evaluated via Spearman correlation coefficient and MDS analyses (Supplemental Figure 3B, C).

Genomic compartment transitions were defined using the following criteria: (i) for a given bin, PC1 scores are available for each sample (i.e., no sample is represented by NA); (ii) for a given bin, at least one sample has a mean PC1 score greater than 0 and at least one sample has a mean PC1 score less than 0. A–B–A–B and B–A–B–A transitions represented less than 1% of genomic compartment-switch regions and were combined, respectively, with A–B- and B–A-transitioning regions for downstream analyses, including enrichment calculations for differentially expressed genes (DEGs) with respect to stable and dynamic genomic compartments.

Saddle plots were generated by assigning each bin to its corresponding percentile value and dividing the genome into deciles. Each interaction was normalized to the average score at the corresponding distance for *cis* interactions and assigned to a decile pair based on the two bins. The plot represents the log_2_ average value for pairs of deciles. Changes between one sample and another are represented by log_2_ values of differences.

### Hi-C: Analyses of topologically associating domains

Using the insulation score method^23^, topologically associating domains (TADs) were called for Hi-C at 40-kb resolution. To do so, cworld (github.com/dekkerlab/cworld-dekker) `matrix2insuiation.pl` was called with the following parameters: `--is 520001 --ids 320001 --ss I60001 --nt 0.01 --im mean.` Insulation scores are defined as the average number of chromatin contacts across a given bin, and TAD boundaries are called at local insulation score minima, which represent areas where the average number of chromatin contacts across a given bin are few. A given boundary was categorized as “shared” if it was observed at the same location ±40 kb (i.e., within up to one bin of the boundary) in another sample; if boundaries did not meet these conditions, then they were categorized as “sample-specific.”

Aggregate TAD heatmaps were generated using FAN-C^24^ `fane aggregate` with the following parameters: `--tads-imakaev —vmin 0.02 --vmax 0.075.`

### Hi-C: Analyses of pairwise point interactions

Pairwise point interactions (PPIs) were called at 10-, 20-, and 40-kb resolutions using HiCCUPS^7^, which was invoked with the following parameters: `-m 512 -r 10000,20000,40000 -k KR -f, 1,.1,.1 -p 4,2,1 -i 8,4,2 -t 0.02,1.5,1.75,2 -d 20000,40000,80000 --cpu.` UpSet plots were generated to score the overlap of loop anchors among samples^25^’^26^.

All calculations, including the proportions of anchors (called at 10-kb resolution) in A and B compartments (called at 100-kb resolution) and the proportions of loop anchors that overlap up- and downregulated DEGs in A and B compartments, were calculated using custom R scripts that will be made available upon publication.

### RNA-seq: Sourcing, alignment, and gene-level quantification of alignments

Reads were mapped to hg38 using HISAT2 (version 2.1.0)^27^ with default settings. Files in`.sam` format were converted into coordinate-sorted `.bam` files using sambamba (version 0.6.6)^28^ with default parameters. The program featureCounts from the Subread package (version 1.6.3)^29^ was used to quantify gene expression levels; the following parameters were specified: `-p -B -a gtf_file -t exon -g gene_id.`

### RNA-seq: Differential gene expression analysis

Differential gene expression analysis was performed with DESeq2 (version 1.32.0)^30^. Genes exhibiting a false discovery rate (FDR)-controlled p-value (adjusted p-value) < 0.05 and an absolute log_2_ fold change > 1 were categorized as DEGs.

### RNA-seq: Gene ontology analyses

Gene ontology term enrichment analyses were performed using the ToppGene Suite ToppFun application^31^; default settings were used, and the full gene set for each category was used as the background set. P-values were obtained from hypergeometric tests and adjusted via Bonferroni corrections.

### RNA-seq: Enrichment of differentially expressed genes with respect to genomic compartments

To calculate enrichment scores for DEGs with respect to compartment-transition types (e.g., A–B, B–A, etc.), we first calculated the ratio of DEGs within a given compartment-transition types to all genes within the compartment-transition type; then, this value was divided by the ratio all DEGs to all genes; i.e.,

~~~
enrichment = (DEGs in compartment type / genes in compartment type) / (all DEGs
/ all genes)
~~~

P-values were obtained from chi-squared tests with Yates correction.

### RNA-seq: Enrichment of differentially expressed genes with respect to topologically associating domain boundaries

To calculate enrichment scores for DEGs with respect to gained (EC-specific), lost (hPSC-specific) and shared TAD boundaries, we calculated the ratio of DEGs within ±40 kb (i.e., ±1 bin) of a given boundary type to all genes within the boundary type; then, this value was divided by the ratio all DEGs to all genes; i.e.,

~~~
enrichment = (DEGs in vicinity of boundary type / genes in vicinity of boundary
type) / (all DEGs / all genes)
~~~

P-values were obtained from chi-squared tests with Yates correction.

### RNA-seq: Enrichment of differentially expressed genes with respect to pairwise point interactions

We used the following equation to calculate enrichment scores for genes that overlap PPI anchors:

~~~
gene enrichment = observed genes overlapping anchors / expected genes
overlapping anchors = (DEGs overlapping anchors / DEGs) / (genes overlapping
anchors / genes)
~~~

We used the following equation to calculate enrichment scores for anchors that overlap genes:

~~~
anchor enrichment = observed anchors in genes / expected anchors in genes =
anchors overlapping DEGs / [anchors x (DEGs / genes)]
~~~

P-values were obtained from chi-squared tests with Yates correction.

### Statistical Analyses

No statistical methods were used to predetermine sample sizes. No data were excluded from analyses. The experiments were not randomized and investigators were not blinded to allocation during experiments and outcome assessment. Where appropriate, statistical tests are described above, in the Results, in Figure Legends, and in Supplemental Figure Legends.

### Figure Preparation

Plots were generated with, alone or in combination, Excel (version 16.60, Microsoft), base R (version 4.1), the R software package ggplot2 (version 3.3.4), and various plotting programs employed by the other software packages used in this study. Illustrator (version 26.0.2, Adobe) was used for composing figures.

### Code Availability

Source code for analyses performed in this study, with documentation, examples, and additional information, will be made available upon publication.

### Data Availability

Sequencing data generated for this study will be deposited in the 4D Nucleome (4DN) Data Portal and made available upon publication (the authors can supply links to the data after being uploaded to the 4DN Data Portal). Published sequencing data have been deposited in the National Center for Biotechnology Information (NCBI) Gene Expression Omnibus (GEO) under the accession numbers GSE106690^9^.

## Results

### Long-range *cis* contacts increase during endothelial cell differentiation

To study the dynamics and functional significance of 3D chromatin organization in endothelial cell development, we developed a novel stepwise protocol to induce endothelial differentiation from hPSCs (see Methods). Using the female RUES2 embryonic stem cell line (Supplemental Figure 1A), we sought to recapitulate key signaling events in endothelial cell development. First, we induced the mesodermal lineage through activation of the WNT signaling pathway; then, we directed the cells to an endothelial cell fate through the addition of bone morphogenic protein 4 (BMP4), basic fibroblast growth factor (bFGF), and vascular endothelial growth factor (VEGF; Figure 1A). We took samples across a time course of differentiation (Figure 1A): day 0, a pluripotent cell type (hPSC); day 2, a mesodermal cell type (MES); day 6, an endothelial-progenitor cell type (EP); and day 14, an endothelial cell type (EC). We obtained high-purity cell populations as determined by flow cytometry with antibodies raised against CD31 and CD34, two endothelial cell markers: EP were >75% pure and EC were >90% pure (Supplemental Figure 1B, C).

To assess the quality of samples resulting from the endothelial cell differentiation protocol, we prepared and analyzed bulk RNA-seq datasets from two independent differentiations for each time point. Alignment of the RNA-seq datasets to the genome resulted in high percentages of uniquely mapped reads (78–90%; Supplemental Table 1). To evaluate relationships among the datasets, we performed principal component analysis (PCA). PCA revealed the tight clustering of biological replicates (Supplemental Figure 2A). The first principal component accounted for 29% of the variance and recapitulated an endothelial cell developmental trajectory (Supplemental Figure 2A). The second principal component accounted for 15.6% of the variance and separated mesenchymal (MES) and epithelial (hPSC, EP, and EC) cell types—evidence of the developmental epithelial-to-mesenchymal transition and subsequent mesenchymal-to-epithelial transition (Supplemental Figure 2A). On a broader scale, we found that transcriptomes underwent overt changes during specification (Supplemental Figure 2B). Taken together, these results support the efficacy and reproducibility of our endothelial cell differentiation protocol.

To examine chromatin organization changes over the course of endothelial cell differentiation, we prepared and analyzed *in situ* DNase Hi-C^8^ datasets from two independent differentiations for each time point. We observed the alignment of Hi-C reads at high rates (68–74%; Supplemental Table 2, 3) and comparable levels of *cis* interactions across differentiation (64–80%; Supplemental Table 2, 3). To assess the experimental reproducibility of the Hi-C datasets, we performed HiCRep analyses^14^. Stratum-adjusted correlation coefficients (SCC) output by HiCRep revealed high levels of reproducibility between biological replicates (Supplemental Table 4). These results indicate that the Hi-C data are of consistent and high quality. Thus, for subsequent analyses of chromatin organization, we pooled replicates to increase the sequencing depth—and thus the resolution of genomic bins—for each cell type.

Next, we surveyed features of chromatin organization across endothelial cell differentiation. In all four time points, chromosome-wide contact maps displayed patterns indicative of local and longer-range chromatin interactions (Figure 1B). These included an abundance of “near” *cis* interactions in hPSCs (e.g., strong interactions along the diagonal of the hPSC panel in Figure 1B, C) that, over the course of differentiation, spread outward (Figure 1B, C), indicating an increase in the numbers of “far” *cis* interactions. Next, we examined the *cis* chromatin contact probability *P(s)* for pairs of genomic loci stratified by genomic distance s, which may be indicative of the general polymer state of chromatin^20,32^. Consistent with our analyses of *cis* contact maps, we found that the proportion of long-range contacts increased over differentiation: hPSC had the lowest proportion of long-range contacts >30 Mb; MES and EP had similar, higher proportions of long-range contacts >30 Mb; and EC exhibited the highest proportion contacts >30 Mb (Figure 1D). These data reveal that long-range *cis* contacts increase over the course of endothelial cell differentiation.

To assess the influence of *cis* chromatin interactions as differentiation progresses, we performed hierarchical clustering of the SCC scores produced by HiCRep for *cis* interactions; clustering paired replicates while separating the time-course datasets into two general groups: an “early” group, composed of hPSC and MES, and a “later” group, composed of EP and EC (Figure 1E). Next, we performed multidimensional scaling (MDS) of *cis* interaction maps using 1 – SCC as a measure of similarity. MDS paired replicates and arranged the samples in order of time point, revealing an apparent endothelial cell differentiation trajectory (Figure 1F). Together, these results indicate that changes in chromatin organization—including a general, gross increase in long-range *cis* chromatin contacts—are a key feature of endothelial cell differentiation, separating and ordering datasets by time point.

### Dynamic compartmentalization reflects endothelial cell-specific changes in gene expression

Given these findings, we sought to understand how chromatin organization changes across differentiation—and the functional significance of such changes. To address these points, we investigated three forms of chromatin organization: genomic compartments, topologically associating domains, and peaks of elevated contact frequency referred to as “pairwise point interactions.” Pairwise point interactions are thought to represent DNA “loops;” however, we avoid using the term “loop” for its multiple meanings and interpretations, as described in a recent review of chromosome organization^33^.

To begin, we focused on genomic compartments, the “plaid” patterns of chromatin interactions evident in Hi-C heat maps^20,34,35^. Genomic compartments represent at least two alternating states of chromatin, A and B, and each state preferentially interacts with other loci of the same state. A compartments are associated with higher gene expression in active chromatin (euchromatin), while B compartments are associated with gene silencing in inactive chromatin (heterochromatin). To segregate genomic bins (500-kb resolution) into A/B compartments, we computed the first principal components (PC1s) of the Pearson correlation-transformed contact matrices. We identified genomic compartments in all samples taken across endothelial cell specification (Figure 2A). To assess changes in compartmentalization as endothelial cells differentiate, we analyzed the proportions of stable and dynamic compartments. We found that approximately 25% of compartments (500-kb resolution) are dynamic, undergoing one or more compartment switches across time points (Figure 2B, C; Supplemental Figure 3A). Hierarchical clustering of Spearman correlation coefficients (ρ) for PC1 scores paired and ordered replicates (Supplemental Figure 3B); likewise, MDS paired replicates while revealing a differentiation trajectory in the separation of time points from each other (Supplemental Figure 3C). Taking these observation together, a substantial number of compartment transitions as endothelial cell specialization progresses and, as with the progressive increase in *cis* chromatin contacts (Figure 1B–D), the dynamics of genome compartmentalization are another key feature of endothelial cell specification.

**Figure 2.**
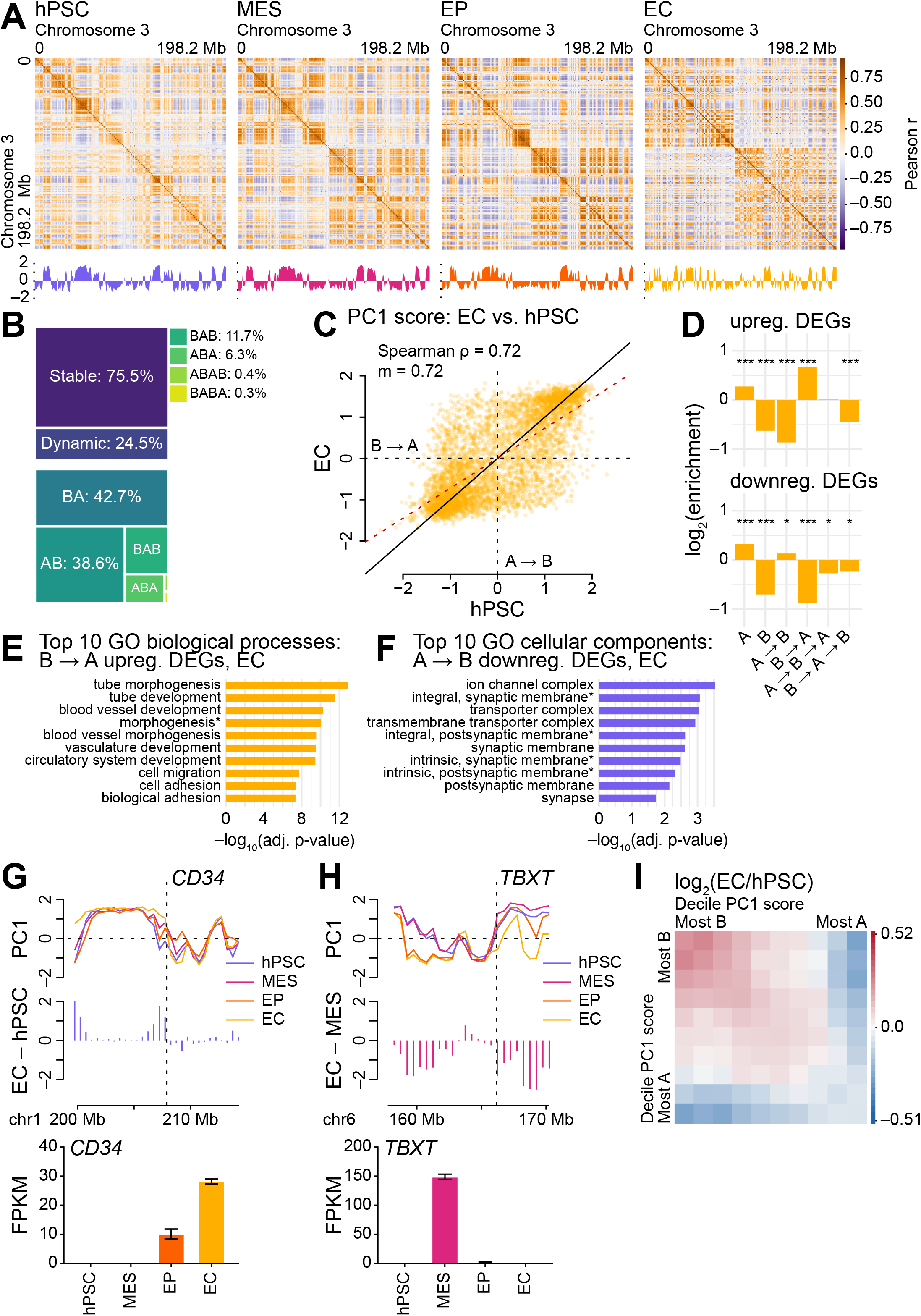
**A**. Pearson correlation coefficient matrices for normalized Hi-C interaction frequencies (500-kb resolution, chromosome 3) for samples from endothelial cell differentiation. Below the matrices, principal component 1 (PC1) from HOMER-implemented principal component analysis (see Methods)^21,22^. **B**. Treemap showing the proportions of stable and dynamic genomic compartments over endothelial cell differentiation (top). Treemap showing proportions of types of compartment switches (bottom). **C**. Scatter plot comparing PC1 scores from hPSC and EC Hi-C samples (500-kb resolution, autosomes). ρ, Spearman correlation coefficient; m, regression slope; red dashed line, regression line; black solid line, x = y; A → B, A-to-B compartment switches; B → A, B-to-A compartment switches. **D**. Bar charts depicting log_2_ enrichment (see Methods) of upregulated (top) and downregulated (bottom) differentially expressed genes (DEGs) with respect to stable and dynamic genomic compartments (500-kb resolution, autosomes). log_2_ values are observed/gene density. DEGs were called via DESeq2 analysis of EC versus hPSC (adjusted p-value < 0.05, absolute log_2_ fold change > 1; see Methods). Enrichment significantly different via chi-squared tests with Yates correction, * < 0.05, ** < 0.01, *** < 0.001. **E, F**. Bar chart depicting adjusted p-values for the top 10 gene ontology (GO) biological process or cellular component terms for genes located within bins (500-kb resolution, autosomes) that undergo (**E**) B-to-A and (**F**) A-to-B compartment transitions in endothelial cell specification. GO biological process terms significantly different from hypergeometric tests with Bonferroni corrections. Terms with asterisks have been abbreviated: (**E**) “morphogenesis” is “anatomical structure formation involved in morphogenesis,” “integral, synaptic membrane” is “integral component of synaptic membrane;” (**F**) “integral, postsynaptic membrane” is “integral component of postsynaptic membrane,” “intrinsic, synaptic membrane” is “intrinsic component of synaptic membrane,” and “intrinsic, postsynaptic membrane” is “intrinsic component of postsynaptic membrane.” **G, H**. Line plots for PC1 scores at and within the vicinity of (**G**) *CD34* (chr1: 207.88–207.91 Mb) and (**H**) *TBXT* (chr6: 166.16–166.17 Mb) for Hi-C samples (500-kb resolution; top). Bar plots indicating change in PC1 scores via the subtraction of (**G**) hPSC and (**H**) MES PC1 scores from EC PC1 scores (middle). Bar plots for the RNA-seq expression levels of (**G**) *CD34* and (**H**) *TBXT*across endothelial cell differentiation (bottom). Expression levels: FPKM; error bars: standard error of the mean (SEM). **I**. log_2_ ratios of “saddle plots,” i.e., 10 × 10 decile-binned matrices quantifying the strength of PC1 scores, for EC and hPSC Hi-C samples (500-kb resolution, autosomes); red, observed/expected interactions higher in EC; blue, observed/expected interactions higher in hPSC.

Next, we focused on the function of dynamic compartmentalization in endothelial cell differentiation. In a recent study of differentiating cardiomyocytes—which share their developmental origin, cardiogenic mesoderm^36^, with endothelial cells—*we* found that compartment switches coincide with cell type-specific transcriptional regulation^9^. Thus, we sought to understand the influence of genomic compartmentalization on the dynamic transcriptomes of endothelial cell differentiation. We calculated the enrichment of differentially expressed genes (EC versus hPSC) with respect to stable and dynamic compartments (Figure 2D). Regions that undergo B-to-A transitions are enriched for differentially expressed genes (DEGs) upregulated in EC, and these are associated with numerous functions in endothelial cell specification (Figure 2E; Supplemental Table 5), including blood vessel development, circulatory system development, cell migration, and cell adhesion, among other related Gene Ontology terms^37^’^38^. Compartment changes are also associated with gene repression; e.g., in regions that undergo A-to-B transitions, DEGs downregulated in EC are associated with neuronal development and function, regions that become “locked down” during endothelial differentiation (Figure 2F; Supplemental Table 6). Consistent with the enrichment of endothelial cell development genes in B-to-A regions, the endothelial cell marker *CD34* is subject to just such a transition prior to its elevated expression in EP and EC (Figure 2G). Similarly, mesodermal markers such as *TBXT* are associated with an A-to-B transition, coincident with downregulation in the EP stage (Figure 2H). Thus, as with differentiating cardiomyocytes^9^’^39^, compartment transitions in differentiating endothelial cells are associated with essential changes in transcription, indicating that dynamic genomic compartmentalization is an important regulator of cell type-specific transcription.

Given that, in the course of specification, long-range *cis* interactions increase (Figure 1B–D) and 25% of the genome undergoes compartment transitions (Figure 2B, C; Supplemental Figure 3A), we sought to examine large-scale changes in genomic compartment strength across endothelial cell differentiation. We generated “saddle plots,” which quantify the strength of compartment segregation, for the hPSC and EC time points. We noted that stronger *cis* contacts occur between homotypic regions (i.e., A–A or B–B compartment interactions) in comparison to heterotypic regions (i.e., A-B compartment interactions; Figure 2I; Supplemental Figure 3D). In the course of endothelial cell differentiation, the strength of *cis* interactions between B compartments increased while the strength between A compartments decreased (Figure 2I; Supplemental Figure 3D). As seen in the differentiation of other tissues^9,40^, these findings show that dynamic, strengthening B compartments repress transcription in endothelial cell differentiation.

### Increasingly strengthened topologically associating domains support endothelial cell-specific gene expression

Next, we shifted focus from megabase-scale genomic compartmentalization to a form of chromatin organization that exists on the kilobase scale: topologically associating domains (TADs). TADs comprise local “neighborhoods” of increased chromatin contact frequency and are often delimited by insulator sequences^41,42^. Because TADs are thought to regulate gene expression^42–45^, we sought to evaluate TAD features across endothelial cell differentiation. Using the insulation score approach to calling TAD boundaries^23^ (see Methods), we identified TADs in all samples taken across endothelial cell differentiation (Supplemental Figure 4A).

Hierarchical clustering of Spearman correlations for TAD insulation scores gave similar results to hierarchical clustering of separate forms of chromatin organization (Figures 1E, Supplemental Figure 3B). Replicates were paired, and the time course datasets were separated into a group of “early” samples, hPSC and MES, and a group of “later” samples, EP and EC, indicating an overall change in insulation scores across differentiation (Figure 3A). We observed a similar dynamism in our analyses of TAD boundaries: approximately 50% of TAD boundaries change during endothelial differentiation (i.e., TAD boundaries change greater than ±40 kb—one bin—in position; Figure 3B). Clustering of the intersections revealed the same branching of early (hPSC and MES) and later (EP and EC) cell types (Figure 3B). Of note, EP and EC share the highest proportion of TAD boundaries: EP shares 77.7% of its boundaries with EC, and EC shares 78.1% of its boundaries with EP (Supplemental Table 7). To evaluate the dynamics of TAD boundaries across differentiation, we sought to understand how insulation scores change at TAD boundaries. Although scatter plots revealed limited changes in insulation scores at TAD boundaries (Supplemental Figure 4B), the differences in insulation to the immediate left and right of boundaries—so-called “TAD boundary strengths”—increased in the course of differentiation (Supplemental Figure 4C; see Methods^23,46^). This is evident in aggregate analyses of TADs across the timepoint datasets (Figure 3C), which reveal that *cis* chromatin interactions are increasingly restricted within TAD boundaries; the restriction effect is far stronger in EC compared to hPSC (Supplemental Figure 4D). Taken together, these results suggest that, although approximately half of all TAD boundaries are conserved in differentiation, dynamic boundaries converge on an EC state; concomitant with this progression is a steady and marked increase in the numbers of intra-TAD chromatin contacts as differentiating cells mature.

**Figure 3.**
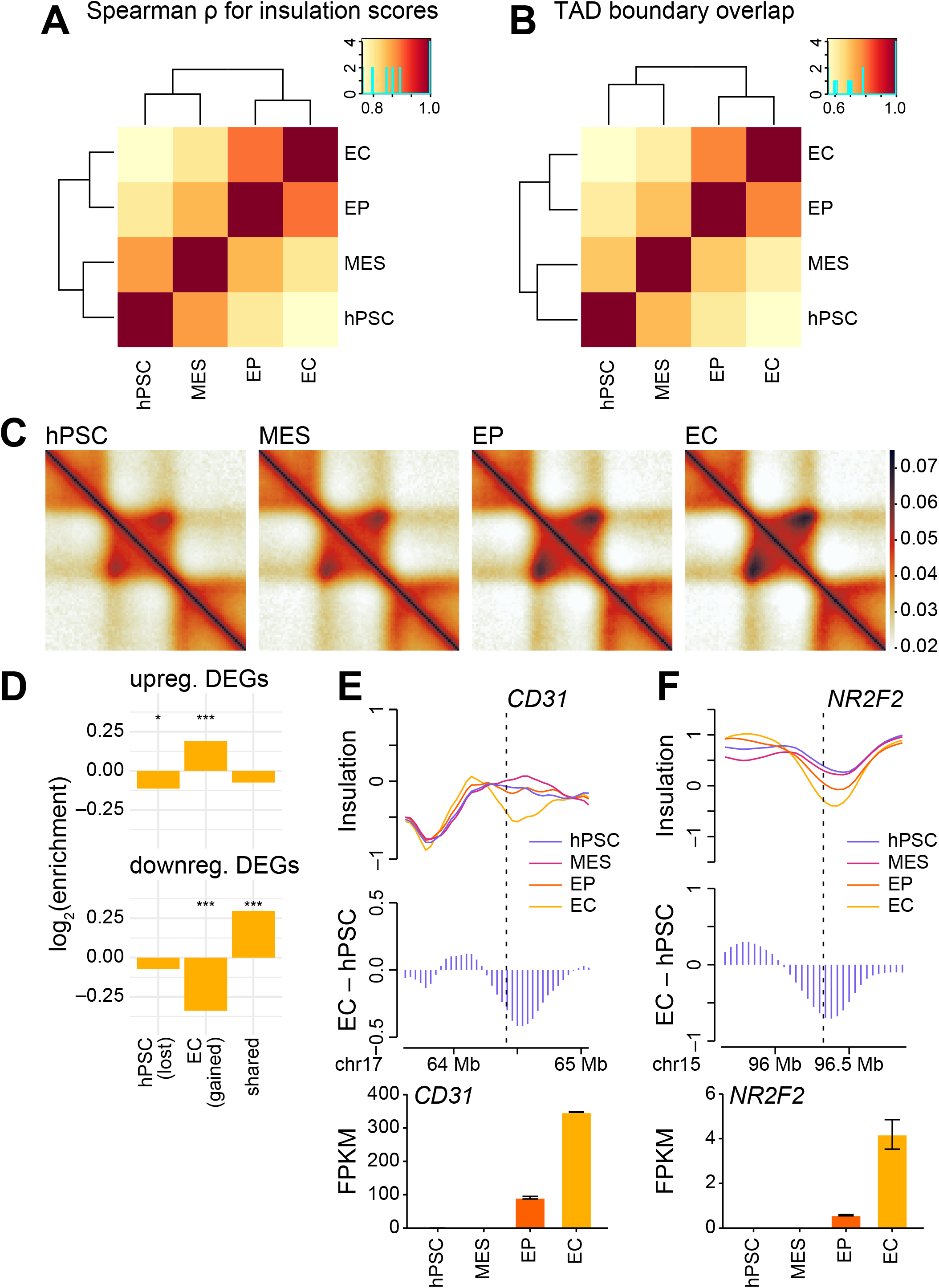
**A**. Hierarchically clustered heatmap of Spearman correlation coefficients (ρ) for insulation scores from Hi-C samples (40-kb resolution, autosomes) taken across endothelial cell differentiation. **B**. Hierarchically clustered heatmap for the TAD-boundary set intersections of Hi-C samples (40-kb resolution, autosomes). **C**. Aggregate heatmaps representing an average *cis* chromatin contact conformation around TADs for Hi-C samples (40-kb resolution, autosomes). **D**. Bar charts depicting log_2_ enrichment (see Methods) of upregulated (top) and downregulated (bottom) differentially expressed genes (DEGs) with respect to TAD boundaries (40-kb resolution, autosomes) that are EC-specific (gained in differentiation), hPSC-specific (lost in differentiation), and shared between hPSC and EC (maintained in differentiation). log_2_ values are observed/gene density. DEGs were called via DESeq2 analysis of EC versus hPSC (adjusted p-value < 0.05, absolute log_2_ fold change > 1; see Methods). Enrichment significantly different via chi-squared tests with Yates correction for set-wise overlaps, * < 0.05, ** < 0.01, *** < 0.001. **E, F**. Line plots for insulation scores at and within the vicinity of (**E**) *CD31* (chr17: 64.32–64.41 Mb) and (**F**) *NR2F2* (chr15: 96.33–96.34 Mb) for Hi-C samples (40-kb resolution; top). Bar plots indicating changes in insulation score via the subtraction of hPSC insulation scores from EC insulation scores (middle). Bar plots for the RNA-seq expression levels of (**E**) *CD31* and (**F**) *NR2F2* across endothelial cell differentiation (bottom). Expression levels: FPKM; error bars: standard error of the mean (SEM).

We hypothesized that nascent boundaries and increased intra-TAD contacts facilitate gene expression necessary for specification. Thus, we analyzed the enrichment of DEGs (EC versus hPSC) proximal to (within ±40 kb of) EC-specific boundaries (i.e., boundaries gained in differentiation), hPSC-specific boundaries (i.e., boundaries lost in differentiation), and boundaries shared between hPSC and EC. We found that boundaries gained during specification are enriched in upregulated DEGs and depleted in downregulated DEGs (Figure 3D; Supplemental Table 8). Boundaries lost in specification, on the other hand, were deficient for DEGs, while shared boundaries were enriched in downregulated DEGs (Figure 3D; Supplemental Table 9). The endothelial cell marker gene *CD31* is an example of an upregulated DEG proximal to an EC-specific boundary: its expression increases as a nearby EC-specific TAD boundary forms (Figure 3E; Supplemental Figure 2B). The orphan nuclear receptor gene *NR2F2*, which is required for embryonic vascular morphogenesis^47^, presents a similar example: its expression increases as it comes to fall within an EC-nascent TAD (Figure 3F; Supplemental Figure 4C). These observations indicate that the acquisition of new TAD boundaries is linked to the expression of genes that support endothelial cell specification.

### Long-range pairwise point interactions increase over differentiation and are associated with both gene activation and repression

Increasing evidence supports the importance of DNA pairwise point interactions (PPIs; local peaks of elevated contact frequency) in transcriptional regulation and suggests the existence of PPIs that function in specific aspects of development^40,48,49^. These interactions are thought to arise from the clustering of regulatory elements and genes through chromatin looping mechanisms^7^. Using the point interaction-calling package HiCCUPs^7^, we sought to investigate the formation of PPIs during endothelial cell differentiation. We identified increasing numbers of PPIs over the course of endothelial cell differentiation: 623 in hPSC to 3,881 in EC (Figure 4A). Most loops are specific to one time point; few loops are shared by more than two time points (Figure 4A). We observed the greatest number of stage-specific loops in EC (Figure 4A), suggesting a role for PPI-mediated transcriptional regulation as the endothelial cell lineage matures. In endothelial cell differenitation, long-range *cis* chromatin contacts increase overtime (Figure 1B–D), compelling us to examine the distances between PPI anchors across differentiation. Consistent with that trend, PPI sizes expand over differentiation, reflecting an increasing prevalence of long-range contacts between specific pairs of loci (Figure 4B). Thus, endothelial cell differentiation sees PPIs increase in both frequency and distance.

**Figure 4.**
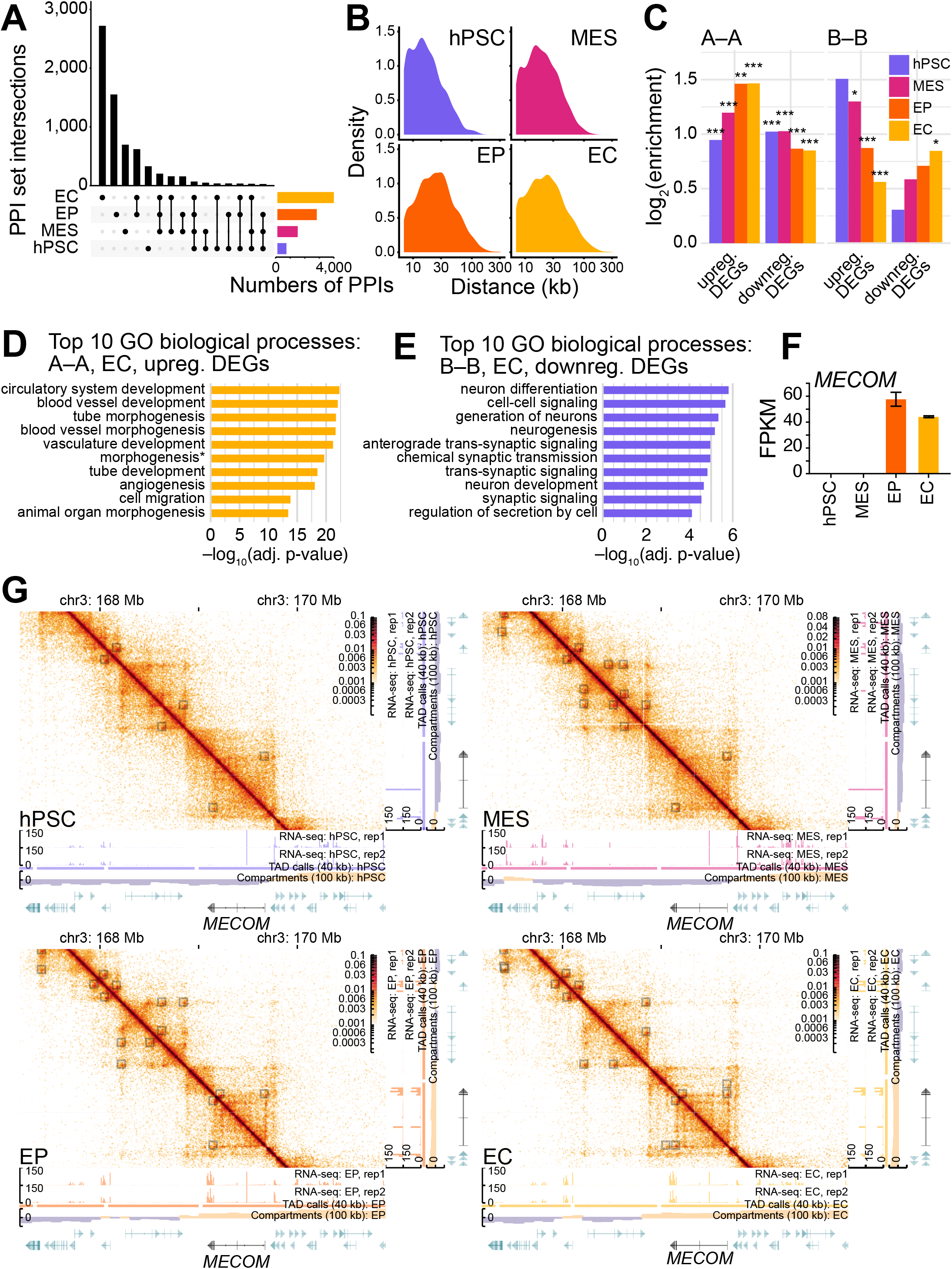
**A**. UpSet plot showing the intersections of HiCCUPS^7^ pairwise point interaction (PPI) anchors from Hi-C samples (10-kb resolution, autosomes) taken across endothelial cell differentiation. Vertical bars, intersection sizes for PPI anchors across datasets; horizontal bars, sample set sizes; solid black circles, anchors present; linked solid black circles, anchors present and shared in the associated samples; solid gray circles, anchors absent. **B**. Density plots showing the distributions of distances between PPI anchors from Hi-C samples (10-kb resolution, autosomes). **C**. Bar plots showing the log_2_ enrichment (see Methods) of differentially expressed genes (DEGs) at PPI anchors stratified by compartment type (A–A, A–B, and B–B; 100-kb resolution, autosomes) for Hi-C samples (10-kb resolution, autosomes). DEGs were called via DESeq2 analysis of EC versus hPSC (adjusted p-value < 0.05, absolute log_2_ fold change > 1; see Methods). Enrichment significantly different via chi-squared tests with Yates correction for set-wise overlaps, * < 0.05, ** < 0.01, *** < 0.001. **D, E**. Bar charts depicting adjusted p-values for the top 10 gene ontology (GO) biological process (BP) terms for (**D**) genes associated with PPIs in A-compartment bins (100-kb resolution, autosomes) and (**E**) genes associated with PPIs in B-compartment bins (500-kb resolution, autosomes). GO biological process terms significantly different from hypergeometric tests with Bonferroni corrections. Terms with asterisks have been abbreviated: “morphogenesis” is “anatomical structure formation involved in morphogenesis.” **F**. Bar plot for the RNA-seq expression levels of *MECOM* across endothelial cell differentiation. Expression levels: FPKM; error bars: standard error of the mean (SEM). **G**. Visualization of PPIs associated with *MECOM* (chr3: 169.08–169.66 Mb) in hPSC (top left), MES (top right), EP (bottom left), and EC (bottom right) Hi-C samples. Heatmaps and tracks for normalized Hi-C interaction frequencies (10-kb resolution), RNA-seq signal (unadjusted), TADs as delimited by the insulation score method (40-kb resolution; see Methods), genomic compartments (100-kb resolution; gold, A compartment; purple, B compartment), and genes (green and black); solid squares overlying the heatmaps, PPIs called by HiCCUPS^7^ (see Methods).

Since PPI dynamics occur amid changes in genomic compartmentalization (Figure 2; Supplemental Figure 3), we investigated the interrelationships between PPIs and genomic compartments. We found that the proportion of PPIs with both anchors in the A compartment (A–A) increases from MES to EC (Supplemental Figure 5A, B). In the same period, the proportion of PPIs with both anchors in the B compartment (B–B) decreases (Supplemental Figure 5A, B). To investigate whether nascent PPIs in A compartments are involved in the transcriptional activation of genes essential to endothelial cell development and differentiation, we evaluated the genome-wide enrichment of DEGs between hPSC and EC at time point-specific PPI anchors in A and B compartments. We observed an enrichment of DEGs, both up- and downregulated, at PPIs in all time points (Figure 4C; Supplemental Table 10). The enrichment of PPIs at upregulated DEGs is elevated in comparison to downregulated DEGs (Figure 4C). PPIs anchored in A compartments are enriched for upregulated DEGs and depleted for downregulated DEGs(Figure 4C); the enriched upregulated DEGs are associated with numerous functions in endothelial cell specification (Figure 4D; Supplemental Table 11). This trend was reversed for PPIs anchored in B compartments, which were enriched for downregulated DEGs and depleted for upregulated DEGs (Figure 4C); the genes associated with B-compartment PPIs have numerous functions in neurogenesis (Figure 4E; Supplemental Table 12), raising the possibility that B-compartment PPIs function in a transcriptional repression mechanism that inhibits neuronal specification. Taken together, our data suggest that PPIs facilitate both transcriptional activation within the A compartment and transcriptional repression within the B compartment.

Several genes with well-established roles in endothelial cell biology associate with PPIs as their transcription increases in development. A prominent example is *MECOM* (MDS1 and EVI1 complex locus), a transcription factor that promotes arterial EC identity^50^, *MECOM* is expressed in EP and EC (Figure 4F), and comes to overlap increasing numbers of PPIs in the course of differentiation (Figure 4G). Additional examples include *VEGFC* (vascular endothelial growth factor C; Supplemental Figure 5C, D), which codes for a protein critical for angiogenesis, endothelial cell growth, and blood vessel permeability in vascular and lymphatic vessels^51,52^; *KDR* (kinase insert domain receptor; Supplemental Figure 5E, F), which encodes a *VEGF* receptor^53^; and *TFPI* (tissue factor pathway inhibitor; Supplemental Figure 5G, H), a gene that encodes a serine protease inhibitor with anti-coagulative effects^54–56^. Numerous genes repressed in endothelial cell differentiation are associated with B-compartment PPIs, including genes associated with tissue patterning and neuronal development. These genes include *EPHB6* (ephrin type-B receptor 6; Supplemental Figure 6A, B), which codes for a pseudokinase member of the Eph receptor family^57–60^; *GABRB3* (gamma-aminobutyric acid type A receptor subunit beta 3) and *GABRA5* (gamma-aminobutyric acid type A receptor subunit alpha 3; Supplemental Figure 6C, D), both of which encode receptor subunits for the neurotransmitter GABA; and *PTPRZ1* (protein tyrosine phosphatase receptor type Z1; Supplemental Figure 6E, F), which codes for a member of the receptor protein tyrosine phosphatase family that is largely restricted to the central nervous system in development^61^. Additionally, we observed examples of B-compartment PPIs associating with genes that code for factors with anti-angiogenic properties. Examples include *FOXC1* (forkhead box C1; Supplemental Figure 6G, H), whose protein product has known antagonistic roles in vascular development^62^; and *ISM1* (isthmin 1; Supplemental Figure 6I, J), which codes for a secreted protein that functions as an endogenous angiogenesis inhibitor^63^.

### Chromatin organization reveals the developmental divergence of endothelial cells and cardiomyocytes

Our stem cell model for endothelial cell development shares initial steps with cardiomyocyte differentiation^36^. Given their common developmental origin from cardiogenic mesoderm, we sought to compare the genome organizational dynamics of the two lineages. We reanalyzed our published time-course cardiomyocyte datasets^9^ alongside our new time-course endothelial cell datasets. To evaluate differences in *cis* contacts between EC and cardiomyocytes (CM), we took the log_2_ ratio of EC and CM contacts for a single chromosome, chromosome 3. We noted clear differences in EC contacts versus CM contacts (Figure 5A), including more near-range cis interactions in EC interactions (red, along the matrix diagonal), while CM exhibited more long-range interactions (blue, spread further away from the diagonal). Surveying the *cis* contact probabilities *P(s)* for EC and CM, we found that, beyond 30 Mb, there was a higher probability for *cis* contacts in CM versus EC (Figure 5B). These results indicate that, although long-range *cis* contacts increase as hPSC differentiate to become EC (Figure 1B–D), longer-range *cis* contacts are present in—and a prominent feature of—genome organization in CM.

**Figure 5.**
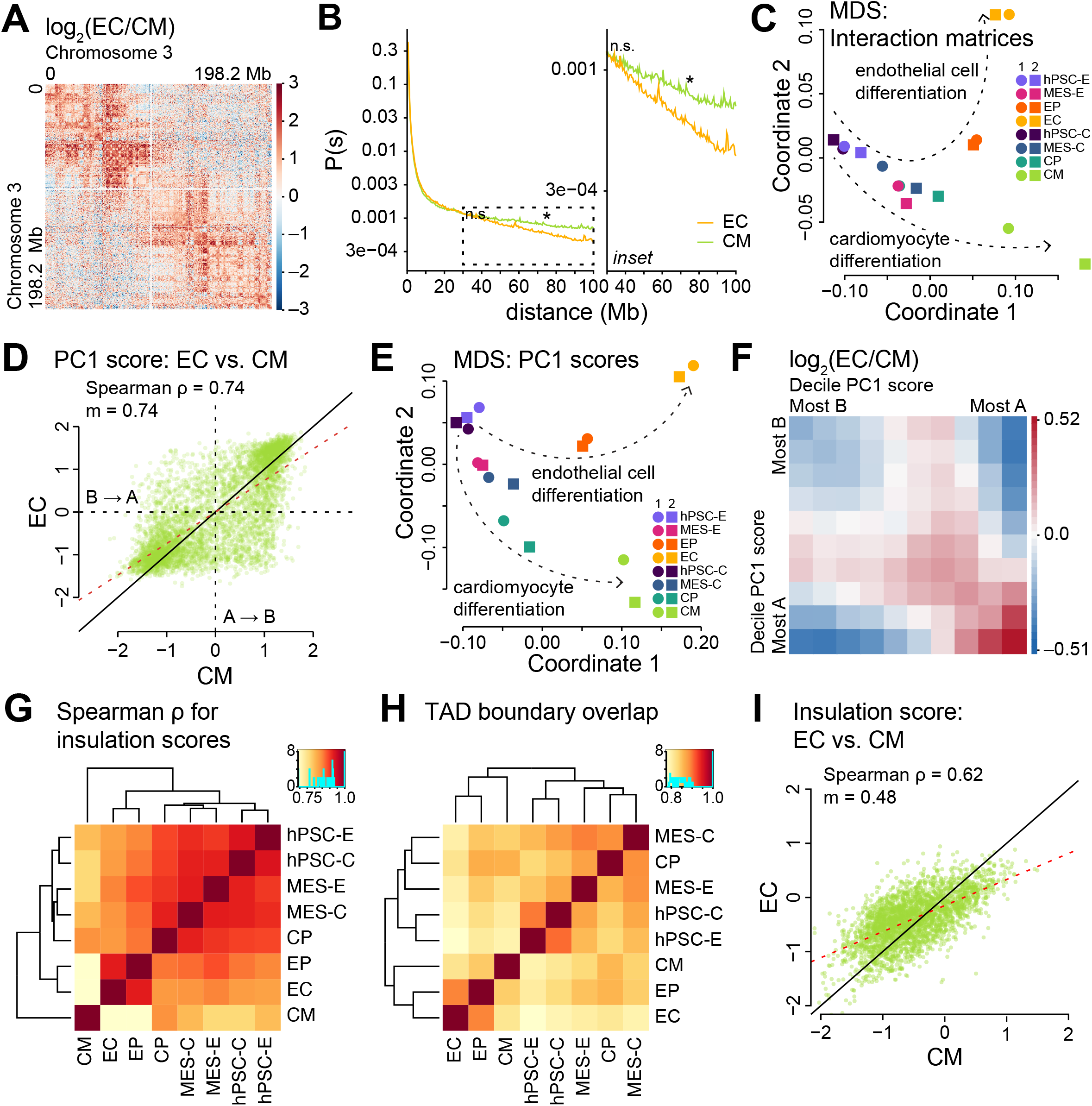
**A**. log_2_ ratios of normalized Hi-C interaction frequencies (500-kb resolution, chromosome 3) for EC vs. cardiomyocytes (CM). Red, observed/expected interactions higher in EC; blue, observed/expected interactions higher in CM. **B**. *Cis* interaction frequency probabilities *P* stratified by genomic distance *s* for EC and CM Hi-C samples (500-kb resolution, all autosomes). *P* stratified by s over 30–100 Mb (inset, right). P-value significantly different at *s* of 30 and 75 Mb via t-tests between EC and CM with false discovery rate correction, * < 0.05, n.s. (not significantly different). **C**. MDS projection of Spearman correlation coefficients (ρ) for Hi-C samples (500-kb resolution, autosomes) taken from across endothelial cell differentiation. Similarity measure: 1 – ρ. hPSC-E: human pluripotent stem cell from endothelial cell differentiation; MES-E: mesoderm cell from endothelial cell differentiation; EP: endothelial cell progenitor; EC: endothelial cell; hPSC-C: human pluripotent stem cell from cardiomyocyte differentiation; MES-C: mesoderm cell from cardiomyocyte differentiation; CP: cardiomyocyte progenitor cell; CM: cardiomyocyte. **D**. Scatter plot comparing PC1 scores derived from EC and CM Hi-C samples (500-kb resolution, autosomes). ρ, Spearman correlation coefficient; m, regression slope; red dashed line, regression line; black solid line, x = y; A → B, A-to-B compartment switches; B → A, B-to-A compartment switches. **E**. MDS projection of PC1 scores for Hi-C samples (500-kb resolution, autosomes) taken from endothelial cell and cardiomyocyte differentiation. Similarity measure: 1 − ρ. **F**. log2 ratios of saddle plots for EC and CM Hi-C samples (500-kb resolution, autosomes); red, observed/expected interactions higher in EC; blue, observed/expected interactions higher in CM. **G**. Hierarchically clustered heatmap of Spearman correlation coefficients for insulation scores from Hi-C samples (40-kb resolution, autosomes) taken across endothelial cell and cardiomyocyte differentiation. **H**. Hierarchically clustered heatmap of the TAD-boundary set intersections of Hi-C samples (40-kb resolution, autosomes) taken across endothelial cell and cardiomyocyte differentiation. **I**. Scatter plot comparing insulation scores derived from CM and EC Hi-C samples (40-kb resolution, autosomes). ρ, Spearman correlation coefficients; m, regression slope; black dashed line, regression line; black solid line, x = y.

To compare and contrast the influence of *cis* chromatin interactions as endothelial cells and cardiomyocytes differentiate from their common origin, we performed MDS of Hi-C interaction maps. MDS of intrachromosomal chromatin interactions pairs revealed two divergent differentiation trajectories, one for endothelial cells and the other for cardiomyocytes (Figure 5C). Consistent with the initial similarity of their stem cell differentiation protocols, the early hPSC and MES cell types clustered together regardless of their differentiation protocol of origin. Also, the cardiomyocyte progenitor (CP) cell type was observed amid the hPSC and MES cell types. Our findings underscore the relevance of varying trends in *cis* contacts as the two cell types differentiate.

Scatter plots for PC1 scores in EC versus CM revealed marked changes in compartment status in EC versus CM (Figure 5D; Supplemental Figure 7A). This led us to consider the influence of genomic compartments on endothelial cell and cardiomyocyte data structure. Similar to MDS of intrachromosomal chromatin interactions (Figure 5C), MDS of PC1 scores paired replicates—including the same cell states from the two protocols, hPSC and MES—and outlined divergent differentiation trajectories (Figure 5E). Hierarchical clustering of Spearman correlation coefficients (ρ) for PC1 scores paired and largely ordered replicates (Supplemental Figure 7B), revealing that EC have the most distinct compartment profiles. To examine gross changes in genomic compartment strength in the divergence of the two cell types, we calculated and plotted the log_2_ ratio of EC and CM saddle plots. We found that, in EC versus CM, *cis* interactions in B compartments are generally weaker, and *cis* interactions between A compartments are generally stronger (Figure 5F; Supplemental Figure 7C). Taken together, these data reveal genomic compartmentalization as an important distinguishing feature of the two cell lineages. Considering their varying transcriptomes^9^, dynamic compartmentalization likely influences the two transcription programs.

Finally, we asked whether TAD dynamics also varied between EC and CM. First, hierarchical clustering of TAD insulation scores grouped “early” cell states such as hPSC and MES—and also CP—while separating out “later” cell states such as EP, EC, and CM (Figure 5G). Of note, insulation scores in CM were markedly different from all other cell types (Figure 5G). Next, we performed hierarchical clustering of TAD boundary set intersections. The set intersections clustered similarly to the TAD insulation scores (Figure 5G), separating “early” cell states (hPSC, MES, and CP) from “later” cell states (EP, EC, and CM; Figure 5H). We sought to home in on TAD dynamics across EC and CM by drawing scatter plots for insulation scores, which revealed marked changes in insulation scores in EC versus CM (Figure 5I; Supplemental Figure 7D)—a trend in which CM insulation scores tend to be higher when EC insulation scores are lower, and vice versa. Together, these results suggest a different chromatin topology in EC versus CM—one in which intra-TAD chromatin interactions are tightly bounded in ECs, preventing longer-range chromatin interactions. On the other hand, in CMs, intra-TAD interactions are less restricted and, thus, open to even longer *cis* chromatin interactions.

## Discussion

In this study, we established a model system in which hPSCs are differentiated to ECs, and we used this system to present a comprehensive look at chromatin organization—alone and with respect to the dynamic transcriptomes of differentiation—in endothelial cell development. To our knowledge, endothelial cell chromatin organization has been examined in only one other study^6^, which compared Hi-C results from human umbilical vein endothelial cells to publicly available datasets from hESCs and mesendoderm cells. In contrast to the work from Niskanen et al., our study makes use of a developmental time course in an isogenic context, allowing for detailed analyses of chromatin organization across endothelial cell specification.

In our examination of genomic compartmentalization, we found that ∼25% of the genome is dynamic in endothelial cell differentiation (Figure 2A, B), transitioning from transcriptionally active, euchromatic A compartments to repressive, heterochromatic B compartments or vice versa. Regions that transition from B to A are enriched in upregulated DEGs related to endothelial cell specification (Figure 2E; Supplemental Table 5); regions that transition from A to B are enriched in downregulated DEGs related to neurogenesis (Figure 2F, Supplemental Table 6). We also noted the increasing strength of B compartments in differentiation (Figure 2I; Supplemental Figure 3D), suggesting the compaction of chromatin^9^. Given the shifting transcriptomes across endothelial cell differentiation (Supplemental Figure 2A, B), it is possible that dynamic, strengthening B compartments function to repress transcription, perhaps through chromatin inaccessibility mechanisms. These findings are in line with recent Hi-C analyses of cardiomyocytes^9^’^65^ and, taken together, indicate that dynamic genomic compartmentalization is an important, conserved regulator of cell type-specific transcription.

In addition to genomic compartmentalization, we assessed two other forms of chromatin organization: TADs (Figure 3) and PPIs (Figure 4). TADs and PPIs are similar in that both arise through a process of chromatin loop extrusion^66–70^. In loop extrusion, the multi-subunit protein complex cohesin entraps one or more small loops of chromatin inside its lumen; then, through progressive extrusion, the loops are enlarged up to the Mb scale, ceasing to grow in size when cohesin colocalizes with the transcription factor/insulator protein CTCF, the CTCF cofactor MAZ^71^, and likely other “architectural proteins”^72^. Such loops of chromatin are formed almost exclusively between convergent consensus sequences bound by CTCF^7,73–75^. It has been suggested that cohesin is subject to rapid turnover^69,76^, limiting the timespan for cohesin-chromatin interactions and, thus, loop extrusion. It is also known that cohesins can extend beyond convergent CTCF-bound consensus sequences: under conditions where cohesin turnover is prolonged or stopped, cohesin appears to move beyond convergent CTCF-bound motifs^77,78^. We observed that most PPIs are unique to each stage of endothelial cell differentiation, with each successive stage in our time point analysis seeing more and more PPIs (Figure 4A; Supplemental Figure 5A). The EC stage exhibits the most PPIs (Figure 4A; Supplemental Figure 5A) in addition to the strongest TAD boundaries and highest number of intra-TAD contacts (Figure 3C; Supplemental Figure 4C, D). Considering these findings together, we speculate that, in the course of maturation, cohesin loading is increased and turnover is prolonged, allowing for strengthened TAD boundaries and increased numbers of PPIs. These findings are consistent with observations from Niskanen and colleagues^6^, who reported high levels of TAD connectivity, identifying endothelial cell-specific long-range interactions between TADs enriched for histone H3 trimethylated at lysine 9 (H3K9me3), a marker of constitutive heterochromatin^6^.

While work remains to understand how the strengthened TADs and numerous PPIs arise in endothelial cells, our findings indicate that these features support transcription necessary for differentiation: strengthened TAD boundaries are accompanied by increased numbers of intra-TAD contacts (Figure 3C; Supplemental Figure 4C, D), nascent TAD boundaries are enriched for upregulated DEGs (Figure 3D–F), and PPI anchors correlate with upregulated DEGs in the A compartment (Figure 4C), consistent with mounting evidence that PPIs play an important role in the regulation of gene expression^40,48,49^. We showed a number of examples of PPIs forming in A compartments coincident with the upregulation of genes essential to endothelial cell biology (Figure 4D, F, G; Supplemental Figure 5C–H; Supplemental Figure 6). These findings suggest that TADs and PPIs, two interrelated forms of chromatin organization, promote gene expression necessary for endothelial function.

Finally, we performed comparative analyses of dynamic chromatin organization in endothelial cell specification versus cardiomyocyte specification (Figure 5). We observed that, although B compartments strengthen in endothelial cell differentiation (Figure 2I; Supplemental Figure 3D), they do not strengthen to the extent seen in cardiomyocyte differentiation (Figure 5F; Supplemental Figure 7C)^9^. Concomitant with this, insulation scores are generally lower at TAD boundaries in EC versus CM (Figure 5I), indicating that *cis* chromatin interactions are less restrained in EC versus CM. Taken together, these observations indicate that high levels of heterochromatinization and long-range chromatin contacts are prevalent features of genome organization in CM versus EC. The CM nuclear environment is notable for, among other things, a *trans*-interaction network of *TTN*-associated genes facilitated by the muscle-specific splicing protein RBM20^9^. Could it be that cardiomyocyte differentiation, with its preponderance of long-range interactions, sees the establishment of a nuclear environment that facilitates functionally significant regulatory *trans* interactions while endothelial cell development sees the development of a nuclear environment that favors regulation through predominantly *cis* forms chromatin organization, e.g., PPIs?

Attempting to address this and other questions will undoubtedly fuel further research. With this study, we provide a comprehensive analysis of 3D chromatin organization in a model of endothelial cell development that will be a key resource for studying endothelial cell biology in health and disease.

## Acknowledgements

We thank Dr. Choli Lee and the lab of Dr. Jay Shendure (Department of Genome Sciences, University of Washington) for assistance with sequencing, Nicole Zeinstra and Dr. Ying Zheng (Department of Bioengineering, University of Washington) for expertise and advice regarding endothelial cells, and the University of Washington Center for Nuclear Organization and Function Group for feedback and discussion. We gratefully acknowledge the Tom and Sue Ellison Stem Cell Core of the Institute for Stem Cell and Regenerative Medicine for use of cell culture space and equipment. This research was supported by the Cell Analysis Facility Flow Cytometry and Imaging Core in the Department of Immunology at the University of Washington.

## Supplemental Material

Figures S1–S7

Tables S1–S12

