## Supplemental Figures for "Dynamic chromatin organization and regulatory interactions in human endothelial cell differentiation"

**A** Karyotype analysis: RUES2 embryonic stem cell line

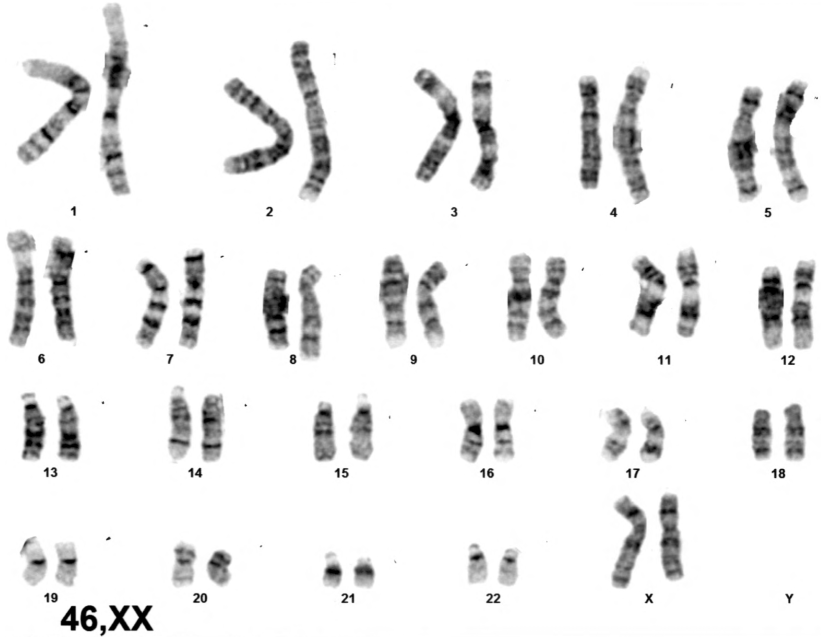


**B** Endothelial cell progenitor (EP)

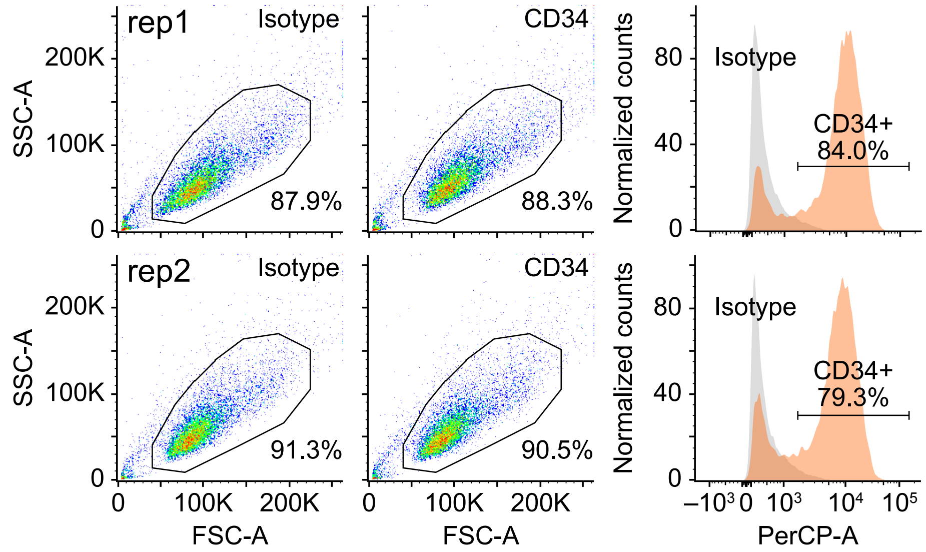


**C** Endothelial cell (EC)

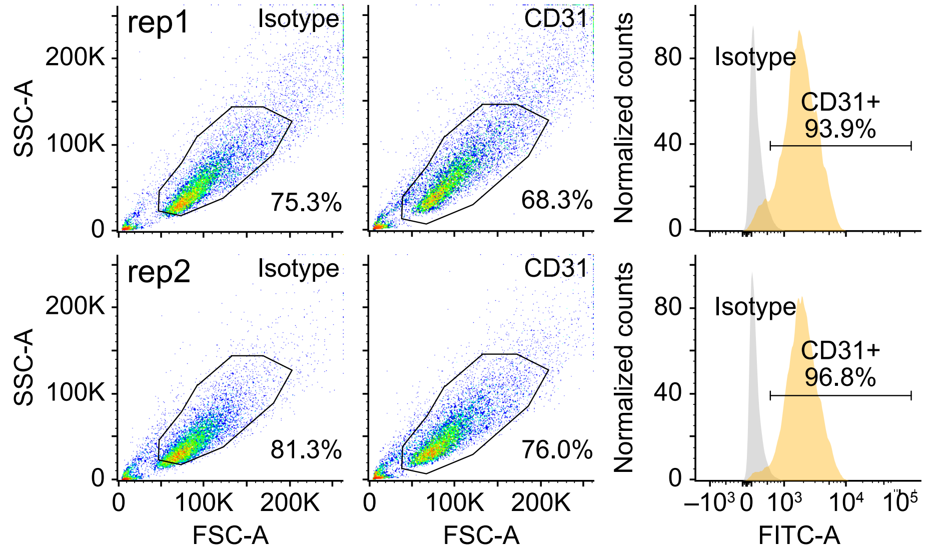

Figure S1 (related to Figure 1)

**A.** Karyotype analysis of RUES2 human embryonic stem cells demonstrating a normal 46,XX pattern.

**B.** Left, flow cytometry scatter plot for endothelial cell progenitors (EP) via immunostaining against human anti-IgG antibody, a negative control. Middle, flow cytometry scatter plot indicating EP purity via immunostaining against mouse anti-human CD34-PerCP antibody. Right, overlapping histograms for EP stained against the IgG isotype (gray) and CD34 (orange). Counts were normalized to modes. Top, replicate 1; bottom, replicate 2.

**C.** Left, flow cytometry scatter plot for endothelial cells (EC) as in panel A. Middle, flow cytometry scatter plot indicating EC purity via immunostaining against mouse anti-human CD31-FITC antibody. Right, overlapping histograms of EC stained against the IgG isotype (gray) and CD31 (yellow). Counts were normalized to modes. Top, replicate 1; bottom, replicate 2.

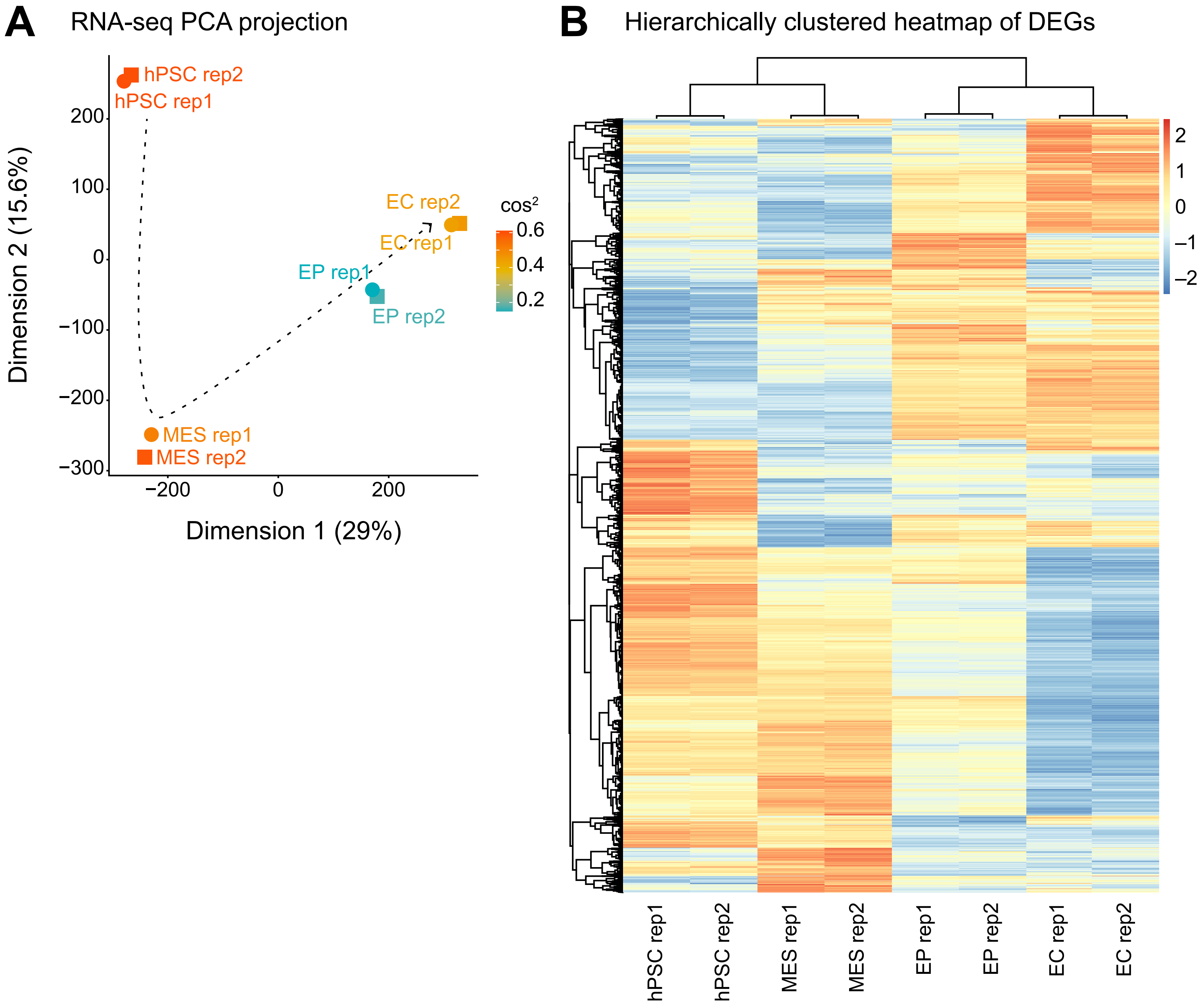
A
RNA-seq PCA projection
hPSC rep2
hPSC rep1
EC rep2
EC rep1
EP rep1
EP rep2
MES rep1
MES rep2
cos²
0.6
0.4
0.2
200
100
0
−100
−200
−300
Dimension 2 (15.6%)
−200
0
200
Dimension 1 (29%)
B
Hierarchically clustered heatmap of DEGs
2
1
0
−1
−2
hPSC rep1
hPSC rep2
MES rep1
MES rep2
EP rep1
EP rep2
EC rep1
EC rep2

Figure S2 (related to Figure 1)

**A.** PCA plots for fragments per kilobase of transcript per million mapped reads (FPKM) from RNA-seq samples taken across endothelial cell differentiation. hPSC: human pluripotent stem cell; MES: mesoderm cell; EP: endothelial cell progenitor; EC: endothelial cell; rep: replicate; PC: principal component. Percentages of variance captured by the principal components are described in axis titles; the scale is cosine squared.

**B.** Hierarchically clustered heatmap of differentially expressed genes (DEGs) across endothelial cell differentiation. Row clustering is based on the four time points of differentiation; the column dendrogram is ordered to match the time point of differentiation. Adjusted p-value < 0.05, absolute $\log_2$ fold change > 1. Color scale: $\log_2$ fold changes in expression (FPKM).

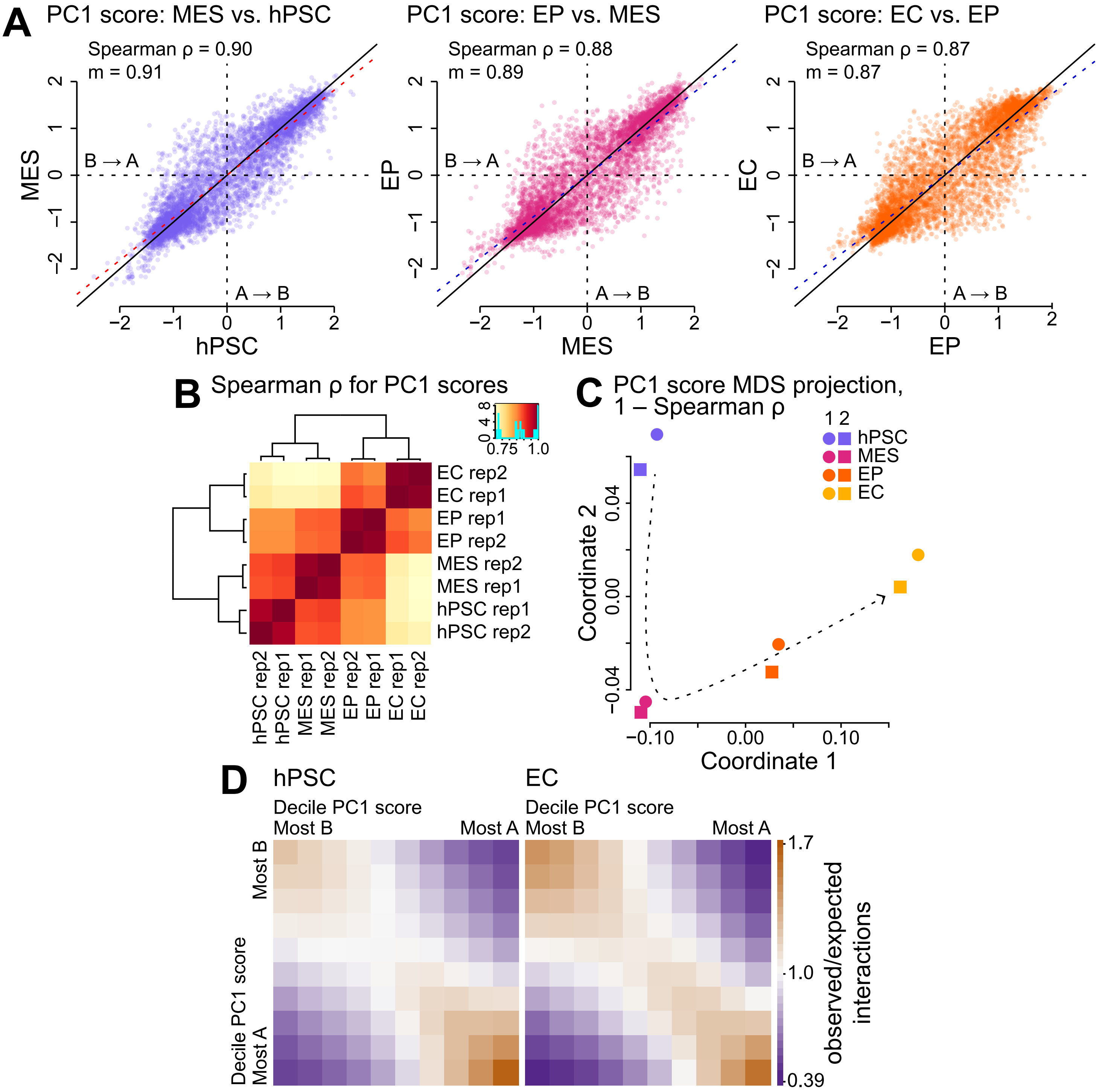




Alavattam, Mitzelfelt et al. – Figure S3

Figure S3 (related to Figure 2)

**A.** Scatter plots comparing Hi-C sample (500-kb resolution, autosomes) PC1 scores from consecutive stages of endothelial cell differentiation: hPSC versus MES, MES versus EP, and EP versus EC. ρ, Spearman correlation coefficient; m, regression slope; red dashed line, regression line; black solid line, x = y; A → B, A-to-B compartment switches; B → A, B-to-A compartment switches.

**B.** Hierarchically clustered heatmap of Spearman correlation coefficients (ρ) for PC1 scores from Hi-C samples (500-kb resolution, autosomes).

**C.** MDS projection of PC1 scores for Hi-C samples (500-kb resolution, autosomes); similarity measure: 1 – Spearman correlation coefficient (ρ).

**D.** Saddle plots—i.e., 10 × 10 decile-binned matrices quantifying the strength of PC1 scores—for EC and hPSC Hi-C samples (500-kb resolution, autosomes).

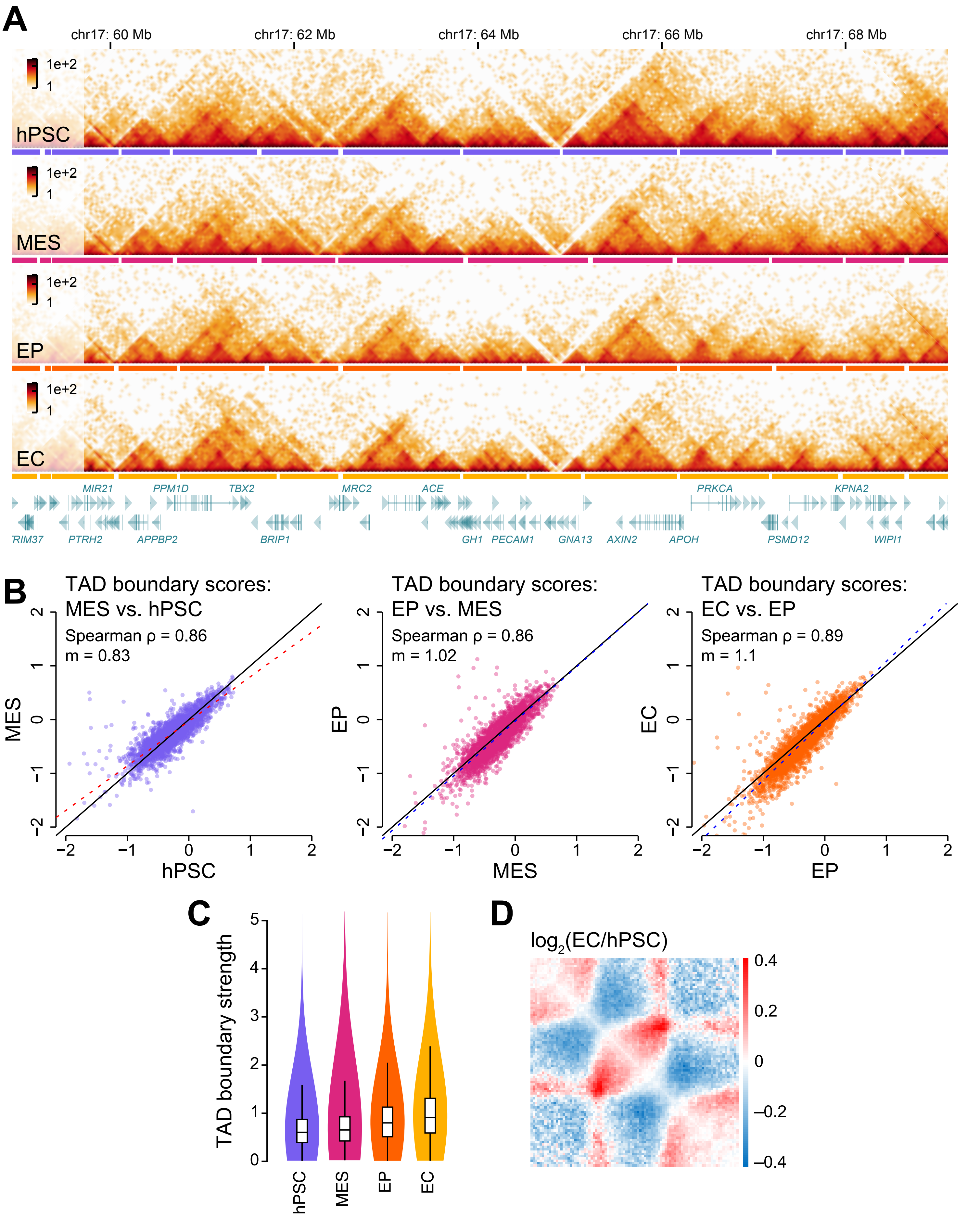
A
chr17: 60 Mb
chr17: 62 Mb
chr17: 64 Mb
chr17: 66 Mb
chr17: 68 Mb
1e+2
1
hPSC
MES
EP
EC
MIR21
PPM1D
TBX2
MRC2
ACE
PRKCA
KPNA2
RIM37
PTRH2
APPBP2
BRIP1
GH1
PECAM1
GNA13
AXIN2
APOH
PSMD12
WIPI1
B
TAD boundary scores: MES vs. hPSC
Spearman ρ = 0.86
m = 0.83
TAD boundary scores: EP vs. MES
Spearman ρ = 0.86
m = 1.02
TAD boundary scores: EC vs. EP
Spearman ρ = 0.89
m = 1.1
MES
EP
EC
hPSC
C
TAD boundary strength
D
log2(EC/hPSC)
0.4
0.2
0
−0.2
−0.4

Figure S4 (related to Figure 3)

**A.** Hi-C interaction heatmaps (40-kb bins, chromosome 17, approximately 59–69 Mb) showing dynamics of local interactions and TADs in hPSC, MES, EP, and EC. Horizontal solid bars, TADs as delimited by the insulation score methods (see Methods); gaps in horizontal solid bars, TAD boundaries; bottom row, genes (green).

**B.** Scatter plots comparing Hi-C sample (40-kb resolution, autosomes) insulation scores at TAD boundaries: hPSC versus MES, MES versus EP, and EP versus EC. ρ, Spearman correlation coefficients; m, regression slope; red or blue dashed line, regression line; black solid line, x = y.

**C.** $\log_2$ ratio heatmap for an EC aggregate TAD plot over an hPSC aggregate TAD plot; red, interaction frequency higher in EC; blue, interaction frequency higher in hPSC.

**D.** Box-and-whisker plots imposed over violin plots showing TAD boundary strengths for Hi-C samples (40-kb resolution, autosomes). Box-and-whisker plots present the 25th percentile, median, and 75th percentile with whiskers extending to 1.5 times the interquartile range.

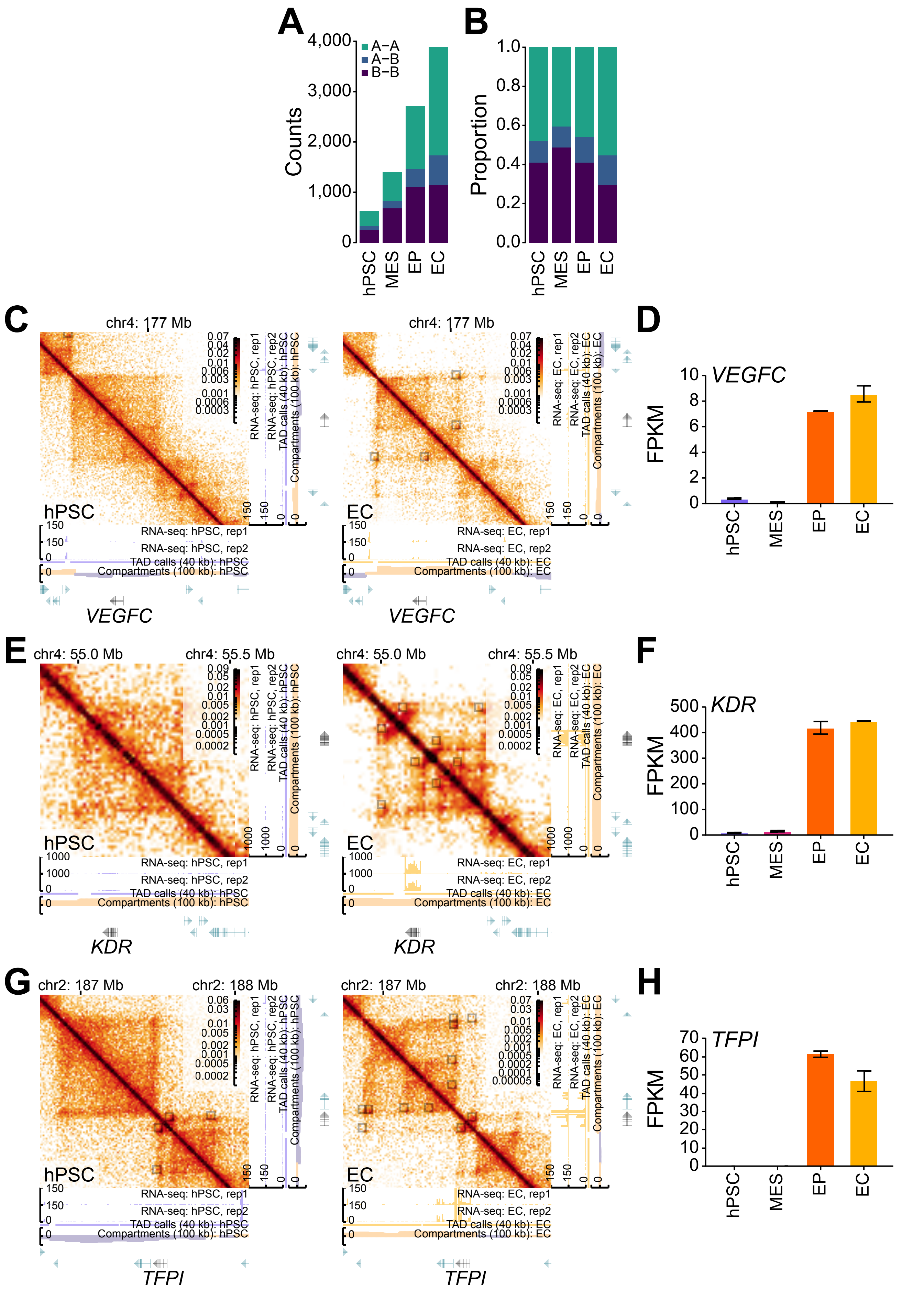




Alavattam, Mitzelfelt et al. – Figure S5

Figure S5 (related to Figure 4)

**A, B.** Stacked bar charts showing the absolute (**A**) and relative (**B**) numbers of PPIs in each Hi-C sample (10-kb resolution, autosomes). Bars are stratified by PPI-anchor compartment (100-kb resolution, autosomes) of origin: A–A, both anchors in A compartments; A–B, one anchor in an A compartment, the other anchor in a B compartment; B–B, both anchors in B compartments.

**C, E, G.** Visualization of PPIs associated with *VEGFC* (**C**), *KDR* (**E**), and *TFPI* (**G**) in hPSC (left) and EC (right) Hi-C samples (10-kb bins, autosomes). Heatmaps of normalized Hi-C interaction frequencies (10-kb resolution), RNA-seq signal (unadjusted), TADs as delimited by the insulation score method (40-kb resolution; see Methods), genomic compartments (100-kb resolution; gold, A compartment; purple, B compartment), and genes (green and black); solid squares overlying the heatmaps, PPIs called by HiCCUPS[7] (see Methods).

**D, F, H.** Bar plots for the RNA-seq expression levels of *VEGFC* (**D**), *KDR* (**F**), and *TFPI* (**H**) across endothelial cell differentiation. Expression levels: FPKM; error bars: standard error of the mean (SEM).

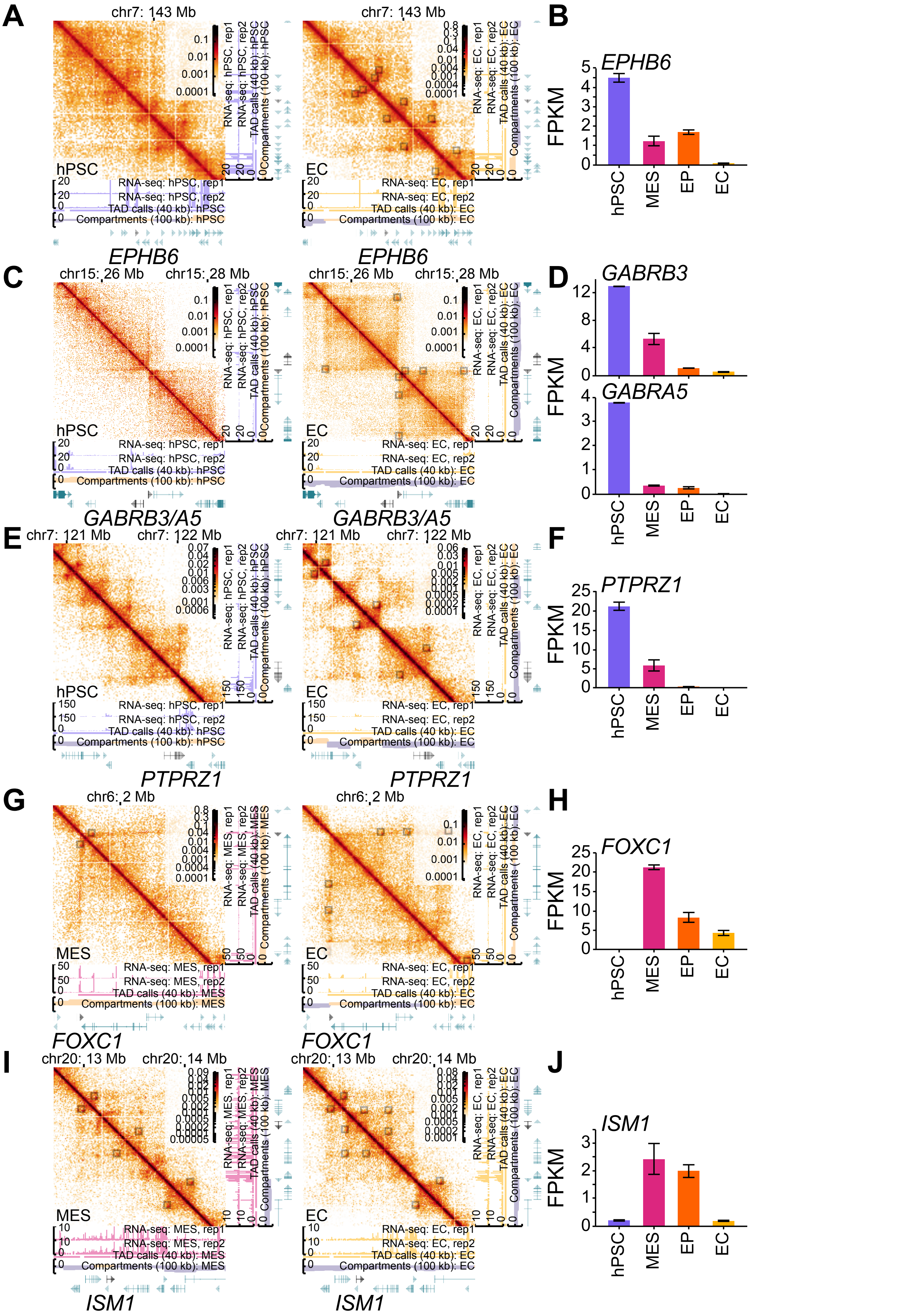
A
chr7: 143 Mb
hPSC
RNA-seq: hPSC, rep1
RNA-seq: hPSC, rep2
TAD calls (40 kb): hPSC
Compartments (100 kb): hPSC
EC
RNA-seq: EC, rep1
RNA-seq: EC, rep2
TAD calls (40 kb): EC
Compartments (100 kb): EC
EPHB6
B
EPHB6
FPKM
hPSC
MES
EP
EC
C
chr15: 26 Mb
chr15: 28 Mb
GABRB3/A5
D
GABRB3
GABRA5
E
chr7: 121 Mb
chr7: 122 Mb
PTPRZ1
F
G
chr6: 2 Mb
MES
RNA-seq: MES, rep1
RNA-seq: MES, rep2
TAD calls (40 kb): MES
Compartments (100 kb): MES
FOXC1
H
I
chr20: 13 Mb
chr20: 14 Mb
ISM1
J

Figure S6 (related to Figure 4)

**A, C, E, G, I.** Visualization of PPIs associated with *EPHB6* (**A**), *GABRB3* and *GABRA5* (**C**), *PTPRZ1* (**E**), *FOXC1* (**G**), and *ISM1* (**I**) in hPSC or MES (left), and EC (right) Hi-C samples (10-kb bins, autosomes). Heatmaps of normalized Hi-C interaction frequencies (10-kb resolution), RNA-seq signal (unadjusted), TADs as delimited by the insulation score method (40-kb resolution; see Methods), genomic compartments (100-kb resolution; gold, A compartment; purple, B compartment), and genes (green and black); solid squares overlying the heatmaps, PPIs called by HiCCUPS[7] (see Methods).

**B, D, F, H, J.** Bar plots for the RNA-seq expression levels of *EPHB6* (**B**), *GABRA3/5* (**D**), *PTPRZ1* (**F**), *FOXC1* (**H**), and *ISM1* (**J**) across endothelial cell differentiation. Expression levels: FPKM; error bars: standard error of the mean (SEM).

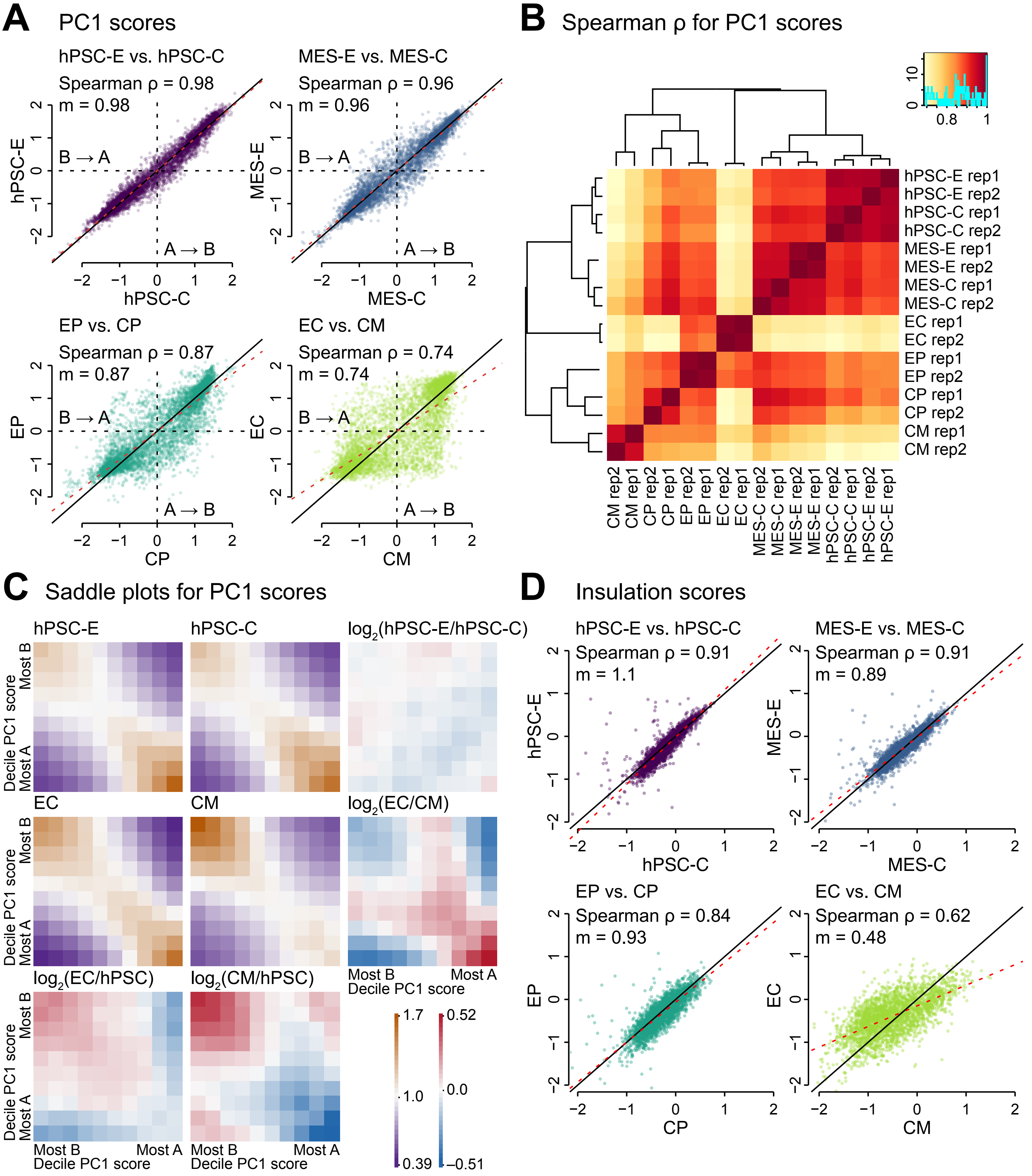
A
PC1 scores
hPSC-E vs. hPSC-C
Spearman ρ = 0.98
m = 0.98
MES-E vs. MES-C
Spearman ρ = 0.96
m = 0.96
EP vs. CP
Spearman ρ = 0.87
m = 0.87
EC vs. CM
Spearman ρ = 0.74
m = 0.74
B → A
A → B
hPSC-E
hPSC-C
MES-E
MES-C
EP
CP
EC
CM
B
Spearman ρ for PC1 scores
hPSC-E rep1
hPSC-E rep2
hPSC-C rep1
hPSC-C rep2
MES-E rep1
MES-E rep2
MES-C rep1
MES-C rep2
EC rep1
EC rep2
EP rep1
EP rep2
CP rep1
CP rep2
CM rep1
CM rep2
C
Saddle plots for PC1 scores
hPSC-E
hPSC-C
log2(hPSC-E/hPSC-C)
EC
CM
log2(EC/CM)
log2(EC/hPSC)
log2(CM/hPSC)
Decile PC1 score
Most B
Most A
1.7
1.0
0.39
0.52
0.0
−0.51
D
Insulation scores
hPSC-E vs. hPSC-C
Spearman ρ = 0.91
m = 1.1
MES-E vs. MES-C
Spearman ρ = 0.91
m = 0.89
EP vs. CP
Spearman ρ = 0.84
m = 0.93
EC vs. CM
Spearman ρ = 0.62
m = 0.48

Figure S7 (related to Figure 5)

**A.** Hierarchically clustered heatmap of Spearman correlation coefficients (ρ) for PC1 scores from Hi-C samples (500-kb resolution, autosomes). hPSC-E: human pluripotent stem cell from endothelial cell differentiation; MES-E: mesoderm cell from endothelial cell differentiation; EP: endothelial cell progenitor; EC: endothelial cell; hPSC-C: human pluripotent stem cell from cardiomyocyte differentiation; MES-C: mesoderm cell from cardiomyocyte differentiation; CP: cardiomyocyte progenitor cell; CM: cardiomyocyte.

**B.** Scatter plots comparing Hi-C sample (500-kb resolution, autosomes) PC1 scores from comparable time points in endothelial cell and cardiomyocyte differentiation: hPSC-E versus hPSC-C, MES-E versus MES-C, EP versus CP, and EC versus CM. ρ, Spearman correlation coefficient; m, regression slope; red dashed line, regression line; black solid line, x = y; A → B, A-to-B compartment switches; B → A, B-to-A compartment switches.

**C.** Saddle plots and for hPSC-E, hPSC-C, EC, and CM Hi-C samples (500-kb resolution, autosomes). Gold-to-purple color bar: observed/expected interactions. Red-to-blue color bar: $\log_2$ ratios of saddle plots; red, observed/expected interactions higher in numerator; blue, observed/expected interactions higher in denominator.

**D.** Scatter plots comparing Hi-C sample (40-kb resolution, autosomes) insulation scores from comparable time points in endothelial cell and cardiomyocyte differentiation: hPSC-E versus hPSC-C, MES-E versus MES-C, EP versus CP, and EC versus CM. ρ, Spearman correlation coefficient; m, regression slope; red dashed line, regression line; black solid line, x = y.
