## Supplemental Tables for "Dynamic chromatin organization and regulatory interactions in human endothelial cell differentiation"

### Table S1

RNA-seq sequencing metrics.

### Table S2

Hi-C sequencing metrics (part 1).

### Table S3

Hi-C sequencing metrics (part 2).

### Table S4

HiCRep scores.

### Table S5

Gene ontology (GO) terms (molecular function, biological process, and cellular component) associated with upregulated differentially expressed genes (DEGs) between EC versus hPSC (i.e., DEGs upregulated in EC, downregulated in hPSC) in regions that undergo B-to-A compartment transitions. P-values were obtained from hypergeometric tests and adjusted via Bonferroni corrections. List was filtered for terms with adjusted p-value < 0.05.

### Table S6

GO terms associated with downregulated DEGs between EC versus hPSC (i.e., DEGs downregulated in EC, upregulated in hPSC) in regions that undergo A-to-B compartment transitions. P-values were obtained from hypergeometric tests and adjusted via Bonferroni corrections. List was filtered for terms with adjusted p-value < 0.05.

### Table S7

Proportions of overlapping topologically associating domain (TAD) boundaries.

### Table S8

GO terms associated with upregulated DEGs between EC versus hPSC (i.e., DEGs upregulated in EC, downregulated in hPSC) at regions of the genome proximal to EC-specific TAD boundaries (i.e., TAD boundaries gained in differentiation). P-values were obtained from hypergeometric tests and adjusted via Bonferroni corrections. List was filtered for terms with adjusted p-value < 0.05.

### Table S9

GO terms associated with downregulated DEGs between EC versus hPSC (i.e., DEGs downregulated in EC, upregulated in hPSC) at regions of the genome proximal to shared TAD boundaries (i.e., TAD boundaries maintained in differentiation). P-values were obtained from hypergeometric tests and adjusted via Bonferroni corrections. List was filtered for terms with adjusted p-value < 0.05.

#### Table S10

Counts for DEGs associated with pairwise point interaction (PPI) anchors.

#### Table S11

GO terms associated with upregulated DEGs between EC versus hPSC (i.e., DEGs upregulated in EC, downregulated in hPSC) at EC A–A PPI anchors.  P-values were obtained from hypergeometric tests and adjusted via Bonferroni corrections. List was filtered for terms with adjusted p-value < 0.05.

#### Table S12

GO terms associated with downregulated DEGs between EC versus hPSC (i.e., DEGs downregulated in EC, upregulated in hPSC) at EC B–B PPI anchors. P-values were obtained from hypergeometric tests and adjusted via Bonferroni corrections. List was filtered for terms with adjusted p-value < 0.05.

| sample | replicate | paired_reads | uniquely_mapped_pairs | percent_uniquely_mapped | pairs_mapped_to_multiple_loci | percent_mapped_to_multiple_loci |
|---|---|---|---|---|---|---|
| hPSC | rep1 | 55,840,122 | 47,422,601 | 84.93 | 3,858,200 | 6.91 |
| hPSC | rep2 | 60,990,868 | 47,651,876 | 78.13 | 7,963,072 | 13.06 |
| MES | rep1 | 71,563,794 | 57,297,844 | 80.07 | 8,275,320 | 11.56 |
| MES | rep2 | 64,653,133 | 51,517,204 | 79.68 | 7,928,694 | 12.26 |
| EP | rep1 | 65,060,462 | 52,029,304 | 79.97 | 7,627,257 | 11.72 |
| EP | rep2 | 61,589,114 | 51,653,553 | 83.87 | 4,868,759 | 7.91 |
| EC | rep1 | 82,628,666 | 68,215,173 | 82.56 | 7,426,107 | 8.99 |
| EC | rep2 | 59,453,709 | 51,350,419 | 86.37 | 3,215,213 | 5.41 |

| sample | replicate | valid_pairs | unique_valid_pairs | percent_unique | trans_pairs | percent_trans_pairs | cis_pairs | percent_cis_pairs | cis_pairs_lt_20_kb | percent_cis_pairs_lt_20_kb | cis_pairs_gt_20_kb | percent_cis_pairs_gt_20_kb | ratio_cis_pairs_to_trans_pairs |
|---|---|---|---|---|---|---|---|---|---|---|---|---|---|
| hPSC | rep1 | 74,370,707 | 70,430,033 | 94.70% | 15,852,694 | 22.51% | 54,577,339 | 77.49% | 11,957,541 | 16.98% | 42,619,798 | 60.51% | 3.44 |
| hPSC | rep2 | 42,377,743 | 41,083,070 | 96.94% | 10,417,752 | 25.36% | 30,665,318 | 74.64% | 9,247,969 | 22.51% | 21,417,349 | 52.13% | 2.94 |
| MES | rep1 | 61,324,069 | 58,129,315 | 94.79% | 15,493,362 | 26.65% | 42,635,953 | 73.35% | 14,409,563 | 24.79% | 28,226,390 | 48.56% | 2.75 |
| MES | rep2 | 64,750,191 | 61,993,009 | 95.74% | 17,061,390 | 27.52% | 44,931,619 | 72.48% | 13,752,296 | 22.18% | 31,179,323 | 50.29% | 2.63 |
| EP | rep1 | 72,043,347 | 67,430,109 | 93.60% | 15,147,301 | 22.46% | 52,282,808 | 77.54% | 15,981,933 | 23.70% | 36,300,875 | 53.83% | 3.45 |
| EP | rep2 | 64,189,111 | 61,378,014 | 95.62% | 11,857,704 | 19.32% | 49,520,310 | 80.68% | 15,180,445 | 24.73% | 34,339,865 | 55.95% | 4.18 |
| EC | rep1 | 67,497,290 | 63,474,609 | 94.04% | 15,107,705 | 23.80% | 48,366,904 | 76.20% | 14,649,787 | 23.08% | 33,717,117 | 53.12% | 3.20 |
| EC | rep2 | 74,553,571 | 69,616,473 | 93.38% | 24,929,031 | 35.81% | 44,687,442 | 64.19% | 13,059,499 | 18.76% | 31,627,943 | 45.43% | 1.79 |

| sample | replicate | forward-forward | reverse-reverse | reverse-forward | forward-reverse |
|---|---|---|---|---|---|
| hPSC | rep1 | 18,606,062 | 18,579,762 | 18,546,266 | 18,638,617 |
| hPSC | rep2 | 10,600,843 | 10,585,697 | 10,554,213 | 10,636,990 |
| MES | rep1 | 15,335,934 | 15,315,408 | 15,235,875 | 15,436,852 |
| MES | rep2 | 16,192,785 | 16,169,040 | 16,092,651 | 16,295,715 |
| EP | rep1 | 18,029,243 | 17,991,746 | 17,919,020 | 18,103,338 |
| EP | rep2 | 16,057,326 | 16,029,086 | 15,948,463 | 16,154,236 |
| EC | rep1 | 16,883,002 | 16,852,483 | 16,786,922 | 16,974,883 |
| EC | rep2 | 18,650,095 | 18,618,145 | 18,566,860 | 18,718,471 |

| | hPSC rep1 | hPSC rep2 | MES rep1 | MES rep2 | EP rep1 | EP rep2 | EC rep1 | EC rep2 |
|---|---|---|---|---|---|---|---|---|
| hPSC rep1 | 1 | 0.97776915 | 0.88736242 | 0.91134917 | 0.79180177 | 0.80364539 | 0.75778713 | 0.77991904 |
| hPSC rep2 | 0.97776915 | 1 | 0.88919682 | 0.91270621 | 0.77742386 | 0.79013307 | 0.75691099 | 0.79203572 |
| MES rep1 | 0.88736242 | 0.88919682 | 1 | 0.98719636 | 0.88840099 | 0.89999116 | 0.79604168 | 0.81160056 |
| MES rep2 | 0.91134917 | 0.91270621 | 0.98719636 | 1 | 0.88098611 | 0.89169317 | 0.80860903 | 0.82887643 |
| EP rep1 | 0.79180177 | 0.77742386 | 0.88840099 | 0.88098611 | 1 | 0.99422268 | 0.9036348 | 0.88366946 |
| EP rep2 | 0.80364539 | 0.79013307 | 0.89999116 | 0.89169317 | 0.99422268 | 1 | 0.902065 | 0.88354703 |
| EC rep1 | 0.75778713 | 0.75691099 | 0.79604168 | 0.80860903 | 0.9036348 | 0.902065 | 1 | 0.98197549 |
| EC rep2 | 0.77991904 | 0.79203572 | 0.81160056 | 0.82887643 | 0.88366946 | 0.88354703 | 0.98197549 | 1 |

| Category | ID | Name | p-value | q-value Bonferroni | q-value FDR B&H | q-value FDR B&Y | Hit Count in Query List | Hit Count in Genome |
|---|---|---|---|---|---|---|---|---|
| GO: Molecular Function | GO:0005102 | signaling receptor binding | 6.87E-07 | 6.61E-04 | 6.61E-04 | 4.93E-03 | 71 | 1842 |
| GO: Molecular Function | GO:0030246 | carbohydrate binding | 2.14E-05 | 2.06E-02 | 7.00E-03 | 5.21E-02 | 19 | 295 |
| GO: Molecular Function | GO:0005161 | platelet-derived growth factor receptor binding | 2.18E-05 | 2.10E-02 | 7.00E-03 | 5.21E-02 | 5 | 17 |
| GO: Molecular Function | GO:0005201 | extracellular matrix structural constituent | 4.63E-05 | 4.46E-02 | 1.11E-02 | 8.30E-02 | 14 | 185 |
| GO: Biological Process | GO:0035239 | tube morphogenesis | 2.28E-17 | 1.32E-13 | 1.32E-13 | 1.22E-12 | 69 | 1068 |
| GO: Biological Process | GO:0035295 | tube development | 5.75E-16 | 3.33E-12 | 1.67E-12 | 1.54E-11 | 75 | 1310 |
| GO: Biological Process | GO:0001568 | blood vessel development | 8.82E-15 | 5.11E-11 | 1.70E-11 | 1.57E-10 | 57 | 866 |
| GO: Biological Process | GO:0048646 | anatomical structure formation involved in morphogenesis | 1.59E-14 | 9.21E-11 | 2.30E-11 | 2.13E-10 | 75 | 1395 |
| GO: Biological Process | GO:0048514 | blood vessel morphogenesis | 4.92E-14 | 2.85E-10 | 5.03E-11 | 4.65E-10 | 52 | 768 |
| GO: Biological Process | GO:0001944 | vasculature development | 5.25E-14 | 3.04E-10 | 5.03E-11 | 4.65E-10 | 57 | 903 |
| GO: Biological Process | GO:0072359 | circulatory system development | 6.08E-14 | 3.52E-10 | 5.03E-11 | 4.65E-10 | 72 | 1340 |
| GO: Biological Process | GO:0016477 | cell migration | 3.15E-12 | 1.82E-08 | 2.28E-09 | 2.11E-08 | 83 | 1812 |
| GO: Biological Process | GO:0007155 | cell adhesion | 6.16E-12 | 3.57E-08 | 3.96E-09 | 3.66E-08 | 75 | 1571 |
| GO: Biological Process | GO:0022610 | biological adhesion | 7.63E-12 | 4.42E-08 | 4.42E-09 | 4.08E-08 | 75 | 1578 |
| GO: Biological Process | GO:0040012 | regulation of locomotion | 9.24E-11 | 5.35E-07 | 4.87E-08 | 4.50E-07 | 61 | 1210 |
| GO: Biological Process | GO:0001525 | angiogenesis | 1.03E-10 | 5.95E-07 | 4.96E-08 | 4.58E-07 | 42 | 658 |
| GO: Biological Process | GO:0030334 | regulation of cell migration | 2.89E-10 | 1.68E-06 | 1.29E-07 | 1.19E-06 | 56 | 1089 |
| GO: Biological Process | GO:0009887 | animal organ morphogenesis | 5.06E-10 | 2.93E-06 | 2.09E-07 | 1.94E-06 | 61 | 1263 |
| GO: Biological Process | GO:0007160 | cell-matrix adhesion | 5.77E-10 | 3.34E-06 | 2.23E-07 | 2.06E-06 | 24 | 252 |
| GO: Biological Process | GO:2000145 | regulation of cell motility | 1.04E-09 | 6.03E-06 | 3.77E-07 | 3.49E-06 | 57 | 1159 |
| GO: Biological Process | GO:0051270 | regulation of cellular component movement | 2.40E-09 | 1.39E-05 | 8.18E-07 | 7.56E-06 | 59 | 1250 |
| GO: Biological Process | GO:0051240 | positive regulation of multicellular organismal process | 2.92E-09 | 1.69E-05 | 9.40E-07 | 8.69E-06 | 72 | 1692 |
| GO: Biological Process | GO:0030155 | regulation of cell adhesion | 3.47E-09 | 2.01E-05 | 1.06E-06 | 9.78E-06 | 45 | 827 |
| GO: Biological Process | GO:0030198 | extracellular matrix organization | 2.91E-08 | 1.69E-04 | 8.44E-06 | 7.80E-05 | 29 | 431 |
| GO: Biological Process | GO:0043062 | extracellular structure organization | 3.06E-08 | 1.78E-04 | 8.45E-06 | 7.81E-05 | 29 | 432 |
| GO: Biological Process | GO:0045229 | external encapsulating structure organization | 3.39E-08 | 1.96E-04 | 8.92E-06 | 8.25E-05 | 29 | 434 |
| GO: Biological Process | GO:0048661 | positive regulation of smooth muscle cell proliferation | 5.96E-08 | 3.45E-04 | 1.44E-05 | 1.33E-04 | 15 | 127 |
| GO: Biological Process | GO:0050900 | leukocyte migration | 5.97E-08 | 3.46E-04 | 1.44E-05 | 1.33E-04 | 33 | 554 |
| GO: Biological Process | GO:0071495 | cellular response to endogenous stimulus | 6.94E-08 | 4.02E-04 | 1.61E-05 | 1.49E-04 | 63 | 1510 |
| GO: Biological Process | GO:0040017 | positive regulation of locomotion | 8.69E-08 | 5.04E-04 | 1.87E-05 | 1.73E-04 | 37 | 678 |
| GO: Biological Process | GO:0032101 | regulation of response to external stimulus | 8.72E-08 | 5.05E-04 | 1.87E-05 | 1.73E-04 | 52 | 1147 |
| GO: Biological Process | GO:0048729 | tissue morphogenesis | 1.40E-07 | 8.10E-04 | 2.88E-05 | 2.66E-04 | 43 | 874 |
| GO: Biological Process | GO:0030335 | positive regulation of cell migration | 1.44E-07 | 8.35E-04 | 2.88E-05 | 2.66E-04 | 35 | 633 |
| GO: Biological Process | GO:0042127 | regulation of cell population proliferation | 1.49E-07 | 8.63E-04 | 2.88E-05 | 2.66E-04 | 75 | 1973 |
| GO: Biological Process | GO:0001775 | cell activation | 1.87E-07 | 1.08E-03 | 3.49E-05 | 3.23E-04 | 65 | 1623 |
| GO: Biological Process | GO:0001953 | negative regulation of cell-matrix adhesion | 2.37E-07 | 1.37E-03 | 4.29E-05 | 3.97E-04 | 9 | 44 |
| GO: Biological Process | GO:0031589 | cell-substrate adhesion | 2.60E-07 | 1.51E-03 | 4.57E-05 | 4.22E-04 | 26 | 397 |
| GO: Biological Process | GO:0042060 | wound healing | 2.98E-07 | 1.73E-03 | 5.06E-05 | 4.68E-04 | 33 | 594 |
| GO: Biological Process | GO:1901701 | cellular response to oxygen-containing compound | 3.06E-07 | 1.77E-03 | 5.06E-05 | 4.68E-04 | 56 | 1330 |
| GO: Biological Process | GO:0009611 | response to wounding | 3.31E-07 | 1.92E-03 | 5.27E-05 | 4.87E-04 | 38 | 746 |
| GO: Biological Process | GO:0010812 | negative regulation of cell-substrate adhesion | 3.37E-07 | 1.95E-03 | 5.27E-05 | 4.87E-04 | 11 | 74 |
| GO: Biological Process | GO:0045321 | leukocyte activation | 4.15E-07 | 2.41E-03 | 6.19E-05 | 5.72E-04 | 59 | 1447 |
| GO: Biological Process | GO:2000147 | positive regulation of cell motility | 4.17E-07 | 2.41E-03 | 6.19E-05 | 5.72E-04 | 35 | 662 |
| GO: Biological Process | GO:0048660 | regulation of smooth muscle cell proliferation | 4.71E-07 | 2.73E-03 | 6.83E-05 | 6.31E-04 | 18 | 212 |
| GO: Biological Process | GO:0007507 | heart development | 5.11E-07 | 2.96E-03 | 7.22E-05 | 6.68E-04 | 36 | 698 |
| GO: Biological Process | GO:0072001 | renal system development | 5.50E-07 | 3.18E-03 | 7.58E-05 | 7.01E-04 | 24 | 360 |
| GO: Biological Process | GO:0032103 | positive regulation of response to external stimulus | 5.69E-07 | 3.30E-03 | 7.67E-05 | 7.09E-04 | 31 | 553 |
| GO: Biological Process | GO:0048659 | smooth muscle cell proliferation | 6.64E-07 | 3.85E-03 | 8.74E-05 | 8.08E-04 | 18 | 217 |
| GO: Biological Process | GO:0002274 | myeloid leukocyte activation | 6.92E-07 | 4.01E-03 | 8.91E-05 | 8.23E-04 | 36 | 707 |
| GO: Biological Process | GO:0006935 | chemotaxis | 7.64E-07 | 4.43E-03 | 9.63E-05 | 8.90E-04 | 36 | 710 |
| GO: Biological Process | GO:0051272 | positive regulation of cellular component movement | 8.03E-07 | 4.65E-03 | 9.90E-05 | 9.15E-04 | 35 | 681 |
| GO: Biological Process | GO:0007167 | enzyme linked receptor protein signaling pathway | 8.74E-07 | 5.06E-03 | 1.04E-04 | 9.65E-04 | 50 | 1168 |

Hit in Query List

HIF1A,TRAF6,PDE4D,ETS2,FABP4,JAM2,SLIT2,EFEMP1,KITLG,PIK3CG,JAG1,AHR,PLCL1,ARRDC3,OSGIN1,VEGFC,COL3A1,VIP,CNRIP1,APP,ASAH1,PDGFD,CLEC2D,LEPROT,IL1R1,CXCL8,DOCK4,IL16,ERAP1,INHBA,GATA3,ITPRID2,ITGA1,ITGA2,PTEN,ACE,ITGAV,ITGB1,[illegible]

ENPP2,CLEC1B,CLEC1A,AGL,CLN5,CD93,CLEC2D,LAYN,GBE1,KLRC4-KLRK1,PTX3,KLRD1,CLEC2B,CLEC12B,CLEC9A,OLR1,CD34,GPCPD1,CD69

PDGFD,IL1R1,PTEN,PDGFC,RASA1

THSD4,SLIT2,FBN2,EFEMP1,COL3A1,COL4A5,COL5A2,NID2,MMRN1,BMPER,IMPG2,LAMA4,LAMB1,THBS1

HIF1A,TRAF6,ENPP2,ETS1,HLX,MECOM,FZD4,SLIT2,DLC1,SAT1,PIK3CA,PIK3CG,HPGD,JAG1,AHR,CC2D2A,ADGRF5,ADGRF4,QKI,VEGFC,COL3A1,ROBO4,CEP290,TIPARP,SLC12A2,PRKACB,CXCL8,PLK2,ERAP1,SDCCAG8,GATA3,PTEN,ACE,ITGAV,ITGB1,GADD45A,ZEB2,PTHLH,[illegible]

HIF1A,TRAF6,ENPP2,ETS1,HLX,MECOM,FZD4,PHEX,SLIT2,DLC1,SAT1,PIK3CA,PIK3CG,HPGD,JAG1,AHR,CC2D2A,ADGRF5,ADGRF4,QKI,TNS3,VEGFC,COL3A1,ROBO4,CEP290,TIPARP,SLC12A2,ASAH1,PRKACB,CXCL8,PLK2,ERAP1,SDCCAG8,GATA3,PTEN,ACE,ITGAV,ITGB1,[illegible]

HIF1A,ENPP2,ETS1,FZD4,SLIT2,SAT1,PIK3CA,PIK3CG,HPGD,JAG1,AHR,ADGRF5,ADGRF4,QKI,VEGFC,COL3A1,ROBO4,TIPARP,SLC12A2,PDGFD,CXCL8,SERPINB7,PLK2,ERAP1,PTEN,ACE,ITGAV,ITGB1,GADD45A,ITGB8,NEDD4,BMP4,JUN,CXCR4,GLI3,BTG1,BMPER,TBX20,[illegible]

HIF1A,TRAF6,ERG,ENPP2,ETS1,CLEC1B,ETS2,ERVW-1,JAM2,SLIT2,FBN2,DLC1,SAT1,PIK3CA,PIK3CG,JAG1,AHR,CC2D2A,VEGFC,ROBO4,COL5A2,CEP290,ITGA8,SLC40A1,SLC12A2,ANKRD1,PRKACB,PRKAR1A,TANC1,CXCL8,PLK2,ERAP1,INHBA,SDCCAG8,GATA3,ITGA2,[illegible]

HIF1A,ENPP2,ETS1,FZD4,SLIT2,SAT1,PIK3CA,PIK3CG,HPGD,JAG1,AHR,ADGRF5,ADGRF4,QKI,VEGFC,COL3A1,ROBO4,TIPARP,SLC12A2,CXCL8,PLK2,ERAP1,PTEN,ACE,ITGAV,ITGB1,GADD45A,ITGB8,NEDD4,BMP4,JUN,CXCR4,BTG1,BMPER,TBX20,TCF4,AKT3,SASH1,RASA1,[illegible]

HIF1A,ENPP2,ETS1,FZD4,SLIT2,SAT1,PIK3CA,PIK3CG,HPGD,JAG1,AHR,ADGRF5,ADGRF4,QKI,VEGFC,COL3A1,ROBO4,TIPARP,SLC12A2,PDGFD,CXCL8,SERPINB7,PLK2,ERAP1,PTEN,ACE,ITGAV,ITGB1,GADD45A,ITGB8,NEDD4,BMP4,JUN,CXCR4,GLI3,BTG1,BMPER,TBX20,[illegible]

HIF1A,ERG,ENPP2,ETS1,MECOM,FZD4,SLIT2,DLC1,SAT1,PIK3CA,PIK3CG,HPGD,JAG1,AHR,CC2D2A,ADGRF5,ADGRF4,QKI,VEGFC,SOX6,COL3A1,ROBO4,CEP290,TIPARP,SLC12A2,ANKRD1,PDGFD,PRKAR1A,CXCL8,SERPINB7,PLK2,ERAP1,GATA3,PTEN,ACE,ITGAV,ITGB1,[illegible]

HIF1A,PDE4D,MOSPD2,PRR5L,ERG,ENPP2,ETS1,JAM2,SLIT2,EFEMP1,DLC1,KITLG,PIK3CA,CD99,PIK3CG,JAG1,ARID5B,MMP1,MMP7,MMP10,TNS3,OSGIN1,VEGFC,COL3A1,VIM,ROBO4,DOCK10,APP,SHROOM2,SLC12A2,PDGFD,CD274,FUT8,IL1R1,CDKN2B-AS1,PALLD,CXCL8,[illegible]

CDKN2A,PRTG,TRAF6,C2CD4B,ENPP2,ETS1,HLX,CD46,JAM2,FZD4,SLIT2,SERPINB8,DLC1,KITLG,PIK3CA,CD99,PIK3CG,JAG1,PCDH10,VEGFC,COL3A1,ROBO4,NID2,PPFIBP1,ITGA8,C2CD4A,APP,MMRN1,CD93,ARHGAP6,PRKAR1A,CD274,PALLD,CXCL8,PAG1,GATA3,CD55,[illegible]

CDKN2A,PRTG,TRAF6,C2CD4B,ENPP2,ETS1,HLX,CD46,JAM2,FZD4,SLIT2,SERPINB8,DLC1,KITLG,PIK3CA,CD99,PIK3CG,JAG1,PCDH10,VEGFC,COL3A1,ROBO4,NID2,PPFIBP1,ITGA8,C2CD4A,APP,MMRN1,CD93,ARHGAP6,PRKAR1A,CD274,PALLD,CXCL8,PAG1,GATA3,CD55,[illegible]

HIF1A,MOSPD2,PRR5L,ENPP2,ETS1,JAM2,SLIT2,EFEMP1,DLC1,KITLG,CD99,PIK3CG,JAG1,ARRDC3,MMP7,MMP10,OSGIN1,VEGFC,COL3A1,VIM,ROBO4,DOCK10,APP,PDGFD,CD274,SNCA,ARHGAP18,IL1R1,CDKN2B-AS1,CXCL8,DOCK4,PLK2,IL16,MYCBP2,GATA3,ITGA2,[illegible]

HIF1A,ENPP2,ETS1,SLIT2,SAT1,PIK3CA,PIK3CG,JAG1,AHR,VEGFC,ROBO4,SLC12A2,CXCL8,PLK2,ERAP1,PTEN,ACE,ITGAV,ITGB1,GADD45A,ITGB8,BMP4,JUN,CXCR4,BTG1,BMPER,TBX20,TCF4,AKT3,SASH1,ESM1,CAV1,S1PR1,LEPR,TGFBR2,THBS1,HEY1,CD34,CD36,ELK3,[illegible]

HIF1A,MOSPD2,PRR5L,ENPP2,ETS1,JAM2,SLIT2,EFEMP1,DLC1,KITLG,CD99,PIK3CG,JAG1,MMP7,MMP10,OSGIN1,VEGFC,COL3A1,VIM,ROBO4,DOCK10,APP,PDGFD,CD274,IL1R1,CDKN2B-AS1,CXCL8,DOCK4,PLK2,GATA3,ITGA2,KLRC4-KLRK1,PTEN,ACE,ITGAV,ITGB1,GADD45A,[illegible]

HIF1A,TRAF6,ERG,SLC26A4,HLX,MECOM,FZD4,PHEX,SLIT2,FBN2,EFEMP1,DLC1,JAG1,ARID5B,AHR,MLLT3,VEGFC,SOX6,COL3A1,FLI1,COL5A2,CEP290,ITGA8,SHROOM2,SLC40A1,TIPARP,SLC12A2,ANKRD1,NR5A2,PRKAR1A,DAAM1,CSGALNACT1,INHBA,GATA3,ITGA2,PTEN,ITGAV,[illegible]

CDKN2A,ENPP2,DLC1,JAG1,VEGFC,COL3A1,NID2,ITGA8,ARHGAP6,ITGA1,ITGA2,PTEN,ITGAV,ITGB1,ITGB8,PTPRK,SGCE,RASA1,THBS1,EFNA5,CD34,CD36,CDH13,PHLDB2

HIF1A,MOSPD2,PRR5L,ENPP2,ETS1,JAM2,SLIT2,EFEMP1,DLC1,KITLG,CD99,PIK3CG,JAG1,MMP7,MMP10,OSGIN1,VEGFC,COL3A1,VIM,ROBO4,DOCK10,APP,PDGFD,CD274,ARHGAP18,IL1R1,CDKN2B-AS1,CXCL8,DOCK4,PLK2,GATA3,ITGA2,KLRC4-KLRK1,PTEN,ACE,ITGAV,ITGB1,[illegible]

HIF1A,PDE4D,MOSPD2,PRR5L,ENPP2,ETS1,JAM2,SLIT2,EFEMP1,DLC1,KITLG,CD99,PIK3CG,JAG1,MMP7,MMP10,OSGIN1,VEGFC,COL3A1,VIM,ROBO4,DOCK10,APP,PDGFD,CD274,ARHGAP18,IL1R1,CDKN2B-AS1,CXCL8,DOCK4,PLK2,MYCBP2,GATA3,ITGA2,KLRC4-KLRK1,PTEN,ACE,[illegible]

HIF1A,TRAF6,PDE4D,ERG,ENPP2,ETS1,HLX,CD46,FABP4,FABP5,SLIT2,FBN2,KITLG,JAG1,QKI,VEGFC,SOX6,VIM,VIP,KLF6,DDX60,BIRC2,BIRC3,ENPP4,APP,CD274,IL1R1,SCN3B,CXCL8,SERPINB7,IL16,ERAP1,INHBA,GATA3,ITGA2,ACE,ITGB1,IL18R1,ZEB2,ITGB8,[illegible]

CDKN2A,TRAF6,C2CD4B,ENPP2,ETS1,HLX,CD46,JAM2,FZD4,DLC1,PIK3CA,PIK3CG,JAG1,VEGFC,C2CD4A,ARHGAP6,PRKAR1A,CD274,CXCL8,PAG1,GATA3,CD55,ITGA2,KLRC4-KLRK1,PTEN,ITGAV,ITGB1,BMP4,NUAK1,CXCR4,GLI3,BCL10,RASA1,LAMA4,LAMB1,CAV1,CBLB,[illegible]

THSD4,ETS1,JAM2,FBN2,MMP1,MMP7,MMP10,COL3A1,COL4A5,COL5A2,NID2,ITGA8,APP,CSGALNACT1,ITGA1,ITGA2,ITGAV,ITGB1,ITGB8,PTX3,LAMA4,LAMB1,CAV1,CAV2,THBS1,ADAMTS6,TLL1,CD47,PHLDB2

THSD4,ETS1,JAM2,FBN2,MMP1,MMP7,MMP10,COL3A1,COL4A5,COL5A2,NID2,ITGA8,APP,CSGALNACT1,ITGA1,ITGA2,ITGAV,ITGB1,ITGB8,PTX3,LAMA4,LAMB1,CAV1,CAV2,THBS1,ADAMTS6,TLL1,CD47,PHLDB2

THSD4,ETS1,JAM2,FBN2,MMP1,MMP7,MMP10,COL3A1,COL4A5,COL5A2,NID2,ITGA8,APP,CSGALNACT1,ITGA1,ITGA2,ITGAV,ITGB1,ITGB8,PTX3,LAMA4,LAMB1,CAV1,CAV2,THBS1,ADAMTS6,TLL1,CD47,PHLDB2

HIF1A,TRAF6,PIK3CA,HPGD,PDGFD,ITGA2,PPARGC1A,BMP4,JUN,S1PR1,TGFBR2,THBS1,NAMPT,CALCRL,CDH13

PDE4D,MOSPD2,JAM2,SLIT2,KITLG,PIK3CA,CD99,PIK3CG,MMP1,VEGFC,APP,SLC12A2,PDGFD,IL1R1,CXCL8,IL16,GATA3,ITGA1,ITGA2,KLRC4-KLRK1,ITGAV,ITGB1,CXCR4,ESAM,CAV1,CXCL1,CXCL2,S1PR1,OLR1,THBD,THBS1,CD34,CD47

PDE4D,GNG2,NREP,PHEX,SLIT2,FBN2,PIK3CA,PIK3CG,HPGD,PIP4K2A,AHR,SOX6,COL3A1,VIM,COL5A2,ITGA8,SESN3,APP,ANKRD1,PDGFD,NR5A2,PRKACB,PRKAR1A,PRKAR2B,LEPROT,SNCA,FUT8,CXCL8,INHBA,GATA3,PTEN,ITGB1,ITGB8,NEDD4,PPARGC1A,BMP4,JUN,TBC1D4,[illegible]

HIF1A,MOSPD2,ENPP2,ETS1,JAM2,SLIT2,KITLG,CD99,PIK3CG,MMP7,VEGFC,APP,PDGFD,CD274,IL1R1,CDKN2B-AS1,CXCL8,DOCK4,PLK2,IL16,GATA3,ITGA2,ITGAV,ITGB1,BMP4,JUN,CXCR4,PDGFC,AKT3,SASH1,LAMB1,CAV1,S1PR1,TGFBR2,THBS1,TMSB4X,CDH13

TRAF6,C2CD4B,MOSPD2,ETS1,FABP4,SLIT2,PIK3CG,VEGFC,DDX60,BIRC2,BIRC3,C2CD4A,ENPP4,APP,CASP12,SLC12A2,PDGFD,PRKACB,SNCA,IL1R1,CXCL8,IL16,MYCBP2,ERAP1,GATA3,CD55,ITGA2,KLRC4-KLRK1,PTEN,ACE,CXCR4,BCL10,KLRD1,CLEC12B,SASH1,CASP4,CAV1,[illegible]

HIF1A,TRAF6,ETS2,FZD4,SLIT2,DLC1,JAG1,AHR,CC2D2A,MLLT3,VEGFC,COL3A1,COL5A2,CEP290,ITGA8,SLC12A2,ANKRD1,PRKACB,PRKAR1A,DAAM1,INHBA,MYO9A,GATA3,ITGA2,PTEN,ITGAV,ITGB1,ZEB2,PTHLH,BMP4,CXCR4,GLI3,BCL10,TBX20,FAT4,LAMB1,NR3C1,S1PR1,[illegible]

HIF1A,MOSPD2,ENPP2,ETS1,JAM2,KITLG,CD99,PIK3CG,MMP7,VEGFC,APP,PDGFD,CD274,IL1R1,CDKN2B-AS1,CXCL8,DOCK4,PLK2,GATA3,ITGA2,ITGAV,ITGB1,BMP4,JUN,CXCR4,PDGFC,AKT3,SASH1,LAMB1,CAV1,S1PR1,TGFBR2,THBS1,TMSB4X,CDH13

CDKN2A,HIF1A,TRAF6,ENPP2,ETS1,HLX,MECOM,CD46,FABP4,SLIT2,EFEMP1,DLC1,KITLG,SAT1,PIK3CA,HPGD,JAG1,AHR,MMP7,TNS3,OSGIN1,VEGFC,VIM,VIP,BIRC2,APP,IFIT3,PDGFD,NR5A2,PRKAR1A,CD274,CXCL8,SERPINB7,TES,INHBA,GATA3,CD55,ITGA1,ITGA2,PTEN,ACE,[illegible]

RAB9B,TRAF6,MOSPD2,GNG2,CLEC1B,HLX,CD46,FABP5,PIK3CA,PIK3CG,AGL,AHR,ADGRF5,COL3A1,KLF6,CEP290,DOCK10,ENPP4,APP,CD93,COMMD9,ASAH1,PRKAR1A,CD274,TMEM30A,SNCA,CXCL8,IL13RA2,PAG1,IL16,INHBA,ACTR10,GATA3,CD55,KLRC4-KLRK1,ITGAV,ITGB1,[illegible]

CDKN2A,ENPP2,DLC1,JAG1,ARHGAP6,PTEN,RASA1,THBS1,PHLDB2

CDKN2A,ENPP2,FZD4,DLC1,JAG1,VEGFC,COL3A1,NID2,ITGA8,ARHGAP6,ITGA1,ITGA2,PTEN,ITGAV,ITGB1,ITGB8,PTPRK,SGCE,RASA1,LAMB1,THBS1,EFNA5,CD34,CD36,CDH13,PHLDB2

HIF1A,GNG2,ETS1,CLEC1B,PIK3CA,PIK3CG,COL3A1,ENPP4,MMRN1,SLC12A2,PRKACB,PRKAR1A,PRKAR2B,GATA3,ITGA2,PTEN,ITGB1,DST,CXCR4,GLI3,C9,CAPZA2,CAV1,PAPSS2,ZFPM2,TFPI,TGFBR2,THBD,THBS1,CD34,CD36,ELK3,PHLDB2

HIF1A,TRAF6,PDE4D,GNG2,ETS1,ABCB4,FZD4,PHEX,SLIT2,PIK3CA,PIK3CG,PIP4K2A,AHR,COL3A1,VIM,COL5A2,SESN3,APP,ANKRD1,PDGFD,PRKACB,PRKAR1A,CD274,PRKAR2B,LEPROT,SNCA,CXCL8,INHBA,CD55,ITGA2,PTEN,PPARGC1A,BMP4,JUN,TBC1D4,PTPRK,BCL10,PDGFC,[illegible]

HIF1A,GNG2,ETS1,CLEC1B,NREP,PIK3CA,PIK3CG,CREB3L2,COL3A1,ENPP4,MMRN1,SLC12A2,PRKACB,PRKAR1A,PRKAR2B,INHBA,GATA3,ITGA2,PTEN,ITGB1,JUN,DST,CXCR4,GLI3,C9,CAPZA2,CAV1,PAPSS2,ZFPM2,TFPI,TGFBR2,THBD,THBS1,CD34,CD36,ELK3,PHLDB2,EPHA4

CDKN2A,ENPP2,FZD4,DLC1,JAG1,ARHGAP6,PTEN,RASA1,THBS1,EFNA5,PHLDB2

RAB9B,TRAF6,MOSPD2,HLX,CD46,FABP5,PIK3CA,PIK3CG,AGL,AHR,ADGRF5,KLF6,CEP290,DOCK10,ENPP4,APP,CD93,COMMD9,ASAH1,PRKAR1A,CD274,TMEM30A,SNCA,CXCL8,IL13RA2,PAG1,IL16,INHBA,ACTR10,GATA3,CD55,KLRC4-KLRK1,ITGAV,ITGB1,IL18R1,ITGB8,NEDD4,[illegible]

HIF1A,MOSPD2,ENPP2,ETS1,JAM2,KITLG,CD99,PIK3CG,MMP7,VEGFC,APP,PDGFD,CD274,IL1R1,CDKN2B-AS1,CXCL8,DOCK4,PLK2,GATA3,ITGA2,ITGAV,ITGB1,BMP4,JUN,CXCR4,PDGFC,AKT3,SASH1,LAMB1,CAV1,S1PR1,TGFBR2,THBS1,TMSB4X,CDH13

HIF1A,TRAF6,PIK3CA,HPGD,VIP,PDGFD,ITGA2,PTEN,PPARGC1A,BMP4,JUN,CAV1,S1PR1,TGFBR2,THBS1,NAMPT,CALCRL,CDH13

HIF1A,ERG,MECOM,SLIT2,DLC1,JAG1,AHR,CC2D2A,SOX6,COL3A1,CEP290,ANKRD1,PRKAR1A,GATA3,PTEN,ACE,ITGB1,NEDD4,BMP4,JUN,CXCR4,GLI3,KCNAB1,TBX20,GNG5,FAT4,CASP7,NR3C1,S1PR1,ZFPM2,ADGRG6,TGFBR2,THBS1,ADAMTS6,HEY1,CALCRL

MECOM,SLIT2,HPGD,JAG1,ARID5B,AHR,CC2D2A,CEP290,ITGA8,TIPARP,PDGFD,PYGO1,SERPINB7,GATA3,ACE,BMP4,CXCR4,GLI3,BMPER,FAT4,ADAMTS6,HEY1,CD34,EPHA4

TRAF6,C2CD4B,MOSPD2,ETS1,FABP4,SLIT2,PIK3CG,VEGFC,DDX60,C2CD4A,APP,PDGFD,PRKACB,SNCA,CXCL8,IL16,ITGA2,KLRC4-KLRK1,ACE,CXCR4,BCL10,KLRD1,SASH1,S1PR1,THBD,THBS1,LPL,CD47,TMSB4X,IL1RL1,CDH13

HIF1A,TRAF6,PIK3CA,HPGD,VIP,PDGFD,ITGA2,PTEN,PPARGC1A,BMP4,JUN,CAV1,S1PR1,TGFBR2,THBS1,NAMPT,CALCRL,CDH13

RAB9B,TRAF6,MOSPD2,FABP5,PIK3CG,AGL,ADGRF5,CEP290,ENPP4,APP,CD93,COMMD9,ASAH1,TMEM30A,SNCA,CXCL8,IL13RA2,IL16,ACTR10,CD55,ITGAV,ITGB8,JUN,PTX3,BCL10,CKAP4,CXCL1,ARPC5,OLR1,TGFBR2,THBS1,NAMPT,ATP6AP2,CD36,CD47,IL1RL1

PRTG,PDE4D,MOSPD2,ENPP2,SLIT2,PIK3CA,PIK3CG,VEGFC,ROBO4,ISL2,APP,SLC12A2,PDGFD,PALLD,CXCL8,DOCK4,IL16,MYCBP2,GATA3,ITGA1,ITGA2,KLRC4-KLRK1,ITGAV,BMP4,CXCR4,NFIB,GLI3,CXCL1,CXCL2,S1PR1,ABLIM1,THBS1,EFNA5,TMSB4X,CDH13,EPHA4

HIF1A,MOSPD2,ENPP2,ETS1,JAM2,KITLG,CD99,PIK3CG,MMP7,VEGFC,APP,PDGFD,CD274,IL1R1,CDKN2B-AS1,CXCL8,DOCK4,PLK2,GATA3,ITGA2,ITGAV,ITGB1,BMP4,JUN,CXCR4,PDGFC,AKT3,SASH1,LAMB1,CAV1,S1PR1,TGFBR2,THBS1,TMSB4X,CDH13

HIF1A,NREP,FZD4,FBN2,EFEMP1,PIK3CA,HPGD,PIP4K2A,ARID5B,VEGFC,COL3A1,VIM,COL4A5,ITGA8,SESN3,TIPARP,PDGFD,LEPROT,SNCA,FUT8,INHBA,GATA3,ITGA1,PTEN,ITGAV,ITGB1,ITGB8,NEDD4,BMP4,JUN,PTPRK,BMPER,TBX20,PDGFC,TCF4,FAT4,RASA1,VEPH1,ESM1,[illegible]

| | | | | | | | | |
|---|---|---|---|---|---|---|---|---|
| GO: Biological Process | GO:0033002 | muscle cell proliferation | 8.87E−07 | 5.14E−03 | 1.04E−04 | 9.65E−04 | 21 | 293 |
| GO: Biological Process | GO:0042330 | taxis | 9.01E−07 | 5.22E−03 | 1.04E−04 | 9.65E−04 | 36 | 715 |
| GO: Biological Process | GO:0001822 | kidney development | 1.05E−06 | 6.07E−03 | 1.19E−04 | 1.10E−03 | 23 | 347 |
| GO: Biological Process | GO:0009719 | response to endogenous stimulus | 1.11E−06 | 6.41E−03 | 1.23E−04 | 1.14E−03 | 67 | 1780 |
| GO: Biological Process | GO:0001570 | vasculogenesis | 1.32E−06 | 7.64E−03 | 1.44E−04 | 1.33E−03 | 12 | 102 |
| GO: Biological Process | GO:0001655 | urogenital system development | 1.37E−06 | 7.95E−03 | 1.47E−04 | 1.36E−03 | 25 | 406 |
| GO: Biological Process | GO:0010631 | epithelial cell migration | 1.73E−06 | 9.99E−03 | 1.82E−04 | 1.68E−03 | 24 | 384 |
| GO: Biological Process | GO:0071560 | cellular response to transforming growth factor beta stimulus | 1.90E−06 | 1.10E−02 | 1.97E−04 | 1.82E−03 | 20 | 282 |
| GO: Biological Process | GO:0090132 | epithelium migration | 1.98E−06 | 1.15E−02 | 2.01E−04 | 1.86E−03 | 24 | 387 |
| GO: Biological Process | GO:0048871 | multicellular organismal homeostasis | 2.14E−06 | 1.24E−02 | 2.14E−04 | 1.98E−03 | 31 | 589 |
| GO: Biological Process | GO:0003179 | heart valve morphogenesis | 2.37E−06 | 1.37E−02 | 2.32E−04 | 2.15E−03 | 9 | 57 |
| GO: Biological Process | GO:0071634 | regulation of transforming growth factor beta production | 2.41E−06 | 1.39E−02 | 2.32E−04 | 2.15E−03 | 8 | 43 |
| GO: Biological Process | GO:0090130 | tissue migration | 2.47E−06 | 1.43E−02 | 2.35E−04 | 2.17E−03 | 24 | 392 |
| GO: Biological Process | GO:0007599 | hemostasis | 2.71E−06 | 1.57E−02 | 2.53E−04 | 2.34E−03 | 23 | 367 |
| GO: Biological Process | GO:0071559 | response to transforming growth factor beta | 2.76E−06 | 1.60E−02 | 2.54E−04 | 2.35E−03 | 20 | 289 |
| GO: Biological Process | GO:0003007 | heart morphogenesis | 3.23E−06 | 1.87E−02 | 2.93E−04 | 2.71E−03 | 20 | 292 |
| GO: Biological Process | GO:0071604 | transforming growth factor beta production | 3.45E−06 | 2.00E−02 | 3.08E−04 | 2.84E−03 | 8 | 45 |
| GO: Biological Process | GO:0098609 | cell−cell adhesion | 4.21E−06 | 2.44E−02 | 3.70E−04 | 3.42E−03 | 42 | 961 |
| GO: Biological Process | GO:0001667 | ameboidal−type cell migration | 4.38E−06 | 2.54E−02 | 3.79E−04 | 3.50E−03 | 28 | 520 |
| GO: Biological Process | GO:0001952 | regulation of cell−matrix adhesion | 5.52E−06 | 3.20E−02 | 4.70E−04 | 4.34E−03 | 13 | 137 |
| GO: Biological Process | GO:0071363 | cellular response to growth factor stimulus | 5.64E−06 | 3.27E−02 | 4.73E−04 | 4.38E−03 | 36 | 775 |
| GO: Biological Process | GO:0061448 | connective tissue development | 6.53E−06 | 3.79E−02 | 5.41E−04 | 5.00E−03 | 20 | 306 |
| GO: Biological Process | GO:0048699 | generation of neurons | 6.76E−06 | 3.92E−02 | 5.52E−04 | 5.10E−03 | 65 | 1802 |
| GO: Biological Process | GO:0008284 | positive regulation of cell population proliferation | 6.88E−06 | 3.98E−02 | 5.53E−04 | 5.11E−03 | 46 | 1117 |
| GO: Biological Process | GO:0007596 | blood coagulation | 7.29E−06 | 4.23E−02 | 5.79E−04 | 5.35E−03 | 22 | 362 |
| GO: Biological Process | GO:0007162 | negative regulation of cell adhesion | 8.04E−06 | 4.65E−02 | 6.29E−04 | 5.81E−03 | 21 | 337 |
| GO: Biological Process | GO:0022603 | regulation of anatomical structure morphogenesis | 8.31E−06 | 4.81E−02 | 6.42E−04 | 5.93E−03 | 47 | 1160 |
| GO: Cellular Component | GO:0009986 | cell surface | 2.93E−09 | 1.79E−06 | 1.79E−06 | 1.25E−05 | 53 | 1053 |
| GO: Cellular Component | GO:0070161 | anchoring junction | 8.74E−08 | 5.35E−05 | 1.26E−05 | 8.84E−05 | 44 | 880 |
| GO: Cellular Component | GO:0005925 | focal adhesion | 9.57E−08 | 5.85E−05 | 1.26E−05 | 8.84E−05 | 28 | 424 |
| GO: Cellular Component | GO:0097478 | leaflet of membrane bilayer | 1.03E−07 | 6.32E−05 | 1.26E−05 | 8.84E−05 | 38 | 704 |
| GO: Cellular Component | GO:0098552 | side of membrane | 1.03E−07 | 6.32E−05 | 1.26E−05 | 8.84E−05 | 38 | 704 |
| GO: Cellular Component | GO:0030055 | cell−substrate junction | 1.62E−07 | 9.94E−05 | 1.66E−05 | 1.16E−04 | 28 | 435 |
| GO: Cellular Component | GO:0015629 | actin cytoskeleton | 1.10E−06 | 6.74E−04 | 8.97E−05 | 6.28E−04 | 40 | 837 |
| GO: Cellular Component | GO:0009897 | external side of plasma membrane | 1.17E−06 | 7.18E−04 | 8.97E−05 | 6.28E−04 | 28 | 480 |
| GO: Cellular Component | GO:0005887 | integral component of plasma membrane | 1.13E−05 | 6.91E−03 | 7.31E−04 | 5.12E−03 | 63 | 1735 |
| GO: Cellular Component | GO:0098636 | protein complex involved in cell adhesion | 1.20E−05 | 7.36E−03 | 7.31E−04 | 5.12E−03 | 7 | 38 |
| GO: Cellular Component | GO:0031226 | intrinsic component of plasma membrane | 1.31E−05 | 8.04E−03 | 7.31E−04 | 5.12E−03 | 65 | 1818 |
| GO: Cellular Component | GO:0031252 | cell leading edge | 2.06E−05 | 1.26E−02 | 9.75E−04 | 6.82E−03 | 25 | 469 |
| GO: Cellular Component | GO:0045121 | membrane raft | 2.15E−05 | 1.31E−02 | 9.75E−04 | 6.82E−03 | 23 | 412 |
| GO: Cellular Component | GO:0098857 | membrane microdomain | 2.23E−05 | 1.36E−02 | 9.75E−04 | 6.82E−03 | 23 | 413 |
| GO: Cellular Component | GO:0005788 | endoplasmic reticulum lumen | 5.54E−05 | 3.39E−02 | 2.12E−03 | 1.49E−02 | 19 | 323 |
| GO: Cellular Component | GO:0008305 | integrin complex | 5.55E−05 | 3.40E−02 | 2.12E−03 | 1.49E−02 | 6 | 33 |

HIF1A,TRAF6,PIK3CA,HPGD,VIP,PDGFD,PRKAR1A,ITGA2,PTEN,PPARGC1A,BMP4,JUN,TBX20,CAV1,S1PR1,ZFPM2,TGFBR2,THBS1,NAMPT,CALCRL,CDH13

PRTG,PDE4D,MOSPD2,ENPP2,SLIT2,PIK3CA,PIK3CG,VEGFC,ROBO4,ISL2,APP,SLC12A2,PDGFD,PALLD,CXCL8,DOCK4,IL16,MYCBP2,GATA3,ITGA1,ITGA2,KLRC4–KLRK1,ITGAV,BMP4,CXCR4,NFIB,GLI3,CXCL1,CXCL2,S1PR1,ABLIM1,THBS1,EFNA5,TMSB4X,CDH13,EPHA4

SLIT2,HPGD,JAG1,ARID5B,AHR,CC2D2A,CEP290,ITGA8,TIPARP,PDGFD,PYGO1,SERPINB7,GATA3,ACE,BMP4,CXCR4,GLI3,BMPER,FAT4,ADAMTS6,HEY1,CD34,EPHA4

PDE4D,GNG2,NREP,PHEX,SLIT2,FBN2,PIK3CA,PIK3CG,HPGD,AGL,PIP4K2A,AHR,SOX6,COL3A1,VIM,COL5A2,ITGA8,SESN3,APP,ANKRD1,PDGFD,NR5A2,PRKACB,PRKAR1A,PRKAR2B,LEPROT,SNCA,FUT8,CXCL8,INHBA,GATA3,ITGA2,PTEN,ITGB1,ITGB8,NEDD4,PPARGC1A,BMP4,JUN,T

FZD4,QKI,TIPARP,ITGAV,ITGB8,TBX20,RASA1,CAV1,ZFPM2,TGFBR2,HEY1,CD34

MECOM,SLIT2,HPGD,JAG1,ARID5B,AHR,CC2D2A,CEP290,ITGA8,TIPARP,PDGFD,PYGO1,SERPINB7,GATA3,PTEN,ACE,BMP4,CXCR4,GLI3,BMPER,FAT4,ADAMTS6,HEY1,CD34,EPHA4

HIF1A,ENPP2,ETS1,SLIT2,PIK3CA,PIK3CG,VEGFC,PLK2,GATA3,ITGA2,PTEN,ITGAV,ITGB1,GADD45A,ZEB2,BMP4,JUN,BMPER,AKT3,SASH1,TGFBR2,THBS1,TMSB4X,CDH13

NREP,FBN2,HPGD,SOX6,COL3A1,ITGA8,ANKRD1,PDGFD,FUT8,ITGB1,ITGB8,PPARGC1A,JUN,PTPRK,VEPH1,CAV1,CAV2,NR3C1,TGFBR2,THBS1

HIF1A,ENPP2,ETS1,SLIT2,PIK3CA,PIK3CG,VEGFC,PLK2,GATA3,ITGA2,PTEN,ITGAV,ITGB1,GADD45A,ZEB2,BMP4,JUN,BMPER,AKT3,SASH1,TGFBR2,THBS1,TMSB4X,CDH13

TRAF6,FABP4,FABP5,JAM2,PIK3CA,ADGRF5,PLCL1,ARRDC3,CEP290,SLC40A1,SLC12A2,PRKACB,PRKAR1A,PRKAR2B,VSIG1,ITGAV,ITGB1,IL18R1,PTHLH,PPARGC1A,CXCR4,PDGFC,AKT3,LAMA4,ESAM,CAV1,S1PR1,LEPR,LNPEP,CD34,CD36

ERG,SLIT2,JAG1,GATA3,BMP4,TBX20,ZFPM2,TGFBR2,HEY1

HIF1A,CD46,SERPINB7,ITGAV,ITGB8,THBS1,ATP6AP2,CD34

HIF1A,ENPP2,ETS1,SLIT2,PIK3CA,PIK3CG,VEGFC,PLK2,GATA3,ITGA2,PTEN,ITGAV,ITGB1,GADD45A,ZEB2,BMP4,JUN,BMPER,AKT3,SASH1,TGFBR2,THBS1,TMSB4X,CDH13

GNG2,CLEC1B,PIK3CA,PIK3CG,COL3A1,FLI1,ENPP4,MMRN1,PRKACB,PRKAR1A,PRKAR2B,GATA3,ITGA2,C9,CAPZA2,CAV1,PAPSS2,ZFPM2,TFPI,THBD,THBS1,CD34,CD36

NREP,FBN2,HPGD,SOX6,COL3A1,ITGA8,ANKRD1,PDGFD,FUT8,ITGB1,ITGB8,PPARGC1A,JUN,PTPRK,VEPH1,CAV1,CAV2,NR3C1,TGFBR2,THBS1

HIF1A,ERG,SLIT2,DLC1,JAG1,AHR,ANKRD1,GATA3,NEDD4,BMP4,JUN,TBX20,GNG5,FAT4,S1PR1,ZFPM2,ADGRG6,TGFBR2,THBS1,HEY1

HIF1A,CD46,SERPINB7,ITGAV,ITGB8,THBS1,ATP6AP2,CD34

PRTG,TRAF6,ETS1,HLX,CD46,JAM2,SLIT2,SERPINB8,PIK3CA,CD99,PIK3CG,JAG1,PCDH10,ROBO4,ITGA8,CD93,PRKAR1A,CD274,PALLD,PAG1,GATA3,CD55,KLRC4–KLRK1,ITGAV,ITGB1,BMP4,GLI3,BCL10,FAT4,LAMB1,ESAM,CAV1,CBLB,OLR1,IL1RAPL1,TGFBR2,THBS1,EFNA5,CD3

HIF1A,PRR5L,ENPP2,ETS1,SLIT2,KITLG,PIK3CA,PIK3CG,ARID5B,VEGFC,PLK2,GATA3,ITGA2,PTEN,ITGAV,ITGB1,GADD45A,ZEB2,BMP4,JUN,CXCR4,BMPER,AKT3,SASH1,TGFBR2,THBS1,TMSB4X,CDH13

CDKN2A,ENPP2,DLC1,JAG1,VEGFC,ARHGAP6,PTEN,RASA1,THBS1,EFNA5,CD36,CDH13,PHLDB2

HIF1A,NREP,FZD4,SLIT2,FBN2,PIK3CA,HPGD,VEGFC,SOX6,COL3A1,ITGA8,APP,ANKRD1,PDGFD,FUT8,CXCL8,GATA3,PTEN,ITGB1,ITGB8,NEDD4,PPARGC1A,BMP4,JUN,PTPRK,BMPER,TCF4,FAT4,RASA1,VEPH1,STMN2,CAV1,CAV2,NR3C1,TGFBR2,THBS1

HIF1A,EFEMP1,PIK3CA,ARID5B,ARRDC3,CREB3L2,SOX6,PDGFD,CSGALNACT1,SERPINB7,PTHLH,ITGB8,PPARGC1A,BMP4,NFIB,GLI3,TGFBR2,THBS1,NAMPT,CD34

PRTG,HIF1A,ENPP2,NREP,FZD4,SLIT2,EFEMP1,PIK3CA,HECW2,CLN5,JAG1,QKI,CREB3L2,VEGFC,COL3A1,VIM,ROBO4,CEP290,DOCK10,ZNF521,ISL2,APP,ANKRD1,TMEM30A,IL1R1,PALLD,TANC2,PLK2,MYCBP2,INHBA,MYO9A,SDCCAG8,GATA3,ITGA1,PTEN,ACE,ITGB1,ZEB2,NEDD4,

HIF1A,TRAF6,ENPP2,ETS1,HLX,CD46,FABP4,EFEMP1,KITLG,PIK3CA,HPGD,TNS3,VEGFC,VIM,VIP,PDGFD,CD274,SERPINB7,CD55,ITGA2,PTEN,ITGAV,ITGB1,PTHLH,PPARGC1A,BMP4,JUN,GLI3,TBX20,PDGFC,GNG5,AKT3,RASA1,SYNE1,LAMB1,ESM1,CAV1,CAV2,S1PR1,ZFPM2,TGFB

GNG2,CLEC1B,PIK3CA,PIK3CG,COL3A1,ENPP4,MMRN1,PRKACB,PRKAR1A,PRKAR2B,GATA3,ITGA2,C9,CAPZA2,CAV1,PAPSS2,ZFPM2,TFPI,THBD,THBS1,CD34,CD36

CDKN2A,ENPP2,HLX,JAM2,FZD4,DLC1,JAG1,ARHGAP6,PRKAR1A,CD274,PAG1,PTEN,BMP4,GLI3,RASA1,LAMB1,CBLB,THBS1,EFNA5,CDH13,PHLDB2

HIF1A,ERG,ENPP2,ETS1,FZD4,SLIT2,DLC1,HECW2,MLLT3,VEGFC,VIM,SLC12A2,DAAM1,ARHGAP18,CXCL8,TANC2,PLK2,ERAP1,MYO9A,GATA3,PALMD,PTEN,ITGB1,GADD45A,ZEB2,ITGB8,NEDD4,PPARGC1A,BMP4,RHOBTB1,CXCR4,BTG1,BMPER,TCF4,AKT3,SASH1,RASA1,SYNE1,IL1RA

CD46,SPA17,JAM2,FZD4,SLIT2,ADGRF5,MMP7,TMEM123,ROBO4,NID2,ITGA8,APP,CD93,CLEC2D,CD274,IL1R1,IL13RA2,LAYN,CD55,ITGA1,ITGA2,KLRC4–KLRK1,ACE,ITGAV,ITGB1,ITGB8,PTPRK,CXCR4,CKAP4,PDGFC,CLEC2B,CLEC12B,CLEC9A,CAV1,CAV2,S1PR1,LEPR,TFPI,ADG

CD46,ABCB4,JAM2,FZD4,DLC1,PIK3CA,CD99,JAG1,TNS3,VIM,PPFIBP1,ITGA8,APP,SHROOM2,PALLD,IL16,TES,LAYN,SDCCAG8,MPDZ,ITGA1,ITGA2,ITGAV,ITGB1,FRMD4B,ITGB8,DST,PTPRK,CXCR4,NHS,PDCD6IP,ESAM,PARD3B,CAV1,CAV2,ARPC5,EFNA5,FRMD6,LPP,IL1RL1,MARC

CD46,ABCB4,DLC1,CD99,TNS3,VIM,PPFIBP1,ITGA8,PALLD,IL16,TES,LAYN,ITGA1,ITGA2,ITGAV,ITGB1,ITGB8,DST,NHS,PDCD6IP,CAV1,CAV2,ARPC5,LPP,IL1RL1,MARCKS,CDH13,PHLDB2

TRAF6,GNG2,SPA17,TMEM123,ROBO4,BIRC2,MYZAP,CLEC2D,CD274,IL1R1,IL13RA2,ITGA1,ITGA2,KLRC4–KLRK1,PTEN,ACE,ITGAV,ITGB1,DST,GNG12,CXCR4,KCNAB1,GNG5,CLEC2B,CLEC12B,CAV2,S1PR1,LEPR,TGFBR2,THBD,THBS1,EFNA5,ATP6AP2,CD34,CD36,CD69,IL1RL1,CDH

TRAF6,GNG2,SPA17,TMEM123,ROBO4,BIRC2,MYZAP,CLEC2D,CD274,IL1R1,IL13RA2,ITGA1,ITGA2,KLRC4–KLRK1,PTEN,ACE,ITGAV,ITGB1,DST,GNG12,CXCR4,KCNAB1,GNG5,CLEC2B,CLEC12B,CAV2,S1PR1,LEPR,TGFBR2,THBD,THBS1,EFNA5,ATP6AP2,CD34,CD36,CD69,IL1RL1,CDH

CD46,ABCB4,DLC1,CD99,TNS3,VIM,PPFIBP1,ITGA8,PALLD,IL16,TES,LAYN,ITGA1,ITGA2,ITGAV,ITGB1,ITGB8,DST,NHS,PDCD6IP,CAV1,CAV2,ARPC5,LPP,IL1RL1,MARCKS,CDH13,PHLDB2

SLC26A4,ABCB4,DLC1,KITLG,KLHL14,PLS3,ARHGAP21,MYZAP,APP,SHROOM2,ARHGAP6,CD274,MYH15,DAAM1,SNCA,PALLD,DOCK4,FILIP1,MYO9A,ACTR10,RGS22,IFIT5,ACE,ITGAV,ITGB1,NEDD4,DST,GNG12,ARSJ,PDCD6IP,CAPZA2,ARPC5,TLNRD1,LMOD3,ABLIM1,CD36,LPP,SEPTI

SPA17,TMEM123,ROBO4,CLEC2D,CD274,IL1R1,IL13RA2,ITGA1,ITGA2,KLRC4–KLRK1,ACE,ITGAV,ITGB1,CXCR4,CLEC2B,CLEC12B,S1PR1,LEPR,TGFBR2,THBD,THBS1,EFNA5,ATP6AP2,CD34,CD36,CD69,IL1RL1,CDH13

SGMS2,TRAF6,MOSPD2,VSIG2,ENPP2,SLC26A4,CLEC1B,CLEC1A,CD46,ABCB4,JAM2,FZD4,PHEX,CLCN4,CD99,RASGRP3,JAG1,OPN5,MIGA1,PCDH10,ITGA8,BIRC2,APP,SLC40A1,SLC12A2,CLEC2D,TSPAN13,IL1R1,SCN3B,PRRG1,ITGA1,ITGA2,KLRC4–KLRK1,ITGAV,ITGB1,IL18R1,IT

JAM2,ITGA8,ITGA1,ITGA2,ITGAV,ITGB1,ITGB8

SGMS2,TRAF6,MOSPD2,VSIG2,ENPP2,SLC26A4,CLEC1B,CLEC1A,CD46,ABCB4,JAM2,FZD4,PHEX,CLCN4,CD99,RASGRP3,JAG1,OPN5,MIGA1,PCDH10,ITGA8,BIRC2,APP,SLC40A1,SLC12A2,CLEC2D,TSPAN13,IL1R1,SCN3B,PRRG1,ITGA1,ITGA2,KLRC4–KLRK1,ITGAV,ITGB1,IL18R1,IT

DLC1,KITLG,PIK3CA,VIM,ITGA8,APP,ARHGAP18,PALLD,LAYN,IFIT5,ITGAV,ITGB1,FRMD4B,BMX,DST,PTPRK,CXCR4,NHS,SGCE,RASA1,STMN2,ARPC5,ABLIM1,ATP6AP2,PHLDB2

ABCB4,DLC1,CAVIN2,BIRC2,APP,PRKAR1A,PRKAR2B,PAG1,CD55,ITGA1,ITGB1,NEDD4,BCL10,CAV1,CAV2,CBLB,S1PR1,OLR1,TFPI,TGFBR2,EFNA5,CD36,CDH13

ABCB4,DLC1,CAVIN2,BIRC2,APP,PRKAR1A,PRKAR2B,PAG1,CD55,ITGA1,ITGB1,NEDD4,BCL10,CAV1,CAV2,CBLB,S1PR1,OLR1,TFPI,TGFBR2,EFNA5,CD36,CDH13

PCSK1,COL3A1,COL4A5,COL5A2,APP,ARSG,EDEM3,STS,ARSD,PDGFD,ERAP1,BMP4,ERAP2,CKAP4,PDGFC,ARSJ,LAMB1,PRSS23,THBS1

ITGA8,ITGA1,ITGA2,ITGAV,ITGB1,ITGB8

| Category | ID | Name | p-value | q-value Bonferroni | q-value FDR B&H | q-value FDR B&Y | Hit Count in Query List | Hit Count in Genome |
|---|---|---|---|---|---|---|---|---|
| GO: Molecular Function | GO:0022836 | gated channel activity | 2.75E-07 | 3.07E-04 | 3.07E-04 | 2.33E-03 | 30 | 352 |
| GO: Molecular Function | GO:0015267 | channel activity | 4.28E-06 | 4.77E-03 | 1.65E-03 | 1.26E-02 | 39 | 597 |
| GO: Molecular Function | GO:0022803 | passive transmembrane transporter activity | 4.45E-06 | 4.96E-03 | 1.65E-03 | 1.26E-02 | 39 | 598 |
| GO: Molecular Function | GO:0005216 | ion channel activity | 5.93E-06 | 6.61E-03 | 1.65E-03 | 1.26E-02 | 35 | 516 |
| GO: Cellular Component | GO:0034702 | ion channel complex | 4.04E-07 | 2.91E-04 | 2.91E-04 | 2.08E-03 | 28 | 319 |
| GO: Cellular Component | GO:0099699 | integral component of synaptic membrane | 1.19E-06 | 8.57E-04 | 2.92E-04 | 2.09E-03 | 22 | 225 |
| GO: Cellular Component | GO:1990351 | transporter complex | 1.24E-06 | 8.94E-04 | 2.92E-04 | 2.09E-03 | 29 | 357 |
| GO: Cellular Component | GO:1902495 | transmembrane transporter complex | 1.62E-06 | 1.17E-03 | 2.92E-04 | 2.09E-03 | 28 | 342 |
| GO: Cellular Component | GO:0099055 | integral component of postsynaptic membrane | 3.31E-06 | 2.38E-03 | 4.08E-04 | 2.92E-03 | 18 | 169 |
| GO: Cellular Component | GO:0097060 | synaptic membrane | 3.40E-06 | 2.45E-03 | 4.08E-04 | 2.92E-03 | 34 | 480 |
| GO: Cellular Component | GO:0099240 | intrinsic component of synaptic membrane | 4.54E-06 | 3.27E-03 | 4.67E-04 | 3.35E-03 | 22 | 244 |
| GO: Cellular Component | GO:0098936 | intrinsic component of postsynaptic membrane | 6.89E-06 | 4.96E-03 | 6.20E-04 | 4.44E-03 | 18 | 178 |
| GO: Cellular Component | GO:0045211 | postsynaptic membrane | 9.93E-06 | 7.15E-03 | 7.94E-04 | 5.69E-03 | 26 | 335 |
| GO: Cellular Component | GO:0045202 | synapse | 2.60E-05 | 1.87E-02 | 1.87E-03 | 1.34E-02 | 76 | 1582 |
| GO: Cellular Component | GO:0034703 | cation channel complex | 3.61E-05 | 2.60E-02 | 2.36E-03 | 1.69E-02 | 20 | 239 |
| GO: Cellular Component | GO:0098878 | neurotransmitter receptor complex | 4.50E-05 | 3.24E-02 | 2.70E-03 | 1.93E-02 | 9 | 57 |
| GO: Cellular Component | GO:0005887 | integral component of plasma membrane | 6.42E-05 | 4.62E-02 | 3.34E-03 | 2.39E-02 | 80 | 1735 |
| GO: Cellular Component | GO:0031226 | intrinsic component of plasma membrane | 6.49E-05 | 4.67E-02 | 3.34E-03 | 2.39E-02 | 83 | 1818 |

| Hit in Query List |
|---|
| TTYH1,KCNIP3,SCN5A,SCNN1G,ASIC4,TMC4,NALF2,CACNA2D3,GABRA3,GABRA5,GABRB3,GABRQ,CACNA1E,GRID2,GRIK3,GRIN2B,P2RX1,P2RX5,TSPOAP1,KCNB2,CLCN1,CNR1,HTR3A,KCNA1,KCNA2,KCNA3,KCNC3,KCNS1,TTYH2,TMC5 |
| TTYH1,GJD2,KCNIP3,SCN5A,SCNN1G,TRPM1,ASIC4,TMC4,AQP4,NALF2,CACNA2D3,GABRA3,GABRA5,GABRB3,GABRQ,NOS1,NPY,CACNA1E,GRID2,GRIK3,GRIN2B,P2RX1,P2RX5,TSPOAP1,KCNB2,SLC9A3R1,CLCN1,PKP2,CNR1,HTR3A,PRKCB,PRSS8,KCNA1,KCNA2 |
| TTYH1,GJD2,KCNIP3,SCN5A,SCNN1G,TRPM1,ASIC4,TMC4,AQP4,NALF2,CACNA2D3,GABRA3,GABRA5,GABRB3,GABRQ,NOS1,NPY,CACNA1E,GRID2,GRIK3,GRIN2B,P2RX1,P2RX5,TSPOAP1,KCNB2,SLC9A3R1,CLCN1,PKP2,CNR1,HTR3A,PRKCB,PRSS8,KCNA1,KCNA2 |
| TTYH1,KCNIP3,SCN5A,SCNN1G,TRPM1,ASIC4,TMC4,NALF2,CACNA2D3,GABRA3,GABRA5,GABRB3,GABRQ,NOS1,CACNA1E,GRID2,GRIK3,GRIN2B,P2RX1,P2RX5,TSPOAP1,KCNB2,SLC9A3R1,CLCN1,PKP2,CNR1,HTR3A,PRSS8,KCNA1,KCNA2,KCNA3,KCNC3,KCNS1,T |
| TTYH1,KCNIP3,SCN5A,SCNN1G,NLGN1,CACNA2D3,GABRA3,GABRA5,GABRB3,GABRQ,NOS1,CACNA1E,GRID2,GRIK3,GRIN2B,SHISA9,OLFM2,KCNB2,DLGAP3,CLCN1,HTR3A,KCNA1,KCNA2,KCNA3,KCNC3,KCNS1,DLG3,TTYH2 |
| LRFN2,NLGN1,GABBR1,GABRA3,GABRA5,GABRB3,SLITRK2,CACNA1E,SYT6,GRID2,GRIN2B,SHISA9,P2RX1,LPAR2,GSG1L,PCDH8,CNR1,HTR3A,KCNA1,KCNA2,KCNA3,KCNC3 |
| TTYH1,KCNIP3,SCN5A,SCNN1G,NLGN1,CACNA2D3,GABRA3,GABRA5,GABRB3,GABRQ,NOS1,CACNA1E,GRID2,GRIK3,GRIN2B,SHISA9,OLFM2,KCNB2,DLGAP3,CLCN1,HTR3A,KCNA1,KCNA2,KCNA3,KCNC3,KCNS1,DLG3,TTYH2,ATP10A |
| TTYH1,KCNIP3,SCN5A,SCNN1G,NLGN1,CACNA2D3,GABRA3,GABRA5,GABRB3,GABRQ,NOS1,CACNA1E,GRID2,GRIK3,GRIN2B,SHISA9,OLFM2,KCNB2,DLGAP3,CLCN1,HTR3A,KCNA1,KCNA2,KCNA3,KCNC3,KCNS1,DLG3,TTYH2 |
| LRFN2,NLGN1,GABBR1,GABRA3,GABRA5,GABRB3,SLITRK2,CACNA1E,GRID2,GRIN2B,SHISA9,P2RX1,GSG1L,PCDH8,HTR3A,KCNA1,KCNA3,KCNC3 |
| LRFN2,NLGN1,MAGEE1,IGSF9B,GABBR1,GABRA3,GABRA5,GABRB3,GABRQ,SLITRK2,ARC,CACNA1E,CALB2,SYT6,GRID2,GRIK3,GRIN2B,PTPN5,GRIP1,SHISA9,P2RX1,LPAR2,OLFM2,GSG1L,IGSF21,PCDH8,CNR1,HTR3A,CADPS2,RIMS2,KCNA1,KCNA2,KCNA3,KCN |
| LRFN2,NLGN1,GABBR1,GABRA3,GABRA5,GABRB3,SLITRK2,CACNA1E,SYT6,GRID2,GRIN2B,SHISA9,P2RX1,LPAR2,GSG1L,PCDH8,CNR1,HTR3A,KCNA1,KCNA2,KCNA3,KCNC3 |
| LRFN2,NLGN1,GABBR1,GABRA3,GABRA5,GABRB3,SLITRK2,CACNA1E,GRID2,GRIN2B,SHISA9,P2RX1,GSG1L,PCDH8,HTR3A,KCNA1,KCNA3,KCNC3 |
| LRFN2,NLGN1,MAGEE1,IGSF9B,GABBR1,GABRA3,GABRA5,GABRB3,GABRQ,SLITRK2,ARC,CACNA1E,GRID2,GRIK3,GRIN2B,GRIP1,SHISA9,P2RX1,GSG1L,IGSF21,PCDH8,HTR3A,CADPS2,KCNA1,KCNA3,KCNC3 |
| TTYH1,RPS4X,ACTC1,RPS17,KCNIP3,ADORA2B,LRFN2,ACAN,FGFR2,NLGN1,MAGEE1,SLC1A3,CADPS,SLC22A3,SLC22A2,ENAH,IGSF9B,FUS,GABBR1,GABRA3,GABRA5,GABRB3,GABRQ,PGRMC1,SLITRK2,ARHGEF9,ARC,NOS1,SDK1,NPY,CACNA1E,NPY1R,CALB2,SY |
| KCNIP3,SCN5A,SCNN1G,NLGN1,CACNA2D3,NOS1,CACNA1E,GRIK3,GRIN2B,SHISA9,OLFM2,KCNB2,DLGAP3,HTR3A,KCNA1,KCNA2,KCNA3,KCNC3,KCNS1,DLG3 |
| NLGN1,GRID2,GRIK3,GRIN2B,SHISA9,OLFM2,DLGAP3,HTR3A,DLG3 |
| RAMP3,KCNIP3,ADCY2,ADORA2B,LRFN2,SCN5A,SCNN1G,TRPM1,FGFR2,ASIC4,SDC4,IL17RB,TMC4,NLGN1,AQP4,SLC1A3,SLC4A3,TENM3,SLC15A2,SLC22A3,SLC22A2,GABBR1,GABRA3,GABRA5,GABRB3,GALR1,GABRQ,GPRC5C,PGRMC1,SLC52A3,SLITRK2,AJAP1 |
| RAMP3,KCNIP3,ADCY2,ADORA2B,LRFN2,SCN5A,SCNN1G,TRPM1,FGFR2,ASIC4,SDC4,IL17RB,TMC4,NLGN1,AQP4,SLC1A3,SLC4A3,TENM3,SLC15A2,SLC22A3,SLC22A2,GABBR1,GABRA3,GABRA5,GABRB3,GALR1,GABRQ,GPRC5C,PGRMC1,SLC52A3,SLITRK2,AJAP1 |

| | hPSC | MES | EP | EC |
|---|---|---|---|---|
| hPSC | 1 | 0.698 | 0.605 | 0.551 |
| MES | 0.682 | 1 | 0.68 | 0.589 |
| EP | 0.599 | 0.691 | 1 | 0.777 |
| EC | 0.549 | 0.601 | 0.781 | 1 |

| Category | ID | Name | p-value | q-value Bonferroni | q-value FDR B&H | q-value FDR B&Y | Hit Count in Query List | Hit Count in Genome |
|---|---|---|---|---|---|---|---|---|
| GO: Biological Process | GO:0050748 | negative regulation of lipoprotein metabolic process | 5.28E-06 | 1.80E-02 | 1.80E-02 | 1.57E-01 | 3 | 5 |
| GO: Biological Process | GO:0007010 | cytoskeleton organization | 1.11E-05 | 3.78E-02 | 1.89E-02 | 1.65E-01 | 32 | 1745 |
| GO: Cellular Component | GO:0098636 | protein complex involved in cell adhesion | 1.36E-05 | 5.65E-03 | 5.65E-03 | 3.74E-02 | 5 | 38 |

| Hit in Query List |
| --- |
| ITGAV,ITGB3,APOD |
| CORO1C,BBS2,TOGARAM1,SYNPO,KATNAL1,ITGB3,GADD45A,FSIP2,TUBB6,PHLDB1,CFAP54,CDC42EP2,GOLGA2,TCTN1,PMP22,RALA,TBC1D30,ANXA1,GOLGA8R,CLIC4,CAVIN3,ZBED3,ARF1,PDLIM4,PLEK2,FAM171A1,KIFAP3,AKAP9,DYNC2LI1,PRKACA,PRKCE,DMTN |
| ITGAV,ITGB3,JAM2,PMP22,ITGA11 |

| Category | ID | Name | p-value | q-value Bonferroni | q-value FDR B&H | q-value FDR B&Y | Hit Count in Query List | Hit Count in Genome | Hit in Query List |
|---|---|---|---|---|---|---|---|---|---|
| GO: Biological Process | GO:0071103 | DNA conformation change | 1.47E-06 | 5.48E-03 | 5.48E-03 | 4.82E-02 | 17 | 363 | MCM6,HMGA1,H2BC4,DNA2,H2BC12,NPM1,H2BC11,H4C15,RFC4,RUVBL1,G3BP1,H1-4,H1-6,SMARCC1,H2BC5,CDK1,XRCC6 |
| GO: Biological Process | GO:0006342 | chromatin silencing | 5.87E-06 | 2.19E-02 | 1.09E-02 | 9.62E-02 | 8 | 83 | HMGA1,H2AC6,H2AC4,H2AC12,ATAD2,DYDC2,H1-4,H1-6 |

| set_loops | compartment–A–A_DEG–up | compartment–A–A_DEG–down | compartment–A–B_DEG–up | compartment–A–B_DEG–down | compartment–B–B_DEG–up | compartment–B–B_DEG–down |
|---|---|---|---|---|---|---|
| hPSC | 105 | 198 | 27 | 27 | 90 | 70 |
| MES | 200 | 318 | 56 | 49 | 178 | 194 |
| EP | 506 | 599 | 130 | 145 | 191 | 305 |
| EC | 867 | 1012 | 178 | 232 | 185 | 403 |

| Category | ID | Name | p-value | q-value Bonferroni | q-value FDR B&H | q-value FDR B&Y | Hit Count in Query List | Hit Count in Genome |
|---|---|---|---|---|---|---|---|---|
| GO: Molecular Function | GO:0003779 | actin binding | 3.44E-08 | 3.92E-05 | 3.92E-05 | 2.99E-04 | 42 | 456 |
| GO: Molecular Function | GO:0008092 | cytoskeletal protein binding | 1.68E-07 | 1.91E-04 | 9.55E-05 | 7.27E-04 | 72 | 1053 |
| GO: Molecular Function | GO:0001228 | DNA-binding transcription activator activity, RNA polymerase II-specific | 4.83E-07 | 5.51E-04 | 1.69E-04 | 1.29E-03 | 43 | 520 |
| GO: Molecular Function | GO:0001216 | DNA-binding transcription activator activity | 5.94E-07 | 6.77E-04 | 1.69E-04 | 1.29E-03 | 43 | 524 |
| GO: Molecular Function | GO:0050839 | cell adhesion molecule binding | 2.60E-06 | 2.96E-03 | 5.20E-04 | 3.96E-03 | 44 | 573 |
| GO: Molecular Function | GO:0030234 | enzyme regulator activity | 3.08E-06 | 3.51E-03 | 5.20E-04 | 3.96E-03 | 79 | 1288 |
| GO: Molecular Function | GO:0019838 | growth factor binding | 3.19E-06 | 3.64E-03 | 5.20E-04 | 3.96E-03 | 20 | 168 |
| GO: Molecular Function | GO:0005520 | insulin-like growth factor binding | 6.33E-06 | 7.22E-03 | 9.03E-04 | 6.87E-03 | 8 | 29 |
| GO: Molecular Function | GO:0016773 | phosphotransferase activity, alcohol group as acceptor | 7.32E-06 | 8.34E-03 | 9.27E-04 | 7.06E-03 | 58 | 875 |
| GO: Molecular Function | GO:0008134 | transcription factor binding | 9.26E-06 | 1.06E-02 | 1.06E-03 | 8.04E-03 | 51 | 740 |
| GO: Molecular Function | GO:0008047 | enzyme activator activity | 1.11E-05 | 1.26E-02 | 1.15E-03 | 8.73E-03 | 42 | 568 |
| GO: Molecular Function | GO:0046332 | SMAD binding | 1.47E-05 | 1.68E-02 | 1.40E-03 | 1.06E-02 | 13 | 87 |
| GO: Molecular Function | GO:0005178 | integrin binding | 1.65E-05 | 1.89E-02 | 1.45E-03 | 1.10E-02 | 18 | 157 |
| GO: Molecular Function | GO:0004672 | protein kinase activity | 1.84E-05 | 2.10E-02 | 1.46E-03 | 1.11E-02 | 51 | 759 |
| GO: Molecular Function | GO:0019904 | protein domain specific binding | 1.92E-05 | 2.19E-02 | 1.46E-03 | 1.11E-02 | 55 | 842 |
| GO: Molecular Function | GO:0016301 | kinase activity | 3.05E-05 | 3.48E-02 | 2.18E-03 | 1.66E-02 | 59 | 940 |
| GO: Biological Process | GO:0072359 | circulatory system development | 6.93E-27 | 4.87E-23 | 4.80E-23 | 4.53E-22 | 128 | 1340 |
| GO: Biological Process | GO:0001568 | blood vessel development | 1.36E-26 | 9.59E-23 | 4.80E-23 | 4.53E-22 | 99 | 866 |
| GO: Biological Process | GO:0035239 | tube morphogenesis | 2.83E-26 | 1.99E-22 | 5.82E-23 | 5.49E-22 | 111 | 1068 |
| GO: Biological Process | GO:0048514 | blood vessel morphogenesis | 3.31E-26 | 2.33E-22 | 5.82E-23 | 5.49E-22 | 92 | 768 |
| GO: Biological Process | GO:0001944 | vasculature development | 8.95E-26 | 6.29E-22 | 1.26E-22 | 1.19E-21 | 100 | 903 |
| GO: Biological Process | GO:0048646 | anatomical structure formation involved in morphogenesis | 3.12E-24 | 2.19E-20 | 3.65E-21 | 3.45E-20 | 126 | 1395 |
| GO: Biological Process | GO:0035295 | tube development | 4.60E-23 | 3.23E-19 | 4.62E-20 | 4.36E-19 | 119 | 1310 |
| GO: Biological Process | GO:0001525 | angiogenesis | 1.29E-22 | 9.10E-19 | 1.14E-19 | 1.07E-18 | 79 | 658 |
| GO: Biological Process | GO:0016477 | cell migration | 2.08E-18 | 1.46E-14 | 1.62E-15 | 1.53E-14 | 135 | 1812 |
| GO: Biological Process | GO:0009887 | animal organ morphogenesis | 5.31E-18 | 3.73E-14 | 3.73E-15 | 3.52E-14 | 106 | 1263 |
| GO: Biological Process | GO:0000902 | cell morphogenesis | 8.71E-18 | 6.13E-14 | 5.57E-15 | 5.26E-14 | 102 | 1197 |
| GO: Biological Process | GO:2000145 | regulation of cell motility | 2.42E-17 | 1.70E-13 | 1.42E-14 | 1.34E-13 | 99 | 1159 |
| GO: Biological Process | GO:0051270 | regulation of cellular component movement | 6.01E-17 | 4.23E-13 | 3.25E-14 | 3.07E-13 | 103 | 1250 |
| GO: Biological Process | GO:0030334 | regulation of cell migration | 8.90E-17 | 6.26E-13 | 4.47E-14 | 4.22E-13 | 94 | 1089 |
| GO: Biological Process | GO:0001667 | ameboidal-type cell migration | 1.51E-16 | 1.06E-12 | 6.62E-14 | 6.24E-13 | 60 | 520 |
| GO: Biological Process | GO:0040012 | regulation of locomotion | 1.51E-16 | 1.06E-12 | 6.62E-14 | 6.24E-13 | 100 | 1210 |
| GO: Biological Process | GO:0022603 | regulation of anatomical structure morphogenesis | 2.15E-16 | 1.51E-12 | 8.89E-14 | 8.39E-13 | 97 | 1160 |
| GO: Biological Process | GO:0090130 | tissue migration | 2.30E-16 | 1.61E-12 | 8.97E-14 | 8.46E-13 | 51 | 392 |
| GO: Biological Process | GO:0010631 | epithelial cell migration | 4.42E-16 | 3.11E-12 | 1.63E-13 | 1.54E-12 | 50 | 384 |
| GO: Biological Process | GO:0090132 | epithelium migration | 6.08E-16 | 4.28E-12 | 2.14E-13 | 2.02E-12 | 50 | 387 |
| GO: Biological Process | GO:2000026 | regulation of multicellular organismal development | 1.65E-15 | 1.16E-11 | 5.52E-13 | 5.21E-12 | 119 | 1632 |
| GO: Biological Process | GO:0045597 | positive regulation of cell differentiation | 2.91E-15 | 2.05E-11 | 9.31E-13 | 8.78E-12 | 87 | 1020 |
| GO: Biological Process | GO:0009611 | response to wounding | 5.99E-15 | 4.21E-11 | 1.83E-12 | 1.73E-11 | 71 | 746 |
| GO: Biological Process | GO:0007167 | enzyme linked receptor protein signaling pathway | 7.21E-15 | 5.07E-11 | 2.11E-12 | 1.99E-11 | 94 | 1168 |
| GO: Biological Process | GO:0051094 | positive regulation of developmental process | 2.00E-14 | 1.41E-10 | 5.63E-12 | 5.31E-11 | 111 | 1525 |
| GO: Biological Process | GO:0045595 | regulation of cell differentiation | 4.95E-14 | 3.48E-10 | 1.34E-11 | 1.26E-10 | 126 | 1859 |
| GO: Biological Process | GO:0010632 | regulation of epithelial cell migration | 1.54E-13 | 1.08E-09 | 4.01E-11 | 3.78E-10 | 41 | 312 |
| GO: Biological Process | GO:0030029 | actin filament-based process | 2.85E-13 | 2.01E-09 | 7.01E-11 | 6.61E-10 | 75 | 879 |
| GO: Biological Process | GO:0051240 | positive regulation of multicellular organismal process | 2.89E-13 | 2.03E-09 | 7.01E-11 | 6.61E-10 | 116 | 1692 |
| GO: Biological Process | GO:0000904 | cell morphogenesis involved in differentiation | 5.03E-13 | 3.54E-09 | 1.18E-10 | 1.11E-09 | 75 | 889 |
| GO: Biological Process | GO:0043542 | endothelial cell migration | 5.61E-13 | 3.94E-09 | 1.27E-10 | 1.20E-09 | 39 | 296 |
| GO: Biological Process | GO:0009790 | embryo development | 1.35E-12 | 9.52E-09 | 2.98E-10 | 2.81E-09 | 96 | 1316 |
| GO: Biological Process | GO:0048729 | tissue morphogenesis | 1.72E-12 | 1.21E-08 | 3.67E-10 | 3.46E-09 | 73 | 874 |
| GO: Biological Process | GO:0007507 | heart development | 2.36E-12 | 1.66E-08 | 4.88E-10 | 4.61E-09 | 63 | 698 |
| GO: Biological Process | GO:0010634 | positive regulation of epithelial cell migration | 2.68E-12 | 1.88E-08 | 5.38E-10 | 5.08E-09 | 30 | 190 |
| GO: Biological Process | GO:0051254 | positive regulation of RNA metabolic process | 3.61E-12 | 2.54E-08 | 7.04E-10 | 6.64E-09 | 120 | 1844 |
| GO: Biological Process | GO:0072001 | renal system development | 4.31E-12 | 3.03E-08 | 8.04E-10 | 7.59E-09 | 42 | 360 |
| GO: Biological Process | GO:0001655 | urogenital system development | 4.35E-12 | 3.06E-08 | 8.04E-10 | 7.59E-09 | 45 | 406 |
| GO: Biological Process | GO:0022008 | neurogenesis | 4.93E-12 | 3.47E-08 | 8.78E-10 | 8.28E-09 | 124 | 1940 |
| GO: Biological Process | GO:0007010 | cytoskeleton organization | 4.99E-12 | 3.51E-08 | 8.78E-10 | 8.28E-09 | 115 | 1745 |
| GO: Biological Process | GO:0030182 | neuron differentiation | 5.16E-12 | 3.63E-08 | 8.84E-10 | 8.35E-09 | 110 | 1639 |
| GO: Biological Process | GO:0022610 | biological adhesion | 5.68E-12 | 4.00E-08 | 9.47E-10 | 8.93E-09 | 107 | 1578 |
| GO: Biological Process | GO:0003158 | endothelium development | 5.79E-12 | 4.07E-08 | 9.47E-10 | 8.93E-09 | 27 | 159 |
| GO: Biological Process | GO:0040017 | positive regulation of locomotion | 6.16E-12 | 4.33E-08 | 9.72E-10 | 9.17E-09 | 61 | 678 |
| GO: Biological Process | GO:0030036 | actin cytoskeleton organization | 6.23E-12 | 4.38E-08 | 9.72E-10 | 9.17E-09 | 66 | 768 |
| GO: Biological Process | GO:0070848 | response to growth factor | 6.36E-12 | 4.47E-08 | 9.72E-10 | 9.17E-09 | 68 | 805 |
| GO: Biological Process | GO:0003170 | heart valve development | 6.70E-12 | 4.71E-08 | 1.00E-09 | 9.45E-09 | 18 | 67 |

| Hit in Query List |
|---|
| ABL2,RCSD1,FMNL2,MRTFB,EVL,FLNB,COBLL1,VASH1,MYO1B,PDLIM5,SMTN,MSN,LIMCH1,FSCN1,PALLD,SNTB2,PHACTR1,MYH9,MYO9B,ITPRID2,IQGAP1,DST,MYPN,COTL1,GSN,FMNL3,AFAP1,TMOD2,PARVB,VPS16,GMFG,SYNE3,PRKCE,PHACTR2,MTSS1,SPIRE1,PXK,CXCR4,ACTR2,ABLIM1,MARCKS,EPB41L2 |
| ABL2,RCSD1,FMNL2,PICALM,MRTFB,DLC1,MTUS1,FES,EVL,FLNB,SORBS2,VEZT,SGK1,COBLL1,VASH1,MYO1B,RAB14,CDC42EP3,PDLIM5,SMTN,MSN,TTLL11,LIMCH1,TPPP3,FSCN1,PALLD,SNTB2,PHACTR1,MYH9,MYO9B,DISC1,RAB11A,ITPRID2,IQGAP1,STIM1,DST,NFKB1,MYPN,DCLK2,CAMSAP2,COTL1,GSN,FMNL3,AFAP1 |
| ETS1,MECOM,MEF2D,ZNF319,EBF4,FOXO1,ATF6,NR5A2,PRDM16,GATA2,GATA3,GATA4,STAT3,STAT5A,TSC22D1,NFATC2,NFE2L2,NFIC,NFKB1,NHLH2,TCF4,SOX17,RUNX2,HHEX,HIVEP2,HOXA1,HOXA5,HOXA13,HOXB7,HOXC4,HOXD3,HOXD8,PITX2,PLAGL1,CREB3L2,KLF6,POU2F2,ZBTB16,ZEB2,JUN,ELK3,EPAS1,SMAD1 |
| ETS1,MECOM,MEF2D,ZNF319,EBF4,FOXO1,ATF6,NR5A2,PRDM16,GATA2,GATA3,GATA4,STAT3,STAT5A,TSC22D1,NFATC2,NFE2L2,NFIC,NFKB1,NHLH2,TCF4,SOX17,RUNX2,HHEX,HIVEP2,HOXA1,HOXA5,HOXA13,HOXB7,HOXC4,HOXD3,HOXD8,PITX2,PLAGL1,CREB3L2,KLF6,POU2F2,ZBTB16,ZEB2,JUN,ELK3,EPAS1,SMAD1 |
| FMNL2,PICALM,MRTFB,SNX9,FLNB,FN1,COBLL1,MYO1B,APP,PDLIM5,MSN,FSCN1,PALLD,MYH9,IQGAP1,CCN4,DST,MYPN,THBS1,SEPTIN7,TNR,CDH5,NRG1,PRTG,CDHR3,CEMIP2,JAM2,AHNAK,COL3A1,VWF,NIBAN2,IGF2,CCN2,ARHGAP18,TES,ILK,ITGAV,ITGB3,PTPRM,KDR,JAM3,ARGLU1,RTN4,PHLDB2 |
| TBC1D19,RIN3,CD109,DTX3L,MGAT5,DLC1,MGST2,ARHGAP31,VAV3,MMP15,DOCK6,FNIP2,CIB1,FN1,BMP2K,ANXA4,RGCC,BIRC2,NLRP1,APP,TRIB3,ATP2B4,PHACTR1,MYO9B,TBC1D2B,CCND1,DENND2B,IQGAP1,CFLAR,RGL1,DNAJC1,NOTCH1,RIPK3,KALRN,ARHGEF12,CAV1,TFPI,CCND3,TBC1D8,PKIG,FURIN,DGKI,CDKN2B |
| CD109,LRRC32,IL18R1,CCN4,TGFBR2,THBS1,FURIN,IL1RL1,COL3A1,COL4A1,IGFBP3,IGFBP5,IGFBP7,CCN2,IL4R,ITGAV,ITGB3,PXDN,KDR,HTRA3 |
| CCN4,IGFBP3,IGFBP5,IGFBP7,CCN2,ITGAV,ITGB3,HTRA3 |
| ABL2,RPS6KA2,MAPK12,FES,MERTK,CIB1,CDC42BPA,BMP2K,SGK1,RGCC,AAK1,HIPK2,CDKL4,TRIB3,RIPK1,CCND1,CDKL1,IQGAP1,MLKL,FCSK,DCLK2,NPR1,RIPK3,KALRN,PAPSS2,CCND3,PKIG,TGFBR2,MAP4K5,DGKI,CDKN2B,CDK17,NRG1,RPS6KA5,STK17A,PFKFB3,TXK,PIK3C2B,PIK3CA,PIP4K2A,MAP4K4,SH3BP5 |
| ETS1,ETS2,DTX3L,MEF2D,AHR,EBF4,FOXO1,ZNF703,ANXA4,APBB2,HIPK2,HDAC7,SUFU,PRDM16,GATA2,GATA3,GATA4,CCND1,BCL2,CREG1,STAT3,NFATC2,NFE2L2,NFKB1,NFKBIA,NHLH2,TACC1,TCF4,SOX17,RUNX2,ZFPM2,HEY1,CD34,HEY2,HHEX,PITX2,HES1,CRTC3,RNF19A,CREM,ZNF516,VGLL4,JUN,PARP9,WIPI1 |
| TBC1D19,RIN3,DTX3L,DLC1,MGST2,ARHGAP31,VAV3,MMP15,FN1,RGCC,NLRP1,APP,MYO9B,TBC1D2B,IQGAP1,CFLAR,DNAJC1,ARHGEF12,CAV1,TBC1D8,NRG1,RABGAP1,PIK3CA,RIN2,ARHGAP29,AGFG2,SMAP2,GMFG,IGF2,RGS6,IGFBP3,SH3PXD2A,PRKCE,RAPGEF2,ARHGAP18,DOCK4,DAD1,TBC1D20,EVI5,JUN,ARHGAP25 |
| RGCC,HIPK2,PRDM16,GATA4,TGFBR2,COL3A1,COL5A2,RNF111,ZEB2,JUN,SMAD1,SMAD6,SMAD9 |
| FN1,APP,MYH9,CCN4,DST,THBS1,TNR,NRG1,JAM2,COL3A1,VWF,IGF2,CCN2,ILK,ITGAV,ITGB3,KDR,JAM3 |
| ABL2,RPS6KA2,MAPK12,FES,MERTK,CIB1,CDC42BPA,BMP2K,SGK1,RGCC,AAK1,HIPK2,CDKL4,TRIB3,RIPK1,CCND1,CDKL1,IQGAP1,MLKL,DCLK2,NPR1,RIPK3,KALRN,CCND3,PKIG,TGFBR2,MAP4K5,CDKN2B,CDK17,NRG1,RPS6KA5,STK17A,TXK,PIK3CA,MAP4K4,SH3BP5,ROCK2,GMFG,PIM3,IGF2,PRKACB,PRKAR2B,PRKCE |
| ETS2,PICALM,FZD8,DLC1,ARHGAP31,EVL,SORBS2,FN1,DLGAP4,NLRP1,APP,SKAP1,ATP2B4,TRADD,PAG1,MYH9,RIPK1,GATA2,GATA3,BCL2,PLXND1,IQGAP1,GNG12,NFE2L2,MYPN,TACC1,RUNX2,AFAP1,HHEX,PDE2A,HOXA3,HOXC4,CLCN3,ARHGAP29,HES1,QKI,SH3BP5,POU2F2,BCAR1,RGS6,PRKAR2B,PRKCE,RAPGEF2 |
| ABL2,RPS6KA2,MAPK12,FES,MERTK,CIB1,CDC42BPA,BMP2K,SGK1,RGCC,AAK1,HIPK2,CDKL4,TRIB3,RIPK1,CCND1,CDKL1,IQGAP1,MLKL,FCSK,DCLK2,NPR1,RIPK3,KALRN,PAPSS2,CCND3,PKIG,TGFBR2,MAP4K5,DGKI,CDKN2B,CDK17,NRG1,SGMS2,RPS6KA5,STK17A,PFKFB3,TXK,PIK3C2B,PIK3CA,PIP4K2A,MAP4K4 |
| ABL2,ACTA2,ETS1,MECOM,MEF2D,GREB1L,FZD8,MRTFB,DLC1,JAG1,AHR,VAV3,AMOTL2,SOX6,FOXO1,SORBS2,CIB1,FN1,RGCC,VASH1,HIPK2,HDAC7,RBM20,PDLIM5,RHOB,SCUBE1,DKK1,SNAI2,SUFU,ATP2B4,MYH9,RIPK1,UNC5B,GATA2,GATA3,GATA4,PLXND1,CDKL1,STAT3,STIM1,CFLAR,CCN4,NFATC2,NFE2L2 |
| ACTA2,ETS1,FZD8,MRTFB,JAG1,AHR,VAV3,AMOTL2,FOXO1,CIB1,FN1,RGCC,VASH1,HIPK2,HDAC7,RHOB,SUFU,ATP2B4,MYH9,UNC5B,GATA2,GATA4,PLXND1,STAT3,STIM1,CFLAR,NFE2L2,NOTCH1,TCF4,NPR1,SOX17,GRN,CAV1,ZFPM2,FMNL3,TGFBR2,COL18A1,THBS1,HEY1,CD34,HEY2,TNFRSF1A,CDH5,HHEX,PDE2A |
| ABL2,ETS1,MECOM,GREB1L,FZD8,MRTFB,DLC1,JAG1,AHR,VAV3,AMOTL2,CIB1,FN1,RGCC,VASH1,HIPK2,HDAC7,RHOB,SUFU,ATP2B4,MYH9,UNC5B,SDCCAG8,GATA2,GATA3,GATA4,BCL2,PLXND1,STAT3,STIM1,NFE2L2,NOTCH1,TCF4,NPR1,SOX17,GRN,CAV1,ZFPM2,FMNL3,TGFBR2,COL18A1,THBS1,HEY1,CD34,HEY2 |
| ETS1,FZD8,MRTFB,JAG1,AHR,VAV3,AMOTL2,CIB1,FN1,RGCC,VASH1,HIPK2,HDAC7,RHOB,ATP2B4,MYH9,UNC5B,GATA2,GATA4,PLXND1,STAT3,STIM1,NFE2L2,NOTCH1,TCF4,NPR1,SOX17,GRN,CAV1,ZFPM2,FMNL3,TGFBR2,COL18A1,THBS1,HEY1,CD34,HEY2,TNFRSF1A,CDH5,HHEX,PECAM1,WNT4,CEMIP2,HOXA1,HOXA3 |
| ACTA2,ETS1,FZD8,MRTFB,JAG1,AHR,VAV3,AMOTL2,FOXO1,CIB1,FN1,RGCC,VASH1,HIPK2,HDAC7,RHOB,SUFU,ATP2B4,MYH9,UNC5B,GATA2,GATA4,PLXND1,STAT3,STIM1,CFLAR,NFATC2,NFE2L2,NOTCH1,TCF4,NPR1,SOX17,GRN,CAV1,ZFPM2,FMNL3,TGFBR2,COL18A1,THBS1,HEY1,CD34,HEY2,TNFRSF1A,CDH5,HHEX,PDE2A |
| ABL2,CD109,ETS1,ETS2,WLS,FZD8,DLC1,SLC24A4,JAG1,AHR,VAV3,AMOTL2,MMP15,FOXO1,CIB1,FN1,RGCC,VASH1,HIPK2,ITGA8,HDAC7,SBNO2,RHOB,DKK1,SNAI2,SUFU,ATP2B4,MYH9,UNC5B,SDCCAG8,GATA2,GATA3,GATA4,PLXND1,BCL6,STAT3,STIM1,CFLAR,NFATC2,NFE2L2,NBEAL2,MYPN,NOTCH1,TCF4,NPR1,SOX17 |
| ABL2,ETS1,MECOM,GREB1L,FZD8,MRTFB,DLC1,ATXN1,JAG1,AHR,VAV3,AMOTL2,CIB1,FN1,RGCC,VASH1,HIPK2,HDAC7,RHOB,SUFU,ATP2B4,MYH9,UNC5B,SDCCAG8,GATA2,GATA3,GATA4,BCL2,PLXND1,STAT3,STIM1,NFE2L2,NOTCH1,TCF4,NPR1,WWP1,SOX17,GRN,CAV1,ZFPM2,FMNL3,TGFBR2,COL18A1,THBS1,HEY1,CD34 |
| ETS1,FZD8,JAG1,AHR,VAV3,AMOTL2,CIB1,FN1,RGCC,VASH1,HIPK2,HDAC7,RHOB,ATP2B4,MYH9,UNC5B,GATA2,GATA4,PLXND1,STAT3,STIM1,NFE2L2,NOTCH1,TCF4,NPR1,SOX17,GRN,CAV1,FMNL3,TGFBR2,COL18A1,THBS1,HEY1,CD34,TNFRSF1A,CDH5,HHEX,PECAM1,CEMIP2,HOXA3,NRARP,HOXA5,SLIT2,PIK3CA,PITX2 |
| RIN3,PRR5L,ABL2,SPNS2,ACTA2,ABHD6,ETS1,FMNL2,MRTFB,MGAT5,DLC1,ATP8A1,MTUS1,JAG1,ARID5B,VAV3,AMOTL2,MERTK,ZNF703,EVL,CIB1,CDC42BPA,FN1,SGK1,RGCC,VASH1,APBB2,APP,HDAC7,MSN,RHOB,RHOG,SNAI2,CD274,LIMCH1,FSCN1,ATP2B4,PALLD,PHACTR1,TRADD,MYH9,MYO9B,DISC1,SDCCAG8,RAB11A |
| ABL2,ACTA2,MECOM,MEF2D,GREB1L,MRTFB,DLC1,CLIC5,SLC24A4,JAG1,ARID5B,AHR,MLLT3,SOX6,FOXO1,EVL,HIPK2,ITGA8,ANKRD6,MSN,LRIG3,DKK1,NR5A2,SNAI2,SUFU,CSGALNACT1,GATA2,GATA3,GATA4,BCL2,PLXND1,STAT3,STAT5A,STIM1,CFLAR,BMP6,NFIC,NOTCH1,SOX17,CAV1,RUNX2,ZFPM2,TGFBR2,COL18A1 |
| ABL2,FMNL2,PICALM,DLC1,CLIC5,HECW2,FES,AHR,MERTK,SOX6,EVL,FLNB,CIB1,FN1,SGK1,APBB2,ITGA8,APP,CPNE5,CDC42EP3,PDLIM5,MSN,RHOB,RHOG,DKK1,PALLD,PHACTR1,MYH9,SPR,MYO9B,UNC5B,DISC1,RAB11A,GATA3,BCL2,PLXND1,BCL6,IQGAP1,DST,NBEAL2,MYPN,NOTCH1,KALRN,SZT2,SOX17,FRYL,FMNL3 |
| RIN3,PRR5L,ABL2,ACTA2,ABHD6,MALAT1,ETS1,MGAT5,DLC1,ATP8A1,MTUS1,JAG1,FES,AMOTL2,ZNF703,EVL,CIB1,FN1,SGK1,RGCC,VASH1,APP,HDAC7,MSN,RHOB,RHOG,SNAI2,CD274,LIMCH1,ATP2B4,PHACTR1,TRADD,RAB11A,GATA2,GATA3,PBLD,BCL2,PLXND1,STAT3,STAT5A,IQGAP1,CCN4,NFE2L2,NOTCH1,RIPK3,ABHD2 |
| RIN3,PRR5L,ABL2,ACTA2,ABHD6,MALAT1,ETS1,MGAT5,DLC1,ATP8A1,MTUS1,JAG1,FES,AMOTL2,ZNF703,EVL,CIB1,FN1,SGK1,RGCC,VASH1,APP,HDAC7,MSN,RHOB,RHOG,SNAI2,CD274,LIMCH1,ATP2B4,PHACTR1,TRADD,RAB11A,GATA2,GATA3,GATA4,PBLD,BCL2,PLXND1,BCL6,STAT3,STAT5A,IQGAP1,CCN4,NFE2L2,NOTCH1 |
| RIN3,PRR5L,ACTA2,ABHD6,ETS1,MGAT5,DLC1,ATP8A1,MTUS1,JAG1,AMOTL2,ZNF703,EVL,CIB1,FN1,SGK1,RGCC,VASH1,APP,HDAC7,MSN,RHOB,RHOG,SNAI2,CD274,LIMCH1,ATP2B4,PHACTR1,TRADD,RAB11A,GATA2,GATA3,PBLD,BCL2,PLXND1,STAT3,STAT5A,IQGAP1,CCN4,NFE2L2,NOTCH1,RIPK3,ABHD2,GRN,CAV1 |
| PRR5L,ACTA2,ETS1,ARID5B,AMOTL2,EVL,CIB1,FN1,RGCC,VASH1,HDAC7,RHOB,SNAI2,ATP2B4,MYH9,RAB11A,GATA2,GATA3,PBLD,PLXND1,STAT5A,IQGAP1,NFE2L2,NOTCH1,SOX17,GRN,TGFBR2,THBS1,CDH5,PECAM1,SLIT2,PIK3CA,RIN2,PITX2,PLCG1,MAP4K4,VEGFC,ROCK2,BCAR1,ADGRA2,IGF2,PRCP,CLEC14A |
| RIN3,PRR5L,ABL2,ACTA2,ABHD6,MALAT1,ETS1,MGAT5,DLC1,ATP8A1,MTUS1,JAG1,FES,AMOTL2,ZNF703,EVL,CIB1,FN1,SGK1,RGCC,VASH1,APP,HDAC7,MSN,RHOB,RHOG,SNAI2,CD274,LIMCH1,ATP2B4,PHACTR1,TRADD,RAB11A,GATA2,GATA3,PBLD,BCL2,PLXND1,STAT3,STAT5A,IQGAP1,CCN4,NFE2L2,NOTCH1,RIPK3,ABHD2 |
| ABL2,ETS1,FMNL2,PICALM,DLC1,HECW2,FES,MLLT3,CIB1,FN1,SGK1,RGCC,VASH1,HIPK2,ANKRD6,CPNE5,CDC42EP3,PDLIM5,MSN,RHOB,RHOG,DKK1,SNAI2,ATP2B4,MYH9,DISC1,RAB11A,GATA2,GATA3,GATA4,PLXND1,BCL6,STAT3,STIM1,NFE2L2,NOTCH1,TCF4,NPR1,KALRN,GRN,RUNX2,FMNL3,TGFBR2,THBS1,CD34,PALM |
| ACTA2,ETS1,EVL,CIB1,RGCC,VASH1,HDAC7,RHOB,SNAI2,ATP2B4,MYH9,RAB11A,GATA2,GATA3,PBLD,PLXND1,STAT5A,NFE2L2,NOTCH1,GRN,TGFBR2,THBS1,CDH5,PECAM1,SLIT2,PIK3CA,RIN2,PLCG1,MAP4K4,VEGFC,ROCK2,BCAR1,ADGRA2,IGF2,PRCP,CLEC14A,MIR21,PRKCE,ITGAV,ITGB3,GADD45A,ZEB2,GAB2,JUN,PTPRM |
| ETS1,EVL,CIB1,RGCC,VASH1,HDAC7,RHOB,SNAI2,ATP2B4,MYH9,RAB11A,GATA2,GATA3,PBLD,PLXND1,STAT5A,NFE2L2,NOTCH1,GRN,TGFBR2,THBS1,CDH5,PECAM1,SLIT2,PIK3CA,RIN2,PLCG1,MAP4K4,VEGFC,ROCK2,BCAR1,ADGRA2,IGF2,PRCP,CLEC14A,MIR21,PRKCE,ITGAV,ITGB3,GADD45A,ZEB2,GAB2,JUN,PTPRM,ADAMTS9 |
| ETS1,EVL,CIB1,RGCC,VASH1,HDAC7,RHOB,SNAI2,ATP2B4,MYH9,RAB11A,GATA2,GATA3,PBLD,PLXND1,STAT5A,NFE2L2,NOTCH1,GRN,TGFBR2,THBS1,CDH5,PECAM1,SLIT2,PIK3CA,RIN2,PLCG1,MAP4K4,VEGFC,ROCK2,BCAR1,ADGRA2,IGF2,PRCP,CLEC14A,MIR21,PRKCE,ITGAV,ITGB3,GADD45A,ZEB2,GAB2,JUN,PTPRM,ADAMTS9 |
| CD109,ETS1,PICALM,ATXN1,JAG1,SLC9B2,FES,AHR,SOX6,CIB1,FN1,BMP2K,SGK1,RGCC,VASH1,HIPK2,RAB14,RHOB,DKK1,SNAI2,PRDM16,ATP2B4,ZDHHC21,TRADD,RIPK1,DISC1,RAB11A,GATA2,GATA3,GATA4,CCND1,PLXND1,BCL6,STAT3,STAT5A,STIM1,CCN4,BMP6,NFATC2,NFE2L2,NFKBIA,NOTCH1,TCF4,NPR1,KALRN |
| ETS1,MECOM,PICALM,MAPK12,ATXN1,JAG1,SLC9B2,FES,SOX6,ZNF703,CIB1,FN1,AAMDC,SGK1,RGCC,APP,DKK1,SNAI2,ZFHX3,PRDM16,CCDC71L,ZHX3,RIPK1,DISC1,RAB11A,GATA2,GATA3,GATA4,BCL2,PLXND1,BCL6,STAT3,STAT5A,CCN4,BMP6,NFKB1,NOTCH1,TCF4,KALRN,SOX17,RUNX2,TGFBR2,HEY1,CD34,TNFRSF1B |
| ACTA2,CD109,ETS1,VAV3,MERTK,DOCK6,EVL,FN1,VASH1,SCUBE1,SNAI2,AHNAK2,NEAT1,MYH9,GATA2,GATA3,GATA4,BCL2,CFLAR,CCN4,LYVE1,DST,NFE2L2,NBEAL2,ABHD2,GRN,CAV1,PAPSS2,GSN,ZFPM2,TFPI,TGFBR2,THBS1,CD34,ENTPD1,DGKI,TNR,NRG1,HHEX,PECAM1,WNT4,LYST,TXK,PIK3CA,CREB3L2,COL3A1,VWF |
| ABL2,CD109,ATXN1,ARID5B,FES,VAV3,ATP6V0D2,MERTK,FOXO1,ZNF703,HIPK2,ITGA8,RAB14,DKK1,PRDM16,TRADD,LRRC32,GATA3,GATA4,PBLD,STAT3,STAT5A,IQGAP1,SHKBP1,BMP6,NOTCH1,TCF4,NPR1,KALRN,TNS2,CAV1,RUNX2,CCND3,TGFBR2,THBS1,FURIN,CDH5,CDKN2B,NRG1,ELAPOR2,HHEX,RPS6KA5,WNT4,LRIT3 |
| ETS1,MECOM,PICALM,MRTFB,MAPK12,ATXN1,JAG1,SLC9B2,FES,SOX6,ZNF703,CIB1,FN1,AAMDC,SGK1,RGCC,HIPK2,APP,CPNE5,RHOB,DKK1,SNAI2,ZFHX3,PRDM16,CCDC71L,ZHX3,TRADD,RIPK1,DISC1,RAB11A,GATA2,GATA3,GATA4,CCND1,BCL2,PLXND1,BCL6,STAT3,STAT5A,IQGAP1,STIM1,CCN4,BMP6,NFATC2,NFE2L2 |
| ABCA1,CD109,ETS1,MECOM,PICALM,MAPK12,ATXN1,JAG1,SLC9B2,FES,AHR,SOX6,FOXO1,ZNF703,CIB1,FN1,AAMDC,SGK1,RGCC,BIRC2,APP,HDAC7,DKK1,TRIB3,SNAI2,ZFHX3,SUFU,PRDM16,ZDHHC21,CCDC71L,ZHX3,RIPK1,DISC1,RAB11A,GATA2,GATA3,GATA4,PBLD,CCND1,BCL2,PLXND1,BCL6,STAT3,STAT5A,CCN4,BMP6 |
| ETS1,EVL,CIB1,RGCC,VASH1,HDAC7,RHOB,SNAI2,ATP2B4,RAB11A,GATA2,GATA3,PBLD,STAT5A,NFE2L2,NOTCH1,GRN,TGFBR2,THBS1,SLIT2,RIN2,PLCG1,MAP4K4,VEGFC,ROCK2,BCAR1,ADGRA2,IGF2,PRCP,MIR21,PRKCE,ITGB3,GADD45A,GAB2,JUN,PTPRM,ADAMTS9,IQSEC1,KDR,RTN4,EDN1 |
| ABL2,ACTA2,FMNL2,MRTFB,DLC1,FES,AMOTL2,SNX9,EVL,FLNB,SORBS2,CDC42BPA,RGCC,APBB2,MYO1B,CDC42EP3,PDLIM5,SMTN,RHOB,RHOG,LIMCH1,FSCN1,PALLD,SCN3B,PHACTR1,MYH9,MYO9B,GATA4,BCL2,BCL6,IQGAP1,CFLAR,MYPN,CAV1,COTL1,GSN,FMNL3,MICALCL,PDE4B,PECAM1,TMOD2,PARVB,WNT4,SLIT2,PIK3CA |
| ABCA1,ABL2,ETS1,PICALM,MRTFB,ATXN1,JAG1,SLC9B2,FES,PIBF1,SOX6,ZNF703,CIB1,FN1,SGK1,RGCC,HIPK2,BIRC2,NLRP1,APP,SMTN,RHOB,DKK1,CD274,PRDM16,SCN3B,TRADD,RIPK1,DISC1,RAB11A,GATA2,GATA3,GATA4,CCND1,BCL2,PLXND1,BCL6,IL18R1,STAT3,STAT5A,STIM1,CCN4,BMP6,NFE2L2,NFKB1,NOTCH1 |
| ABL2,PICALM,CLIC5,HECW2,MERTK,EVL,FLNB,CIB1,FN1,SGK1,APBB2,ITGA8,APP,PDLIM5,RHOG,DKK1,PALLD,PHACTR1,MYH9,SPR,UNC5B,DISC1,RAB11A,GATA3,BCL2,PLXND1,DST,NBEAL2,MYPN,NOTCH1,KALRN,SZT2,SOX17,COL18A1,TTL,SEPTIN7,TNR,NRG1,PRTG,RPS6KA5,PECAM1,MGLL,PARVB,HOXA1,HOXA13,SLIT2 |
| ETS1,CIB1,RGCC,VASH1,HDAC7,RHOB,SNAI2,ATP2B4,MYH9,GATA2,GATA3,PLXND1,STAT5A,NFE2L2,NOTCH1,GRN,THBS1,CDH5,PECAM1,SLIT2,PIK3CA,RIN2,PLCG1,VEGFC,ROCK2,BCAR1,ADGRA2,IGF2,PRCP,CLEC14A,MIR21,ITGAV,ITGB3,GADD45A,PTPRM,ADAMTS9,KDR,EDN1,EFNB2 |
| ABL2,SLC25A34,ETS2,MECOM,WLS,MRTFB,DLC1,CLIC5,AHR,MMP15,SOX6,FN1,VEZT,VASH1,HIPK2,ITGA8,RAB14,LRIG3,DKK1,NR5A2,SUFU,MYH9,GATA2,GATA3,GATA4,NFE2L2,NOTCH1,SOX17,GRN,RUNX2,ZFPM2,TGFBR2,HEY1,FURIN,HEY2,SEPTIN7,HHEX,HLX,WNT4,HOXA1,HOXA3,NRARP,HOXA4,HOXA5,HOXA6,HOXA13,SLIT2 |
| ABL2,ACTA2,ETS2,WLS,GREB1L,DLC1,JAG1,AHR,MLLT3,MMP15,FN1,ITGA8,ANKRD6,MSN,RHOB,DKK1,SNAI2,SUFU,GATA3,GATA4,BCL2,PLXND1,STAT5A,NOTCH1,SOX17,ZFPM2,TGFBR2,HEY1,HEY2,NRG1,HHEX,WNT4,NRARP,HOXA5,HOXA13,SLIT2,HOXB7,AP2S1,PITX2,HES1,VEGFC,COL3A1,COL4A1,ROCK2,COL4A2,COL5A2 |
| MECOM,MEF2D,GREB1L,MRTFB,DLC1,JAG1,AHR,SOX6,SORBS2,FN1,RBM20,PDLIM5,SCUBE1,DKK1,SNAI2,SUFU,RIPK1,GATA3,GATA4,PLXND1,CDKL1,CCN4,NOTCH1,SOX17,ZFPM2,TGFBR2,THBS1,HEY1,HEY2,TNFRSF1A,TNFRSF1B,NRG1,HHEX,PDE2A,HOXA13,SLIT2,PITX2,HES1,AKAP6,COL3A1,ROCK2,MIR21,VGLL4,ILK,SMYD2 |
| ETS1,CIB1,HDAC7,RHOB,SNAI2,RAB11A,GATA2,GATA3,STAT5A,NFE2L2,GRN,TGFBR2,THBS1,RIN2,PLCG1,MAP4K4,VEGFC,ROCK2,BCAR1,ADGRA2,IGF2,MIR21,PRKCE,ITGB3,GAB2,JUN,IQSEC1,KDR,RTN4,EDN1 |
| PRR5L,ACTA2,ETS1,ETS2,MECOM,DTX3L,PICALM,MEF2D,MRTFB,ATXN1,JAG1,ARID5B,AHR,AFF1,MLLT3,ZNF319,MAML3,EBF4,SOX6,FOXO1,AAMDC,RGCC,HIPK2,ITGA8,BIRC2,ZNF407,APP,PCGF5,RBM20,SBNO2,RHOG,ATF6,SKAP1,NR5A2,ZFHX3,SSBP4,PRDM16,ARID4B,RIPK1,GATA2,GATA3,GATA4,PLXND1,STAT3,STAT5A,BMP6 |
| ACTA2,MECOM,GREB1L,JAG1,ARID5B,AHR,ANXA4,ITGA8,GATA3,BCL2,PLXND1,IQGAP1,CFLAR,BMP6,TMED10,NOTCH1,SOX17,TNS2,HEY1,CD34,PECAM1,WNT4,SLIT2,HOXB7,HES1,COL4A1,WNT2B,WWTR1,ZBTB16,ILK,MTSS1,ITGB3,CXCR4,FAT4,KIF26B,RDH10,EFNB2,ENPEP,SMAD1,EPHA4,SMAD6,SMAD9 |
| ACTA2,MECOM,GREB1L,JAG1,ARID5B,AHR,ANXA4,ITGA8,GATA2,GATA3,BCL2,PLXND1,STAT5A,IQGAP1,CFLAR,BMP6,TMED10,NOTCH1,SOX17,TNS2,HEY1,CD34,PECAM1,WNT4,HOXA13,SLIT2,HOXB7,HES1,COL4A1,WNT2B,WWTR1,ZBTB16,ILK,MTSS1,ITGB3,CXCR4,FAT4,KIF26B,RDH10,EFNB2,ENPEP,SMAD1,EPHA4,SMAD6,SMAD9 |
| ABL2,PICALM,FZD8,MRTFB,CLIC5,ATXN1,HECW2,JAG1,FES,SOX6,EVL,CIB1,FN1,SGK1,APBB2,HIPK2,APP,CPNE5,PDLIM5,RHOG,DKK1,ZFHX3,SUFU,PRDM16,PALLD,PHACTR1,SPR,UNC5B,DISC1,SDCCAG8,RAB11A,GATA2,GATA3,BCL2,PLXND1,BCL6,STAT3,IQGAP1,CFLAR,BMP6,DST,NFE2L2,MYPN,TACC1,DCLK2,CAMSAP2 |
| TBC1D19,ABL2,FMNL2,MRTFB,DLC1,FES,AMOTL2,PIBF1,SNX9,EVL,FLNB,SORBS2,CIB1,CDC42BPA,SGK1,RGCC,FIGNL2,MYO1B,CDC42EP3,PDLIM5,SMTN,MSN,RHOB,RHOG,TTLL11,LIMCH1,TPPP3,FSCN1,VAMP4,CFAP44,PALLD,PHACTR1,MYH9,DISC1,SDCCAG8,RAB11A,TBC1D2B,BCL2,BCL6,CDKL1,IQGAP1,CFLAR,DST,MYPN |
| ABL2,PICALM,FZD8,MRTFB,CLIC5,HECW2,JAG1,FES,EVL,CIB1,FN1,SGK1,APBB2,HIPK2,APP,CPNE5,PDLIM5,RHOG,DKK1,ZFHX3,SUFU,PALLD,PHACTR1,SPR,UNC5B,DISC1,RAB11A,GATA2,GATA3,BCL2,PLXND1,BCL6,STAT3,IQGAP1,CFLAR,BMP6,DST,NFE2L2,MYPN,DCLK2,CAMSAP2,NOTCH1,TCF4,KALRN,SZT2,CTDSP1,GRN |
| ABL2,ETS1,DLC1,JAG1,FES,VAV3,MERTK,ZNF703,CIB1,FN1,VEZT,RGCC,ITGA8,APP,PDLIM5,MSN,RHOB,CD93,SKAP1,SNAI2,CD274,ZFHX3,LIMCH1,PALLD,ZDHHC21,PAG1,MYH9,DISC1,LRRC32,CD164,GATA3,BCL2,PLXND1,BCL6,LRRC4,PCDH17,STAT5A,IQGAP1,CCN4,LYVE1,BMP6,DST,MYPN,NOTCH1,NT5E,CAV1,GSN,TGFBR2 |
| JAG1,MSN,RHOB,SCUBE1,ZDHHC21,BMP6,NOTCH1,SOX17,COL18A1,HEY1,CD34,HEY2,TNFRSF1A,CDH5,NRG1,PDE2A,PECAM1,HOXA13,ROCK2,COL15A1,CLIC4,MIR21,RAPGEF2,CXCR4,KDR,RTN4,S1PR1 |
| ACTA2,MALAT1,ETS1,MGAT5,ATP8A1,ZNF703,CIB1,FN1,APP,HDAC7,RHOB,SNAI2,CD274,TRADD,RAB11A,GATA2,GATA3,BCL2,STAT3,STAT5A,IQGAP1,CCN4,NFE2L2,NOTCH1,GRN,CAV1,TGFBR2,COL18A1,THBS1,FURIN,CDH5,PECAM1,JAM2,SLIT2,RIN2,PLCG1,MAP4K4,VEGFC,ROCK2,BCAR1,ADGRA2,IGF2,IGFBP5,MIR21,PRKCE |
| ABL2,FMNL2,MRTFB,DLC1,FES,AMOTL2,SNX9,EVL,FLNB,SORBS2,CDC42BPA,RGCC,MYO1B,CDC42EP3,PDLIM5,SMTN,RHOB,RHOG,LIMCH1,FSCN1,PALLD,PHACTR1,MYH9,BCL2,BCL6,IQGAP1,CFLAR,MYPN,COTL1,GSN,FMNL3,MICALCL,PECAM1,TMOD2,PARVB,WNT4,SLIT2,PIK3CA,ROCK2,GMFG,BCAR1,PALM2AKAP2,MIR21,PRKCE |
| CD109,SOX6,ZNF703,CIB1,HIPK2,ITGA8,APP,RAB14,DKK1,SNAI2,ZFHX3,PRDM16,ATP2B4,RIPK1,LRRC32,GATA3,GATA4,PBLD,IQGAP1,CFLAR,BMP6,NFKB1,NOTCH1,TCF4,CAV1,RUNX2,TGFBR2,THBS1,FURIN,TNFRSF1B,CDH5,CDKN2B,ELAPOR2,HHEX,PDE2A,PDE3A,WNT4,LRIT3,HOXA13,SLIT2,PIK3CA,HES1,PLCG1,VEGFC |
| JAG1,SNAI2,GATA3,GATA4,NOTCH1,ZFPM2,TGFBR2,HEY1,HEY2,TNFRSF1A,TNFRSF1B,PDE2A,SLIT2,PITX2,ROCK2,ADAMTS9,RB1,SMAD6 |

| | | | | | | | | |
|---|---|---|---|---|---|---|---|---|
| GO: Biological Process | GO:2000147 | positive regulation of cell motility | 6.84E-12 | 4.81E-08 | 1.00E-09 | 9.45E-09 | 60 | 662 |
| GO: Biological Process | GO:0051272 | positive regulation of cellular component movement | 7.42E-12 | 5.22E-08 | 1.07E-09 | 1.01E-08 | 61 | 681 |
| GO: Biological Process | GO:0048699 | generation of neurons | 8.26E-12 | 5.81E-08 | 1.16E-09 | 1.10E-08 | 117 | 1802 |
| GO: Biological Process | GO:0045944 | positive regulation of transcription by RNA polymerase II | 8.77E-12 | 6.17E-08 | 1.21E-09 | 1.14E-08 | 95 | 1339 |
| GO: Biological Process | GO:0071363 | cellular response to growth factor stimulus | 9.30E-12 | 6.54E-08 | 1.21E-09 | 1.14E-08 | 66 | 775 |
| GO: Biological Process | GO:0030198 | extracellular matrix organization | 9.36E-12 | 6.58E-08 | 1.21E-09 | 1.14E-08 | 46 | 431 |
| GO: Biological Process | GO:0045446 | endothelial cell differentiation | 9.56E-12 | 6.72E-08 | 1.21E-09 | 1.14E-08 | 25 | 139 |
| GO: Biological Process | GO:0007155 | cell adhesion | 9.81E-12 | 6.90E-08 | 1.21E-09 | 1.14E-08 | 106 | 1571 |
| GO: Biological Process | GO:0051093 | negative regulation of developmental process | 9.81E-12 | 6.90E-08 | 1.21E-09 | 1.14E-08 | 83 | 1100 |
| GO: Biological Process | GO:0048534 | hematopoietic or lymphoid organ development | 9.92E-12 | 6.98E-08 | 1.21E-09 | 1.14E-08 | 84 | 1120 |
| GO: Biological Process | GO:0030335 | positive regulation of cell migration | 1.01E-11 | 7.09E-08 | 1.21E-09 | 1.14E-08 | 58 | 633 |
| GO: Biological Process | GO:0043062 | extracellular structure organization | 1.01E-11 | 7.13E-08 | 1.21E-09 | 1.14E-08 | 46 | 432 |
| GO: Biological Process | GO:1903508 | positive regulation of nucleic acid-templated transcription | 1.06E-11 | 7.44E-08 | 1.22E-09 | 1.15E-08 | 114 | 1744 |
| GO: Biological Process | GO:0045893 | positive regulation of transcription, DNA-templated | 1.06E-11 | 7.44E-08 | 1.22E-09 | 1.15E-08 | 114 | 1744 |
| GO: Biological Process | GO:1902680 | positive regulation of RNA biosynthetic process | 1.10E-11 | 7.72E-08 | 1.25E-09 | 1.18E-08 | 114 | 1745 |
| GO: Biological Process | GO:0045229 | external encapsulating structure organization | 1.19E-11 | 8.36E-08 | 1.33E-09 | 1.25E-08 | 46 | 434 |
| GO: Biological Process | GO:0001822 | kidney development | 2.06E-11 | 1.45E-07 | 2.26E-09 | 2.13E-08 | 40 | 347 |
| GO: Biological Process | GO:0032970 | regulation of actin filament-based process | 2.41E-11 | 1.69E-07 | 2.61E-09 | 2.46E-08 | 46 | 443 |
| GO: Biological Process | GO:0002520 | immune system development | 3.04E-11 | 2.14E-07 | 3.24E-09 | 3.06E-08 | 86 | 1185 |
| GO: Biological Process | GO:0001775 | cell activation | 3.11E-11 | 2.19E-07 | 3.26E-09 | 3.08E-08 | 107 | 1623 |
| GO: Biological Process | GO:0030097 | hemopoiesis | 4.57E-11 | 3.21E-07 | 4.73E-09 | 4.46E-08 | 80 | 1074 |
| GO: Biological Process | GO:0090287 | regulation of cellular response to growth factor stimulus | 5.59E-11 | 3.93E-07 | 5.69E-09 | 5.37E-08 | 39 | 343 |
| GO: Biological Process | GO:0042060 | wound healing | 7.26E-11 | 5.11E-07 | 7.29E-09 | 6.88E-08 | 54 | 594 |
| GO: Biological Process | GO:0042127 | regulation of cell population proliferation | 1.37E-10 | 9.62E-07 | 1.36E-08 | 1.28E-07 | 121 | 1973 |
| GO: Biological Process | GO:1901342 | regulation of vasculature development | 1.76E-10 | 1.24E-06 | 1.72E-08 | 1.62E-07 | 41 | 388 |
| GO: Biological Process | GO:0097435 | supramolecular fiber organization | 1.88E-10 | 1.32E-06 | 1.81E-08 | 1.70E-07 | 64 | 793 |
| GO: Biological Process | GO:0048598 | embryonic morphogenesis | 2.00E-10 | 1.40E-06 | 1.90E-08 | 1.79E-07 | 59 | 701 |
| GO: Biological Process | GO:0032956 | regulation of actin cytoskeleton organization | 2.60E-10 | 1.83E-06 | 2.44E-08 | 2.30E-07 | 41 | 393 |
| GO: Biological Process | GO:0007219 | Notch signaling pathway | 2.77E-10 | 1.95E-06 | 2.57E-08 | 2.42E-07 | 29 | 214 |
| GO: Biological Process | GO:0010594 | regulation of endothelial cell migration | 3.14E-10 | 2.21E-06 | 2.86E-08 | 2.70E-07 | 31 | 243 |
| GO: Biological Process | GO:0001501 | skeletal system development | 3.28E-10 | 2.31E-06 | 2.96E-08 | 2.79E-07 | 52 | 583 |
| GO: Biological Process | GO:0031589 | cell-substrate adhesion | 3.54E-10 | 2.49E-06 | 3.16E-08 | 2.98E-07 | 41 | 397 |
| GO: Biological Process | GO:0045765 | regulation of angiogenesis | 3.87E-10 | 2.72E-06 | 3.40E-08 | 3.21E-07 | 40 | 382 |
| GO: Biological Process | GO:0030030 | cell projection organization | 5.33E-10 | 3.75E-06 | 4.63E-08 | 4.36E-07 | 116 | 1904 |
| GO: Biological Process | GO:0071495 | cellular response to endogenous stimulus | 5.63E-10 | 3.96E-06 | 4.83E-08 | 4.56E-07 | 98 | 1510 |
| GO: Biological Process | GO:1905314 | semi-lunar valve development | 5.88E-10 | 4.13E-06 | 4.98E-08 | 4.70E-07 | 13 | 41 |
| GO: Biological Process | GO:0007160 | cell-matrix adhesion | 7.80E-10 | 5.48E-06 | 6.53E-08 | 6.16E-07 | 31 | 252 |
| GO: Biological Process | GO:0009952 | anterior/posterior pattern specification | 9.00E-10 | 6.33E-06 | 7.45E-08 | 7.02E-07 | 30 | 239 |
| GO: Biological Process | GO:0030155 | regulation of cell adhesion | 1.03E-09 | 7.26E-06 | 8.44E-08 | 7.96E-07 | 64 | 827 |
| GO: Biological Process | GO:0072073 | kidney epithelium development | 1.06E-09 | 7.47E-06 | 8.59E-08 | 8.10E-07 | 24 | 159 |
| GO: Biological Process | GO:0010557 | positive regulation of macromolecule biosynthetic process | 1.13E-09 | 7.92E-06 | 9.00E-08 | 8.50E-07 | 119 | 1996 |
| GO: Biological Process | GO:0051493 | regulation of cytoskeleton organization | 1.21E-09 | 8.51E-06 | 9.56E-08 | 9.02E-07 | 50 | 569 |
| GO: Biological Process | GO:0003176 | aortic valve development | 1.41E-09 | 9.93E-06 | 1.10E-07 | 1.04E-06 | 12 | 36 |
| GO: Biological Process | GO:0031175 | neuron projection development | 1.47E-09 | 1.03E-05 | 1.14E-07 | 1.07E-06 | 82 | 1197 |
| GO: Biological Process | GO:0001570 | vasculogenesis | 1.60E-09 | 1.13E-05 | 1.23E-07 | 1.16E-06 | 19 | 102 |
| GO: Biological Process | GO:0120036 | plasma membrane bounded cell projection organization | 2.41E-09 | 1.70E-05 | 1.82E-07 | 1.72E-06 | 112 | 1863 |
| GO: Biological Process | GO:0051241 | negative regulation of multicellular organismal process | 2.50E-09 | 1.76E-05 | 1.87E-07 | 1.77E-06 | 86 | 1295 |
| GO: Biological Process | GO:0010595 | positive regulation of endothelial cell migration | 3.19E-09 | 2.25E-05 | 2.36E-07 | 2.23E-06 | 22 | 142 |
| GO: Biological Process | GO:0061061 | muscle structure development | 3.37E-09 | 2.37E-05 | 2.47E-07 | 2.33E-06 | 59 | 755 |
| GO: Biological Process | GO:0007169 | transmembrane receptor protein tyrosine kinase signaling pathway | 3.61E-09 | 2.54E-05 | 2.62E-07 | 2.47E-06 | 61 | 795 |
| GO: Biological Process | GO:0003179 | heart valve morphogenesis | 5.35E-09 | 3.76E-05 | 3.83E-07 | 3.61E-06 | 14 | 57 |
| GO: Biological Process | GO:0044087 | regulation of cellular component biogenesis | 5.39E-09 | 3.79E-05 | 3.83E-07 | 3.61E-06 | 75 | 1085 |
| GO: Biological Process | GO:0009719 | response to endogenous stimulus | 5.91E-09 | 4.16E-05 | 4.16E-07 | 3.92E-06 | 107 | 1780 |
| GO: Biological Process | GO:0010941 | regulation of cell death | 8.73E-09 | 6.14E-05 | 6.08E-07 | 5.73E-06 | 114 | 1952 |
| GO: Biological Process | GO:0045596 | negative regulation of cell differentiation | 9.22E-09 | 6.48E-05 | 6.35E-07 | 6.00E-06 | 59 | 776 |
| GO: Biological Process | GO:0006468 | protein phosphorylation | 1.03E-08 | 7.26E-05 | 7.05E-07 | 6.65E-06 | 106 | 1776 |
| GO: Biological Process | GO:0060562 | epithelial tube morphogenesis | 1.05E-08 | 7.40E-05 | 7.07E-07 | 6.67E-06 | 39 | 411 |
| GO: Biological Process | GO:0002521 | leukocyte differentiation | 1.06E-08 | 7.44E-05 | 7.07E-07 | 6.67E-06 | 51 | 626 |
| GO: Biological Process | GO:0048871 | multicellular organismal homeostasis | 1.07E-08 | 7.49E-05 | 7.07E-07 | 6.67E-06 | 49 | 589 |
| GO: Biological Process | GO:0048666 | neuron development | 1.10E-08 | 7.71E-05 | 7.20E-07 | 6.80E-06 | 87 | 1358 |
| GO: Biological Process | GO:0008360 | regulation of cell shape | 1.13E-08 | 7.95E-05 | 7.36E-07 | 6.94E-06 | 23 | 165 |
| GO: Biological Process | GO:0007596 | blood coagulation | 1.16E-08 | 8.16E-05 | 7.49E-07 | 7.06E-06 | 36 | 362 |
| GO: Biological Process | GO:0045321 | leukocyte activation | 1.18E-08 | 8.27E-05 | 7.52E-07 | 7.10E-06 | 91 | 1447 |
| GO: Biological Process | GO:0072006 | nephron development | 1.27E-08 | 8.93E-05 | 8.05E-07 | 7.59E-06 | 23 | 166 |
| GO: Biological Process | GO:0002009 | morphogenesis of an epithelium | 1.46E-08 | 1.03E-04 | 9.17E-07 | 8.66E-06 | 53 | 670 |
| GO: Biological Process | GO:0022604 | regulation of cell morphogenesis | 1.61E-08 | 1.13E-04 | 1.00E-06 | 9.46E-06 | 35 | 350 |
| GO: Biological Process | GO:0007015 | actin filament organization | 1.64E-08 | 1.15E-04 | 1.01E-06 | 9.55E-06 | 42 | 470 |
| GO: Biological Process | GO:0007599 | hemostasis | 1.66E-08 | 1.17E-04 | 1.01E-06 | 9.57E-06 | 36 | 367 |
| GO: Biological Process | GO:0010648 | negative regulation of cell communication | 1.72E-08 | 1.21E-04 | 1.04E-06 | 9.81E-06 | 100 | 1658 |
| GO: Biological Process | GO:0050817 | coagulation | 1.78E-08 | 1.25E-04 | 1.07E-06 | 1.01E-05 | 36 | 368 |
| GO: Biological Process | GO:0023057 | negative regulation of signaling | 1.88E-08 | 1.32E-04 | 1.12E-06 | 1.06E-05 | 100 | 1661 |

ACTA2,MALAT1,ETS1,MGAT5,ATP8A1,ZNF703,CIB1,FN1,APP,HDAC7,RHOB,SNAI2,CD274,TRADD,RAB11A,GATA2,GATA3,BCL2,STAT3,STAT5A,IQGAP1,CCN4,NFE2L2,NOTCH1,GRN,CAV1,TGFBR2,COL18A1,THBS1,FURIN,CDH5,PECAM1,JAM2,RIN2,PLCG1,MAP4K4,VEGFC,ROCK2,BCAR1,ADGRA2,IGF2,IGFBP5,MIR21,PRKCE,[illegible]

ACTA2,MALAT1,ETS1,MGAT5,ATP8A1,ZNF703,CIB1,FN1,APP,HDAC7,RHOB,SNAI2,CD274,TRADD,RAB11A,GATA2,GATA3,BCL2,BCL6,STAT3,STAT5A,IQGAP1,CCN4,NFE2L2,NOTCH1,GRN,CAV1,TGFBR2,COL18A1,THBS1,FURIN,CDH5,PECAM1,JAM2,RIN2,PLCG1,MAP4K4,VEGFC,ROCK2,BCAR1,ADGRA2,IGF2,IGFBP5,MIR21,PRKCE,CA

ABL2,PICALM,FZD8,MRTFB,CLIC5,ATXN1,HECW2,JAG1,FES,EVL,CIB1,FN1,SGK1,APBB2,HIPK2,APP,CPNE5,PDLIM5,RHOG,DKK1,ZFHX3,SUFU,PALLD,PHACTR1,SPR,UNC5B,DISC1,SDCCAG8,RAB11A,GATA2,GATA3,BCL2,PLXND1,BCL6,STAT3,IQGAP1,CFLAR,BMP6,DST,NFE2L2,MYPN,DCLK2,CAMSAP2,NOTCH1,TCF4,KALRN,SZT2,C

ETS1,ETS2,MECOM,MEF2D,MRTFB,ATXN1,JAG1,AHR,ZNF319,MAML3,EBF4,SOX6,FOXO1,AAMDC,RGCC,HIPK2,ITGA8,ZNF407,APP,PCGF5,SBNO2,ATF6,SKAP1,NR5A2,ZFHX3,SSBP4,PRDM16,ARID4B,RIPK1,GATA2,GATA3,GATA4,PLXND1,STAT3,STAT5A,BMP6,TSC22D1,NFATC2,NFE2L2,NFIC,NFKB1,NFKBIA,NHLH2,NOTCH1,NPAS2,T

CD109,SOX6,ZNF703,CIB1,HIPK2,ITGA8,APP,RAB14,DKK1,SNAI2,PRDM16,ATP2B4,RIPK1,LRRC32,GATA3,GATA4,PBLD,IQGAP1,CFLAR,BMP6,NFKB1,NOTCH1,TCF4,CAV1,RUNX2,TGFBR2,THBS1,FURIN,TNFRSF1B,CDH5,CDKN2B,ELAPOR2,HHEX,PDE2A,PDE3A,WNT4,LRIT3,HOXA13,SLIT2,PIK3CA,HES1,PLCG1,VEGFC,COL3A1,COL

ABL2,ETS1,MMP15,FN1,RGCC,APBB2,ITGA8,APP,SCUBE1,FSCN1,CSGALNACT1,CFLAR,NOTCH1,CAV1,GSN,COL18A1,THBS1,TLL1,FURIN,TNFRSF1A,TNFRSF1B,TNR,PECAM1,JAM2,COL3A1,COL4A1,COL4A2,COL5A2,VWF,COL15A1,COL27A1,SH3PXD2A,SCUBE3,CCN2,ITGAV,ITGB3,PXDN,ADAMTS9,GFOD2,KDR,JAM3,RB1,LAMA4,LAMB3

JAG1,MSN,SCUBE1,ZDHHC21,BMP6,NOTCH1,SOX17,COL18A1,HEY1,HEY2,TNFRSF1A,CDH5,NRG1,PDE2A,PECAM1,HOXA13,ROCK2,COL15A1,CLIC4,MIR21,RAPGEF2,CXCR4,KDR,RTN4,S1PR1

ABL2,ETS1,DLC1,JAG1,FES,VAV3,MERTK,ZNF703,CIB1,FN1,VEZT,RGCC,ITGA8,APP,PDLIM5,MSN,RHOB,CD93,SKAP1,SNAI2,CD274,ZFHX3,LIMCH1,PALLD,ZDHHC21,PAG1,MYH9,DISC1,LRRC32,CD164,GATA3,BCL2,PLXND1,BCL6,LRRC4,PCDH17,STAT5A,IQGAP1,CCN4,LYVE1,BMP6,DST,MYPN,NOTCH1,NT5E,CAV1,GSN,TGFBR2,C

ABCA1,JAG1,AHR,SOX6,FOXO1,CIB1,RGCC,VASH1,APP,HDAC7,DKK1,TRIB3,SNAI2,ZFHX3,SUFU,PRDM16,ATP2B4,RIPK1,GATA2,GATA3,PBLD,CCND1,BCL2,BCL6,STAT3,STAT5A,CFLAR,CCN4,NFATC2,NFE2L2,NFKBIA,NOTCH1,TCF4,NPR1,SOX17,CTDSP1,CAV1,ZFPM2,TGFBR2,THBS1,HEY1,HEY2,TNR,ELAPOR2,HHEX,PRTG,HLX,WN

ABL2,SPNS2,CD109,ETS1,MECOM,PICALM,FZD8,JAG1,SLC9B2,FES,AHR,MLLT3,MERTK,SOX6,CIB1,OSTM1,HIPK2,ST3GAL1,APP,HDAC7,SBNO2,PRDM16,MYH9,RIPK1,CD164,GATA2,GATA3,BCL2,BCL6,IL18R1,STAT3,STAT5A,CCN4,NFATC2,NFE2L2,NBEAL2,NFKB1,NFKBIA,NOTCH1,RIPK3,WWP1,HS1BP3,RUNX2,TGFBR2,THBS1,CD3

ACTA2,ETS1,MGAT5,ATP8A1,ZNF703,CIB1,FN1,APP,HDAC7,RHOB,SNAI2,CD274,TRADD,RAB11A,GATA2,GATA3,BCL2,STAT3,STAT5A,IQGAP1,CCN4,NFE2L2,NOTCH1,GRN,CAV1,TGFBR2,COL18A1,THBS1,FURIN,CDH5,PECAM1,JAM2,RIN2,PLCG1,MAP4K4,VEGFC,ROCK2,BCAR1,ADGRA2,IGF2,IGFBP5,MIR21,PRKCE,RAPGEF2,CDKN2B

ABL2,ETS1,MMP15,FN1,RGCC,APBB2,ITGA8,APP,SCUBE1,FSCN1,CSGALNACT1,CFLAR,NOTCH1,CAV1,GSN,COL18A1,THBS1,TLL1,FURIN,TNFRSF1A,TNFRSF1B,TNR,PECAM1,JAM2,COL3A1,COL4A1,COL4A2,COL5A2,VWF,COL15A1,COL27A1,SH3PXD2A,SCUBE3,CCN2,ITGAV,ITGB3,PXDN,ADAMTS9,GFOD2,KDR,JAM3,RB1,LAMA4,LAMB3

ACTA2,ETS1,ETS2,MECOM,DTX3L,PICALM,MEF2D,MRTFB,ATXN1,JAG1,ARID5B,AHR,AFF1,MLLT3,ZNF319,MAML3,EBF4,SOX6,FOXO1,AAMDC,RGCC,HIPK2,ITGA8,BIRC2,ZNF407,APP,PCGF5,SBNO2,RHOG,ATF6,SKAP1,NR5A2,ZFHX3,SSBP4,PRDM16,ARID4B,RIPK1,GATA2,GATA3,GATA4,PLXND1,STAT3,STAT5A,BMP6,TSC22D1,NFAT

ACTA2,ETS1,ETS2,MECOM,DTX3L,PICALM,MEF2D,MRTFB,ATXN1,JAG1,ARID5B,AHR,AFF1,MLLT3,ZNF319,MAML3,EBF4,SOX6,FOXO1,AAMDC,RGCC,HIPK2,ITGA8,BIRC2,ZNF407,APP,PCGF5,SBNO2,RHOG,ATF6,SKAP1,NR5A2,ZFHX3,SSBP4,PRDM16,ARID4B,RIPK1,GATA2,GATA3,GATA4,PLXND1,STAT3,STAT5A,BMP6,TSC22D1,NFAT

ACTA2,ETS1,ETS2,MECOM,DTX3L,PICALM,MEF2D,MRTFB,ATXN1,JAG1,ARID5B,AHR,AFF1,MLLT3,ZNF319,MAML3,EBF4,SOX6,FOXO1,AAMDC,RGCC,HIPK2,ITGA8,BIRC2,ZNF407,APP,PCGF5,SBNO2,RHOG,ATF6,SKAP1,NR5A2,ZFHX3,SSBP4,PRDM16,ARID4B,RIPK1,GATA2,GATA3,GATA4,PLXND1,STAT3,STAT5A,BMP6,TSC22D1,NFAT

ABL2,ETS1,MMP15,FN1,RGCC,APBB2,ITGA8,APP,SCUBE1,FSCN1,CSGALNACT1,CFLAR,NOTCH1,CAV1,GSN,COL18A1,THBS1,TLL1,FURIN,TNFRSF1A,TNFRSF1B,TNR,PECAM1,JAM2,COL3A1,COL4A1,COL4A2,COL5A2,VWF,COL15A1,COL27A1,SH3PXD2A,SCUBE3,CCN2,ITGAV,ITGB3,PXDN,ADAMTS9,GFOD2,KDR,JAM3,RB1,LAMA4,LAMB3

ACTA2,GREB1L,JAG1,ARID5B,AHR,ANXA4,ITGA8,GATA3,BCL2,PLXND1,IQGAP1,CFLAR,BMP6,TMED10,NOTCH1,SOX17,TNS2,HEY1,CD34,PECAM1,WNT4,SLIT2,HOXB7,HES1,WNT2B,WWTR1,ZBTB16,ILK,MTSS1,ITGB3,CXCR4,FAT4,KIF26B,RDH10,EFNB2,ENPEP,SMAD1,EPHA4,SMAD6,SMAD9

ABL2,ACTA2,DLC1,FES,SNX9,EVL,RGCC,CDC42EP3,RHOB,RHOG,LIMCH1,FSCN1,MYH9,GATA4,IQGAP1,CAV1,COTL1,GSN,PDE4B,PECAM1,TMOD2,WNT4,SLIT2,PIK3CA,AKAP6,ROCK2,GMFG,MIR21,PRKCE,CCN2,ARHGAP18,ILK,MTSS1,SPIRE1,ITGB3,SIGLEC15,CYRIA,FCHSD2,RHOBTB1,JAM3,S1PR1,ACTR2,EDN1,SORBS3,ARHGAP28,

ABL2,SPNS2,CD109,ETS1,MECOM,PICALM,FZD8,JAG1,SLC9B2,FES,AHR,MLLT3,MERTK,SOX6,CIB1,OSTM1,HIPK2,ST3GAL1,APP,HDAC7,SBNO2,CD274,PRDM16,MYH9,RIPK1,CD164,GATA2,GATA3,BCL2,BCL6,IL18R1,STAT3,STAT5A,CCN4,NFATC2,NFE2L2,NBEAL2,NFKB1,NFKBIA,NOTCH1,RIPK3,WWP1,HS1BP3,RUNX2,TGFBR2,THB

ABL2,ACTA2,FZD8,ATP8A1,FES,AHR,VAV3,MERTK,PIBF1,EVL,FN1,RGCC,ST3GAL1,APP,RAB14,HDAC7,SBNO2,MSN,CD93,RHOG,COMMD9,CD274,PAG1,MYH9,LRRC32,GATA2,GATA3,BCL2,BCL6,CREG1,IL18R1,STAT3,STAT5A,IQGAP1,GLB1,NFATC2,NBEAL2,NFKB1,AP1M1,SKAP2,TCF4,DNAJC13,RIPK3,WWP1,GRN,HS1BP3,CAV1,RUN

ABL2,CD109,ETS1,MECOM,PICALM,FZD8,JAG1,SLC9B2,FES,AHR,MLLT3,MERTK,SOX6,CIB1,OSTM1,HIPK2,ST3GAL1,APP,HDAC7,SBNO2,PRDM16,MYH9,RIPK1,CD164,GATA2,GATA3,BCL2,BCL6,IL18R1,STAT3,STAT5A,CCN4,NFATC2,NFE2L2,NBEAL2,NFKBIA,NOTCH1,RIPK3,WWP1,HS1BP3,RUNX2,TGFBR2,THBS1,CD34,CDKN2B,HHE

CD109,ZNF703,HIPK2,ITGA8,DKK1,PRDM16,ATP2B4,GATA3,GATA4,PBLD,CFLAR,NOTCH1,TCF4,CAV1,RUNX2,TGFBR2,THBS1,FURIN,CDH5,CDKN2B,ELAPOR2,HHEX,WNT4,LRIT3,HOXA13,SLIT2,HES1,VEGFC,NIBAN2,ADGRA2,MIR21,RNF111,ILK,ITGB3,KDR,VEPH1,HTRA3,MYOF,SMAD6

ACTA2,CD109,ETS1,VAV3,MERTK,DOCK6,EVL,FN1,SCUBE1,SNAI2,AHNAK2,MYH9,GATA2,GATA3,GATA4,CFLAR,CCN4,DST,NFE2L2,NBEAL2,CAV1,PAPSS2,GSN,ZFPM2,TFPI,TGFBR2,THBS1,CD34,ENTPD1,DGKI,NRG1,PECAM1,WNT4,LYST,TXK,PIK3CA,COL3A1,VWF,GNA14,PRCP,PRKACB,PRKAR2B,PRKCE,PRKCH,ILK,DGKG,ITGB3,IT

RPS6KA2,CD109,MALAT1,ETS1,MECOM,MEF2D,DLC1,ATXN1,JAG1,FES,AHR,VAV3,FOXO1,ZNF703,CIB1,FN1,P3H3,SGK1,RGCC,VASH1,HIPK2,BIRC2,APP,RHOG,NR5A2,SNAI2,CD274,NEAT1,RIPK1,DISC1,LRRC32,CD164,GATA2,GATA3,GATA4,PBLD,CCND1,BCL2,BCL6,STAT3,STAT5A,CFLAR,CCN4,BMP6,TSC22D1,NFATC2,NFKBIA,

ETS1,RGCC,VASH1,HIPK2,RHOB,ATP2B4,GATA2,GATA4,PLXND1,STAT3,STIM1,NFE2L2,TCF4,NPR1,GRN,TGFBR2,THBS1,CD34,TNFRSF1A,CDH5,HHEX,WNT4,CEMIP2,HOXA5,PLCG1,VEGFC,ROCK2,COL4A2,NIBAN2,ADGRA2,IGF2,MIR21,ITGB3,GADD45A,JAK1,PTPRM,ADAMTS9,CXCR4,KDR,RTN4,SMAD1

ABL2,DLC1,FES,SNX9,EVL,SORBS2,CIB1,SGK1,RGCC,FIGNL2,MYO1B,APP,CDC42EP3,RHOB,RHOG,LIMCH1,TPPP3,FSCN1,PHACTR1,MYH9,RIPK1,RAB11A,BCL2,CFLAR,DST,MYPN,CAMSAP2,RIPK3,CAV1,COTL1,GSN,CDH5,PECAM1,TMOD2,WNT4,SLIT2,PIK3CA,COL3A1,ROCK2,COL5A2,CLIP2,GMFG,BCAR1,PALM2AKAP2,MIR21,PRKCE

ABL2,ETS2,MECOM,WLS,DLC1,CLIC5,MMP15,FN1,HIPK2,ITGA8,LRIG3,DKK1,SUFU,GATA2,GATA3,GATA4,NOTCH1,SOX17,RUNX2,TGFBR2,HHEX,HLX,WNT4,HOXA1,HOXA3,HOXA4,HOXA5,HOXA6,HOXA13,HOXB7,HOXB8,HOXC4,HOXC9,HOXD3,HOXD9,PITX2,VAX2,HES1,VEGFC,ROCK2,COL4A2,COL5A2,WNT2B,IGF2,PRKACB,ZBTB16,INH

ABL2,DLC1,FES,SNX9,EVL,RGCC,CDC42EP3,RHOB,RHOG,LIMCH1,FSCN1,MYH9,IQGAP1,COTL1,GSN,PECAM1,TMOD2,WNT4,SLIT2,PIK3CA,ROCK2,GMFG,MIR21,PRKCE,CCN2,ARHGAP18,ILK,MTSS1,SPIRE1,ITGB3,SIGLEC15,CYRIA,FCHSD2,RHOBTB1,JAM3,S1PR1,ACTR2,EDN1,SORBS3,ARHGAP28,PHLDB2

DTX3L,JAG1,MAML3,SORBS2,BMP2K,ANXA4,AAK1,APP,SNAI2,GATA2,PLXND1,BCL6,STAT3,NFKBIA,NOTCH1,TGFBR2,HEY1,HEY2,TSPAN15,HHEX,NRARP,HOXD3,HES1,NIBAN2,JAG2,GDPD5,TSPAN14,FAT4,RCAN2

ETS1,CIB1,RGCC,VASH1,HDAC7,RHOB,SNAI2,ATP2B4,GATA2,GATA3,STAT5A,NFE2L2,NOTCH1,GRN,THBS1,SLIT2,RIN2,PLCG1,VEGFC,ROCK2,BCAR1,ADGRA2,IGF2,PRCP,MIR21,ITGB3,GADD45A,PTPRM,ADAMTS9,KDR,EDN1

SPNS2,ETS2,MEF2D,ARID5B,SLC9B2,SOX6,SNAI2,SUFU,CSGALNACT1,CCN4,BMP6,NBEAL2,RUNX2,PAPSS2,TGFBR2,THBS1,TLL1,HOXA1,HOXA3,HOXA4,HOXA5,HOXA6,HOXA13,HOXB7,HOXB8,HOXC4,HOXC6,HOXC8,HOXC9,HOXD3,HOXD8,HOXD9,PIP4K2A,PITX2,CREB3L2,COL3A1,COL5A2,COL27A1,WNT2B,IGF2,MIR21,CCN2,ZBTB16,

DLC1,JAG1,MERTK,CIB1,FN1,ITGA8,SKAP1,LIMCH1,DISC1,BCL2,BCL6,IQGAP1,LYVE1,NOTCH1,THBS1,CD34,PECAM1,PARVB,WNT4,HOXD3,RIN2,MAP4K4,VEGFC,COL3A1,ROCK2,VWF,BCAR1,PRKCE,CCN2,ILK,ITGAV,ITGB3,ADAMTS9,KDR,JAM3,DUSP3,LAMB3,LAMC1,SORBS3,PHLDB2,SMAD6

ETS1,RGCC,VASH1,HIPK2,RHOB,ATP2B4,GATA2,GATA4,PLXND1,STAT3,STIM1,NFE2L2,TCF4,NPR1,GRN,TGFBR2,THBS1,CD34,TNFRSF1A,CDH5,HHEX,CEMIP2,HOXA5,PLCG1,VEGFC,ROCK2,COL4A2,NIBAN2,ADGRA2,IGF2,MIR21,ITGB3,GADD45A,JAK1,PTPRM,ADAMTS9,CXCR4,KDR,RTN4,SMAD1

TBC1D19,ABL2,QRICH2,PICALM,MRTFB,DLC1,CLIC5,HECW2,FES,VAV3,PIBF1,EVL,CIB1,FN1,SGK1,APBB2,ITGA8,APP,CPNE5,CDC42EP3,PDLIM5,RHOG,DKK1,FSCN1,CFAP44,PALLD,PHACTR1,MYH9,MYO9B,UNC5B,DISC1,TTC21A,SDCCAG8,RAB11A,TBC1D2B,GATA3,BCL2,PLXND1,CDKL1,IQGAP1,CFLAR,DST,NFE2L2,MYPN,CAMSAP

CD109,AHR,ATP6V0D2,SOX6,FOXO1,ZNF703,ADCY4,CIB1,SGK1,HIPK2,ITGA8,APP,RAB14,DKK1,TRIB3,NR5A2,SNAI2,PRDM16,ATP2B4,LRRC32,GATA3,GATA4,PBLD,STAT3,STAT5A,IQGAP1,CFLAR,BMP6,NFE2L2,NFKB1,NOTCH1,ABHD2,TNS2,CAV1,RUNX2,TFPI,CCND3,TGFBR2,THBS1,HEY1,FURIN,PALM,CDH5,CDKN2B,ELAPOR2,P

JAG1,SNAI2,GATA3,GATA4,NOTCH1,HEY1,HEY2,TNFRSF1A,TNFRSF1B,SLIT2,ROCK2,RB1,SMAD6

DLC1,JAG1,CIB1,FN1,ITGA8,SKAP1,LIMCH1,DISC1,BCL2,BCL6,IQGAP1,LYVE1,THBS1,CD34,PECAM1,WNT4,HOXD3,RIN2,MAP4K4,VEGFC,COL3A1,ROCK2,CCN2,ILK,ITGAV,ITGB3,ADAMTS9,KDR,JAM3,DUSP3,PHLDB2

ETS2,WLS,MLLT3,HIPK2,DKK1,GATA4,NOTCH1,SOX17,HEY1,HEY2,HHEX,HOXA1,HOXA3,NRARP,HOXA4,HOXA5,HOXA6,HOXB7,HOXB8,HOXC4,HOXC6,HOXC8,HOXC9,HOXD3,HOXD8,HOXD9,HES1,WNT2B,ZBTB16,ZEB2

ABL2,ETS1,DLC1,JAG1,FES,VAV3,ZNF703,CIB1,FN1,RGCC,SKAP1,SNAI2,CD274,ZFHX3,LIMCH1,ZDHHC21,PAG1,DISC1,LRRC32,CD164,GATA3,BCL2,PLXND1,BCL6,STAT5A,IQGAP1,BMP6,NOTCH1,CAV1,GSN,TGFBR2,THBS1,TNR,HLX,WNT4,JAM2,NRARP,PIK3CA,RIN2,HES1,MAP4K4,VEGFC,ROCK2,BCAR1,IGF2,MIR21,PRKCE,IL4

GREB1L,JAG1,GATA3,BCL2,IQGAP1,NOTCH1,HEY1,CD34,PECAM1,WNT4,SLIT2,HOXB7,HES1,WNT2B,WWTR1,ILK,MTSS1,FAT4,KIF26B,EFNB2,SMAD1,EPHA4,SMAD6,SMAD9

ACTA2,ETS1,ETS2,MECOM,DTX3L,PICALM,MEF2D,MRTFB,ATXN1,JAG1,ARID5B,AHR,AFF1,MLLT3,ZNF319,MAML3,EBF4,SOX6,FOXO1,AAMDC,RGCC,HIPK2,ITGA8,BIRC2,ZNF407,APP,PCGF5,SBNO2,RHOG,ATF6,SKAP1,NR5A2,ZFHX3,SSBP4,PRDM16,SAMD4A,ARID4B,RIPK1,GATA2,GATA3,GATA4,PLXND1,STAT3,STAT5A,BMP6,TSC22

ABL2,DLC1,FES,SNX9,EVL,CIB1,SGK1,RGCC,CDC42EP3,RHOB,RHOG,LIMCH1,FSCN1,MYH9,IQGAP1,DST,CAMSAP2,CAV1,COTL1,GSN,AFAP1,RASSF1,CDH5,PECAM1,TMOD2,WNT4,SLIT2,PIK3CA,ROCK2,GMFG,CLIC4,MIR21,PRKCE,CCN2,ARHGAP18,ILK,MTSS1,SPIRE1,ITGB3,SIGLEC15,CYRIA,FCHSD2,RHOBTB1,JAM3,S1PR1,ACTR2

JAG1,SNAI2,GATA3,GATA4,NOTCH1,HEY2,TNFRSF1A,TNFRSF1B,SLIT2,ROCK2,RB1,SMAD6

ABL2,PICALM,MRTFB,CLIC5,HECW2,FES,EVL,CIB1,FN1,SGK1,APBB2,APP,CPNE5,PDLIM5,RHOG,DKK1,PALLD,PHACTR1,UNC5B,DISC1,RAB11A,GATA3,BCL2,PLXND1,IQGAP1,CFLAR,DST,NFE2L2,MYPN,CAMSAP2,NOTCH1,KALRN,SZT2,GRN,FRYL,HEY1,CD38,TTL,PALM,SEPTIN7,TNR,NRG1,PRTG,RPS6KA5,MGLL,HOXA1,SLIT2,PIK3

HDAC7,GATA4,NOTCH1,SOX17,CAV1,ZFPM2,TGFBR2,HEY1,CD34,HEY2,HHEX,HOXA13,RIN2,PITX2,QKI,RAPGEF2,ITGAV,KDR,RTN4

TBC1D19,ABL2,PICALM,MRTFB,DLC1,CLIC5,HECW2,FES,VAV3,PIBF1,EVL,CIB1,FN1,SGK1,APBB2,APP,CPNE5,CDC42EP3,PDLIM5,RHOG,DKK1,FSCN1,CFAP44,PALLD,PHACTR1,MYH9,MYO9B,UNC5B,DISC1,TTC21A,SDCCAG8,RAB11A,TBC1D2B,GATA3,BCL2,PLXND1,CDKL1,IQGAP1,CFLAR,DST,NFE2L2,MYPN,CAMSAP2,NOTCH1,KALR

ABHD6,AHR,MERTK,PIBF1,SOX6,FOXO1,EVL,FN1,ANXA4,RGCC,VASH1,APP,HDAC7,DKK1,SNAI2,CD274,PRDM16,ATP2B4,RIPK1,LRRC32,GATA2,GATA3,GATA4,PBLD,BCL2,BCL6,IL18R1,CCN4,NFKB1,NOTCH1,TCF4,NPR1,SOX17,CTDSP1,GRN,ZFPM2,TFPI,TGFBR2,THBS1,HEY1,CD34,FURIN,CD38,HEY2,IL1RL1,TNFRSF1A,TNFRSF1

ETS1,CIB1,HDAC7,RHOB,SNAI2,GATA2,GATA3,STAT5A,NFE2L2,GRN,THBS1,RIN2,PLCG1,VEGFC,ROCK2,BCAR1,ADGRA2,IGF2,MIR21,ITGB3,KDR,EDN1

MEF2D,MRTFB,MAPK12,JAG1,ARID5B,SOX6,FLNB,SORBS2,ITGA8,HDAC7,PDLIM5,SMTN,DKK1,ZFHX3,MYH9,CD164,GATA4,BCL2,STIM1,CFLAR,CCN4,NFATC2,MYPN,NOTCH1,CAV1,ZFPM2,HEY1,GPCPD1,HEY2,NRG1,HLX,TMOD2,WNT4,JAM2,HOXD9,PITX2,PLAGL1,HES1,QKI,AKAP6,COL3A1,NIBAN2,IGF2,IGFBP3,IGFBP5,MIR21,IL4

ABL2,ATXN1,ARID5B,FES,VAV3,ATP6V0D2,MERTK,FOXO1,RAB14,TRADD,GATA3,STAT3,STAT5A,IQGAP1,SHKBP1,NOTCH1,TCF4,KALRN,TNS2,RUNX2,CCND3,THBS1,NRG1,HHEX,RPS6KA5,WNT4,LRIT3,AP2S1,PIK3CA,PIP4K2A,PLCG1,VEGFC,COL4A1,ROCK2,COL4A2,NIBAN2,BCAR1,ADGRA2,IGF2,IGFBP3,IGFBP5,CLEC14A,MIR21,C

JAG1,SNAI2,GATA3,GATA4,NOTCH1,ZFPM2,TGFBR2,HEY1,HEY2,SLIT2,ROCK2,ADAMTS9,RB1,SMAD6

TBC1D19,ABCA1,DLC1,FES,AHR,SNX9,FNIP2,EVL,SGK1,RGCC,BIRC2,APP,CDC42EP3,PDLIM5,MSN,RHOG,DKK1,SKAP1,SNAI2,LIMCH1,TPPP3,FSCN1,PALLD,MYH9,SDCCAG8,TBC1D2B,CDKL1,IQGAP1,CAMSAP2,NOTCH1,CAV1,COTL1,GSN,TBC1D8,THBS1,PALM,TNFRSF1A,SEPTIN7,CDH5,NRG1,PECAM1,TMOD2,RABGAP1,WNT4,ZDHHC1

ABCA1,CD109,AHR,ATP6V0D2,SOX6,FOXO1,ZNF703,ADCY4,CIB1,SGK1,HIPK2,ITGA8,APP,RAB14,DKK1,TRIB3,NR5A2,SNAI2,ZFHX3,CATSPERG,PRDM16,ATP2B4,LRRC32,GATA3,GATA4,PBLD,CCND1,BCL2,STAT3,STAT5A,IQGAP1,CFLAR,BMP6,GLB1,NFE2L2,NFKB1,NOTCH1,ABHD2,TNS2,CAV1,RUNX2,TFPI,CCND3,TGFBR2,THBS1,

ABL2,RPS6KA2,ETS1,MECOM,PICALM,DLC1,AHR,VAV3,MERTK,FOXO1,CIB1,FN1,PHLDA1,AAMDC,SGK1,ANXA4,RGCC,APBB2,HIPK2,BIRC2,NLRP1,ST3GAL1,APP,HDAC7,RHOB,ATF6,DKK1,SNAI2,CD274,ACSL5,TRADD,RIPK1,UNC5B,GATA2,GATA3,GATA4,BCL2,TNFRSF10D,BCL6,STAT3,STAT5A,CFLAR,CCN4,TSC22D1,IER3,NFE2L2,

ABCA1,JAG1,AHR,SOX6,FOXO1,CIB1,APP,HDAC7,DKK1,TRIB3,SNAI2,ZFHX3,SUFU,PRDM16,GATA2,GATA3,PBLD,CCND1,BCL6,STAT3,STAT5A,CCN4,NFATC2,NFE2L2,NFKBIA,NOTCH1,CTDSP1,CAV1,ZFPM2,HEY1,HEY2,TNR,PRTG,HLX,WNT4,NRARP,HOXA5,SLIT2,HOXB8,GORASP1,HES1,VEGFC,COL5A2,WWTR1,BCAR1,IGF2,IGFBP5,

PRR5L,ABL2,RPS6KA2,CD109,FZD8,MAPK12,FES,MERTK,PIBF1,SNX9,FNIP2,ADCY4,CIB1,CDC42BPA,FN1,BMP2K,SGK1,RGCC,AAK1,HIPK2,CDKL4,ST3GAL1,APP,DKK1,TRIB3,LIMCH1,ATP2B4,RIPK1,GATA4,PBLD,CCND1,BCL2,CDKL1,STAT3,STAT5A,IQGAP1,MLKL,BMP6,DCLK2,NPR1,RIPK3,KALRN,CTDSP1,CAV1,CCND3,PKIG,TG

ABL2,GREB1L,DLC1,AHR,RHOB,SUFU,GATA3,GATA4,BCL2,PLXND1,NOTCH1,SOX17,TGFBR2,HEY1,HHEX,WNT4,NRARP,HOXA5,SLIT2,HOXB7,PITX2,HES1,COL4A1,ROCK2,WNT2B,CLIC4,PRKACB,MIR21,ILK,MTSS1,ZEB2,CXCR4,KDR,FAT4,KIF26B,RDH10,EDN1,EFNB2,EPHA4

ABL2,CD109,FZD8,SLC9B2,FES,AHR,MERTK,OSTM1,ST3GAL1,APP,HDAC7,SBNO2,PRDM16,MYH9,RIPK1,GATA2,GATA3,BCL2,BCL6,IL18R1,STAT3,STAT5A,CCN4,NFATC2,RIPK3,WWP1,HS1BP3,RUNX2,TGFBR2,HHEX,HLX,WNT4,TXK,NRARP,KLF6,POU2F2,SH3PXD2A,MIR21,IL4R,IRF2BP2,ZBTB16,INHBA,ITGB3,SIGLEC15,GAB2,JAG

ABCA4,ABHD6,SCTR,FOXO1,ADCY4,PIWIL4,PRDM16,RAB11A,GATA2,PBLD,BCL2,IL18R1,STAT3,BMP6,NOTCH1,TNS2,CAV1,CD34,CD38,CDH5,PECAM1,JAM2,HOXA13,CLCN3,PIK3CA,TPP1,CRTC3,WWTR1,PRCP,ZNF516,PRKACB,PRKAR2B,CCN2,IL4R,ITGAV,ITGB3,SIGLEC15,CXCR4,SLC27A1,KDR,JAM3,RB1,LAMA4,LAMC1,CUBN,S1P

ABL2,PICALM,MRTFB,CLIC5,HECW2,FES,EVL,CIB1,FN1,SGK1,APBB2,APP,CPNE5,PDLIM5,RHOG,DKK1,PALLD,PHACTR1,SPR,UNC5B,DISC1,RAB11A,GATA2,GATA3,BCL2,PLXND1,IQGAP1,CFLAR,DST,NFE2L2,MYPN,DCLK2,CAMSAP2,NOTCH1,KALRN,SZT2,GRN,FRYL,HEY1,CD38,TTL,PALM,SEPTIN7,TNR,NRG1,PRTG,RPS6KA5,MGLL,

FMNL2,DLC1,FES,FN1,CDC42EP3,MSN,RHOB,RHOG,MYH9,PLXND1,FMNL3,PALM,SEPTIN7,PARVB,PALM2AKAP2,SYNE3,MIR21,ARHGAP18,PALMD,RHOBTB1,KDR,PLEKHO1,EPB41L2

VAV3,MERTK,DOCK6,EVL,FN1,SCUBE1,MYH9,GATA2,GATA3,GATA4,NFE2L2,NBEAL2,CAV1,PAPSS2,ZFPM2,TFPI,THBS1,CD34,ENTPD1,DGKI,LYST,TXK,PIK3CA,COL3A1,VWF,GNA14,PRCP,PRKACB,PRKAR2B,PRKCE,PRKCH,ILK,DGKG,ITGB3,ITPR1,EDN1

ABL2,FZD8,ATP8A1,FES,AHR,VAV3,MERTK,PIBF1,ST3GAL1,APP,RAB14,HDAC7,SBNO2,MSN,CD93,RHOG,COMMD9,CD274,PAG1,MYH9,LRRC32,GATA2,GATA3,BCL2,BCL6,CREG1,IL18R1,STAT3,STAT5A,IQGAP1,GLB1,NFATC2,NBEAL2,NFKB1,AP1M1,SKAP2,DNAJC13,RIPK3,WWP1,GRN,HS1BP3,CAV1,RUNX2,COTL1,GSN,CCND3,TGFBR

ACTA2,GREB1L,JAG1,AHR,GATA3,BCL2,IQGAP1,CFLAR,NOTCH1,HEY1,CD34,PECAM1,WNT4,HOXB7,HES1,WNT2B,WWTR1,ILK,MTSS1,ITGB3,FAT4,KIF26B,ENPEP

ABL2,GREB1L,DLC1,JAG1,AHR,MLLT3,ANKRD6,MSN,RHOB,DKK1,SNAI2,SUFU,GATA3,GATA4,BCL2,PLXND1,STAT5A,NOTCH1,SOX17,TGFBR2,HEY1,HHEX,WNT4,NRARP,HOXA5,HOXA13,SLIT2,HOXB7,AP2S1,PITX2,HES1,VEGFC,COL4A1,ROCK2,WNT2B,CLIC4,CRYGS,IGFBP5,PRKACB,MIR21,ILK,MTSS1,ZEB2,JAG2,CXCR4,KDR,FAT4,

FMNL2,PICALM,DLC1,FES,CIB1,FN1,SGK1,CPNE5,CDC42EP3,MSN,RHOB,RHOG,MYH9,PLXND1,KALRN,FMNL3,PALM,SEPTIN7,PARVB,BCAR1,PALM2AKAP2,SYNE3,MIR21,ARHGAP18,ILK,PALMD,ITGB3,RHOBTB1,CXCR4,KDR,RTN4,ACTR2,PLEKHO1,EPB41L2,EPHA4

ABL2,DLC1,SNX9,EVL,SORBS2,RGCC,MYO1B,CDC42EP3,RHOB,RHOG,LIMCH1,FSCN1,PHACTR1,MYH9,BCL2,COTL1,GSN,PECAM1,TMOD2,WNT4,SLIT2,PIK3CA,ROCK2,GMFG,BCAR1,PALM2AKAP2,MIR21,PRKCE,CCN2,ARHGAP18,MTSS1,SPIRE1,CYRIA,FCHSD2,RHOBTB1,ARHGAP25,S1PR1,ACTR2,SORBS3,ARHGAP28,MARCKS,PHLDB2

VAV3,MERTK,DOCK6,EVL,FN1,SCUBE1,MYH9,GATA2,GATA3,GATA4,NFE2L2,NBEAL2,CAV1,PAPSS2,ZFPM2,TFPI,THBS1,CD34,ENTPD1,DGKI,LYST,TXK,PIK3CA,COL3A1,VWF,GNA14,PRCP,PRKACB,PRKAR2B,PRKCE,PRKCH,ILK,DGKG,ITGB3,ITPR1,EDN1

PRR5L,ABL2,CD109,ABHD6,MECOM,DLC1,ATXN1,MLLT3,PIBF1,FOXO1,ADCY4,CIB1,HIPK2,APP,ANKRD6,HDAC7,DKK1,TRIB3,SNAI2,SUFU,PRDM16,ATP2B4,TRADD,RIPK1,UNC5B,GATA2,GATA3,GATA4,PBLD,BCL2,BCL6,PCDH17,CFLAR,IER3,NFE2L2,NFKBIA,NOTCH1,KALRN,SZT2,SOX17,TNS2,CAV1,RUNX2,TGFBR2,THBS1,HEY1,C

VAV3,MERTK,DOCK6,EVL,FN1,SCUBE1,MYH9,GATA2,GATA3,GATA4,NFE2L2,NBEAL2,CAV1,PAPSS2,ZFPM2,TFPI,THBS1,CD34,ENTPD1,DGKI,LYST,TXK,PIK3CA,COL3A1,VWF,GNA14,PRCP,PRKACB,PRKAR2B,PRKCE,PRKCH,ILK,DGKG,ITGB3,ITPR1,EDN1

PRR5L,ABL2,CD109,ABHD6,MECOM,DLC1,ATXN1,MLLT3,PIBF1,FOXO1,ADCY4,CIB1,HIPK2,APP,ANKRD6,HDAC7,DKK1,TRIB3,SNAI2,SUFU,PRDM16,ATP2B4,TRADD,RIPK1,UNC5B,GATA2,GATA3,GATA4,PBLD,BCL2,BCL6,PCDH17,CFLAR,IER3,NFE2L2,NFKBIA,NOTCH1,KALRN,SZT2,SOX17,TNS2,CAV1,RUNX2,TGFBR2,THBS1,HEY1,C

| | | | | | | | | |
|---|---|---|---|---|---|---|---|---|
| GO: Biological Process | GO:0048667 | cell morphogenesis involved in neuron differentiation | 2.02E-08 | 1.42E-04 | 1.20E-06 | 1.13E-05 | 55 | 715 |
| GO: Biological Process | GO:0071559 | response to transforming growth factor beta | 2.13E-08 | 1.50E-04 | 1.25E-06 | 1.18E-05 | 31 | 289 |
| GO: Biological Process | GO:0060429 | epithelium development | 2.17E-08 | 1.53E-04 | 1.26E-06 | 1.19E-05 | 94 | 1532 |
| GO: Biological Process | GO:1902903 | regulation of supramolecular fiber organization | 3.48E-08 | 2.45E-04 | 2.00E-06 | 1.89E-05 | 38 | 412 |
| GO: Biological Process | GO:0009967 | positive regulation of signal transduction | 4.16E-08 | 2.92E-04 | 2.38E-06 | 2.24E-05 | 102 | 1733 |
| GO: Biological Process | GO:0048858 | cell projection morphogenesis | 4.25E-08 | 2.99E-04 | 2.39E-06 | 2.25E-05 | 58 | 790 |
| GO: Biological Process | GO:0009792 | embryo development ending in birth or egg hatching | 4.25E-08 | 2.99E-04 | 2.39E-06 | 2.25E-05 | 61 | 850 |
| GO: Biological Process | GO:0001952 | regulation of cell-matrix adhesion | 4.63E-08 | 3.26E-04 | 2.58E-06 | 2.44E-05 | 20 | 137 |
| GO: Biological Process | GO:0003180 | aortic valve morphogenesis | 4.86E-08 | 3.42E-04 | 2.69E-06 | 2.54E-05 | 10 | 31 |
| GO: Biological Process | GO:0150115 | cell-substrate junction organization | 6.36E-08 | 4.47E-04 | 3.49E-06 | 3.29E-05 | 18 | 114 |
| GO: Biological Process | GO:0035904 | aorta development | 7.35E-08 | 5.17E-04 | 4.00E-06 | 3.78E-05 | 14 | 69 |
| GO: Biological Process | GO:0030099 | myeloid cell differentiation | 7.52E-08 | 5.29E-04 | 4.07E-06 | 3.84E-05 | 42 | 496 |
| GO: Biological Process | GO:0061448 | connective tissue development | 7.95E-08 | 5.59E-04 | 4.27E-06 | 4.03E-05 | 31 | 306 |
| GO: Biological Process | GO:0120039 | plasma membrane bounded cell projection morphogenesis | 8.12E-08 | 5.71E-04 | 4.33E-06 | 4.08E-05 | 57 | 785 |
| GO: Biological Process | GO:0043009 | chordate embryonic development | 8.78E-08 | 6.17E-04 | 4.64E-06 | 4.38E-05 | 59 | 827 |
| GO: Biological Process | GO:0003002 | regionalization | 8.90E-08 | 6.26E-04 | 4.67E-06 | 4.41E-05 | 36 | 392 |
| GO: Biological Process | GO:0003007 | heart morphogenesis | 9.44E-08 | 6.64E-04 | 4.92E-06 | 4.64E-05 | 30 | 292 |
| GO: Biological Process | GO:0008593 | regulation of Notch signaling pathway | 9.58E-08 | 6.73E-04 | 4.93E-06 | 4.65E-05 | 18 | 117 |
| GO: Biological Process | GO:0032990 | cell part morphogenesis | 9.61E-08 | 6.76E-04 | 4.93E-06 | 4.65E-05 | 58 | 809 |
| GO: Biological Process | GO:0033043 | regulation of organelle organization | 1.04E-07 | 7.32E-04 | 5.30E-06 | 5.00E-05 | 82 | 1317 |
| GO: Biological Process | GO:0003281 | ventricular septum development | 1.08E-07 | 7.60E-04 | 5.47E-06 | 5.16E-05 | 15 | 82 |
| GO: Biological Process | GO:0032989 | cellular component morphogenesis | 1.09E-07 | 7.68E-04 | 5.49E-06 | 5.18E-05 | 63 | 914 |
| GO: Biological Process | GO:1904018 | positive regulation of vasculature development | 1.13E-07 | 7.97E-04 | 5.61E-06 | 5.30E-05 | 24 | 201 |
| GO: Biological Process | GO:0045766 | positive regulation of angiogenesis | 1.13E-07 | 7.97E-04 | 5.61E-06 | 5.30E-05 | 24 | 201 |
| GO: Biological Process | GO:0090066 | regulation of anatomical structure size | 1.25E-07 | 8.81E-04 | 6.16E-06 | 5.81E-05 | 47 | 599 |
| GO: Biological Process | GO:0060548 | negative regulation of cell death | 1.26E-07 | 8.87E-04 | 6.16E-06 | 5.81E-05 | 77 | 1214 |
| GO: Biological Process | GO:0060537 | muscle tissue development | 1.33E-07 | 9.36E-04 | 6.46E-06 | 6.09E-05 | 41 | 488 |
| GO: Biological Process | GO:0071560 | cellular response to transforming growth factor beta stimulus | 1.51E-07 | 1.06E-03 | 7.26E-06 | 6.85E-05 | 29 | 282 |
| GO: Biological Process | GO:0034330 | cell junction organization | 1.52E-07 | 1.07E-03 | 7.26E-06 | 6.85E-05 | 58 | 820 |
| GO: Biological Process | GO:0007044 | cell-substrate junction assembly | 1.54E-07 | 1.08E-03 | 7.33E-06 | 6.91E-05 | 17 | 108 |
| GO: Biological Process | GO:0048585 | negative regulation of response to stimulus | 1.67E-07 | 1.17E-03 | 7.88E-06 | 7.44E-05 | 109 | 1945 |
| GO: Biological Process | GO:0072009 | nephron epithelium development | 2.09E-07 | 1.47E-03 | 9.78E-06 | 9.23E-05 | 18 | 123 |
| GO: Biological Process | GO:0032835 | glomerulus development | 2.20E-07 | 1.55E-03 | 1.03E-05 | 9.68E-05 | 14 | 75 |
| GO: Biological Process | GO:0007178 | transmembrane receptor protein serine/threonine kinase signaling pathway | 2.26E-07 | 1.59E-03 | 1.04E-05 | 9.85E-05 | 36 | 407 |
| GO: Biological Process | GO:0048812 | neuron projection morphogenesis | 2.36E-07 | 1.66E-03 | 1.08E-05 | 1.02E-04 | 55 | 770 |
| GO: Biological Process | GO:0003177 | pulmonary valve development | 2.51E-07 | 1.77E-03 | 1.15E-05 | 1.08E-04 | 8 | 21 |
| GO: Biological Process | GO:0043067 | regulation of programmed cell death | 2.65E-07 | 1.87E-03 | 1.20E-05 | 1.14E-04 | 101 | 1777 |
| GO: Biological Process | GO:0040007 | growth | 2.70E-07 | 1.90E-03 | 1.22E-05 | 1.15E-04 | 74 | 1172 |
| GO: Biological Process | GO:0048589 | developmental growth | 2.95E-07 | 2.07E-03 | 1.32E-05 | 1.25E-04 | 57 | 816 |
| GO: Biological Process | GO:1901700 | response to oxygen-containing compound | 3.04E-07 | 2.14E-03 | 1.35E-05 | 1.28E-04 | 105 | 1875 |
| GO: Biological Process | GO:0031344 | regulation of cell projection organization | 3.51E-07 | 2.47E-03 | 1.55E-05 | 1.47E-04 | 56 | 800 |
| GO: Biological Process | GO:0050878 | regulation of body fluid levels | 3.55E-07 | 2.50E-03 | 1.56E-05 | 1.47E-04 | 44 | 563 |
| GO: Biological Process | GO:0045667 | regulation of osteoblast differentiation | 3.66E-07 | 2.58E-03 | 1.60E-05 | 1.51E-04 | 20 | 155 |
| GO: Biological Process | GO:0003151 | outflow tract morphogenesis | 3.85E-07 | 2.71E-03 | 1.67E-05 | 1.57E-04 | 15 | 90 |
| GO: Biological Process | GO:0120035 | regulation of plasma membrane bounded cell projection organization | 3.86E-07 | 2.72E-03 | 1.67E-05 | 1.57E-04 | 55 | 782 |
| GO: Biological Process | GO:0048568 | embryonic organ development | 3.92E-07 | 2.76E-03 | 1.68E-05 | 1.59E-04 | 42 | 527 |
| GO: Biological Process | GO:0043534 | blood vessel endothelial cell migration | 4.09E-07 | 2.87E-03 | 1.74E-05 | 1.64E-04 | 22 | 185 |
| GO: Biological Process | GO:0090092 | regulation of transmembrane receptor protein serine/threonine kinase signaling pathway | 4.40E-07 | 3.09E-03 | 1.86E-05 | 1.76E-04 | 28 | 280 |
| GO: Biological Process | GO:0110053 | regulation of actin filament organization | 4.85E-07 | 3.41E-03 | 2.04E-05 | 1.93E-04 | 29 | 298 |
| GO: Biological Process | GO:0060840 | artery development | 5.02E-07 | 3.53E-03 | 2.10E-05 | 1.98E-04 | 17 | 117 |
| GO: Biological Process | GO:0003231 | cardiac ventricle development | 5.05E-07 | 3.55E-03 | 2.10E-05 | 1.98E-04 | 19 | 144 |
| GO: Biological Process | GO:0042981 | regulation of apoptotic process | 5.61E-07 | 3.94E-03 | 2.32E-05 | 2.19E-04 | 98 | 1736 |
| GO: Biological Process | GO:0042692 | muscle cell differentiation | 5.85E-07 | 4.12E-03 | 2.41E-05 | 2.27E-04 | 38 | 460 |
| GO: Biological Process | GO:0043085 | positive regulation of catalytic activity | 5.94E-07 | 4.17E-03 | 2.43E-05 | 2.29E-04 | 92 | 1600 |
| GO: Biological Process | GO:0061326 | renal tubule development | 7.30E-07 | 5.13E-03 | 2.97E-05 | 2.80E-04 | 16 | 107 |
| GO: Biological Process | GO:2001212 | regulation of vasculogenesis | 7.78E-07 | 5.47E-03 | 3.15E-05 | 2.97E-04 | 7 | 17 |
| GO: Biological Process | GO:0051247 | positive regulation of protein metabolic process | 8.14E-07 | 5.72E-03 | 3.27E-05 | 3.09E-04 | 96 | 1704 |
| GO: Biological Process | GO:0030855 | epithelial cell differentiation | 8.49E-07 | 5.97E-03 | 3.39E-05 | 3.20E-04 | 60 | 906 |
| GO: Biological Process | GO:0090100 | positive regulation of transmembrane receptor protein serine/threonine kinase signaling pathway | 9.20E-07 | 6.47E-03 | 3.65E-05 | 3.45E-04 | 17 | 122 |
| GO: Biological Process | GO:0061564 | axon development | 9.38E-07 | 6.60E-03 | 3.71E-05 | 3.50E-04 | 45 | 603 |
| GO: Biological Process | GO:0003013 | circulatory system process | 1.00E-06 | 7.04E-03 | 3.93E-05 | 3.71E-04 | 50 | 704 |
| GO: Biological Process | GO:0048732 | gland development | 1.01E-06 | 7.07E-03 | 3.93E-05 | 3.71E-04 | 41 | 527 |
| GO: Biological Process | GO:0000122 | negative regulation of transcription by RNA polymerase II | 1.02E-06 | 7.14E-03 | 3.95E-05 | 3.72E-04 | 64 | 996 |
| GO: Biological Process | GO:0048754 | branching morphogenesis of an epithelial tube | 1.10E-06 | 7.76E-03 | 4.26E-05 | 4.02E-04 | 22 | 196 |
| GO: Biological Process | GO:0010647 | positive regulation of cell communication | 1.15E-06 | 8.07E-03 | 4.41E-05 | 4.16E-04 | 106 | 1952 |
| GO: Biological Process | GO:1903053 | regulation of extracellular matrix organization | 1.19E-06 | 8.38E-03 | 4.55E-05 | 4.30E-04 | 11 | 52 |
| GO: Biological Process | GO:0023056 | positive regulation of signaling | 1.26E-06 | 8.88E-03 | 4.80E-05 | 4.53E-04 | 106 | 1956 |
| GO: Biological Process | GO:0003279 | cardiac septum development | 1.30E-06 | 9.15E-03 | 4.92E-05 | 4.64E-04 | 17 | 125 |
| GO: Biological Process | GO:0007163 | establishment or maintenance of cell polarity | 1.35E-06 | 9.47E-03 | 5.06E-05 | 4.78E-04 | 25 | 246 |
| GO: Biological Process | GO:0050673 | epithelial cell proliferation | 1.42E-06 | 9.99E-03 | 5.31E-05 | 5.01E-04 | 40 | 515 |
| GO: Biological Process | GO:2000146 | negative regulation of cell motility | 1.62E-06 | 1.14E-02 | 6.03E-05 | 5.69E-04 | 33 | 387 |

ABL2,PICALM,CLIC5,HECW2,EVL,FN1,SGK1,APBB2,APP,PDLIM5,RHOG,PALLD,PHACTR1,SPR,UNC5B,DISC1,RAB11A,GATA3,BCL2,PLXND1,DST,MYPN,NOTCH1,KALRN,SZT2,TTL,SEPTIN7,TNR,NRG1,PRTG,RPS6KA5,MGLL,HOXA1,SLIT2,PIK3CA,GORASP1,VAX2,HES1,PLCG1,ROCK2,NIBAN2,FEZ2,RAPGEF2,TANC2,ILK,ZEB2,[illegible]

CD109,SOX6,ZNF703,HIPK2,ITGA8,ZFHX3,PRDM16,LRRC32,PBLD,CFLAR,CAV1,TGFBR2,THBS1,FURIN,CDH5,CDKN2B,PDE2A,PDE3A,WNT4,COL3A1,ROCK2,COL4A2,MIR21,RNF111,JUN,VEPH1,HTRA3,EDN1,SMAD1,SMAD6,SMAD9

ABL2,ACTA2,CD109,MECOM,GREB1L,DLC1,CLIC5,JAG1,AHR,MLLT3,ZNF703,FLNB,ANXA4,ANKRD6,MSN,RHOB,SCUBE1,DKK1,GSTK1,NR5A2,SNAI2,SUFU,PALLD,ZDHHC21,TRADD,ARID4B,GATA3,GATA4,CCND1,BCL2,PLXND1,STAT5A,IQGAP1,BMP6,NOTCH1,SOX17,CAV1,TGFBR2,COL18A1,HEY1,CD34,FURIN,HEY2,TNFRSF1A,CDH5,[illegible]

DLC1,FES,SNX9,EVL,CIB1,SGK1,RGCC,APP,CDC42EP3,LIMCH1,MYH9,CAMSAP2,CAV1,COTL1,GSN,CDH5,PECAM1,TMOD2,WNT4,SLIT2,PIK3CA,ROCK2,GMFG,MIR21,PRKCE,CCN2,ARHGAP18,MTSS1,SPIRE1,CYRIA,FCHSD2,RB1,S1PR1,ACTR2,EDN1,SORBS3,ARHGAP28,PHLDB2

PRR5L,ABL2,ACTA2,WLS,FZD8,MGAT5,JAG1,USP12,SCTR,MLLT3,RELL1,PIBF1,MAML3,CIB1,FN1,BMP2K,AAK1,HIPK2,ITGA8,BIRC2,APP,ANKRD6,RHOG,ATF6,SCUBE1,DKK1,KLHDC10,SH3RF3,TRADD,RIPK1,UNC5B,DISC1,GATA3,GATA4,BCL2,IL18R1,DENND2B,STAT3,IQGAP1,SHKBP1,CFLAR,CCN4,MLKL,BMP6,NFKB1,SERINC3,[illegible]

ABL2,PICALM,HECW2,EVL,FN1,SGK1,APBB2,APP,CPNE5,PDLIM5,RHOG,PALLD,PHACTR1,MYO9B,UNC5B,DISC1,RAB11A,GATA3,BCL2,PLXND1,IQGAP1,DST,MYPN,NOTCH1,KALRN,SZT2,TTL,SEPTIN7,TNR,NRG1,PRTG,RPS6KA5,MGLL,HOXA1,SLIT2,PIK3CA,GORASP1,VAX2,PLCG1,MAP4K4,ROCK2,NIBAN2,FEZ2,RAPGEF2,TANC2,ILK,[illegible]

ABL2,SLC25A34,MECOM,MRTFB,DLC1,SOX6,VEZT,VASH1,DKK1,NR5A2,SUFU,MYH9,GATA2,GATA3,GATA4,NOTCH1,SOX17,GRN,RUNX2,ZFPM2,TGFBR2,HEY1,FURIN,HEY2,HHEX,HOXA1,HOXA3,NRARP,HOXA4,HOXA5,HOXA6,SLIT2,HOXB7,HOXB8,HOXC4,HOXC6,HOXC9,HOXD3,HOXD9,PITX2,HES1,ABCG2,PLCG1,ROCK2,IGF2,SH3PXD2A,[illegible]

DLC1,JAG1,CIB1,SKAP1,LIMCH1,DISC1,BCL2,BCL6,IQGAP1,THBS1,WNT4,RIN2,MAP4K4,VEGFC,ROCK2,ILK,ITGB3,KDR,DUSP3,PHLDB2

JAG1,SNAI2,GATA3,GATA4,NOTCH1,HEY2,SLIT2,ROCK2,RB1,SMAD6

DLC1,FN1,LIMCH1,BCL2,IQGAP1,DST,THBS1,WNT4,MAP4K4,ROCK2,ITGAV,ITGB3,IQSEC1,KDR,DUSP3,LAMB3,LAMC1,PHLDB2

JAG1,SUFU,PLXND1,NOTCH1,HEY1,HEY2,PDE2A,HES1,COL3A1,ADAMTS9,CXCR4,KAT6A,EFNB2,SMAD6

CD109,ETS1,JAG1,SLC9B2,FES,SOX6,CIB1,OSTM1,HIPK2,APP,SBNO2,PRDM16,MYH9,RIPK1,GATA2,GATA3,BCL6,STAT3,STAT5A,CCN4,NBEAL2,NFKBIA,TGFBR2,THBS1,CDKN2B,HHEX,HOXA5,HOXB7,HOXB8,PIP4K2A,SH3PXD2A,ZBTB16,INHBA,SNRK,ITGB3,SIGLEC15,GAB2,JUN,RB1,KAT6A,LMO2,EPAS1

ACTA2,MEF2D,ARID5B,SOX6,SNAI2,CSGALNACT1,CFLAR,CCN4,BMP6,NOTCH1,RUNX2,TGFBR2,THBS1,CD34,HOXA3,HOXA5,HOXC4,HOXD3,PIK3CA,CREB3L2,COL27A1,WNT2B,ZNF516,MIR21,CCN2,ZBTB16,ITGB3,EDN1,LMNA,SMAD1,SMAD9

ABL2,PICALM,HECW2,EVL,FN1,SGK1,APBB2,APP,CPNE5,PDLIM5,RHOG,PALLD,PHACTR1,MYO9B,UNC5B,DISC1,RAB11A,GATA3,BCL2,PLXND1,IQGAP1,DST,MYPN,NOTCH1,KALRN,SZT2,TTL,SEPTIN7,TNR,NRG1,PRTG,RPS6KA5,MGLL,HOXA1,SLIT2,PIK3CA,GORASP1,VAX2,PLCG1,MAP4K4,ROCK2,NIBAN2,FEZ2,RAPGEF2,TANC2,ILK,[illegible]

ABL2,SLC25A34,MECOM,MRTFB,DLC1,SOX6,VEZT,VASH1,DKK1,SUFU,MYH9,GATA2,GATA3,GATA4,NOTCH1,SOX17,GRN,RUNX2,ZFPM2,TGFBR2,HEY1,FURIN,HEY2,HHEX,HOXA1,HOXA3,NRARP,HOXA4,HOXA5,HOXA6,SLIT2,HOXB7,HOXB8,HOXC4,HOXC6,HOXC9,HOXD3,HOXD9,PITX2,HES1,PLCG1,ROCK2,IGF2,SH3PXD2A,PRKACB,[illegible]

ETS2,WLS,MLLT3,HIPK2,DKK1,SUFU,GATA4,NOTCH1,SOX17,HEY1,HEY2,HHEX,HOXA1,HOXA3,NRARP,HOXA4,HOXA5,HOXA6,HOXB7,HOXB8,HOXC4,HOXC6,HOXC8,HOXC9,HOXD3,HOXD8,HOXD9,PITX2,VAX2,HES1,WNT2B,PRKACB,ZBTB16,ZEB2,EDN1,SMAD6

MRTFB,DLC1,JAG1,AHR,DKK1,SNAI2,SUFU,GATA3,GATA4,PLXND1,NOTCH1,SOX17,ZFPM2,TGFBR2,THBS1,HEY1,HEY2,NRG1,SLIT2,PITX2,HES1,MIR21,ILK,JUN,FAT4,RTN4,KAT6A,S1PR1,ADGRG6,SMAD6

JAG1,MAML3,BMP2K,AAK1,GATA2,BCL6,STAT3,NFKBIA,NOTCH1,HEY1,HEY2,TSPAN15,NRARP,HES1,NIBAN2,JAG2,GDPD5,TSPAN14

ABL2,PICALM,HECW2,EVL,FN1,SGK1,APBB2,APP,CPNE5,PDLIM5,RHOG,PALLD,PHACTR1,MYO9B,UNC5B,DISC1,RAB11A,GATA3,BCL2,PLXND1,IQGAP1,DST,MYPN,NOTCH1,KALRN,SZT2,TTL,SEPTIN7,TNR,NRG1,PRTG,RPS6KA5,MGLL,HOXA1,SLIT2,PIK3CA,GORASP1,VAX2,PLCG1,MAP4K4,ROCK2,NIBAN2,FEZ2,RAPGEF2,TANC2,ILK,[illegible]

TBC1D19,ABL2,DLC1,HECW2,FES,SNX9,EVL,CIB1,SGK1,RGCC,APP,CDC42EP3,MSN,RHOB,RHOG,SNAI2,LIMCH1,FSCN1,PALLD,MYH9,SDCCAG8,TBC1D2B,GATA3,BCL2,BCL6,CDKL1,IQGAP1,DST,IER3,CAMSAP2,CLEC16A,GRN,CAV1,COTL1,GSN,TBC1D8,AFAP1,RASSF1,SEPTIN7,CDH5,PDE2A,PDE3A,RPS6KA5,PECAM1,TMOD2,[illegible]

SUFU,GATA3,GATA4,NOTCH1,ZFPM2,TGFBR2,HEY1,HEY2,PDE2A,HOXA13,SLIT2,PITX2,HES1,CXCR4,SMAD6

ABL2,PICALM,HECW2,EVL,FN1,SGK1,APBB2,APP,CPNE5,PDLIM5,RHOG,PALLD,PHACTR1,MYO9B,UNC5B,DISC1,RAB11A,GATA3,BCL2,PLXND1,IQGAP1,CFLAR,DST,MYPN,NOTCH1,KALRN,SZT2,TTL,SEPTIN7,TNR,NRG1,PRTG,RPS6KA5,TMOD2,MGLL,HOXA1,SLIT2,PIK3CA,GORASP1,VAX2,PLCG1,MAP4K4,ROCK2,NIBAN2,FEZ2,[illegible]

ETS1,HIPK2,RHOB,GATA2,GATA4,STAT3,STIM1,NFE2L2,GRN,TGFBR2,THBS1,CD34,TNFRSF1A,CDH5,PLCG1,VEGFC,IGF2,MIR21,ITGB3,JAK1,CXCR4,KDR,RTN4,SMAD1

ETS1,HIPK2,RHOB,GATA2,GATA4,STAT3,STIM1,NFE2L2,GRN,TGFBR2,THBS1,CD34,TNFRSF1A,CDH5,PLCG1,VEGFC,IGF2,MIR21,ITGB3,JAK1,CXCR4,KDR,RTN4,SMAD1

RIN3,ACTA2,PICALM,AHR,SCTR,VAV3,SNX9,EVL,FN1,CDC42EP3,MSN,ZDHHC21,MYH9,DISC1,RAB11A,TBXAS1,NPR1,CAV1,COTL1,GSN,CD38,TTL,TNR,NRG1,PECAM1,TMOD2,MGLL,SLIT2,CLCN3,PIK3CA,ROCK2,GMFG,MIR21,PRKCE,ARHGAP18,DOCK4,ILK,SPIRE1,CYRIA,FCHSD2,PTPRM,RTN4,ECE1,ACTR2,EDN1,ARHGAP28,SMAD6

ABL2,MECOM,AHR,MERTK,FOXO1,CIB1,FN1,AAMDC,SGK1,ANXA4,APBB2,HIPK2,BIRC2,ST3GAL1,APP,HDAC7,DKK1,SNAI2,TRADD,RIPK1,UNC5B,GATA2,GATA3,GATA4,BCL2,TNFRSF10D,BCL6,STAT3,STAT5A,CFLAR,CCN4,TSC22D1,IER3,NFE2L2,NFKB1,NFKBIA,NOTCH1,NPAS2,GPX4,GRN,CAV1,ZFPM2,THBS1,CD34,CD38,NANOS3,[illegible]

MEF2D,MRTFB,AHR,SOX6,FLNB,SORBS2,ITGA8,HDAC7,PDLIM5,DKK1,RIPK1,GATA4,BCL2,CFLAR,CCN4,NOTCH1,CAV1,ZFPM2,TGFBR2,HEY1,GPCPD1,HEY2,NRG1,HLX,HOXD9,PITX2,PLAGL1,AKAP6,COL3A1,NIBAN2,IGF2,IGFBP5,VGLL4,ADAMTS9,KCNAB1,RB1,S1PR1,EDN1,EFNB2,LMNA,SMAD1

CD109,SOX6,ZNF703,HIPK2,ITGA8,PRDM16,LRRC32,PBLD,CFLAR,CAV1,TGFBR2,THBS1,FURIN,CDH5,CDKN2B,PDE2A,PDE3A,WNT4,COL3A1,COL4A2,MIR21,RNF111,JUN,VEPH1,HTRA3,EDN1,SMAD1,SMAD6,SMAD9

ABL2,MALAT1,PICALM,DLC1,ZNF703,SORBS2,FN1,APBB2,APP,HDAC7,PDLIM5,DKK1,SNAI2,LIMCH1,FSCN1,PALLD,FILIP1,DISC1,BCL2,PLXND1,LRRC4,PCDH17,IQGAP1,BMP6,DST,KALRN,CAV1,THBS1,PALM,RASSF8,SEPTIN7,TNR,CDH5,NRG1,CDHR3,PECAM1,WNT4,MAP4K4,COL4A1,ROCK2,SRGN,PRKCH,RAPGEF2,TANC2,ILK,[illegible]

DLC1,FN1,LIMCH1,BCL2,IQGAP1,DST,THBS1,WNT4,MAP4K4,ROCK2,ITGAV,ITGB3,KDR,DUSP3,LAMB3,LAMC1,PHLDB2

RIN3,PRR5L,ABL2,CD109,ETS1,MECOM,DLC1,ATXN1,MLLT3,PIBF1,FOXO1,ADCY4,CIB1,HIPK2,APP,ANKRD6,HDAC7,DKK1,TRIB3,SNAI2,SUFU,PRDM16,ATP2B4,TRADD,RIPK1,UNC5B,GATA2,GATA3,GATA4,PBLD,BCL2,BCL6,CFLAR,IER3,NFE2L2,NFKB1,NFKBIA,NOTCH1,SZT2,NT5E,SOX17,TNS2,GRN,CAV1,RUNX2,TFPI,TGFBR2,[illegible]

GREB1L,JAG1,GATA3,BCL2,IQGAP1,NOTCH1,HEY1,CD34,PECAM1,WNT4,HOXB7,HES1,WNT2B,WWTR1,ILK,MTSS1,FAT4,KIF26B

ACTA2,JAG1,AHR,BCL2,IQGAP1,CFLAR,NOTCH1,CD34,PECAM1,HES1,WWTR1,MTSS1,ITGB3,ENPEP

CD109,ZNF703,HIPK2,ITGA8,DKK1,PRDM16,LRRC32,GATA4,PBLD,BMP6,NOTCH1,CAV1,RUNX2,TGFBR2,THBS1,FURIN,CDH5,CDKN2B,ELAPOR2,HOXA13,HES1,COL3A1,WWTR1,PALM2AKAP2,MIR21,RNF111,ILK,INHBA,INHBB,JUN,KDR,VEPH1,HTRA3,SMAD1,SMAD6,SMAD9

ABL2,PICALM,HECW2,EVL,FN1,SGK1,APBB2,APP,CPNE5,PDLIM5,RHOG,PALLD,PHACTR1,UNC5B,DISC1,RAB11A,GATA3,BCL2,PLXND1,IQGAP1,DST,MYPN,NOTCH1,KALRN,SZT2,TTL,SEPTIN7,TNR,NRG1,PRTG,RPS6KA5,MGLL,HOXA1,SLIT2,PIK3CA,GORASP1,VAX2,PLCG1,MAP4K4,ROCK2,NIBAN2,FEZ2,RAPGEF2,TANC2,ILK,[illegible]

JAG1,NOTCH1,HEY1,HEY2,TNFRSF1A,TNFRSF1B,SLIT2,SMAD6

ABL2,RPS6KA2,ETS1,MECOM,DLC1,VAV3,MERTK,FOXO1,CIB1,FN1,PHLDA1,AAMDC,SGK1,ANXA4,RGCC,APBB2,HIPK2,BIRC2,NLRP1,ST3GAL1,APP,RHOB,ATF6,DKK1,SNAI2,CD274,ACSL5,TRADD,RIPK1,UNC5B,GATA2,GATA3,GATA4,BCL2,TNFRSF10D,BCL6,STAT5A,CFLAR,TSC22D1,IER3,NFE2L2,NFKB1,NFKBIA,SERINC3,RASSF3,[illegible]

ABL2,PICALM,ARID5B,AHR,SORBS2,CIB1,FN1,SGK1,APBB2,APP,CPNE5,PDLIM5,RIPK1,SPR,DISC1,RAB11A,GATA3,GATA4,BCL2,BCL6,CREG1,STAT3,STAT5A,IQGAP1,CFLAR,CCN4,NOTCH1,NPR1,SOX17,TNS2,GSN,ZFPM2,TGFBR2,ZMAT3,THBS1,CD38,TTL,HEY2,SEPTIN7,TNR,NRG1,HHEX,HLX,MGLL,TXK,HOXA5,SLIT2,PIK3CA,[illegible]

ABL2,PICALM,ARID5B,AHR,SORBS2,FN1,APP,CPNE5,PDLIM5,RIPK1,SPR,DISC1,RAB11A,GATA3,GATA4,BCL2,STAT3,STAT5A,IQGAP1,CFLAR,CCN4,NOTCH1,TNS2,GSN,ZFPM2,TGFBR2,THBS1,TTL,HEY2,SEPTIN7,TNR,NRG1,HHEX,HLX,MGLL,TXK,HOXA5,SLIT2,PIK3CA,AKAP6,COL27A1,CLIC4,WWTR1,IGF2,VGLL4,ILK,ZEB2,[illegible]

ABCA1,ABL2,ETS1,MGST2,JAG1,SLC9B2,FES,AHR,ATP6V0D2,MMP15,FOXO1,ZNF703,ADCY4,SORBS2,SGK1,BIRC2,PIWIL4,NLRP1,APP,SBNO2,MSN,RHOB,DKK1,TRIB3,SNAI2,CD274,CATSPERG,ATP2B4,RIPK1,GATA3,GATA4,CCND1,BCL2,STAT3,STAT5A,IQGAP1,CFLAR,FCSK,BMP6,GLB1,GNG12,NFE2L2,NFKB1,NFKBIA,NOTCH1,[illegible]

TBC1D19,ABL2,PICALM,HECW2,FES,EVL,CIB1,FN1,SGK1,APP,CDC42EP3,PDLIM5,RHOG,DKK1,FSCN1,DISC1,SDCCAG8,RAB11A,TBC1D2B,GATA3,PLXND1,CDKL1,IQGAP1,CFLAR,NFE2L2,CAMSAP2,KALRN,GRN,CAV1,TBC1D8,CD38,PALM,SEPTIN7,TNR,NRG1,RABGAP1,SLIT2,PIK3CA,GORASP1,HES1,QKI,MAP4K4,CREB3L2,MIR21,RA

SCTR,VAV3,MERTK,DOCK6,EVL,ADCY4,FN1,RAB14,SCUBE1,MYH9,RAB11A,GATA2,GATA3,GATA4,CCND1,STAT5A,NFE2L2,NBEAL2,NPR1,KALRN,CAV1,PAPSS2,ZFPM2,TFPI,THBS1,CD34,ENTPD1,DGKI,LYST,TXK,PIK3CA,COL3A1,VWF,GNA14,PRCP,PRKACB,PRKAR2B,PRKCE,PRKCH,ILK,DGKG,ITGB3,ITPR1,EDN1

JAG1,AHR,HDAC7,SNAI2,SUFU,ZHX3,CCN4,BMP6,NOTCH1,RUNX2,HEY1,WNT4,VEGFC,WWTR1,IGFBP5,MIR21,ILK,LMNA,SMAD1,SMAD6

JAG1,GATA4,PLXND1,NOTCH1,SOX17,ZFPM2,TGFBR2,THBS1,HEY1,HEY2,PITX2,HES1,ILK,JUN,SMAD6

TBC1D19,ABL2,PICALM,HECW2,FES,EVL,CIB1,FN1,SGK1,APP,CDC42EP3,PDLIM5,RHOG,DKK1,FSCN1,DISC1,SDCCAG8,RAB11A,TBC1D2B,GATA3,PLXND1,CDKL1,IQGAP1,CFLAR,NFE2L2,CAMSAP2,KALRN,GRN,CAV1,TBC1D8,CD38,PALM,SEPTIN7,TNR,NRG1,RABGAP1,SLIT2,PIK3CA,GORASP1,HES1,QKI,MAP4K4,CREB3L2,RAPGEF2,

MRTFB,CLIC5,AHR,VASH1,HIPK2,ITGA8,LRIG3,SUFU,GATA2,GATA3,GATA4,NOTCH1,SOX17,RUNX2,ZFPM2,TGFBR2,HEY1,HEY2,HHEX,HLX,HOXA1,HOXA3,HOXA4,HOXA5,HOXA6,HOXB7,HOXB8,HOXC4,HOXC9,HOXD3,HOXD9,PITX2,VAX2,HES1,IGF2,ITGAV,KDR,KAT6A,RDH10,EDN1,LMO2,EPAS1

ETS1,CIB1,RGCC,VASH1,HDAC7,ATP2B4,MYH9,GATA2,STAT5A,NFE2L2,NOTCH1,THBS1,CDH5,SLIT2,PLCG1,VEGFC,IGF2,PRCP,CLEC14A,GADD45A,KDR,EFNB2

CD109,ZNF703,HIPK2,ITGA8,DKK1,PRDM16,GATA4,PBLD,BMP6,NOTCH1,CAV1,TGFBR2,THBS1,FURIN,CDH5,CDKN2B,ELAPOR2,HOXA13,HES1,MIR21,RNF111,ILK,INHBA,INHBB,KDR,VEPH1,HTRA3,SMAD6

DLC1,SNX9,EVL,RGCC,CDC42EP3,LIMCH1,MYH9,COTL1,GSN,PECAM1,TMOD2,WNT4,SLIT2,PIK3CA,ROCK2,GMFG,MIR21,PRKCE,CCN2,ARHGAP18,MTSS1,SPIRE1,CYRIA,FCHSD2,S1PR1,ACTR2,SORBS3,ARHGAP28,PHLDB2

JAG1,SUFU,PLXND1,NOTCH1,HEY1,HEY2,PDE2A,HOXA1,HOXA13,HES1,COL3A1,ADAMTS9,CXCR4,RTN4,KAT6A,EFNB2,SMAD6

GREB1L,JAG1,AHR,SUFU,GATA3,GATA4,NOTCH1,ZFPM2,TGFBR2,HEY1,HEY2,NRG1,PDE2A,HOXA13,SLIT2,PITX2,HES1,CXCR4,SMAD6

ABL2,RPS6KA2,ETS1,DLC1,VAV3,MERTK,FOXO1,CIB1,FN1,PHLDA1,AAMDC,SGK1,ANXA4,RGCC,APBB2,HIPK2,BIRC2,NLRP1,ST3GAL1,APP,RHOB,ATF6,DKK1,SNAI2,CD274,ACSL5,TRADD,RIPK1,UNC5B,GATA2,GATA3,GATA4,BCL2,TNFRSF10D,BCL6,STAT5A,CFLAR,TSC22D1,IER3,NFE2L2,NFKB1,NFKBIA,SERINC3,RASSF3,NOTCH1

MRTFB,MAPK12,SOX6,SORBS2,ITGA8,PDLIM5,DKK1,MYH9,GATA4,BCL2,STIM1,CFLAR,CCN4,NFATC2,MYPN,NOTCH1,HEY1,HEY2,NRG1,TMOD2,WNT4,JAM2,PITX2,HES1,QKI,AKAP6,NIBAN2,IGF2,IGFBP5,MIR21,IL4R,RB1,EDN1,MYOF,EFNB2,LMNA,PLEKHO1,SMAD6

TBC1D19,RIN3,ABL2,DTX3L,PICALM,FZD8,DLC1,MAPK12,ARHGAP31,VAV3,MERTK,PIBF1,SNX9,ADCY4,CIB1,FN1,RGCC,NLRP1,APP,RHOG,DKK1,TRIB3,ATP2B4,TRADD,RIPK1,MYO9B,TBC1D2B,CCND1,BCL2,STAT3,IQGAP1,STIM1,CFLAR,MLKL,DNAJC1,RIPK3,KALRN,ARHGEF12,GRN,CAV1,GSN,CCND3,TBC1D8,TGFBR2,THBS1,MAP4

GREB1L,JAG1,GATA3,BCL2,NOTCH1,HEY1,WNT4,HOXB7,HES1,COL4A1,WNT2B,WWTR1,ILK,MTSS1,FAT4,KIF26B

HEY1,CD34,HEY2,RIN2,RAPGEF2,KDR,RTN4

PRR5L,PICALM,FZD8,DLC1,MAPK12,HECW2,PIBF1,SNX9,FNIP2,FOXO1,ADCY4,CIB1,FN1,RGCC,HIPK2,BIRC2,NLRP1,APP,MSN,DKK1,TRIB3,SNAI2,LIMCH1,ATP2B4,SAMD4A,TRADD,MYH9,RIPK1,DISC1,GATA3,GATA4,CCND1,BCL2,BCL6,STAT3,IQGAP1,CFLAR,MLKL,BMP6,IER3,NFE2L2,NFKBIA,RIPK3,WWP1,SOX17,GRN,CAV1,GS

ABL2,ACTA2,CD109,CLIC5,JAG1,ZNF703,FLNB,ANXA4,MSN,SCUBE1,GSTK1,NR5A2,PALLD,ZDHHC21,ARID4B,GATA3,GATA4,CCND1,STAT5A,IQGAP1,BMP6,NOTCH1,SOX17,CAV1,COL18A1,HEY1,CD34,FURIN,HEY2,TNFRSF1A,CDH5,NRG1,HHEX,PDE2A,PECAM1,WNT4,HOXA5,HOXA13,TPP1,PITX2,HES1,COL4A1,ROCK2,COL15A1,CLIC

HIPK2,ITGA8,GATA4,BMP6,NOTCH1,THBS1,FURIN,CDH5,CDKN2B,ELAPOR2,HES1,MIR21,RNF111,ILK,INHBA,INHBB,KDR

PICALM,EVL,FN1,APBB2,APP,RHOG,PALLD,UNC5B,DISC1,RAB11A,GATA3,BCL2,PLXND1,DST,MYPN,CAMSAP2,NOTCH1,KALRN,SZT2,GRN,TTL,SEPTIN7,TNR,NRG1,PRTG,RPS6KA5,MGLL,HOXA1,SLIT2,PIK3CA,VAX2,PLCG1,NIBAN2,FEZ2,ILK,ZEB2,GAB2,JUN,PTPRM,CXCR4,JAM3,RTN4,ABLIM1,EFNB2,EPHA4

ACTA2,ATP8A1,AHR,SGK1,APP,SMTN,ATP2B4,SCN3B,ZDHHC21,GATA4,STIM1,BMP6,IER3,TBXAS1,NPR1,CAV1,GSN,TBC1D8,CD34,CD38,HEY2,CDH5,PDE2A,PDE3A,PDE4B,SLIT2,PIK3CA,ABCG2,VEGFC,AKAP6,ROCK2,WWTR1,CACNA2D4,PRCP,MIR21,CCN2,DOCK4,SLC6A17,ITPR1,PTPRM,CXCR4,SLC27A1,KCNE3,ECE1,LEPR,EDN1,M

ETS1,WLS,MRTFB,ARID5B,AHR,ZNF703,MSN,SNAI2,ZDHHC21,GATA2,GATA3,CCND1,BCL2,PLXND1,STAT5A,CFLAR,NOTCH1,RIPK3,KALRN,CAV1,TG,TGFBR2,NRG1,HHEX,HLX,WNT4,HOXA3,HOXA5,HOXA13,SLIT2,HOXD3,HOXD9,PIK3CA,PITX2,HES1,VWF,IGF2,IGFBP5,ITPR1,JUN,RTN4

ETS2,ATXN1,ARID5B,AHR,FNIP2,SOX6,HIPK2,APP,HDAC7,DKK1,TRIB3,SNAI2,ZFHX3,SUFU,PRDM16,ZHX3,GATA2,GATA3,CCND1,BCL6,STAT3,BMP6,NFATC2,NFIC,NFKB1,NOTCH1,TCF4,CAV1,ZFPM2,CCND3,PKIG,HEY1,HEY2,KLF12,HHEX,HIC1,PDE2A,NRARP,HOXB8,HOXC8,HOXD8,HOXD9,PITX2,PLAGL1,VAX2,HES1,WWTR1,CREM

GREB1L,AHR,BCL2,PLXND1,NOTCH1,TGFBR2,HHEX,WNT4,NRARP,HOXA5,SLIT2,HOXB7,PITX2,COL4A1,WNT2B,CLIC4,ILK,CXCR4,KDR,FAT4,RDH10,EDN1

PRR5L,ABL2,ACTA2,WLS,FZD8,MGAT5,JAG1,USP12,SCTR,MLLT3,RELL1,PIBF1,MAML3,CIB1,FN1,BMP2K,AAK1,HIPK2,ITGA8,BIRC2,APP,ANKRD6,RHOG,ATF6,SCUBE1,DKK1,KLHDC10,SH3RF3,TRADD,RIPK1,UNC5B,DISC1,GATA3,GATA4,BCL2,IL18R1,DENND2B,STAT3,IQGAP1,SHKBP1,CFLAR,CCN4,MLKL,BMP6,NFKB1,SERINC3,C

ABL2,ETS1,RGCC,FSCN1,CFLAR,NOTCH1,TNFRSF1A,TNFRSF1B,ITGB3,RB1,PHLDB2

PRR5L,ABL2,ACTA2,WLS,FZD8,MGAT5,JAG1,USP12,SCTR,MLLT3,RELL1,PIBF1,MAML3,CIB1,FN1,BMP2K,AAK1,HIPK2,ITGA8,BIRC2,APP,ANKRD6,RHOG,ATF6,SCUBE1,DKK1,KLHDC10,SH3RF3,TRADD,RIPK1,UNC5B,DISC1,GATA3,GATA4,BCL2,IL18R1,DENND2B,STAT3,IQGAP1,SHKBP1,CFLAR,CCN4,MLKL,BMP6,NFKB1,SERINC3,C

JAG1,SUFU,GATA3,GATA4,PLXND1,NOTCH1,ZFPM2,TGFBR2,HEY1,HEY2,PDE2A,HOXA13,SLIT2,PITX2,HES1,CXCR4,SMAD6

AMOTL2,MSN,RHOB,RHOG,FRMD4A,FSCN1,MYH9,MYO9B,SDCCAG8,GATA3,FRMD4B,DST,GSN,HEY1,CDH5,PARVB,HES1,ROCK2,CLIC4,ILK,RHOBTB1,JAM3,KIF26B,ACTR2,LMNA

CD109,ZNF703,RGCC,VASH1,SNAI2,GATA2,GATA3,PBLD,CCND1,STAT3,STAT5A,CFLAR,BMP6,NOTCH1,GRN,CAV1,THBS1,HEY1,CD34,CDKN2B,NRARP,HOXA5,HES1,PLCG1,VEGFC,IGF2,IGFBP3,IGFBP5,MIR21,MTSS1,ITGB3,ITPR1,JUN,PTPRM,KDR,RB1,RTN4,LAMC1,EFNB2,SERPINB1

RIN3,ABHD6,DLC1,JAG1,EVL,RGCC,VASH1,RHOB,LIMCH1,ATP2B4,GATA3,PBLD,BCL2,STAT3,NFE2L2,NOTCH1,ABHD2,THBS1,NRG1,WNT4,SLIT2,COL3A1,CLIC4,IGFBP3,IGFBP5,MIR21,ILK,DACH1,GADD45A,PTPRM,ADAMTS9,DUSP3,PHLDB2

| | | | | | | | | |
|---|---|---|---|---|---|---|---|---|
| GO: Biological Process | GO:0014897 | striated muscle hypertrophy | 1.63E-06 | 1.15E-02 | 6.03E-05 | 5.69E-04 | 17 | 127 |
| GO: Biological Process | GO:0061138 | morphogenesis of a branching epithelium | 1.72E-06 | 1.21E-02 | 6.33E-05 | 5.97E-04 | 24 | 233 |
| GO: Biological Process | GO:0032270 | positive regulation of cellular protein metabolic process | 1.81E-06 | 1.27E-02 | 6.61E-05 | 6.24E-04 | 91 | 1619 |
| GO: Biological Process | GO:0030336 | negative regulation of cell migration | 1.82E-06 | 1.28E-02 | 6.64E-05 | 6.27E-04 | 32 | 371 |
| GO: Biological Process | GO:0014706 | striated muscle tissue development | 1.91E-06 | 1.34E-02 | 6.92E-05 | 6.53E-04 | 37 | 464 |
| GO: Biological Process | GO:0001823 | mesonephros development | 1.96E-06 | 1.38E-02 | 7.05E-05 | 6.65E-04 | 16 | 115 |
| GO: Biological Process | GO:0014896 | muscle hypertrophy | 2.03E-06 | 1.43E-02 | 7.28E-05 | 6.87E-04 | 17 | 129 |
| GO: Biological Process | GO:0051130 | positive regulation of cellular component organization | 2.05E-06 | 1.44E-02 | 7.30E-05 | 6.89E-04 | 77 | 1304 |
| GO: Biological Process | GO:0008285 | negative regulation of cell population proliferation | 2.09E-06 | 1.47E-02 | 7.39E-05 | 6.97E-04 | 58 | 889 |
| GO: Biological Process | GO:0001763 | morphogenesis of a branching structure | 2.09E-06 | 1.47E-02 | 7.39E-05 | 6.97E-04 | 25 | 252 |
| GO: Biological Process | GO:0043087 | regulation of GTPase activity | 2.11E-06 | 1.48E-02 | 7.39E-05 | 6.98E-04 | 39 | 504 |
| GO: Biological Process | GO:0001503 | ossification | 2.11E-06 | 1.49E-02 | 7.39E-05 | 6.98E-04 | 37 | 466 |
| GO: Biological Process | GO:0050678 | regulation of epithelial cell proliferation | 2.20E-06 | 1.55E-02 | 7.67E-05 | 7.24E-04 | 36 | 448 |
| GO: Biological Process | GO:1905245 | regulation of aspartic-type peptidase activity | 2.22E-06 | 1.56E-02 | 7.69E-05 | 7.26E-04 | 6 | 13 |
| GO: Biological Process | GO:0060284 | regulation of cell development | 2.24E-06 | 1.58E-02 | 7.74E-05 | 7.30E-04 | 44 | 603 |
| GO: Biological Process | GO:0072080 | nephron tubule development | 2.27E-06 | 1.60E-02 | 7.80E-05 | 7.36E-04 | 15 | 103 |
| GO: Biological Process | GO:0007389 | pattern specification process | 2.32E-06 | 1.63E-02 | 7.91E-05 | 7.47E-04 | 39 | 506 |
| GO: Biological Process | GO:0009968 | negative regulation of signal transduction | 2.36E-06 | 1.66E-02 | 8.03E-05 | 7.58E-04 | 87 | 1537 |
| GO: Biological Process | GO:0007423 | sensory organ development | 2.86E-06 | 2.01E-02 | 9.67E-05 | 9.12E-04 | 47 | 669 |
| GO: Biological Process | GO:0031399 | regulation of protein modification process | 2.87E-06 | 2.02E-02 | 9.67E-05 | 9.12E-04 | 99 | 1825 |
| GO: Biological Process | GO:0001932 | regulation of protein phosphorylation | 2.93E-06 | 2.06E-02 | 9.81E-05 | 9.26E-04 | 79 | 1362 |
| GO: Biological Process | GO:0051271 | negative regulation of cellular component movement | 2.98E-06 | 2.10E-02 | 9.94E-05 | 9.38E-04 | 33 | 398 |
| GO: Biological Process | GO:0003171 | atrioventricular valve development | 3.10E-06 | 2.18E-02 | 1.03E-04 | 9.70E-04 | 8 | 28 |
| GO: Biological Process | GO:0140352 | export from cell | 3.13E-06 | 2.20E-02 | 1.03E-04 | 9.75E-04 | 88 | 1571 |
| GO: Biological Process | GO:0003205 | cardiac chamber development | 3.18E-06 | 2.24E-02 | 1.05E-04 | 9.86E-04 | 21 | 193 |
| GO: Biological Process | GO:0048705 | skeletal system morphogenesis | 3.20E-06 | 2.25E-02 | 1.05E-04 | 9.87E-04 | 25 | 258 |
| GO: Biological Process | GO:0007409 | axonogenesis | 3.22E-06 | 2.27E-02 | 1.05E-04 | 9.90E-04 | 41 | 552 |
| GO: Biological Process | GO:0032535 | regulation of cellular component size | 3.45E-06 | 2.43E-02 | 1.12E-04 | 1.06E-03 | 35 | 438 |
| GO: Biological Process | GO:0045892 | negative regulation of transcription, DNA-templated | 3.50E-06 | 2.46E-02 | 1.13E-04 | 1.07E-03 | 82 | 1437 |
| GO: Biological Process | GO:0001885 | endothelial cell development | 3.55E-06 | 2.49E-02 | 1.14E-04 | 1.07E-03 | 13 | 81 |
| GO: Biological Process | GO:0043254 | regulation of protein-containing complex assembly | 3.77E-06 | 2.65E-02 | 1.21E-04 | 1.14E-03 | 38 | 497 |
| GO: Biological Process | GO:1903507 | negative regulation of nucleic acid-templated transcription | 3.79E-06 | 2.67E-02 | 1.21E-04 | 1.14E-03 | 82 | 1440 |
| GO: Biological Process | GO:0019220 | regulation of phosphate metabolic process | 3.84E-06 | 2.70E-02 | 1.22E-04 | 1.15E-03 | 97 | 1790 |
| GO: Biological Process | GO:0051174 | regulation of phosphorus metabolic process | 3.93E-06 | 2.77E-02 | 1.24E-04 | 1.17E-03 | 97 | 1791 |
| GO: Biological Process | GO:0046649 | lymphocyte activation | 3.97E-06 | 2.79E-02 | 1.25E-04 | 1.18E-03 | 56 | 865 |
| GO: Biological Process | GO:1902679 | negative regulation of RNA biosynthetic process | 4.00E-06 | 2.81E-02 | 1.25E-04 | 1.18E-03 | 82 | 1442 |
| GO: Biological Process | GO:0042325 | regulation of phosphorylation | 4.06E-06 | 2.85E-02 | 1.26E-04 | 1.19E-03 | 87 | 1558 |
| GO: Biological Process | GO:0007417 | central nervous system development | 4.18E-06 | 2.94E-02 | 1.30E-04 | 1.22E-03 | 70 | 1172 |
| GO: Biological Process | GO:0010810 | regulation of cell-substrate adhesion | 4.45E-06 | 3.13E-02 | 1.37E-04 | 1.30E-03 | 24 | 246 |
| GO: Biological Process | GO:0034329 | cell junction assembly | 4.62E-06 | 3.25E-02 | 1.42E-04 | 1.34E-03 | 37 | 482 |
| GO: Biological Process | GO:0003300 | cardiac muscle hypertrophy | 4.79E-06 | 3.37E-02 | 1.47E-04 | 1.38E-03 | 16 | 123 |
| GO: Biological Process | GO:0008015 | blood circulation | 5.00E-06 | 3.51E-02 | 1.52E-04 | 1.44E-03 | 44 | 622 |
| GO: Biological Process | GO:0035909 | aorta morphogenesis | 5.23E-06 | 3.68E-02 | 1.57E-04 | 1.48E-03 | 9 | 39 |
| GO: Biological Process | GO:0001657 | ureteric bud development | 5.24E-06 | 3.69E-02 | 1.57E-04 | 1.48E-03 | 15 | 110 |
| GO: Biological Process | GO:0003190 | atrioventricular valve formation | 5.25E-06 | 3.69E-02 | 1.57E-04 | 1.48E-03 | 5 | 9 |
| GO: Biological Process | GO:1905247 | positive regulation of aspartic-type peptidase activity | 5.25E-06 | 3.69E-02 | 1.57E-04 | 1.48E-03 | 5 | 9 |
| GO: Biological Process | GO:0061005 | cell differentiation involved in kidney development | 5.27E-06 | 3.71E-02 | 1.57E-04 | 1.48E-03 | 11 | 60 |
| GO: Biological Process | GO:0030856 | regulation of epithelial cell differentiation | 5.51E-06 | 3.88E-02 | 1.64E-04 | 1.54E-03 | 20 | 184 |
| GO: Biological Process | GO:1903131 | mononuclear cell differentiation | 5.57E-06 | 3.92E-02 | 1.65E-04 | 1.55E-03 | 37 | 486 |
| GO: Biological Process | GO:0072164 | mesonephric tubule development | 5.88E-06 | 4.13E-02 | 1.72E-04 | 1.62E-03 | 15 | 111 |
| GO: Biological Process | GO:0072163 | mesonephric epithelium development | 5.88E-06 | 4.13E-02 | 1.72E-04 | 1.62E-03 | 15 | 111 |
| GO: Biological Process | GO:0030098 | lymphocyte differentiation | 5.97E-06 | 4.20E-02 | 1.74E-04 | 1.64E-03 | 34 | 430 |
| GO: Biological Process | GO:2000377 | regulation of reactive oxygen species metabolic process | 6.20E-06 | 4.36E-02 | 1.80E-04 | 1.70E-03 | 23 | 234 |
| GO: Biological Process | GO:0045444 | fat cell differentiation | 6.28E-06 | 4.42E-02 | 1.82E-04 | 1.72E-03 | 25 | 268 |
| GO: Biological Process | GO:0006935 | chemotaxis | 6.36E-06 | 4.47E-02 | 1.83E-04 | 1.73E-03 | 48 | 710 |
| GO: Biological Process | GO:0007517 | muscle organ development | 6.46E-06 | 4.54E-02 | 1.85E-04 | 1.75E-03 | 32 | 394 |
| GO: Biological Process | GO:0046903 | secretion | 6.71E-06 | 4.72E-02 | 1.92E-04 | 1.81E-03 | 91 | 1672 |
| GO: Biological Process | GO:0001934 | positive regulation of protein phosphorylation | 6.87E-06 | 4.83E-02 | 1.96E-04 | 1.85E-03 | 59 | 946 |
| GO: Cellular Component | GO:0031252 | cell leading edge | 2.10E-09 | 1.55E-06 | 1.55E-06 | 1.11E-05 | 44 | 469 |
| GO: Cellular Component | GO:0070161 | anchoring junction | 6.50E-09 | 4.78E-06 | 2.39E-06 | 1.72E-05 | 65 | 880 |
| GO: Cellular Component | GO:0015629 | actin cytoskeleton | 7.43E-08 | 5.47E-05 | 1.82E-05 | 1.31E-04 | 60 | 837 |
| GO: Cellular Component | GO:0005794 | Golgi apparatus | 1.14E-07 | 8.36E-05 | 2.09E-05 | 1.50E-04 | 100 | 1710 |
| GO: Cellular Component | GO:0030027 | lamellipodium | 4.19E-07 | 3.08E-04 | 6.17E-05 | 4.43E-04 | 24 | 214 |
| GO: Cellular Component | GO:0005884 | actin filament | 9.03E-07 | 6.65E-04 | 9.87E-05 | 7.09E-04 | 42 | 540 |
| GO: Cellular Component | GO:0045121 | membrane raft | 1.02E-06 | 7.47E-04 | 9.87E-05 | 7.09E-04 | 35 | 412 |
| GO: Cellular Component | GO:0098857 | membrane microdomain | 1.07E-06 | 7.90E-04 | 9.87E-05 | 7.09E-04 | 35 | 413 |
| GO: Cellular Component | GO:0005925 | focal adhesion | 1.96E-06 | 1.44E-03 | 1.60E-04 | 1.15E-03 | 35 | 424 |
| GO: Cellular Component | GO:0048471 | perinuclear region of cytoplasm | 2.54E-06 | 1.87E-03 | 1.87E-04 | 1.34E-03 | 56 | 846 |
| GO: Cellular Component | GO:0030055 | cell-substrate junction | 3.48E-06 | 2.56E-03 | 2.33E-04 | 1.67E-03 | 35 | 435 |
| GO: Cellular Component | GO:0030659 | cytoplasmic vesicle membrane | 1.00E-05 | 7.36E-03 | 5.80E-04 | 4.17E-03 | 53 | 822 |
| GO: Cellular Component | GO:0012506 | vesicle membrane | 1.03E-05 | 7.54E-03 | 5.80E-04 | 4.17E-03 | 54 | 844 |

FOXO1,SORBS2,PDLIM5,ATP2B4,GATA4,CCN4,NOTCH1,HEY2,TNFRSF1A,TNFRSF1B,AKAP6,ROCK2,IGFBP5,MIR21,ECE1,EDN1,LMNA
GREB1L,AHR,SNAI2,BCL2,PLXND1,NOTCH1,TGFBR2,HHEX,WNT4,NRARP,HOXA5,HOXA13,SLIT2,HOXB7,PITX2,COL4A1,WNT2B,CLIC4,ILK,CXCR4,KDR,FAT4,RDH10,EDN1
PRR5L,PICALM,FZD8,DLC1,MAPK12,PIBF1,SNX9,FNIP2,ADCY4,CIB1,FN1,RGCC,HIPK2,BIRC2,NLRP1,APP,MSN,DKK1,TRIB3,SNAI2,LIMCH1,ATP2B4,SAMD4A,TRADD,MYH9,RIPK1,DISC1,GATA3,GATA4,CCND1,BCL2,BCL6,STAT3,IQGAP1,CFLAR,MLKL,BMP6,NFE2L2,NFKBIA,RIPK3,GRN,CAV1,GSN,CCND3,TGFBR2,THBS1,[illegible]
RIN3,ABHD6,DLC1,JAG1,EVL,RGCC,VASH1,RHOB,LIMCH1,ATP2B4,PBLD,BCL2,STAT3,NFE2L2,NOTCH1,ABHD2,THBS1,NRG1,WNT4,SLIT2,COL3A1,CLIC4,IGFBP3,IGFBP5,MIR21,ILK,DACH1,GADD45A,PTPRM,ADAMTS9,DUSP3,PHLDB2
MEF2D,MRTFB,SOX6,FLNB,SORBS2,HDAC7,PDLIM5,DKK1,RIPK1,GATA4,BCL2,CFLAR,CCN4,NOTCH1,CAV1,ZFPM2,TGFBR2,HEY1,GPCPD1,HEY2,NRG1,HLX,HOXD9,PITX2,PLAGL1,AKAP6,NIBAN2,IGF2,VGLL4,ADAMTS9,KCNAB1,RB1,S1PR1,EDN1,EFNB2,LMNA,SMAD1
GREB1L,GATA3,BCL2,HEY1,WNT4,SLIT2,HOXB7,HES1,WNT2B,ZBTB16,ILK,FAT4,KIF26B,SMAD1,SMAD6,SMAD9
FOXO1,SORBS2,PDLIM5,ATP2B4,GATA4,CCN4,NOTCH1,HEY2,TNFRSF1A,TNFRSF1B,AKAP6,ROCK2,IGFBP5,MIR21,ECE1,EDN1,LMNA
ABCA1,ABL2,PICALM,ATP8A1,FES,AHR,SNX9,FNIP2,EVL,FN1,SGK1,RGCC,APP,CDC42EP3,MSN,SKAP1,SNAI2,LIMCH1,TPPP3,FSCN1,PALLD,DISC1,RAB11A,GATA2,GATA3,BCL2,PLXND1,BCL6,IQGAP1,CFLAR,NFATC2,NFE2L2,CLEC16A,KALRN,GRN,CAV1,GSN,PALM,SEPTIN7,CDH5,NRG1,RPS6KA5,WNT4,ZDHHC1,HOXA13,[illegible]
RPS6KA2,CD109,ETS1,DLC1,AHR,CIB1,P3H3,RGCC,VASH1,APP,SNAI2,CD274,RIPK1,LRRC32,CD164,GATA2,GATA3,PBLD,BCL2,BCL6,STAT3,SKAP2,NOTCH1,NPR1,TNS2,CAV1,TGFBR2,COL18A1,THBS1,IL1RL1,CDH5,CDKN2B,SLIT2,HES1,VEGFC,TP53I11,NIBAN2,IGFBP3,IGFBP5,IGFBP7,MIR21,VGLL4,RAPGEF2,ZBTB16,[illegible]
GREB1L,AHR,SNAI2,BCL2,PLXND1,NOTCH1,TGFBR2,HHEX,WNT4,NRARP,HOXA5,HOXA13,SLIT2,HOXB7,PITX2,COL4A1,WNT2B,CLIC4,ILK,CXCR4,KDR,FAT4,RTN4,RDH10,EDN1
TBC1D19,RIN3,PICALM,DLC1,ARHGAP31,VAV3,SNX9,RHOG,MYO9B,TBC1D2B,PLXND1,BCL6,IQGAP1,KALRN,ARHGEF12,TBC1D8,DGKI,PECAM1,RABGAP1,WNT4,SLIT2,RIN2,ARHGAP29,AGFG2,MAP4K4,SMAP2,RGS6,MIR21,RAPGEF2,ARHGAP18,DOCK4,TBC1D20,EVI5,JUN,IQSEC1,ARHGAP25,S1PR1,ARHGAP28,EPHA4
MEF2D,JAG1,AHR,BMP2K,HDAC7,SBNO2,DKK1,SNAI2,SUFU,CSGALNACT1,ZHX3,BCL2,CCN4,BMP6,NOTCH1,DNAJC13,RUNX2,HEY1,SGMS2,WNT4,VEGFC,COL5A2,WWTR1,IGF2,IGFBP3,IGFBP5,SRGN,MIR21,CCN2,ZBTB16,ILK,MTSS1,FAT4,S1PR1,LMNA,SMAD1,SMAD6
CD109,ZNF703,RGCC,VASH1,SNAI2,GATA2,GATA3,PBLD,CCND1,STAT3,STAT5A,CFLAR,BMP6,NOTCH1,GRN,CAV1,THBS1,HEY1,CDKN2B,NRARP,HOXA5,HES1,PLCG1,VEGFC,IGF2,MIR21,MTSS1,ITGB3,ITPR1,JUN,PTPRM,KDR,RB1,RTN4,LAMC1,EFNB2
PICALM,APP,TMED10,GRN,ROCK2,EPHA4
PICALM,ATXN1,SLC9B2,CIB1,FN1,SGK1,ZDHHC21,DISC1,RAB11A,BCL2,PLXND1,NOTCH1,KALRN,CTDSP1,GRN,HEY1,HEY2,TNFRSF1A,TNFRSF1B,TNR,CDH5,NRG1,PRTG,PDE3A,SLIT2,GORASP1,HES1,QKI,VEGFC,ROCK2,NIBAN2,BCAR1,ADGRA2,PRKCH,RAPGEF2,ILK,ITGB3,SIGLEC15,ZEB2,CXCR4,KDR,RTN4,ACTR2,EPHA4
GREB1L,JAG1,GATA3,BCL2,NOTCH1,HEY1,WNT4,HOXB7,HES1,WNT2B,WWTR1,ILK,MTSS1,FAT4,KIF26B
ETS2,WLS,MLLT3,HIPK2,DKK1,SUFU,GATA4,NOTCH1,SOX17,TGFBR2,HEY1,HEY2,HHEX,HOXA1,HOXA3,NRARP,HOXA4,HOXA5,HOXA6,HOXB7,HOXB8,HOXC4,HOXC6,HOXC8,HOXC9,HOXD3,HOXD8,HOXD9,PITX2,VAX2,HES1,WNT2B,PRKACB,RNF111,ZBTB16,ZEB2,EDN1,SMAD1,SMAD6
PRR5L,ABL2,CD109,MECOM,DLC1,ATXN1,MLLT3,PIBF1,FOXO1,ADCY4,CIB1,HIPK2,APP,ANKRD6,HDAC7,DKK1,TRIB3,SNAI2,SUFU,PRDM16,ATP2B4,TRADD,RIPK1,UNC5B,GATA2,GATA3,GATA4,PBLD,BCL2,BCL6,CFLAR,IER3,NFE2L2,NFKBIA,NOTCH1,SZT2,SOX17,TNS2,CAV1,RUNX2,TGFBR2,THBS1,HEY1,NANOS3,HEY2,[illegible]
CLIC5,JAG1,AHR,OPN5,MERTK,HIPK2,ITGA8,LRIG3,ATF6,PRDM16,ATP2B4,GATA2,GATA3,BCL2,STAT3,BMP6,NOTCH1,GRN,TGFBR2,HEY1,HEY2,HOXA1,HOXA13,PITX2,VAX2,HES1,COL4A1,COL5A2,WNT2B,CLIC4,CRYGS,IGFBP7,INHBA,INHBB,FJX1,ZEB2,TBC1D20,JAG2,JUN,PTPRM,KDR,FAT4,RDH10,ECE1,EDN1,[illegible]
PRR5L,CD109,DTX3L,FZD8,MGAT5,DLC1,PIBF1,SNX9,FNIP2,ADCY4,CIB1,FN1,RGCC,HIPK2,BIRC2,APP,DKK1,TRIB3,SNAI2,LIMCH1,ATP2B4,PHACTR1,RIPK1,GATA2,GATA3,GATA4,PBLD,CCND1,BCL2,BCL6,STAT3,IQGAP1,MLKL,BMP6,RIPK3,CTDSP1,CAV1,CCND3,PKIG,TGFBR2,THBS1,MAP4K5,RASSF1,DGKI,[illegible]
PRR5L,CD109,FZD8,PIBF1,SNX9,FNIP2,ADCY4,CIB1,FN1,RGCC,HIPK2,APP,DKK1,TRIB3,LIMCH1,ATP2B4,RIPK1,GATA4,PBLD,CCND1,BCL2,STAT3,IQGAP1,MLKL,BMP6,RIPK3,CTDSP1,CAV1,CCND3,PKIG,TGFBR2,THBS1,MAP4K5,DGKI,TNFRSF1A,CDH5,CDKN2B,NRG1,HHEX,RPS6KA5,PECAM1,SLIT2,PIK3CA,HES1,[illegible]
RIN3,ABHD6,DLC1,JAG1,EVL,RGCC,VASH1,RHOB,LIMCH1,ATP2B4,GATA3,PBLD,BCL2,STAT3,NFE2L2,NOTCH1,ABHD2,THBS1,NRG1,WNT4,SLIT2,COL3A1,CLIC4,IGFBP3,IGFBP5,MIR21,ILK,DACH1,GADD45A,PTPRM,ADAMTS9,DUSP3,PHLDB2
GATA4,NOTCH1,ZFPM2,TGFBR2,HEY1,HEY2,PITX2,SMAD6
ABCA1,CD109,WLS,ATP8A1,TEX264,SLC9B2,FES,MERTK,FN1,COPG1,RGCC,PIWIL4,APP,RAB14,CD93,RHOG,FRMD4A,COMMD9,GAD1,MYH9,RAB11A,GATA2,GATA3,CREG1,IQGAP1,BMP6,GLB1,NBEAL2,NFKB1,AP1M1,DNAJC1,TMED10,NOTCH1,DNAJC13,KALRN,GRN,COTL1,GSN,THBS1,FURIN,CD38,ENTPD1,DGKI,TNFRSF1A,[illegible]
GREB1L,JAG1,AHR,SUFU,GATA3,GATA4,PLXND1,NOTCH1,ZFPM2,TGFBR2,HEY1,HEY2,NRG1,PDE2A,HOXA13,SLIT2,PITX2,HES1,CXCR4,ADGRG6,SMAD6
MEF2D,ARID5B,SOX6,CSGALNACT1,BMP6,RUNX2,TGFBR2,THBS1,HOXA1,HOXA3,HOXA4,HOXA5,HOXA6,HOXB7,HOXB8,HOXC4,HOXC8,HOXC9,HOXD3,HOXD8,HOXD9,COL27A1,CCN2,RDH10,PLEKHA1
PICALM,EVL,FN1,APBB2,APP,RHOG,PALLD,UNC5B,DISC1,RAB11A,GATA3,BCL2,PLXND1,DST,MYPN,NOTCH1,KALRN,SZT2,TTL,SEPTIN7,TNR,NRG1,PRTG,RPS6KA5,MGLL,HOXA1,SLIT2,PIK3CA,VAX2,PLCG1,NIBAN2,FEZ2,ILK,ZEB2,GAB2,PTPRM,CXCR4,RTN4,ABLIM1,EFNB2,EPHA4
RIN3,PICALM,AHR,SCTR,VAV3,SNX9,EVL,FN1,CDC42EP3,MSN,MYH9,DISC1,RAB11A,COTL1,GSN,TTL,TNR,NRG1,PECAM1,TMOD2,MGLL,SLIT2,CLCN3,PIK3CA,GMFG,PRKCE,ARHGAP18,ILK,SPIRE1,CYRIA,FCHSD2,RTN4,ACTR2,EDN1,ARHGAP28
ETS2,MECOM,ATXN1,ARID5B,AHR,FNIP2,SOX6,FOXO1,ZNF703,HIPK2,APP,HDAC7,SBNO2,DKK1,TRIB3,SNAI2,ZFHX3,SUFU,PRDM16,ZHX3,GATA2,GATA3,CCND1,BCL6,CREG1,STAT3,BMP6,NFATC2,NFIC,NFKB1,NFKBIE,NOTCH1,TCF4,WWP1,CAV1,RUNX2,ZFPM2,CCND3,PKIG,HEY1,CD38,HEY2,NRG1,KLF12,HHEX,HIC1,PDE2A,RPS6
MSN,ZDHHC21,COL18A1,TNFRSF1A,CDH5,PDE2A,PECAM1,HOXA13,ROCK2,COL15A1,CLIC4,RAPGEF2,RTN4
ABCA1,FES,AHR,SNX9,FNIP2,EVL,SGK1,BIRC2,CDC42EP3,MSN,DKK1,SKAP1,TPPP3,FSCN1,PALLD,MYH9,CAMSAP2,CAV1,COTL1,GSN,CDH5,NRG1,PECAM1,TMOD2,ZDHHC1,SLIT2,HES1,GMFG,PRKCE,ARHGAP18,SPIRE1,CYRIA,FCHSD2,JAM3,RB1,ACTR2,ARHGAP28,SMAD6
ETS2,MECOM,ATXN1,ARID5B,AHR,FNIP2,SOX6,FOXO1,ZNF703,HIPK2,APP,HDAC7,SBNO2,DKK1,TRIB3,SNAI2,ZFHX3,SUFU,PRDM16,ZHX3,GATA2,GATA3,CCND1,BCL6,CREG1,STAT3,BMP6,NFATC2,NFIC,NFKB1,NFKBIE,NOTCH1,TCF4,WWP1,CAV1,RUNX2,ZFPM2,CCND3,PKIG,HEY1,CD38,HEY2,NRG1,KLF12,HHEX,HIC1,PDE2A,RPS6
PRR5L,CD109,FZD8,MGAT5,DLC1,ATXN1,AHR,VAV3,MERTK,PIBF1,SNX9,FNIP2,ADCY4,CIB1,FN1,RGCC,HIPK2,APP,DKK1,TRIB3,LIMCH1,ATP2B4,PHACTR1,RIPK1,GATA4,PBLD,CCND1,BCL2,STAT3,IQGAP1,MLKL,BMP6,IER3,RIPK3,CTDSP1,CAV1,CCND3,PKIG,TGFBR2,THBS1,MAP4K5,ENTPD1,DGKI,TNFRSF1A,CDH5,CDKN2B,NRG
PRR5L,CD109,FZD8,MGAT5,DLC1,ATXN1,AHR,VAV3,MERTK,PIBF1,SNX9,FNIP2,ADCY4,CIB1,FN1,RGCC,HIPK2,APP,DKK1,TRIB3,LIMCH1,ATP2B4,PHACTR1,RIPK1,GATA4,PBLD,CCND1,BCL2,STAT3,IQGAP1,MLKL,BMP6,IER3,RIPK3,CTDSP1,CAV1,CCND3,PKIG,TGFBR2,THBS1,MAP4K5,ENTPD1,DGKI,TNFRSF1A,CDH5,CDKN2B,NRG
ABL2,FZD8,AHR,VAV3,MERTK,PIBF1,ST3GAL1,HDAC7,MSN,CD274,PAG1,MYH9,LRRC32,GATA3,BCL2,BCL6,IL18R1,STAT3,STAT5A,NFATC2,SKAP2,RIPK3,WWP1,HS1BP3,CAV1,RUNX2,GSN,CCND3,TGFBR2,CD38,TNFRSF1B,HHEX,HLX,WNT4,LYST,TXK,NRARP,PIK3CA,HES1,KLF6,POU2F2,IGF2,MIR21,IL4R,IRF2BP2,ZBTB16,[illegible]
ETS2,MECOM,ATXN1,ARID5B,AHR,FNIP2,SOX6,FOXO1,ZNF703,HIPK2,APP,HDAC7,SBNO2,DKK1,TRIB3,SNAI2,ZFHX3,SUFU,PRDM16,ZHX3,GATA2,GATA3,CCND1,BCL6,CREG1,STAT3,BMP6,NFATC2,NFIC,NFKB1,NFKBIE,NOTCH1,TCF4,WWP1,CAV1,RUNX2,ZFPM2,CCND3,PKIG,HEY1,CD38,HEY2,NRG1,KLF12,HHEX,HIC1,PDE2,[illegible]
PRR5L,CD109,FZD8,ATXN1,VAV3,MERTK,PIBF1,SNX9,FNIP2,ADCY4,CIB1,FN1,RGCC,HIPK2,APP,DKK1,TRIB3,LIMCH1,ATP2B4,PHACTR1,RIPK1,GATA4,PBLD,CCND1,BCL2,STAT3,IQGAP1,MLKL,BMP6,IER3,RIPK3,CTDSP1,CAV1,CCND3,PKIG,TGFBR2,THBS1,MAP4K5,DGKI,TNFRSF1A,CDH5,CDKN2B,NRG1,HHEX,PDE2A,RPS6KA5,[illegible]
ABL2,ETS1,MECOM,WLS,MRTFB,DLC1,ATXN1,SCTR,SOX6,ITGA8,APP,DKK1,ZFHX3,SUFU,ATP2B4,PHACTR1,DISC1,GATA2,BCL2,STAT3,GNG12,NHLH2,TACC1,DCLK2,NOTCH1,NPAS2,NRGN,KALRN,SZT2,WWP1,GRN,GSN,TGFBR2,TNFRSF1B,TNR,NRG1,HHEX,WNT4,HOXA1,SLIT2,HOXB8,TPP1,PITX2,VAX2,HES1,QKI,COL3A1,COL4A1,[illegible]
DLC1,JAG1,CIB1,FN1,SKAP1,LIMCH1,DISC1,BCL2,BCL6,IQGAP1,NOTCH1,THBS1,WNT4,RIN2,MAP4K4,VEGFC,ROCK2,BCAR1,PRKCE,ILK,ITGB3,KDR,DUSP3,PHLDB2
DLC1,ZNF703,FN1,APP,HDAC7,PDLIM5,DKK1,SNAI2,LIMCH1,FSCN1,BCL2,PLXND1,LRRC4,PCDH17,IQGAP1,DST,CAV1,THBS1,CDH5,NRG1,CDHR3,PECAM1,WNT4,MAP4K4,ROCK2,PRKCH,RAPGEF2,ILK,ITGAV,ITGB3,KDR,JAM3,DUSP3,LAMB3,LAMC1,EFNB2,PHLDB2
FOXO1,SORBS2,PDLIM5,ATP2B4,GATA4,CCN4,NOTCH1,HEY2,TNFRSF1A,TNFRSF1B,AKAP6,ROCK2,MIR21,ECE1,EDN1,LMNA
ACTA2,AHR,SGK1,APP,SMTN,ATP2B4,SCN3B,ZDHHC21,GATA4,STIM1,BMP6,IER3,TBXAS1,NPR1,CAV1,GSN,TBC1D8,CD34,CD38,HEY2,CDH5,PDE2A,PDE3A,PDE4B,SLIT2,PIK3CA,VEGFC,AKAP6,ROCK2,CACNA2D4,PRCP,MIR21,CCN2,DOCK4,ITPR1,PTPRM,CXCR4,KCNE3,ECE1,EDN1,MYOF,ENPEP,EPAS1,SMAD6
JAG1,NOTCH1,HEY1,HEY2,HES1,COL3A1,ADAMTS9,KAT6A,EFNB2
GREB1L,GATA3,BCL2,HEY1,WNT4,SLIT2,HOXB7,HES1,WNT2B,ILK,FAT4,KIF26B,SMAD1,SMAD6,SMAD9
GATA4,NOTCH1,ZFPM2,HEY1,HEY2
PICALM,APP,GRN,ROCK2,EPHA4
ACTA2,JAG1,GATA3,IQGAP1,NOTCH1,CD34,WNT4,HES1,WWTR1,MTSS1,FAT4
CD109,JAG1,ZDHHC21,GATA3,CCND1,STAT5A,BMP6,NOTCH1,CAV1,HEY1,HEY2,TNFRSF1A,CDH5,HES1,ROCK2,WWTR1,MIR21,PRKCH,ZEB2,FAT4
ABL2,FZD8,FES,AHR,MERTK,ST3GAL1,HDAC7,MYH9,GATA3,BCL2,BCL6,IL18R1,STAT3,STAT5A,NFATC2,RIPK3,WWP1,HS1BP3,RUNX2,TGFBR2,HHEX,HLX,WNT4,TXK,NRARP,KLF6,POU2F2,MIR21,IL4R,IRF2BP2,ZBTB16,INHBA,JAG2,JUN,PPP2R3C,LEPR,LYL1
GREB1L,GATA3,BCL2,HEY1,WNT4,SLIT2,HOXB7,HES1,WNT2B,ILK,FAT4,KIF26B,SMAD1,SMAD6,SMAD9
GREB1L,GATA3,BCL2,HEY1,WNT4,SLIT2,HOXB7,HES1,WNT2B,ILK,FAT4,KIF26B,SMAD1,SMAD6,SMAD9
ABL2,FZD8,AHR,MERTK,ST3GAL1,HDAC7,GATA3,BCL2,BCL6,IL18R1,STAT3,STAT5A,NFATC2,RIPK3,WWP1,HS1BP3,RUNX2,TGFBR2,HHEX,HLX,WNT4,TXK,NRARP,KLF6,POU2F2,MIR21,IL4R,IRF2BP2,ZBTB16,INHBA,JAG2,PPP2R3C,LEPR,LYL1
FOXO1,BIRC2,APP,ATP2B4,RIPK1,BCL2,STAT3,CFLAR,IER3,NFE2L2,RIPK3,CAV1,TGFBR2,THBS1,CD34,TLR5,STK17A,CLCN3,ROCK2,PRCP,MIR21,GADD45A,EDN1
MECOM,JAG1,ARID5B,FOXO1,AAMDC,TRIB3,SNAI2,PRDM16,CCDC71L,GATA2,GATA3,CCND1,CCN4,ZFPM2,HES1,WWTR1,ZNF516,MIR21,ZBTB16,INHBB,ZADH2,LAMA4,LAMB3,SDF4,SMAD6
RIN3,MTUS1,FES,VAV3,EVL,FN1,APBB2,APP,RHOG,SNAI2,PALLD,MYO9B,UNC5B,GATA3,PLXND1,MYPN,NOTCH1,KALRN,THBS1,TNR,NRG1,PRTG,PDE4B,RPS6KA5,LYST,HOXA1,SLIT2,PIK3CA,PLCG1,VEGFC,NIBAN2,BCAR1,ADGRA2,FEZ2,DOCK4,ITGAV,ITGB3,GAB2,PTPRM,CXCR4,KDR,JAM3,DUSP3,S1PR1,EDN1,ABLIM1,EFNB2,EPH
MEF2D,MRTFB,MAPK12,ARID5B,SOX6,FLNB,HDAC7,SMTN,ZFHX3,CD164,GATA4,BCL2,CFLAR,NOTCH1,CAV1,ZFPM2,HEY1,GPCPD1,HEY2,NRG1,HLX,HOXD9,PITX2,PLAGL1,COL3A1,NIBAN2,IGF2,MTSS1,KCNAB1,RB1,S1PR1,LMNA
ABCA1,CD109,WLS,ATP8A1,TEX264,SLC9B2,FES,MERTK,FN1,COPG1,SGK1,RGCC,PIWIL4,APP,RAB14,CD93,RHOG,FRMD4A,COMMD9,GAD1,MYH9,RAB11A,GATA2,GATA3,CCND1,CREG1,STAT5A,IQGAP1,BMP6,GLB1,NBEAL2,NFKB1,AP1M1,DNAJC1,TMED10,NOTCH1,NPR1,DNAJC13,KALRN,GRN,CAV1,COTL1,GSN,THBS1,FURIN,CD38,EN
PRR5L,FZD8,PIBF1,SNX9,FNIP2,ADCY4,CIB1,RGCC,HIPK2,APP,DKK1,LIMCH1,ATP2B4,RIPK1,GATA4,CCND1,BCL2,STAT3,IQGAP1,MLKL,BMP6,RIPK3,CAV1,CCND3,TGFBR2,THBS1,MAP4K5,DGKI,TNFRSF1A,NRG1,RPS6KA5,PECAM1,PIK3CA,HES1,MAP4K4,VEGFC,AKAP6,ROCK2,IGF2,PRKACB,PRKAR2B,MIR21,CCN2,RAPGEF2,ILK,
ABL2,ACTA2,WLS,DLC1,ARHGAP31,SNX9,EVL,SORBS2,CIB1,CDC42BPA,VEZT,AAK1,APBB2,ITGA8,APP,FSCN1,PALLD,MYH9,MYO9B,PLXND1,FRMD4B,IQGAP1,DST,GSN,PALM,NRG1,PECAM1,PARVB,CLCN3,PIK3CA,PLCG1,BCAR1,ARHGAP18,ILK,MTSS1,ITGAV,ITGB3,PTPRM,CXCR4,ACTR2,ABLIM1,PLEKHA1,PLEKHO1,PHLDB2
DLC1,ARHGAP31,JAG1,FES,AMOTL2,EVL,FLNB,SORBS2,CDC42BPA,VEZT,ITGA8,APP,PDLIM5,MSN,RHOB,RHOG,FRMD4A,SKAP1,FSCN1,PALLD,SNTB2,MYH9,SDCCAG8,CCND1,FRMD4B,IQGAP1,DST,NOTCH1,TNS2,CAV1,GSN,AFAP1,IL1RL1,CDH5,CDHR3,PECAM1,PARVB,JAM2,PIK3CA,AHNAK,PLCG1,MAP4K4,AKAP6,USP53,CLIC4,NIBA
ABL2,ACTA2,RCSD1,DLC1,CLIC5,SNX9,FLNB,SORBS2,CIB1,CDC42BPA,VEZT,MYO1B,APP,CDC42EP3,PDLIM5,SMTN,MSN,CD274,LIMCH1,FSCN1,PALLD,FILIP1,MYH9,MYO9B,IQGAP1,DST,GNG12,KALRN,COTL1,GSN,AFAP1,PALM,MICALCL,SEPTIN7,TMOD2,PARVB,VPS16,AHNAK,ABCG2,CLIC4,SLC34A3,BCAR1,ADGRA2,SH3PXD2A,DC
ABCA1,SLC35E1,FAM91A1,WLS,PICALM,FZD8,MGAT5,ATP8A1,CLIC5,ST3GAL6,MTUS1,COG6,FES,COG5,SNX9,GOLGA7B,CIB1,COPG1,MYO1B,ST3GAL1,ST3GAL2,CHST1,ST3GAL3,APP,RAB14,PCGF5,GPR107,ATF6,COMMD9,VAMP4,CSGALNACT1,SNTB2,ZDHHC21,SLC16A13,RAB11A,BCL6,GLB1,ST3GAL5,NFE2L2,NFKBIE,GOLIM4,AP1M
ABL2,ACTA2,ARHGAP31,EVL,SORBS2,CIB1,CDC42BPA,APBB2,APP,FSCN1,PALLD,MYO9B,PLXND1,GSN,PARVB,PIK3CA,PLCG1,BCAR1,ILK,ITGAV,ITGB3,PTPRM,ACTR2,ABLIM1
ACTA2,RCSD1,DLC1,CLIC5,FLNB,CIB1,VEZT,MYO1B,APP,PDLIM5,SMTN,MSN,LIMCH1,FSCN1,PALLD,MYH9,MYO9B,IQGAP1,DST,GNG12,COTL1,AFAP1,PALM,MICALCL,SEPTIN7,VPS16,ABCG2,CLIC4,SLC34A3,ADGRA2,DOCK4,ILK,ITGAV,ITGB3,FCHSD2,JAM3,RALGDS,CUBN,ABLIM1,ENPEP,MARCKS,EPHA4
ABCA1,DLC1,BIRC2,APP,SKAP1,ATP2B4,TRADD,PAG1,RIPK1,UNC5B,IQGAP1,STIM1,CFLAR,CAV1,TFPI,TGFBR2,FURIN,TNFRSF1A,TNFRSF1B,TNR,PDE2A,PECAM1,TPP1,AHNAK,ABCG2,AKAP6,PRKAR2B,CAVIN1,CTSB,ITPR1,SLC27A1,KDR,KCNE3,S1PR1,MYOF
ABCA1,DLC1,BIRC2,APP,SKAP1,ATP2B4,TRADD,PAG1,RIPK1,UNC5B,IQGAP1,STIM1,CFLAR,CAV1,TFPI,TGFBR2,FURIN,TNFRSF1A,TNFRSF1B,TNR,PDE2A,PECAM1,TPP1,AHNAK,ABCG2,AKAP6,PRKAR2B,CAVIN1,CTSB,ITPR1,SLC27A1,KDR,KCNE3,S1PR1,MYOF
DLC1,ARHGAP31,FES,EVL,FLNB,SORBS2,ITGA8,MSN,RHOB,RHOG,PALLD,SNTB2,MYH9,IQGAP1,DST,TNS2,CAV1,GSN,AFAP1,IL1RL1,PARVB,AHNAK,MAP4K4,BCAR1,TES,ILK,ITGAV,ITGB3,JAK1,ACTR2,EFNB2,SORBS3,MARCKS,EPB41L2,PHLDB2
ABCA1,PICALM,DOCK6,SORBS2,CIB1,SGK1,ANXA4,MYO1B,APP,RAB14,MSN,TPPP3,MYO9B,DISC1,RAB11A,BCL2,DST,GLB1,NFKBIE,SERINC3,KALRN,CAV1,GSN,TG,CD34,NANOS3,DGKI,TNFRSF1B,PDE2A,PDE4B,PIK3CA,TMEM192,CREB3L2,AKAP6,CLIC4,SLC30A7,PRKACB,PRKAR2B,PRKCE,CCN2,RAPGEF2,CTSB,INHBA,INHBB,SPIR
DLC1,ARHGAP31,FES,EVL,FLNB,SORBS2,ITGA8,MSN,RHOB,RHOG,PALLD,SNTB2,MYH9,IQGAP1,DST,TNS2,CAV1,GSN,AFAP1,IL1RL1,PARVB,AHNAK,MAP4K4,BCAR1,TES,ILK,ITGAV,ITGB3,JAK1,ACTR2,EFNB2,SORBS3,MARCKS,EPB41L2,PHLDB2
CD109,WLS,PICALM,ATP8A1,SLC9B2,ATP6V0D2,SNX9,COPG1,RAB14,MSN,CD93,RHOG,LRIG3,VAMP4,SNTB2,AHNAK2,GAD1,RAB11A,IQGAP1,NBEAL2,AP1M1,TMED10,DNAJC13,NRGN,CAV1,FURIN,CD38,ZNRF1,DGKI,TNFRSF1B,PECAM1,WNT4,AP2S1,CLCN3,SH3BP5,CLIC4,SEC31B,PRCP,RAB11FIP5,PHACTR2,SLC6A17,RNF144A,SPI
CD109,WLS,PICALM,ATP8A1,SLC9B2,ATP6V0D2,SNX9,COPG1,ANXA4,RAB14,MSN,CD93,RHOG,LRIG3,VAMP4,SNTB2,AHNAK2,GAD1,RAB11A,IQGAP1,NBEAL2,AP1M1,TMED10,DNAJC13,NRGN,CAV1,FURIN,CD38,ZNRF1,DGKI,TNFRSF1B,PECAM1,WNT4,AP2S1,CLCN3,SH3BP5,CLIC4,SEC31B,PRCP,RAB11FIP5,PHACTR2,SLC6A17,RNF14

| | | | | | | | | |
|---|---|---|---|---|---|---|---|---|
| GO: Cellular Component | GO:0099503 | secretory vesicle | 1.28E-05 | 9.40E-03 | 6.72E-04 | 4.82E-03 | 66 | 1113 |
| GO: Cellular Component | GO:0031091 | platelet alpha granule | 1.61E-05 | 1.19E-02 | 7.91E-04 | 5.68E-03 | 13 | 92 |
| GO: Cellular Component | GO:0005938 | cell cortex | 1.78E-05 | 1.31E-02 | 8.21E-04 | 5.89E-03 | 29 | 355 |
| GO: Cellular Component | GO:0001726 | ruffle | 1.98E-05 | 1.45E-02 | 8.55E-04 | 6.14E-03 | 20 | 199 |
| GO: Cellular Component | GO:0010008 | endosome membrane | 2.27E-05 | 1.67E-02 | 8.91E-04 | 6.40E-03 | 37 | 514 |
| GO: Cellular Component | GO:0005788 | endoplasmic reticulum lumen | 2.30E-05 | 1.69E-02 | 8.91E-04 | 6.40E-03 | 27 | 323 |
| GO: Cellular Component | GO:0042641 | actomyosin | 3.58E-05 | 2.64E-02 | 1.32E-03 | 9.46E-03 | 13 | 99 |
| GO: Cellular Component | GO:0098685 | Schaffer collateral – CA1 synapse | 3.88E-05 | 2.86E-02 | 1.36E-03 | 9.77E-03 | 14 | 114 |
| GO: Cellular Component | GO:0097517 | contractile actin filament bundle | 4.87E-05 | 3.58E-02 | 1.56E-03 | 1.12E-02 | 12 | 88 |
| GO: Cellular Component | GO:0001725 | stress fiber | 4.87E-05 | 3.58E-02 | 1.56E-03 | 1.12E-02 | 12 | 88 |
| GO: Cellular Component | GO:0005764 | lysosome | 5.90E-05 | 4.34E-02 | 1.74E-03 | 1.25E-02 | 46 | 726 |
| GO: Cellular Component | GO:0000323 | lytic vacuole | 5.90E-05 | 4.34E-02 | 1.74E-03 | 1.25E-02 | 46 | 726 |

CD109,PICALM,ATP8A1,TEX264,SLC9B2,FN1,VEZT,APP,RAB14,MSN,CD93,RHOG,COMMD9,VAMP4,DISC1,CREG1,IQGAP1,GLB1,NBEAL2,NFKB1,AP1M1,TMED10,NOTCH1,DNAJC13,ABHD2,GRN,CAV1,COTL1,GSN,THBS1,CD38,ENTPD1,ZNRF1,DGKI,TNFRSF1B,PDE4B,PECAM1,CLCN3,EXOC3L1,VEGFC,DPP7,VWF,CLIP2,GMFG,IGF2,IGFB
CD109,TEX264,FN1,APP,THBS1,PECAM1,VEGFC,VWF,IGF2,IGFBP3,SRGN,PHACTR2,ITGB3
DLC1,CLIC5,SNX9,FLNB,RHOB,RHOG,FSCN1,GAD1,ZHX3,MYH9,MYO9B,DENND2B,IQGAP1,STIM1,DST,FRYL,CAV1,COTL1,GSN,SEPTIN7,EXOC3L1,CCN2,SPIRE1,RHOBTB1,ACTR2,ACTR1A,MARCKS,EPB41L2,PHLDB2
DLC1,SNX9,CIB1,FSCN1,PALLD,MYH9,MYO9B,FRMD4B,IQGAP1,GSN,PECAM1,CLCN3,PLCG1,BCAR1,ARHGAP18,MTSS1,ITGAV,ITGB3,PLEKHA1,PLEKHO1
SPNS2,ABHD6,DTX3L,WLS,SLC9B2,ATP6V0D2,MYO1B,RAB14,SLC48A1,RHOB,DKK1,CD274,VAMP4,SLC9A9,RIPK1,CD164,RAB11A,GOLIM4,CLEC16A,DNAJC13,CAV1,FURIN,TSPAN15,VPS16,ZDHHC1,AP2S1,CLCN3,RAB11FIP5,LEPROT,RNF144A,TMEM163,MVB12B,WIPI1,CUBN,ATP9A,ECE1,EPHA4
EVA1A,FN1,APP,ARSG,UGGT2,DNAJC1,PRSS23,COL18A1,THBS1,FURIN,TSPAN15,MGAT4A,WNT4,COL3A1,COL4A1,COL4A2,COL5A2,SUMF2,COL15A1,COL27A1,IGFBP3,IGFBP5,IGFBP7,TXNDC5,TSPAN14,LAMC1,EDN1
ACTA2,DLC1,FLNB,CDC42BPA,PDLIM5,LIMCH1,FSCN1,PALLD,MYH9,DST,SEPTIN7,ILK,ABLIM1
PICALM,RAB11A,LRRC4,STAT3,DGKI,TNR,PLCG1,ROCK2,SRGN,SLC30A1,ITPR1,IQSEC1,EFNB2,EPHA4
ACTA2,DLC1,FLNB,PDLIM5,LIMCH1,FSCN1,PALLD,MYH9,DST,SEPTIN7,ILK,ABLIM1
ACTA2,DLC1,FLNB,PDLIM5,LIMCH1,FSCN1,PALLD,MYH9,DST,SEPTIN7,ILK,ABLIM1
ABHD6,DTX3L,ATP8A1,EVA1A,ATP6V0D2,FNIP2,OSTM1,APP,RAB14,SLC48A1,ARSG,VAMP4,CD164,CREG1,SHKBP1,GLB1,AP1M1,CLEC16A,DNAJC13,SZT2,GRN,CD34,ZNRF1,VPS16,PLA2G15,SIAE,AP2S1,CLCN3,TPP1,AHNAK,PIP4K2A,TMEM192,DPP7,PRCP,SRGN,CTSB,FYCO1,TXNDC5,CXCR4,RAB2A,CUBN,ECE1,ACTR2,SLC1[illegible],CA
ABHD6,DTX3L,ATP8A1,EVA1A,ATP6V0D2,FNIP2,OSTM1,APP,RAB14,SLC48A1,ARSG,VAMP4,CD164,CREG1,SHKBP1,GLB1,AP1M1,CLEC16A,DNAJC13,SZT2,GRN,CD34,ZNRF1,VPS16,PLA2G15,SIAE,AP2S1,CLCN3,TPP1,AHNAK,PIP4K2A,TMEM192,DPP7,PRCP,SRGN,CTSB,FYCO1,TXNDC5,CXCR4,RAB2A,CUBN,ECE1,ACTR2,SLC17[illegible],CA

| Category | ID | Name | p-value | q-value Bonferroni | q-value FDR B&H | q-value FDR B&Y | Hit Count in Query List | Hit Count in Genome |
|---|---|---|---|---|---|---|---|---|
| GO: Molecular Function | GO:0016880 | acid-ammonia (or amide) ligase activity | 2.39E-05 | 2.09E-02 | 1.63E-02 | 1.20E-01 | 3 | 4 |
| GO: Molecular Function | GO:0016917 | GABA receptor activity | 4.05E-05 | 3.54E-02 | 1.63E-02 | 1.20E-01 | 5 | 22 |
| GO: Biological Process | GO:0030182 | neuron differentiation | 3.39E-10 | 1.54E-06 | 1.09E-06 | 9.79E-06 | 65 | 1639 |
| GO: Biological Process | GO:0007267 | cell-cell signaling | 4.78E-10 | 2.18E-06 | 1.09E-06 | 9.79E-06 | 73 | 1971 |
| GO: Biological Process | GO:0048699 | generation of neurons | 1.00E-09 | 4.56E-06 | 1.52E-06 | 1.37E-05 | 68 | 1802 |
| GO: Biological Process | GO:0022008 | neurogenesis | 1.47E-09 | 6.68E-06 | 1.67E-06 | 1.50E-05 | 71 | 1940 |
| GO: Biological Process | GO:0098916 | anterograde trans-synaptic signaling | 2.38E-09 | 1.08E-05 | 1.81E-06 | 1.63E-05 | 43 | 899 |
| GO: Biological Process | GO:0007268 | chemical synaptic transmission | 2.38E-09 | 1.08E-05 | 1.81E-06 | 1.63E-05 | 43 | 899 |
| GO: Biological Process | GO:0099537 | trans-synaptic signaling | 3.21E-09 | 1.46E-05 | 2.09E-06 | 1.88E-05 | 43 | 908 |
| GO: Biological Process | GO:0048666 | neuron development | 4.73E-09 | 2.16E-05 | 2.69E-06 | 2.42E-05 | 55 | 1358 |
| GO: Biological Process | GO:0099536 | synaptic signaling | 6.11E-09 | 2.78E-05 | 3.09E-06 | 2.78E-05 | 43 | 928 |
| GO: Biological Process | GO:1903530 | regulation of secretion by cell | 1.67E-08 | 7.58E-05 | 7.58E-06 | 6.82E-05 | 36 | 719 |
| GO: Biological Process | GO:0023061 | signal release | 2.26E-08 | 1.03E-04 | 9.36E-06 | 8.42E-05 | 33 | 629 |
| GO: Biological Process | GO:0006836 | neurotransmitter transport | 9.28E-08 | 4.22E-04 | 3.52E-05 | 3.17E-04 | 20 | 275 |
| GO: Biological Process | GO:0051046 | regulation of secretion | 1.45E-07 | 6.59E-04 | 5.07E-05 | 4.56E-04 | 36 | 784 |
| GO: Biological Process | GO:0001505 | regulation of neurotransmitter levels | 1.66E-07 | 7.55E-04 | 5.39E-05 | 4.85E-04 | 20 | 285 |
| GO: Biological Process | GO:0007417 | central nervous system development | 2.50E-07 | 1.14E-03 | 7.59E-05 | 6.83E-04 | 46 | 1172 |
| GO: Biological Process | GO:0031344 | regulation of cell projection organization | 1.88E-06 | 8.55E-03 | 5.23E-04 | 4.70E-03 | 34 | 800 |
| GO: Biological Process | GO:0050808 | synapse organization | 1.95E-06 | 8.88E-03 | 5.23E-04 | 4.70E-03 | 26 | 522 |
| GO: Biological Process | GO:0010975 | regulation of neuron projection development | 2.13E-06 | 9.71E-03 | 5.39E-04 | 4.85E-03 | 27 | 558 |
| GO: Biological Process | GO:0042391 | regulation of membrane potential | 2.32E-06 | 1.06E-02 | 5.57E-04 | 5.01E-03 | 26 | 527 |
| GO: Biological Process | GO:0031175 | neuron projection development | 2.68E-06 | 1.22E-02 | 5.83E-04 | 5.25E-03 | 44 | 1197 |
| GO: Biological Process | GO:0120036 | plasma membrane bounded cell projection organization | 2.70E-06 | 1.23E-02 | 5.83E-04 | 5.25E-03 | 60 | 1863 |
| GO: Biological Process | GO:0051050 | positive regulation of transport | 2.82E-06 | 1.28E-02 | 5.83E-04 | 5.25E-03 | 41 | 1081 |
| GO: Biological Process | GO:0120035 | regulation of plasma membrane bounded cell projection organization | 3.12E-06 | 1.42E-02 | 6.18E-04 | 5.56E-03 | 33 | 782 |
| GO: Biological Process | GO:0007610 | behavior | 4.90E-06 | 2.23E-02 | 9.29E-04 | 8.36E-03 | 32 | 762 |
| GO: Biological Process | GO:0030030 | cell projection organization | 5.35E-06 | 2.44E-02 | 9.75E-04 | 8.77E-03 | 60 | 1904 |
| GO: Biological Process | GO:0099003 | vesicle-mediated transport in synapse | 6.42E-06 | 2.92E-02 | 1.13E-03 | 1.01E-02 | 17 | 271 |
| GO: Biological Process | GO:1903532 | positive regulation of secretion by cell | 7.05E-06 | 3.21E-02 | 1.18E-03 | 1.07E-02 | 20 | 363 |
| GO: Biological Process | GO:0046883 | regulation of hormone secretion | 7.28E-06 | 3.32E-02 | 1.18E-03 | 1.07E-02 | 19 | 333 |
| GO: Biological Process | GO:0099643 | signal release from synapse | 7.91E-06 | 3.60E-02 | 1.20E-03 | 1.08E-02 | 15 | 219 |
| GO: Biological Process | GO:0007269 | neurotransmitter secretion | 7.91E-06 | 3.60E-02 | 1.20E-03 | 1.08E-02 | 15 | 219 |
| GO: Biological Process | GO:0021889 | olfactory bulb interneuron differentiation | 8.34E-06 | 3.79E-02 | 1.22E-03 | 1.10E-02 | 5 | 17 |
| GO: Cellular Component | GO:0045202 | synapse | 1.20E-15 | 7.89E-13 | 7.89E-13 | 5.57E-12 | 75 | 1582 |
| GO: Cellular Component | GO:0097060 | synaptic membrane | 2.20E-13 | 1.45E-10 | 7.23E-11 | 5.11E-10 | 36 | 480 |
| GO: Cellular Component | GO:0043005 | neuron projection | 2.20E-12 | 1.44E-09 | 4.81E-10 | 3.40E-09 | 71 | 1679 |
| GO: Cellular Component | GO:0098793 | presynapse | 3.94E-12 | 2.58E-09 | 6.46E-10 | 4.56E-09 | 41 | 673 |
| GO: Cellular Component | GO:0045211 | postsynaptic membrane | 9.05E-12 | 5.94E-09 | 1.19E-09 | 8.39E-09 | 28 | 335 |
| GO: Cellular Component | GO:0098794 | postsynapse | 1.49E-11 | 9.74E-09 | 1.62E-09 | 1.15E-08 | 44 | 795 |
| GO: Cellular Component | GO:0098978 | glutamatergic synapse | 3.24E-11 | 2.12E-08 | 3.03E-09 | 2.14E-08 | 34 | 513 |
| GO: Cellular Component | GO:0099572 | postsynaptic specialization | 7.50E-11 | 4.92E-08 | 6.15E-09 | 4.35E-08 | 32 | 473 |
| GO: Cellular Component | GO:0014069 | postsynaptic density | 2.37E-10 | 1.56E-07 | 1.44E-08 | 1.02E-07 | 30 | 439 |
| GO: Cellular Component | GO:0032279 | asymmetric synapse | 2.37E-10 | 1.56E-07 | 1.44E-08 | 1.02E-07 | 30 | 439 |
| GO: Cellular Component | GO:0098984 | neuron to neuron synapse | 2.41E-10 | 1.58E-07 | 1.44E-08 | 1.02E-07 | 31 | 467 |
| GO: Cellular Component | GO:0099240 | intrinsic component of synaptic membrane | 3.87E-10 | 2.54E-07 | 2.12E-08 | 1.50E-07 | 22 | 244 |
| GO: Cellular Component | GO:0036477 | somatodendritic compartment | 7.85E-10 | 5.15E-07 | 3.96E-08 | 2.80E-07 | 51 | 1145 |
| GO: Cellular Component | GO:0098936 | intrinsic component of postsynaptic membrane | 2.54E-09 | 1.67E-06 | 1.19E-07 | 8.41E-07 | 18 | 178 |
| GO: Cellular Component | GO:0098982 | GABA-ergic synapse | 7.07E-09 | 4.64E-06 | 3.09E-07 | 2.18E-06 | 14 | 109 |
| GO: Cellular Component | GO:0098948 | intrinsic component of postsynaptic specialization membrane | 1.01E-08 | 6.63E-06 | 4.14E-07 | 2.93E-06 | 14 | 112 |
| GO: Cellular Component | GO:0030424 | axon | 1.34E-08 | 8.77E-06 | 5.16E-07 | 3.65E-06 | 40 | 848 |
| GO: Cellular Component | GO:0099699 | integral component of synaptic membrane | 1.82E-08 | 1.20E-05 | 6.64E-07 | 4.69E-06 | 19 | 225 |
| GO: Cellular Component | GO:0042734 | presynaptic membrane | 3.45E-08 | 2.26E-05 | 1.19E-06 | 8.41E-06 | 18 | 210 |
| GO: Cellular Component | GO:0099060 | integral component of postsynaptic specialization membrane | 4.31E-08 | 2.82E-05 | 1.41E-06 | 9.98E-06 | 13 | 106 |
| GO: Cellular Component | GO:0099055 | integral component of postsynaptic membrane | 5.03E-08 | 3.30E-05 | 1.57E-06 | 1.11E-05 | 16 | 169 |
| GO: Cellular Component | GO:0099634 | postsynaptic specialization membrane | 7.62E-08 | 5.00E-05 | 2.27E-06 | 1.60E-05 | 15 | 152 |
| GO: Cellular Component | GO:0099146 | intrinsic component of postsynaptic density membrane | 1.27E-07 | 8.34E-05 | 3.63E-06 | 2.56E-05 | 11 | 79 |
| GO: Cellular Component | GO:0030425 | dendrite | 1.90E-07 | 1.25E-04 | 4.98E-06 | 3.52E-05 | 37 | 828 |
| GO: Cellular Component | GO:0097447 | dendritic tree | 1.90E-07 | 1.25E-04 | 4.98E-06 | 3.52E-05 | 37 | 828 |
| GO: Cellular Component | GO:0099061 | integral component of postsynaptic density membrane | 5.59E-07 | 3.66E-04 | 1.41E-05 | 9.96E-05 | 10 | 73 |
| GO: Cellular Component | GO:0098839 | postsynaptic density membrane | 7.71E-07 | 5.06E-04 | 1.87E-05 | 1.32E-04 | 12 | 114 |
| GO: Cellular Component | GO:0098889 | intrinsic component of presynaptic membrane | 2.66E-06 | 1.74E-03 | 6.23E-05 | 4.40E-04 | 12 | 128 |
| GO: Cellular Component | GO:0044297 | cell body | 6.16E-06 | 4.04E-03 | 1.39E-04 | 9.84E-04 | 34 | 845 |
| GO: Cellular Component | GO:0043025 | neuronal cell body | 1.03E-05 | 6.75E-03 | 2.25E-04 | 1.59E-03 | 31 | 753 |
| GO: Cellular Component | GO:0031226 | intrinsic component of plasma membrane | 1.12E-05 | 7.34E-03 | 2.37E-04 | 1.67E-03 | 57 | 1818 |
| GO: Cellular Component | GO:0005887 | integral component of plasma membrane | 1.20E-05 | 7.84E-03 | 2.45E-04 | 1.73E-03 | 55 | 1735 |
| GO: Cellular Component | GO:0044306 | neuron projection terminus | 1.32E-05 | 8.67E-03 | 2.63E-04 | 1.86E-03 | 16 | 257 |
| GO: Cellular Component | GO:0034702 | ion channel complex | 1.47E-05 | 9.67E-03 | 2.84E-04 | 2.01E-03 | 18 | 319 |
| GO: Cellular Component | GO:0043679 | axon terminus | 1.83E-05 | 1.20E-02 | 3.44E-04 | 2.43E-03 | 15 | 235 |
| GO: Cellular Component | GO:0043197 | dendritic spine | 2.32E-05 | 1.52E-02 | 4.22E-04 | 2.98E-03 | 16 | 269 |
| GO: Cellular Component | GO:0044309 | neuron spine | 2.90E-05 | 1.90E-02 | 5.14E-04 | 3.63E-03 | 16 | 274 |
| GO: Cellular Component | GO:1902495 | transmembrane transporter complex | 3.71E-05 | 2.43E-02 | 6.34E-04 | 4.48E-03 | 18 | 342 |
| GO: Cellular Component | GO:0099056 | integral component of presynaptic membrane | 3.77E-05 | 2.47E-02 | 6.34E-04 | 4.48E-03 | 10 | 116 |
| GO: Cellular Component | GO:0032809 | neuronal cell body membrane | 4.07E-05 | 2.67E-02 | 6.67E-04 | 4.71E-03 | 6 | 37 |
| GO: Cellular Component | GO:0044298 | cell body membrane | 6.42E-05 | 4.21E-02 | 1.01E-03 | 7.12E-03 | 6 | 40 |
| GO: Cellular Component | GO:1990351 | transporter complex | 6.46E-05 | 4.24E-02 | 1.01E-03 | 7.12E-03 | 18 | 357 |

Hit in Query List
LGSN,GLUL,DPH6
GABBR1,GABRA5,GABRB1,GABRB3,GPR156
EPHB6,ERBB4,BCL11A,KLF4,EOMES,NRXN3,SALL1,NRXN1,NEGR1,UGT8,USH2A,PLP1,CXCL12,CNTN1,SDK2,ANK3,CHODL,PPFIA2,SH3GL3,LMO4,CASK,EFHC2,TENM4,DNM3,RSPO2,TMEM108,CTNNA2,GABRA5,GABRB1,GABRB3,DOK5,SOX11,SALL3,MYO5B,ATP8A2,SRRM4,RAP1GAP2,KIAA0319,XYLT1,RGMA,MAGI2,ANOS1,BLOC1S6,PTPRG,PTPRO,PTPRZ1,NTM,PEX5,DLG2,GNAO1,DMD,NT[illegible]
ERBB4,SYN3,TSPOAP1,SYT9,CDK14,KLF4,CHRM1,ADCY8,FAT1,ADRA2A,NRXN3,SALL1,NRXN1,GPC4,TRPV6,PLP1,CXCL12,BICC1,VSNL1,ANK3,PPFIA2,MYRIP,PEX5L,CASK,SLC6A2,CADPS,RSPO2,SORCS3,PRKCB,TMEM108,CADPS2,MCTP2,GABBR1,GABRA5,GABRB1,GABRB3,SOX11,GAD2,MYO5B,RSPO4,PSMD11,SST,BCL9,BCR,ENY2,MAGI2,NELL2,BLOC1S6,KCNA2,GPC3,KCND2,PTPRO,[illegible]
EPHB6,ERBB4,BCL11A,KLF4,EOMES,NRXN3,SALL1,NRXN1,NEGR1,FKTN,UGT8,USH2A,PLP1,CXCL12,CNTN1,SDK2,ANK3,CHODL,PPFIA2,SH3GL3,LMO4,CASK,EFHC2,TENM4,DNM3,RSPO2,TMEM108,CTNNA2,GABRA5,GABRB1,GABRB3,DOK5,SOX11,SALL3,MYO5B,ATP8A2,SRRM4,RAP1GAP2,KIAA0319,XYLT1,RGMA,MAGI2,ANOS1,BLOC1S6,PTPRG,PTPRO,PTPRZ1,NTM,PEX5,DLG2,ASTN2,[illegible]
EPHB6,ERBB4,BCL11A,KLF4,CHRM1,EOMES,NRXN3,SALL1,NRXN1,NEGR1,FKTN,UGT8,USH2A,PLP1,CXCL12,CNTN1,SDK2,ANK3,CHODL,PPFIA2,SH3GL3,LMO4,ADAM22,CASK,EFHC2,TENM4,DNM3,RSPO2,TMEM108,CTNNA2,GABRA5,GABRB1,GABRB3,DOK5,SOX11,SALL3,MYO5B,ATP8A2,SRRM4,RAP1GAP2,KIAA0319,XYLT1,RGMA,MAGI2,ANOS1,BLOC1S6,PTPRG,PTPRO,PTPRZ1,NTM,NAV2,PEX5[illegible]
ERBB4,SYN3,TSPOAP1,SYT9,CHRM1,ADCY8,ADRA2A,NRXN3,NRXN1,PLP1,ANK3,PPFIA2,CASK,SLC6A2,CADPS,SORCS3,PRKCB,TMEM108,CADPS2,MCTP2,GABBR1,GABRA5,GABRB1,GABRB3,GAD2,MYO5B,SST,BCR,BLOC1S6,KCNA2,KCND2,GLUL,DLG2,GNAO1,NOS1,CACNA1E,CACNB2,NTRK3,GRID2,IGSF11,GRIN2B,CCKBR,SHISA9
ERBB4,SYN3,TSPOAP1,SYT9,CHRM1,ADCY8,ADRA2A,NRXN3,NRXN1,PLP1,ANK3,PPFIA2,CASK,SLC6A2,CADPS,SORCS3,PRKCB,TMEM108,CADPS2,MCTP2,GABBR1,GABRA5,GABRB1,GABRB3,GAD2,MYO5B,SST,BCR,BLOC1S6,KCNA2,KCND2,GLUL,DLG2,GNAO1,NOS1,CACNA1E,CACNB2,NTRK3,GRID2,IGSF11,GRIN2B,CCKBR,SHISA9
ERBB4,SYN3,TSPOAP1,SYT9,CHRM1,ADCY8,ADRA2A,NRXN3,NRXN1,PLP1,ANK3,PPFIA2,CASK,SLC6A2,CADPS,SORCS3,PRKCB,TMEM108,CADPS2,MCTP2,GABBR1,GABRA5,GABRB1,GABRB3,GAD2,MYO5B,SST,BCR,BLOC1S6,KCNA2,KCND2,GLUL,DLG2,GNAO1,NOS1,CACNA1E,CACNB2,NTRK3,GRID2,IGSF11,GRIN2B,CCKBR,SHISA9
EPHB6,BCL11A,KLF4,NRXN3,NRXN1,NEGR1,UGT8,PLP1,CXCL12,CNTN1,ANK3,CHODL,PPFIA2,CASK,EFHC2,TENM4,DNM3,TMEM108,CTNNA2,GABRA5,GABRB1,GABRB3,DOK5,SALL3,MYO5B,ATP8A2,SRRM4,RAP1GAP2,KIAA0319,XYLT1,RGMA,MAGI2,ANOS1,BLOC1S6,PTPRG,PTPRO,PTPRZ1,NTM,DLG2,GNAO1,DMD,NTRK3,NR4A2,GRID2,DICER1,OPCML,CCK,GRIP1,THRB,ROBO1,RORB,[illegible]
ERBB4,SYN3,TSPOAP1,SYT9,CHRM1,ADCY8,ADRA2A,NRXN3,NRXN1,PLP1,ANK3,PPFIA2,CASK,SLC6A2,CADPS,SORCS3,PRKCB,TMEM108,CADPS2,MCTP2,GABBR1,GABRA5,GABRB1,GABRB3,GAD2,MYO5B,SST,BCR,BLOC1S6,KCNA2,KCND2,GLUL,DLG2,GNAO1,NOS1,CACNA1E,CACNB2,NTRK3,GRID2,IGSF11,GRIN2B,CCKBR,SHISA9
SYT9,ADCY8,ADRA2A,NRXN3,NRXN1,TRPV6,CXCL12,VSNL1,PPFIA2,MYRIP,PEX5L,CASK,STXBP6,CADPS,PRKCB,EXOC2,CADPS2,MCTP2,GABBR1,SOX11,MYO5B,BCR,ENY2,NELL2,KCNA2,SERGEF,GLUL,NOS1,NOS2,CACNA1E,CACNB2,NPY1R,GPR39,GRIN2B,CCK,CCKBR
SYN3,TSPOAP1,SYT9,ADCY8,ADRA2A,NRXN3,NRXN1,TRPV6,CXCL12,VSNL1,PPFIA2,MYRIP,PEX5L,CASK,CADPS,PRKCB,CADPS2,MCTP2,GABBR1,SOX11,GAD2,ENY2,NELL2,BLOC1S6,KCNA2,GLUL,NOS1,NOS2,CACNA1E,CACNB2,NPY1R,GPR39,CCKBR
SYN3,TSPOAP1,SYT9,ADRA2A,NRXN3,NRXN1,PPFIA2,CASK,SLC6A2,CADPS,SLC22A3,PRKCB,CADPS2,MCTP2,GAD2,BLOC1S6,GLUL,NOS1,CACNB2,SLC6A15
SYT9,ADCY8,ADRA2A,NRXN3,NRXN1,TRPV6,CXCL12,VSNL1,PPFIA2,MYRIP,PEX5L,CASK,STXBP6,CADPS,PRKCB,EXOC2,CADPS2,MCTP2,GABBR1,SOX11,MYO5B,BCR,ENY2,NELL2,KCNA2,SERGEF,GLUL,NOS1,NOS2,CACNA1E,CACNB2,NPY1R,GPR39,GRIN2B,CCK,CCKBR
SYN3,MAOA,TSPOAP1,SYT9,ADRA2A,NRXN3,NRXN1,PPFIA2,CASK,SLC6A2,CADPS,SLC22A3,PRKCB,CADPS2,MCTP2,GAD2,BLOC1S6,GLUL,NOS1,CACNB2
EPHB6,MKKS,ERBB4,CENPF,EOMES,SALL1,NRXN1,ACAN,UGP2,UGT8,ALDH1A3,PLP1,CXCL12,CNTN1,SH3GL3,LMO4,ADAM22,ZIC3,TENM4,TMEM108,CTNNA2,JRKL,GABRA5,GABRB1,GABRB3,SOX11,SALL3,MYO1D,BCR,CA10,PTPRG,PTPRZ1,ARNT2,NAV2,PEX5,GNAO1,DMD,NTRK3,NR4A2,GRID2,GRIN2B,DICER1,CHST8,ROBO1,PBX1,DCLK1
BCL11A,KLF4,NRXN1,NEGR1,CXCL12,CNTN1,CHODL,PPFIA2,CARMIL1,CASK,EFHC2,DNM3,CTNNA2,MYO5B,BCAS3,ATP8A2,TBC1D5,RAP1GAP2,KIAA0319,XYLT1,RGMA,MAGI2,PTPRG,PTPRO,PTPRZ1,NTM,DMD,NTRK3,GRID2,GRIN2B,GRIP1,ROBO1,EPHA3,PLCE1
ERBB4,NRXN3,NRXN1,NEGR1,GPC4,SDK2,ANK3,PPFIBP2,PPFIA2,CNKSR2,CASK,DNM3,TMEM108,CTNNA2,PDZRN3,GABRB3,MYO5B,MAGI2,PTPRO,CACNB2,NTRK3,GRID2,GRIN2B,GPC6,PRMT3,EPHA3
BCL11A,KLF4,NEGR1,CXCL12,CNTN1,CHODL,PPFIA2,CASK,EFHC2,DNM3,CTNNA2,MYO5B,ATP8A2,RAP1GAP2,KIAA0319,XYLT1,RGMA,MAGI2,PTPRG,PTPRO,PTPRZ1,NTM,DMD,NTRK3,GRID2,ROBO1,EPHA3
KCNH8,TAFA4,CHRM1,TRDN,NRXN1,NALCN,ANK3,PEX5L,TMEM108,GABBR1,KCNH5,GABRA5,GABRB1,GABRB3,KCNA2,KCND2,DMD,PID1,CACNA1E,CACNB2,GPR39,NTRK3,GRID2,IGSF11,GRIN2B,CCK
EPHB6,BCL11A,KLF4,NRXN3,NRXN1,NEGR1,UGT8,PLP1,CXCL12,CNTN1,ANK3,CHODL,PPFIA2,CASK,EFHC2,DNM3,TMEM108,CTNNA2,DOK5,MYO5B,ATP8A2,RAP1GAP2,KIAA0319,XYLT1,RGMA,MAGI2,ANOS1,BLOC1S6,PTPRG,PTPRO,PTPRZ1,NTM,GNAO1,DMD,NTRK3,NR4A2,GRID2,DICER1,CCK,GRIP1,ROBO1,PRMT3,DCLK1,EPHA3
EPHB6,MKKS,BCL11A,KLF4,ONECUT1,TUB,NRXN3,NRXN1,NEGR1,UGT8,PLP1,CXCL12,CNTN1,ANK3,CHODL,OFD1,PPFIA2,CARMIL1,RTTN,CASK,EFHC2,DYNLL2,CEP70,DNM3,TMEM108,CEP152,CTNNA2,DOK5,MYO5B,BCAS3,ATP8A2,TBC1D5,STK26,RAP1GAP2,KIAA0319,XYLT1,RGMA,MAGI2,ANOS1,BLOC1S6,PTPRG,PTPRO,PTPRZ1,NTM,EXOG,GNAO1,DMD,NTRK3,NR4A2,GRID2,GRIN2[illegible]
LRAT,ERBB4,OPN3,SYT9,CHRM1,TRDN,TUB,ADCY8,ADRA2A,PLP1,CXCL12,CNTN1,VSNL1,ANK3,MYRIP,PEX5L,CASK,CADPS,DNM3,EXOC2,CADPS2,GABBR1,SOX11,MYO5B,BCAS3,ATP8A2,TBC1D5,BCR,MAGI2,NELL2,GPC3,GLUL,DMD,NOS1,CACNB2,CALCR,GPR39,GRIN2B,CCK,CCKBR,CEMIP
BCL11A,KLF4,NRXN1,NEGR1,CXCL12,CNTN1,CHODL,PPFIA2,CARMIL1,CASK,EFHC2,DNM3,CTNNA2,MYO5B,BCAS3,ATP8A2,TBC1D5,RAP1GAP2,KIAA0319,XYLT1,RGMA,MAGI2,PTPRG,PTPRO,PTPRZ1,NTM,DMD,NTRK3,GRID2,GRIN2B,ROBO1,EPHA3,PLCE1
MKKS,ADCY8,NRXN3,NRXN1,NEGR1,ALDH1A3,UNC79,CXCL12,ADAM22,SORCS3,DMRTA1,GABRA5,GABRB3,ATP8A2,KCNA2,KCND2,PTPRZ1,NAV2,GNAO1,NOS1,CACNA1E,NPY1R,GPR39,PAK5,NTS,NR4A2,GRIN2B,GRPR,CCK,CCKBR,THRB,OTOG
EPHB6,MKKS,BCL11A,KLF4,ONECUT1,TUB,NRXN3,NRXN1,NEGR1,UGT8,PLP1,CXCL12,CNTN1,ANK3,CHODL,OFD1,PPFIA2,CARMIL1,RTTN,CASK,EFHC2,DYNLL2,CEP70,DNM3,TMEM108,CEP152,CTNNA2,DOK5,MYO5B,BCAS3,ATP8A2,TBC1D5,STK26,RAP1GAP2,KIAA0319,XYLT1,RGMA,MAGI2,ANOS1,BLOC1S6,PTPRG,PTPRO,PTPRZ1,NTM,EXOG,GNAO1,DMD,NTRK3,NR4A2,GRID2,GRIN2B,RFX2,[illegible]
SYN3,SYT9,ADRA2A,NRXN3,NRXN1,PPFIA2,SH3GL3,CASK,CADPS,DNM3,PRKCB,CADPS2,MYO5B,MAGI2,BLOC1S6,CACNB2,GRIP1
SYT9,ADCY8,CXCL12,VSNL1,MYRIP,PEX5L,CASK,CADPS,EXOC2,CADPS2,GABBR1,SOX11,MYO5B,NELL2,GLUL,CACNB2,GPR39,GRIN2B,CCK,CCKBR
SYT9,ADCY8,ADRA2A,NRXN1,VSNL1,MYRIP,PEX5L,CASK,GABBR1,SOX11,ENY2,NELL2,GLUL,NOS1,NOS2,CACNA1E,NPY1R,GPR39,CCKBR
SYN3,TSPOAP1,SYT9,ADRA2A,NRXN3,NRXN1,PPFIA2,CASK,CADPS,PRKCB,CADPS2,MCTP2,GAD2,BLOC1S6,CACNB2
SYN3,TSPOAP1,SYT9,ADRA2A,NRXN3,NRXN1,PPFIA2,CASK,CADPS,PRKCB,CADPS2,MCTP2,GAD2,BLOC1S6,CACNB2
ERBB4,SALL1,SALL3,DICER1,ROBO1
ERBB4,SYN3,TSPOAP1,SYT9,BCL11A,CHRM1,ADCY8,PPM1H,ADRA2A,NRXN3,NRXN1,ACAN,GPC4,USH2A,PLP1,CNTN1,SDK2,ANK3,PPFIBP2,PPFIA2,SH3GL3,MYRIP,CNKSR2,ADAM22,CASK,SLC6A2,CADPS,SLC22A3,DYNLL2,DNM3,SORCS3,PRKCB,TMEM108,SLC2A13,CTNNA2,CADPS2,MCTP2,PDZRN3,GABBR1,GABRA5,GABRB1,GABRB3,GAD2,ARHGAP32,MYO5B,SST,BCR,MAGI2,BLOC1S6,[illegible]
ERBB4,CHRM1,ADCY8,ADRA2A,NRXN3,NRXN1,GPC4,CNTN1,ANK3,CNKSR2,ADAM22,CASK,SLC6A2,DNM3,SORCS3,CTNNA2,CADPS2,GABBR1,GABRA5,GABRB1,GABRB3,GAD2,MAGI2,KCNA2,KCND2,PTPRO,PTPRZ1,DLG2,DMD,CACNA1E,NTRK3,GRID2,GRIN2B,GRIP1,SHISA9,GPR156
EPHB6,MKKS,MAP9,TSPOAP1,OPN3,SYT9,KCNB2,CHRM1,ADCY8,ADRA2A,NRXN1,NEGR1,ACAN,USH2A,ANK3,PPFIA2,MYRIP,CNKSR2,PEX5L,ADAM22,CASK,SLC6A2,TENM4,DNM3,PRKCB,TMEM108,SLC2A13,CTNNA2,UNC80,GABBR1,GABRA5,GABRB1,GABRB3,GAD2,ARHGAP32,MYO1D,MYO5B,ATP8A2,RAP1GAP2,GC,BCR,RGMA,MAGI2,NELL2,BLOC1S6,KCNA2,KCND2,PTPRO,PTPRZ1,NTM,GLUL,DL[illegible]
ERBB4,SYN3,TSPOAP1,SYT9,CHRM1,ADCY8,ADRA2A,NRXN3,NRXN1,GPC4,USH2A,CNTN1,PPFIBP2,PPFIA2,SH3GL3,CASK,SLC6A2,CADPS,SLC22A3,DNM3,PRKCB,SLC2A13,CTNNA2,CADPS2,MCTP2,GABBR1,GABRA5,GABRB3,GAD2,BLOC1S6,KCNA2,GLUL,DLG2,NOS1,CACNA1E,CACNB2,NPY1R,NTS,GRIN2B,CCK,EPHA3
ERBB4,CHRM1,ADRA2A,CNTN1,ANK3,CNKSR2,ADAM22,DNM3,SORCS3,CTNNA2,CADPS2,GABBR1,GABRA5,GABRB1,GABRB3,MAGI2,KCND2,PTPRO,PTPRZ1,DLG2,DMD,CACNA1E,NTRK3,GRID2,GRIN2B,GRIP1,SHISA9,GPR156
ERBB4,SYN3,SYT9,BCL11A,CHRM1,ADCY8,ADRA2A,CNTN1,ANK3,PPFIA2,SH3GL3,CNKSR2,ADAM22,CASK,DYNLL2,DNM3,SORCS3,TMEM108,CTNNA2,CADPS2,GABBR1,GABRA5,GABRB1,GABRB3,ARHGAP32,MYO5B,BCR,MAGI2,KCND2,PTPRO,PTPRZ1,DLG2,DMD,NOS1,CACNA1E,GOPC,NTRK3,GRID2,IGSF11,GRIN2B,GRIP1,SHISA9,GPR156,DCLK1
ERBB4,SYN3,TSPOAP1,CHRM1,ADCY8,PPM1H,ADRA2A,NRXN1,ACAN,GPC4,PPFIA2,SH3GL3,CNKSR2,ADAM22,CADPS,DYNLL2,DNM3,SORCS3,CADPS2,GABBR1,MYO5B,BCR,MAGI2,KCND2,PTPRO,PTPRZ1,DLG2,NOS1,NTRK3,GRID2,GRIN2B,GPC6,GRIP1,SHISA9
ERBB4,SYN3,BCL11A,CHRM1,ADCY8,ADRA2A,SH3GL3,CNKSR2,ADAM22,CASK,DYNLL2,DNM3,SORCS3,TMEM108,CTNNA2,GABRA5,GABRB3,ARHGAP32,BCR,MAGI2,KCND2,PTPRO,PTPRZ1,DLG2,NOS1,GOPC,GRID2,IGSF11,GRIN2B,GRIP1,SHISA9,DCLK1
ERBB4,SYN3,BCL11A,CHRM1,ADCY8,ADRA2A,SH3GL3,CNKSR2,ADAM22,CASK,DYNLL2,DNM3,SORCS3,TMEM108,CTNNA2,ARHGAP32,BCR,MAGI2,KCND2,PTPRO,PTPRZ1,DLG2,NOS1,GOPC,GRID2,IGSF11,GRIN2B,GRIP1,SHISA9,DCLK1
ERBB4,SYN3,BCL11A,CHRM1,ADCY8,ADRA2A,SH3GL3,CNKSR2,ADAM22,CASK,DYNLL2,DNM3,SORCS3,TMEM108,CTNNA2,ARHGAP32,BCR,MAGI2,KCND2,PTPRO,PTPRZ1,DLG2,NOS1,GOPC,GRID2,IGSF11,GRIN2B,GRIP1,SHISA9,DCLK1
ERBB4,SYN3,SYT9,BCL11A,CHRM1,ADCY8,ADRA2A,SH3GL3,CNKSR2,ADAM22,CASK,DYNLL2,DNM3,SORCS3,TMEM108,CTNNA2,ARHGAP32,BCR,MAGI2,KCND2,PTPRO,PTPRZ1,DLG2,NOS1,GOPC,GRID2,IGSF11,GRIN2B,GRIP1,SHISA9,DCLK1
ERBB4,CHRM1,ADCY8,ADRA2A,NRXN1,GPC4,CNTN1,ADAM22,SORCS3,GABBR1,GABRA5,GABRB3,KCNA2,KCND2,PTPRO,PTPRZ1,DLG2,CACNA1E,NTRK3,GRID2,GRIN2B,SHISA9
SYT9,KCNB2,CHRM1,ADCY8,ADRA2A,NRXN1,NEGR1,ACAN,USH2A,ANK3,PPFIA2,CNKSR2,PEX5L,CASK,SLC6A2,SLC22A3,DNM3,TMEM108,CTNNA2,GABBR1,GABRA5,GABRB1,ARHGAP32,MYO1D,MYO5B,SST,BCR,MAGI2,NELL2,KCNA2,KCND2,PTPRO,PTPRZ1,GLUL,DLG2,ASTN2,GNAO1,GNAZ,NOS1,CACNA1E,CALCR,GOPC,NTS,GRID2,GRIN2B,DICER1,CCK,GRIP1,SHISA9,ROBO1,HCFC1
ERBB4,CHRM1,ADRA2A,CNTN1,ADAM22,SORCS3,GABBR1,GABRA5,GABRB3,KCND2,PTPRO,PTPRZ1,DLG2,CACNA1E,NTRK3,GRID2,GRIN2B,SHISA9
ERBB4,ADRA2A,NRXN3,NRXN1,ACAN,GABBR1,GABRA5,GABRB1,GABRB3,MAGI2,KCND2,PTPRO,CACNA1E,EPHA3
ERBB4,CHRM1,ADRA2A,ADAM22,SORCS3,GABRA5,GABRB3,KCND2,PTPRO,PTPRZ1,DLG2,GRID2,GRIN2B,SHISA9
MAP9,TSPOAP1,SYT9,CHRM1,ADCY8,ADRA2A,NRXN1,USH2A,ANK3,PPFIA2,ADAM22,DNM3,PRKCB,TMEM108,SLC2A13,CTNNA2,UNC80,GABBR1,GABRB3,GAD2,MYO1D,GC,BCR,BLOC1S6,KCNA2,PTPRO,PTPRZ1,NTM,GLUL,DLG2,NOS1,NPY1R,CALCR,NTRK3,NTS,GRIN2B,DICER1,CCK,ROBO1,HCFC1
ERBB4,CHRM1,ADCY8,ADRA2A,NRXN1,ADAM22,SORCS3,GABBR1,GABRA5,GABRB3,KCNA2,KCND2,PTPRO,PTPRZ1,CACNA1E,NTRK3,GRID2,GRIN2B,SHISA9
ERBB4,CHRM1,ADCY8,ADRA2A,NRXN3,NRXN1,GPC4,CNTN1,CASK,SLC6A2,CTNNA2,CADPS2,GABBR1,GABRA5,GAD2,KCNA2,CACNA1E,GRIN2B
ERBB4,CHRM1,ADRA2A,ADAM22,SORCS3,GABRA5,GABRB3,KCND2,PTPRO,PTPRZ1,GRID2,GRIN2B,SHISA9
ERBB4,CHRM1,ADRA2A,ADAM22,SORCS3,GABBR1,GABRA5,GABRB3,KCND2,PTPRO,PTPRZ1,CACNA1E,NTRK3,GRID2,GRIN2B,SHISA9
ERBB4,CHRM1,ADRA2A,CNKSR2,ADAM22,SORCS3,GABRA5,GABRB3,KCND2,PTPRO,PTPRZ1,DLG2,GRID2,GRIN2B,SHISA9
ERBB4,CHRM1,ADRA2A,ADAM22,SORCS3,PTPRO,PTPRZ1,DLG2,GRID2,GRIN2B,SHISA9
SYT9,KCNB2,CHRM1,ADCY8,NEGR1,ANK3,PPFIA2,PEX5L,CASK,DNM3,TMEM108,CTNNA2,GABBR1,GABRA5,GABRB1,ARHGAP32,MYO1D,MYO5B,BCR,MAGI2,NELL2,KCNA2,KCND2,PTPRO,PTPRZ1,DLG2,GNAO1,GNAZ,NOS1,GOPC,GRID2,GRIN2B,DICER1,CCK,GRIP1,SHISA9,HCFC1
SYT9,KCNB2,CHRM1,ADCY8,NEGR1,ANK3,PPFIA2,PEX5L,CASK,DNM3,TMEM108,CTNNA2,GABBR1,GABRA5,GABRB1,ARHGAP32,MYO1D,MYO5B,BCR,MAGI2,NELL2,KCNA2,KCND2,PTPRO,PTPRZ1,DLG2,GNAO1,GNAZ,NOS1,GOPC,GRID2,GRIN2B,DICER1,CCK,GRIP1,SHISA9,HCFC1
ERBB4,CHRM1,ADRA2A,ADAM22,SORCS3,PTPRO,PTPRZ1,GRID2,GRIN2B,SHISA9
ERBB4,CHRM1,ADRA2A,CNKSR2,ADAM22,SORCS3,PTPRO,PTPRZ1,DLG2,GRID2,GRIN2B,SHISA9
ERBB4,CHRM1,ADCY8,ADRA2A,NRXN1,GPC4,CNTN1,GABBR1,GABRA5,KCNA2,CACNA1E,GRIN2B
KCNB2,ADCY8,ADRA2A,NRXN1,NEGR1,ACAN,USH2A,CNKSR2,CASK,SLC6A2,SLC22A3,SLC2A13,GABBR1,GABRA5,MYO1D,MYO5B,SST,NELL2,KCNA2,KCND2,PTPRZ1,GLUL,DLG2,ASTN2,GNAO1,GNAZ,CACNA1E,CALCR,NTS,GRIN2B,CCK,GRIP1,ROBO1,HCFC1
KCNB2,ADCY8,ADRA2A,NRXN1,NEGR1,ACAN,USH2A,CNKSR2,CASK,SLC6A2,SLC22A3,GABBR1,GABRA5,MYO1D,MYO5B,SST,NELL2,KCNA2,KCND2,PTPRZ1,GLUL,DLG2,ASTN2,CACNA1E,CALCR,NTS,GRIN2B,CCK,GRIP1,ROBO1,HCFC1
EPHB6,ERBB4,KCNH8,OPN3,SLC24A3,KCNB2,CHRM1,ADCY8,FAT1,ADRA2A,NRXN3,NRXN1,GPC4,TRPV6,PLP1,CNTN1,STX8,ADAM22,SLC6A2,TENM4,SLC22A3,SORCS3,SLC2A13,GABBR1,KCNH5,GABRA5,GABRB1,GABRB3,SLCO3A1,GPRC5C,ST14,KCNA2,GPC3,PTPRG,KCND2,PTPRO,PTPRZ1,DLG2,CACNA1E,CACNB2,NPY1R,CALCR,GPR39,NTRK3,SLC6A15,GRID2,GRIN2B,GPC6,GRPR,SHISA9,SL[illegible]
EPHB6,ERBB4,KCNH8,OPN3,SLC24A3,KCNB2,CHRM1,ADCY8,FAT1,ADRA2A,NRXN3,NRXN1,TRPV6,PLP1,STX8,ADAM22,SLC6A2,TENM4,SLC22A3,SORCS3,SLC2A13,GABBR1,KCNH5,GABRA5,GABRB1,GABRB3,SLCO3A1,GPRC5C,ST14,KCNA2,GPC3,PTPRG,KCND2,PTPRO,PTPRZ1,DLG2,CACNA1E,CACNB2,NPY1R,CALCR,GPR39,NTRK3,SLC6A15,GRID2,GRIN2B,GPC6,GRPR,SHISA9,SLCO5A1,GUCY2[illegible]
TSPOAP1,SYT9,CHRM1,ADCY8,ADRA2A,USH2A,PRKCB,CTNNA2,GABRB3,KCNA2,GLUL,DMD,NOS1,NTS,GRIN2B,CCK
KCNB2,TRDN,TRPV6,PEX5L,UNC80,CACNA2D3,GABRA5,GABRB1,GABRB3,KCNA2,KCND2,DLG2,NOS1,CACNA1E,CACNB2,GRID2,GRIN2B,SHISA9
TSPOAP1,SYT9,CHRM1,ADCY8,ADRA2A,USH2A,PRKCB,CTNNA2,GABRB3,KCNA2,GLUL,NOS1,NTS,GRIN2B,CCK
SYT9,ADCY8,PPFIA2,DNM3,CTNNA2,GABBR1,ARHGAP32,MYO5B,BCR,KCND2,PTPRO,PTPRZ1,NOS1,GRID2,GRIN2B,SHISA9
SYT9,ADCY8,PPFIA2,DNM3,CTNNA2,GABBR1,ARHGAP32,MYO5B,BCR,KCND2,PTPRO,PTPRZ1,NOS1,GRID2,GRIN2B,SHISA9
KCNB2,TRDN,TRPV6,PEX5L,UNC80,CACNA2D3,GABRA5,GABRB1,GABRB3,KCNA2,KCND2,DLG2,NOS1,CACNA1E,CACNB2,GRID2,GRIN2B,SHISA9
ERBB4,CHRM1,ADCY8,ADRA2A,NRXN1,GABBR1,GABRA5,KCNA2,CACNA1E,GRIN2B
KCNB2,ADCY8,SLC6A2,GABRA5,KCNA2,KCND2
KCNB2,ADCY8,SLC6A2,GABRA5,KCNA2,KCND2
KCNB2,TRDN,TRPV6,PEX5L,UNC80,CACNA2D3,GABRA5,GABRB1,GABRB3,KCNA2,KCND2,DLG2,NOS1,CACNA1E,CACNB2,GRID2,GRIN2B,SHISA9
